## Extended Discussion for "SyntheMol-RL: a flexible reinforcement learning framework for designing novel and synthesizable antibiotics"

We compared SyntheMol-RL to GFlowNet^1^, a state-of-the-art generative model for drug design that employs an RL-like generative model with sampling for diverse molecule generation, which inspired some of the elements of SyntheMol-RL. We trained a multi-objective GFlowNet^2^ to design molecules using three reward functions: (1) our antibacterial activity Chemprop-RDKit model, (2) our aqueous solubility Chemprop-RDKit model, and (3) the synthetic accessibility score SAScore^3^. The former two rewards are the same models used to reward SyntheMol-RL, while the latter objective accounts for the fact that GFlowNet models do not inherently generate easily synthesizable molecules.

The GFlowNet model outperforms SyntheMol-RL in terms of predicted antibacterial activity and aqueous solubility, even when filtered to only include generated molecules with SAScore ≤ 4 (i.e., predicted easy synthesizability) (Extended Data Fig. 3a-b). However, the molecules designed by GFlowNet are bulky (Extended Data Fig. 3c) with complex multi-ring structures and would likely suffer from poor whole cell activity and synthesizability, despite the high predicted SAScores (Supplementary Data 12, Extended Data Fig. 3d-f). Indeed, consultations with medicinal chemists at Enamine suggested that synthesis of any of the GFlowNet-generated compounds would be complex, resulting in a cost at least 40x that of the SyntheMol-RL compounds, with long synthesis times. Similarly, WuXi chemists offered synthesis for 8 / 14 (57%) of the compounds but with costs 30-50x higher and synthesis times 1.5-2x longer than the SyntheMol-RL compounds. This result highlights the value of SyntheMol-RL; while other models may generate molecules with better properties *in silico*, the lack of readily available synthesis of those molecules hinders their real-world value for drug discovery purposes. SyntheMol-RL, in contrast, can translate promising *in silico* compound designs to actual, synthesized molecules for rapid and inexpensive laboratory validation.

**Extended Discussion References**

1.	Bengio, E., Jain, M., Korablyov, M., Precup, D. & Bengio, Y. Flow Network based Generative Models for Non-Iterative Diverse Candidate Generation. in *Advances in Neural Information Processing Systems* vol. 34 27381–27394 (Curran Associates, Inc., 2021).

2.	Jain, M. *et al.* Multi-Objective GFlowNets. in *Proceedings of the 40th International Conference on Machine Learning* (eds. Krause, A. et al.) vol. 202 14631–14653 (PMLR, 2023).

3.	Ertl, P. & Schuffenhauer, A. Estimation of synthetic accessibility score of drug-like molecules based on molecular complexity and fragment contributions. *J. Cheminformatics* **1**, 8 (2009).
