## Supplementary Data 13 for "SyntheMol-RL: a flexible reinforcement learning framework for designing novel and synthesizable antibiotics": Z365789618.PDF

# BE177056$3

MaxPeak: 100.00%
Ret_Time: 0.892 min

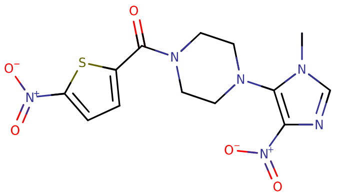

**Mol Wt** 366.35
**Exact Mass** 366.06

| # | Time | Area% |
|---|---|---|
| 1 | 0.892 | 100.00 |

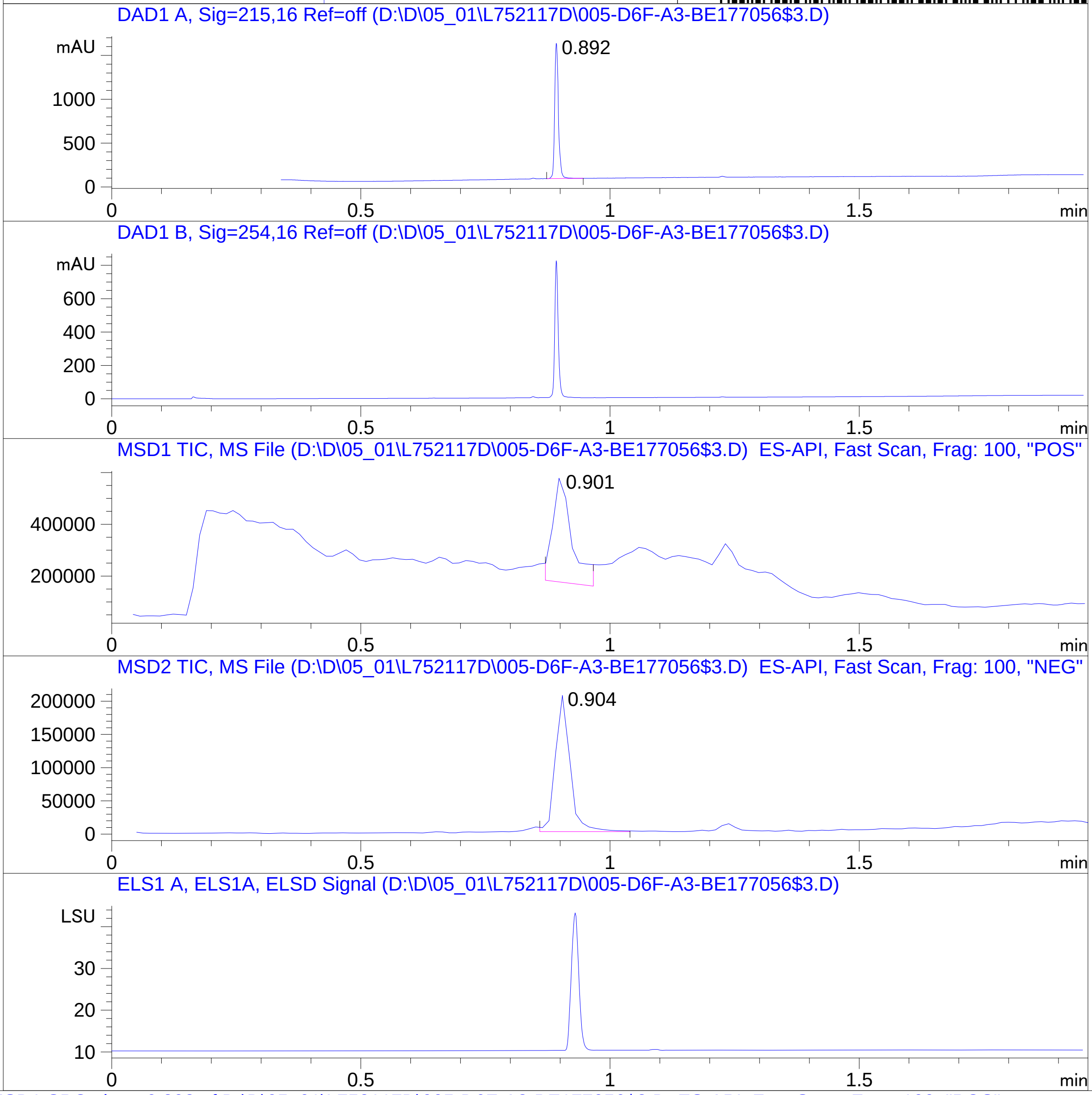


RT 0.901

RT 0.904

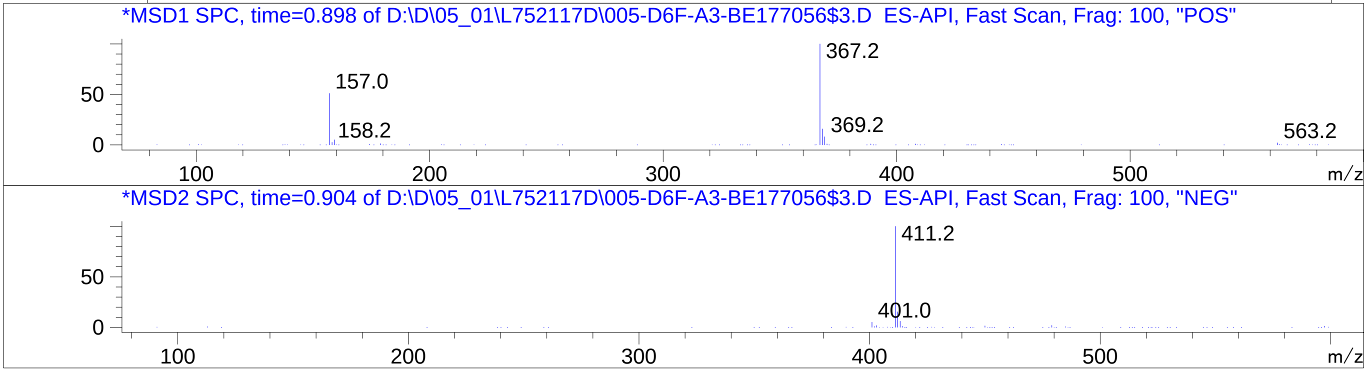


Inj.Date 5/1/2024 N 35
C:\Users\Public\Documents\ChemStation\1\Data\05_01\L752117D\SUPOR_30.M
