## Supplementary Data 13 for "SyntheMol-RL: a flexible reinforcement learning framework for designing novel and synthesizable antibiotics": Z1329167212.PDF

MaxPeak: 100.00%
Ret_Time: 1.340 min

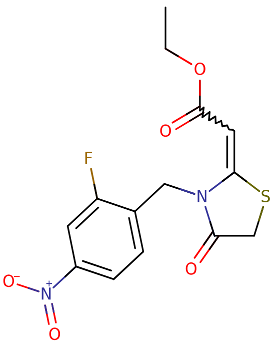

# BD772647$2

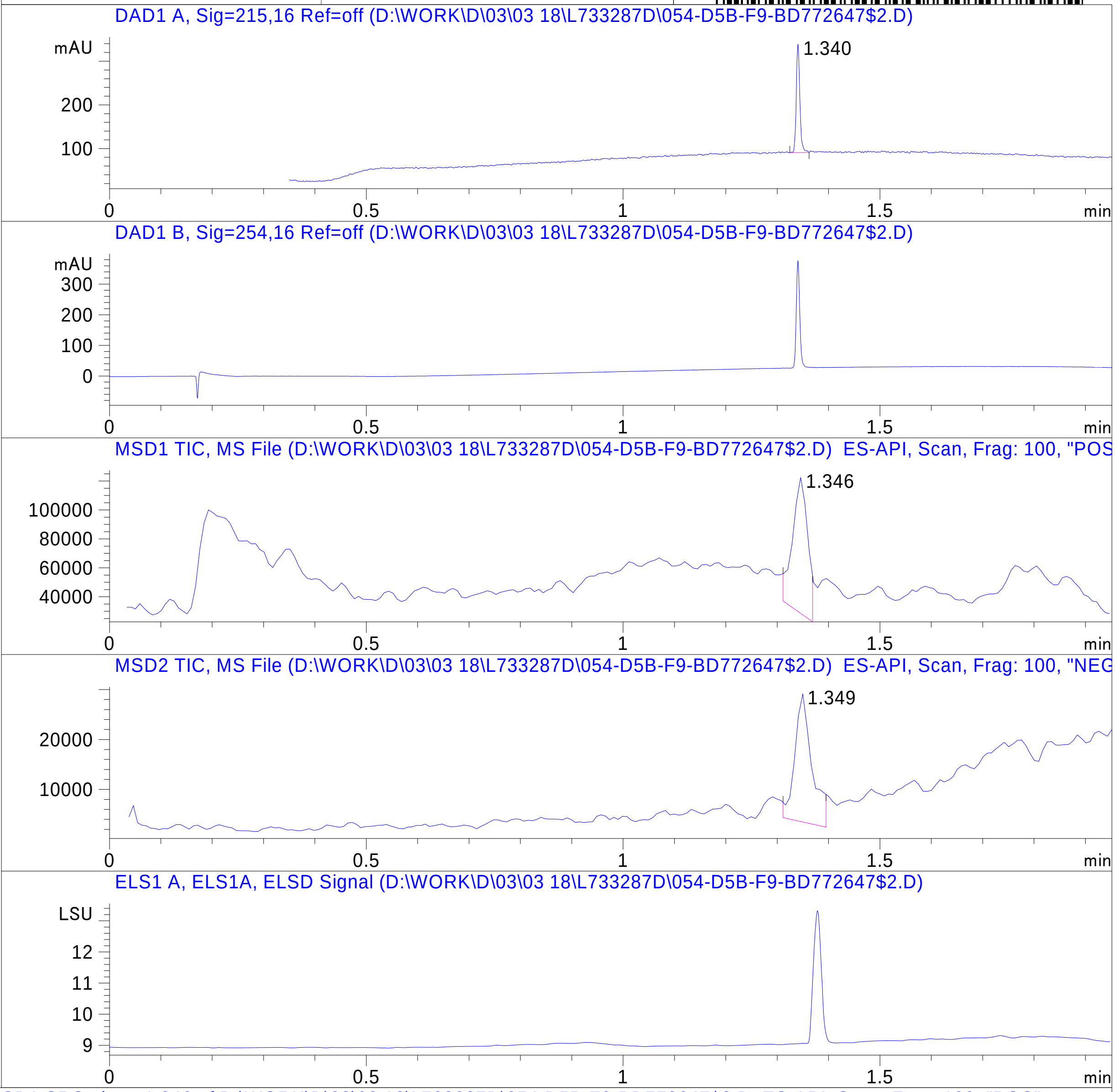


**Mol Wt** 340.33

**Exact Mass** 340.05

| # | Time | Area% |
|---|---|---|
| 1 | 1.340 | 100.00 |

RT 1.346

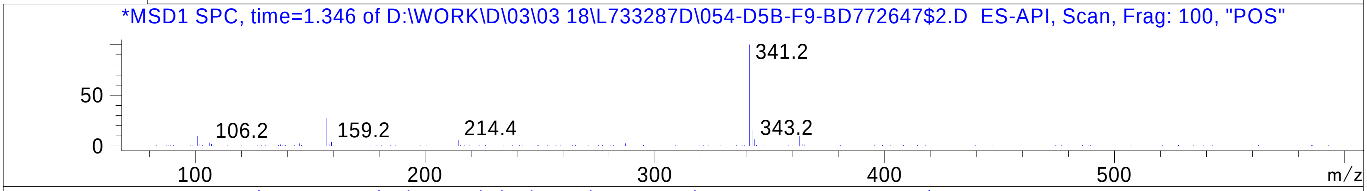


RT 1.349

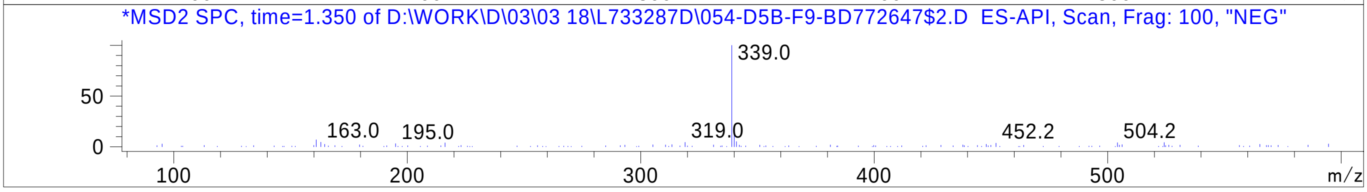


Inj.Date 3/18/2024 K



C:\Users\Public\Documents\ChemStation\1\Data\03_18\L733287D\SUPOR_30.M
