## Supplementary Data 13 for "SyntheMol-RL: a flexible reinforcement learning framework for designing novel and synthesizable antibiotics": Z1468376758.PDF

MaxPeak: 100.00%
Ret_Time: 0.952 min

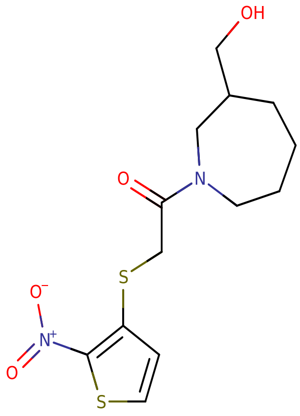


**Mol Wt** 330.42

**Exact Mass** 330.08

| # | Time | Area% |
|---|---|---|
| 1 | 0.952 | 100.00 |

# BE177024$1

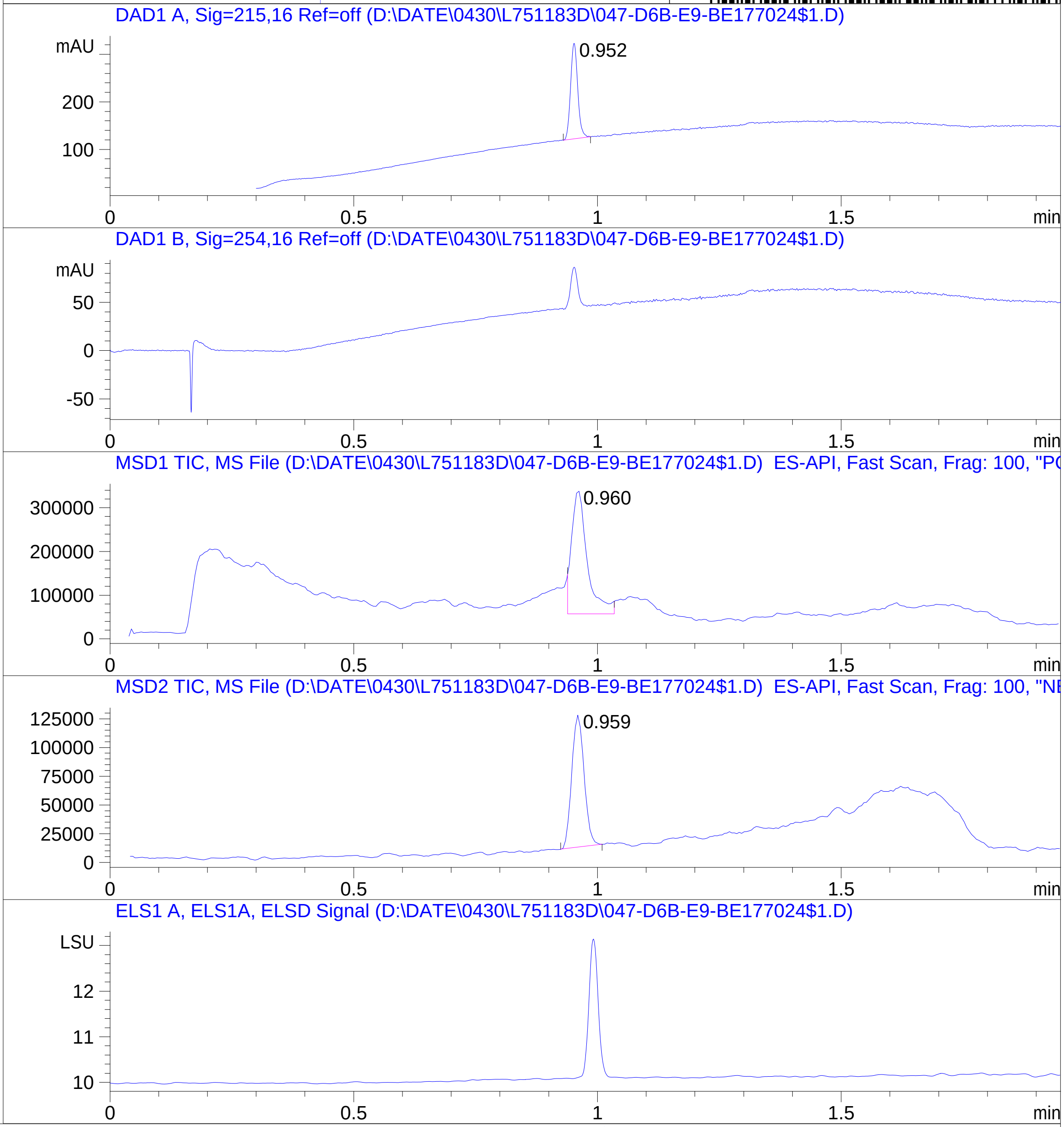


RT 0.960

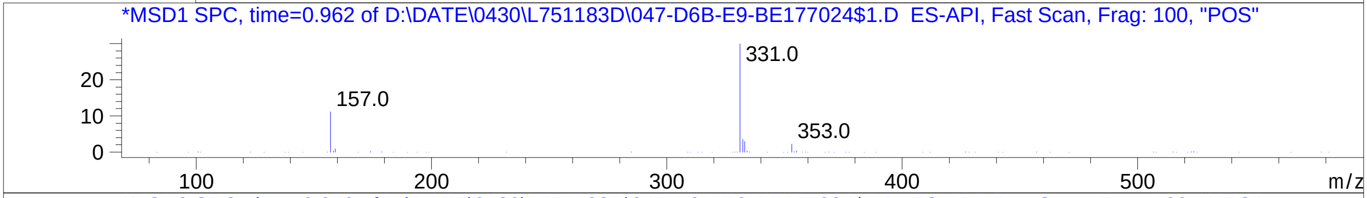


RT 0.959

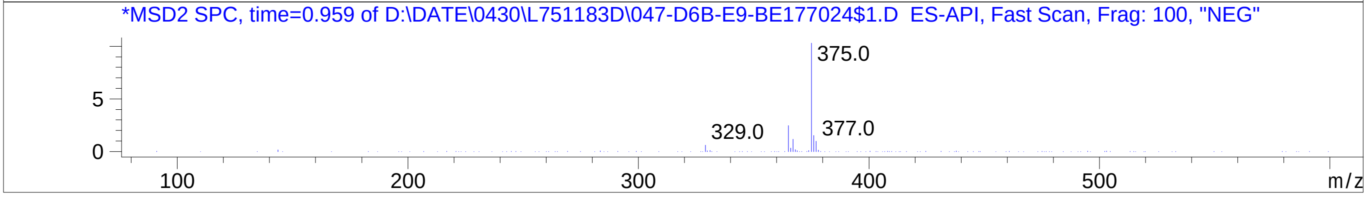


Inj.Date 4/30/2024 0
C:\Users\Public\Documents\ChemStation\1\Data\04_29\L751183D\SUPOR_30.M
