## Supplementary Data 13 for "SyntheMol-RL: a flexible reinforcement learning framework for designing novel and synthesizable antibiotics": Z1720541944.PDF

MaxPeak: 92.40%
Ret_Time: 1.232 min

# BD252444$1

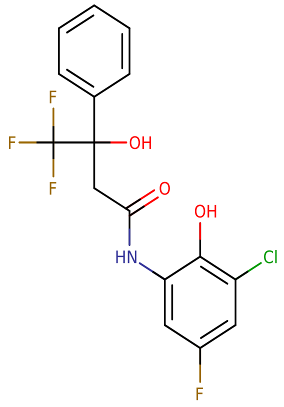


**Mol Wt** 377.72

**Exact Mass** 377.06

| # | Time | Area% |
|---|---|---|
| 1 | 0.937 | 2.26 |
| 2 | 0.979 | 1.14 |
| 3 | 1.080 | 1.30 |
| 4 | 1.115 | 2.91 |
| 5 | 1.232 | 92.40 |

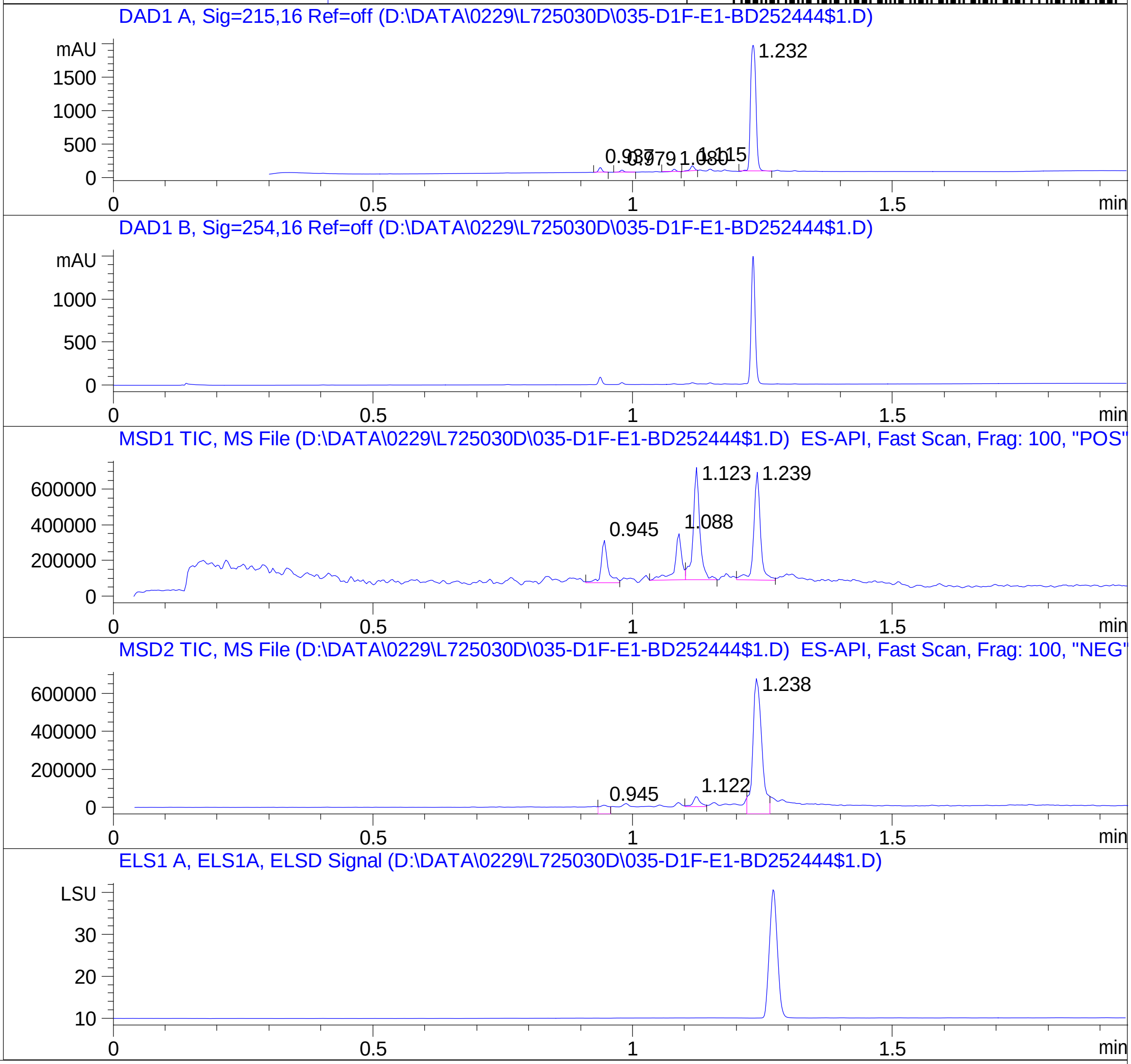


RT 0.945

RT 1.088

RT 1.123

RT 1.239

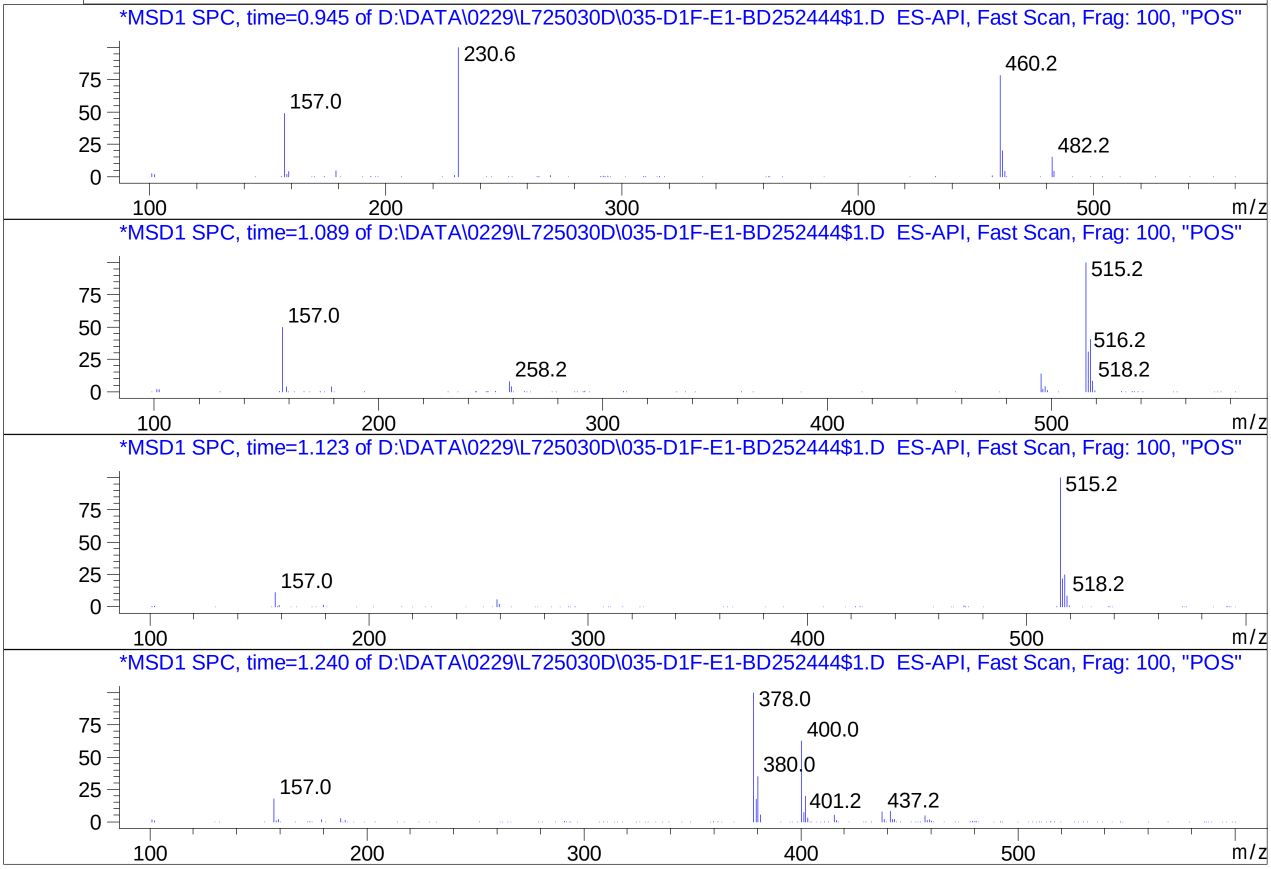


Inj.Date 2/29/2024 LB

C:\Users\Public\Documents\ChemStation\1\Data\02_29\L725030D\SUPOR_96.M

RT 0.945

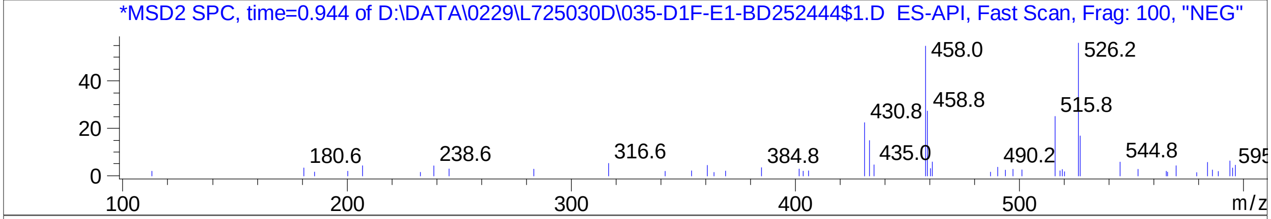


RT 1.122

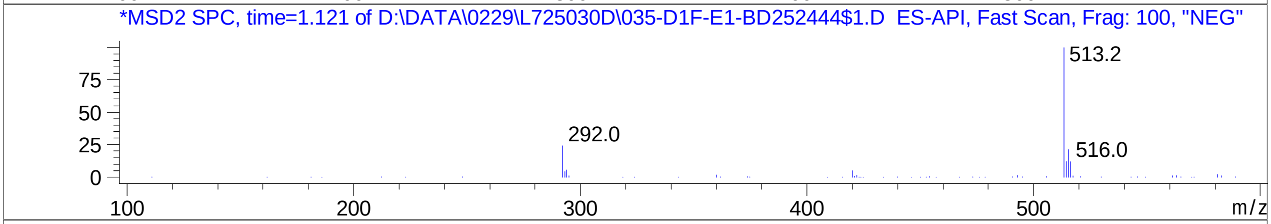


RT 1.238

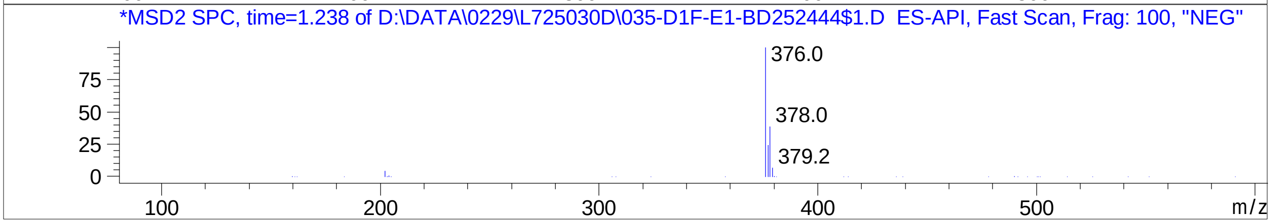


Inj.Date 2/29/2024 LB

C:\Users\Public\Documents\ChemStation\1\Data\02_29\L725030D\SUPOR_96.M
