## Supplementary Data 13 for "SyntheMol-RL: a flexible reinforcement learning framework for designing novel and synthesizable antibiotics": Z1985659907.PDF

# BE177022$2

MaxPeak: 100.00%
Ret_Time: 1.245 min

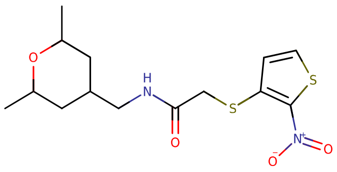

**Mol Wt** 344.45

**Exact Mass** 344.1

| # | Time | Area% |
|---|---|---|
| 1 | 1.245 | 100.00 |

DAD1 A, Sig=215,16 Ref=off (D:\D\04_30\L751360D\SAMPLE000033.D)

DAD1 B, Sig=254,16 Ref=off (D:\D\04_30\L751360D\SAMPLE000033.D)

MSD1 TIC, MS File (D:\D\04_30\L751360D\SAMPLE000033.D) ES-API, Scan, Frag: 100, "POS"

MSD2 TIC, MS File (D:\D\04_30\L751360D\SAMPLE000033.D) ES-API, Scan, Frag: 100, "NEG"

ADC1 A, ADC1 (D:\D\04_30\L751360D\SAMPLE000033.D)

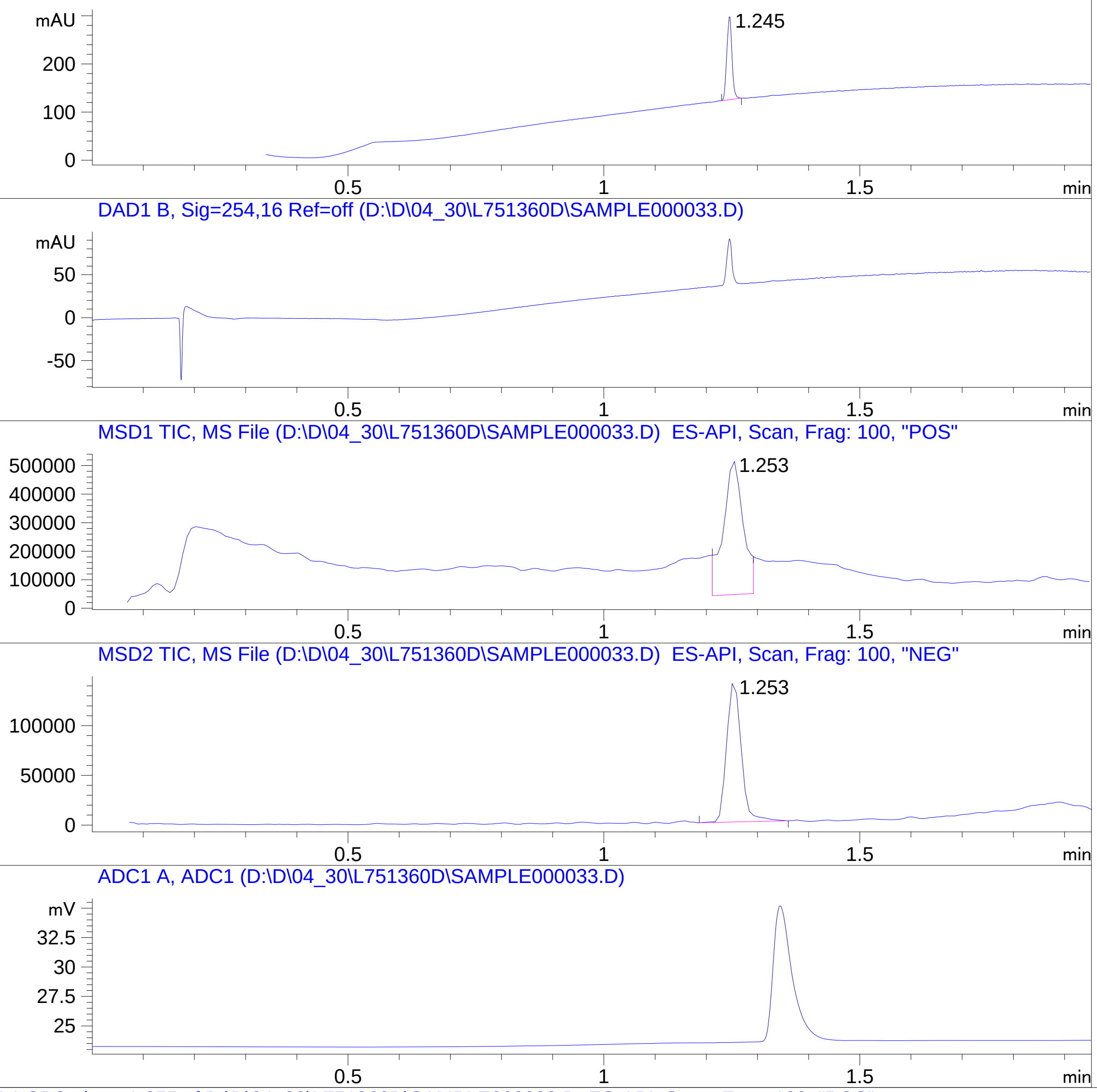


*MSD1 SPC, time=1.255 of D:\D\04_30\L751360D\SAMPLE000033.D ES-API, Scan, Frag: 100, "POS"

RT 1.253

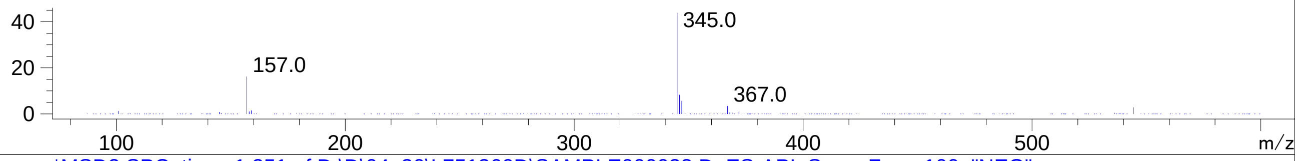


*MSD2 SPC, time=1.251 of D:\D\04_30\L751360D\SAMPLE000033.D ES-API, Scan, Frag: 100, "NEG"

RT 1.253

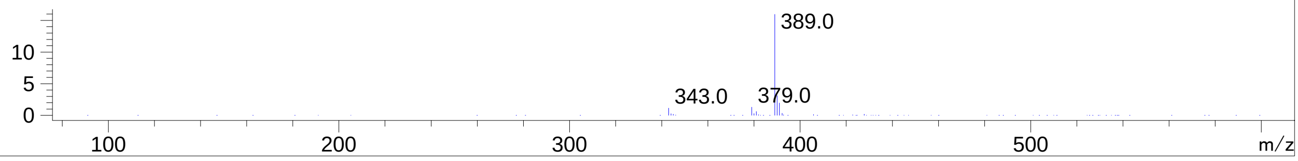


Inj.Date 4/30/2024
C:\CHEM32\1\METHODS\SUPOR_30.M

N
