## Supplementary Data 13 for "SyntheMol-RL: a flexible reinforcement learning framework for designing novel and synthesizable antibiotics": Z2273991831.PDF

MaxPeak: 63.70%
Ret_Time: 0.921 min

# BE177043$1

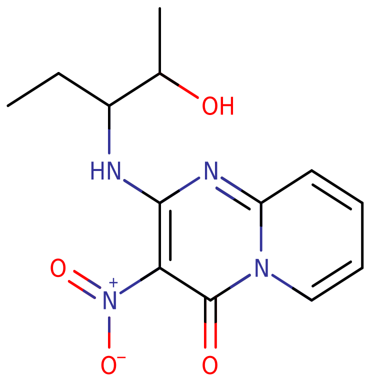


**Mol Wt** 292.29

**Exact Mass** 292.12

| # | Time | Area% |
|---|---|---|
| 1 | 0.908 | 36.30 |
| 2 | 0.921 | 63.70 |

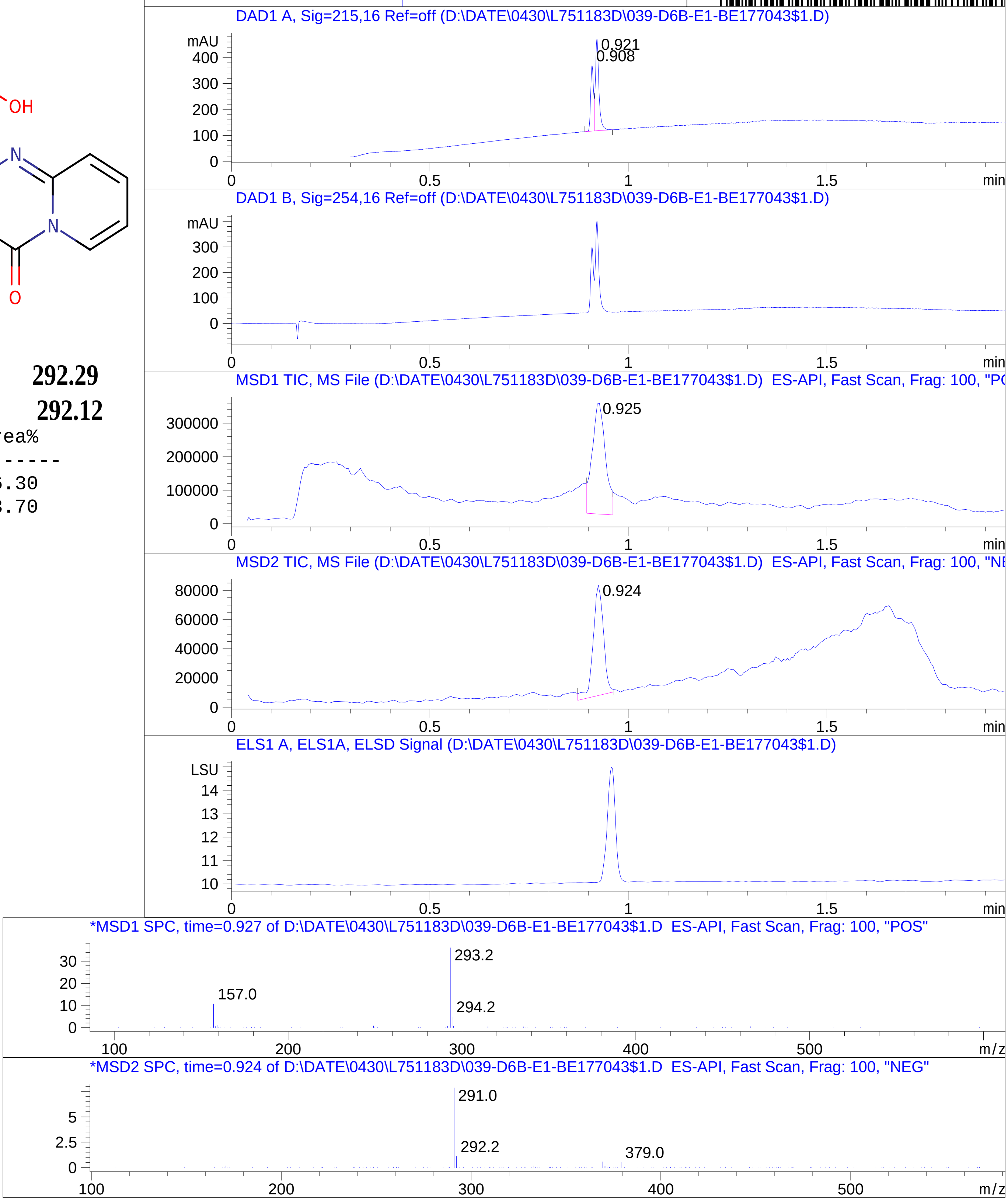


Inj.Date 4/30/2024 0
C:\Users\Public\Documents\ChemStation\1\Data\04_29\L751183D\SUPOR_30.M
