## Supplementary Data 13 for "SyntheMol-RL: a flexible reinforcement learning framework for designing novel and synthesizable antibiotics": Z2274035786.PDF

MaxPeak: 93.79%
Ret_Time: 0.833 min

# BE177026$2

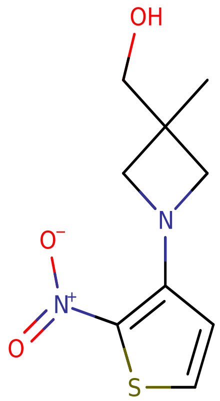


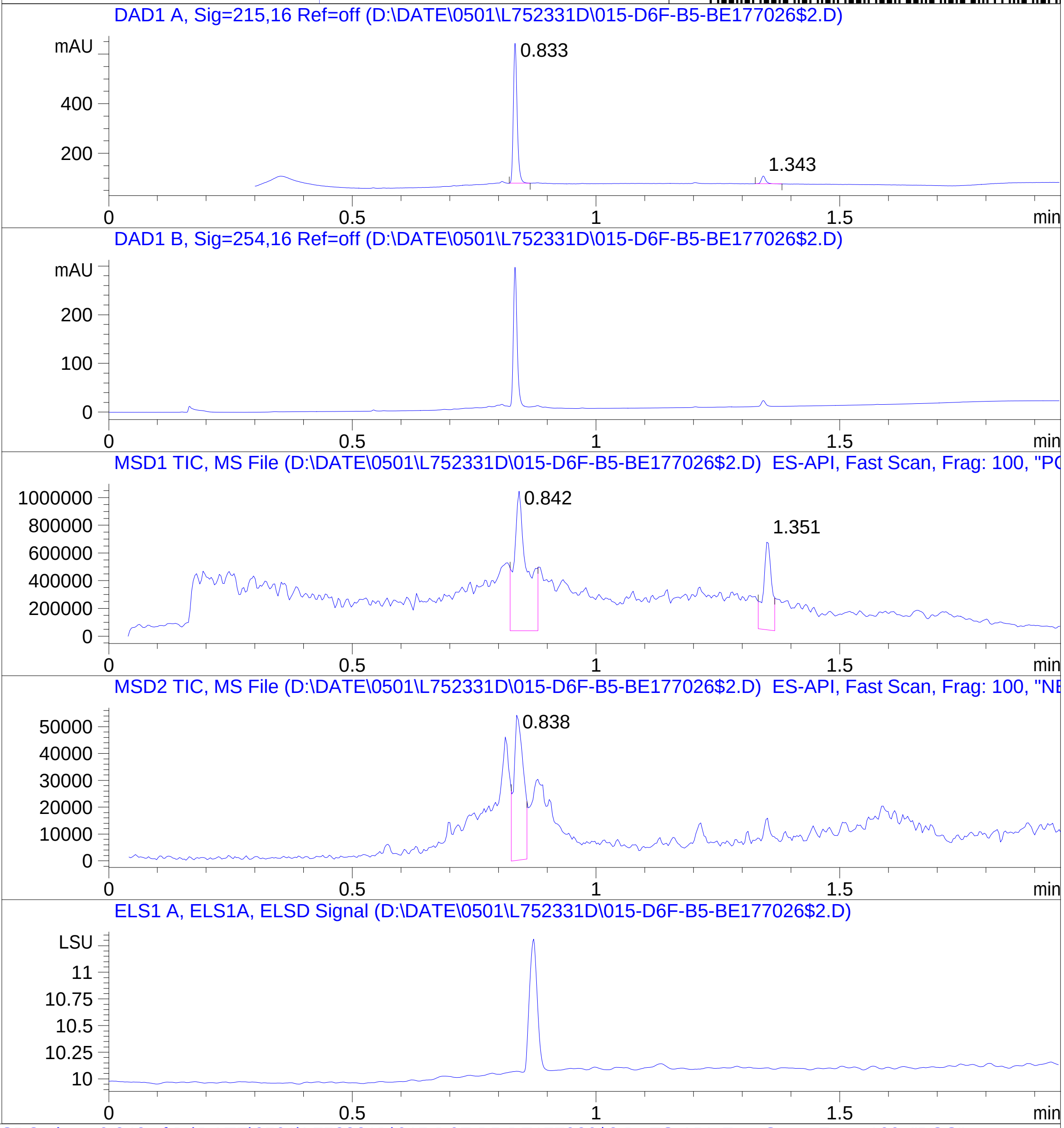


**Mol Wt** 228.27

**Exact Mass** 228.06

| # | Time | Area% |
|---|---|---|
| 1 | 0.833 | 93.79 |
| 2 | 1.343 | 6.21 |

RT 0.842

RT 1.351

RT 0.838

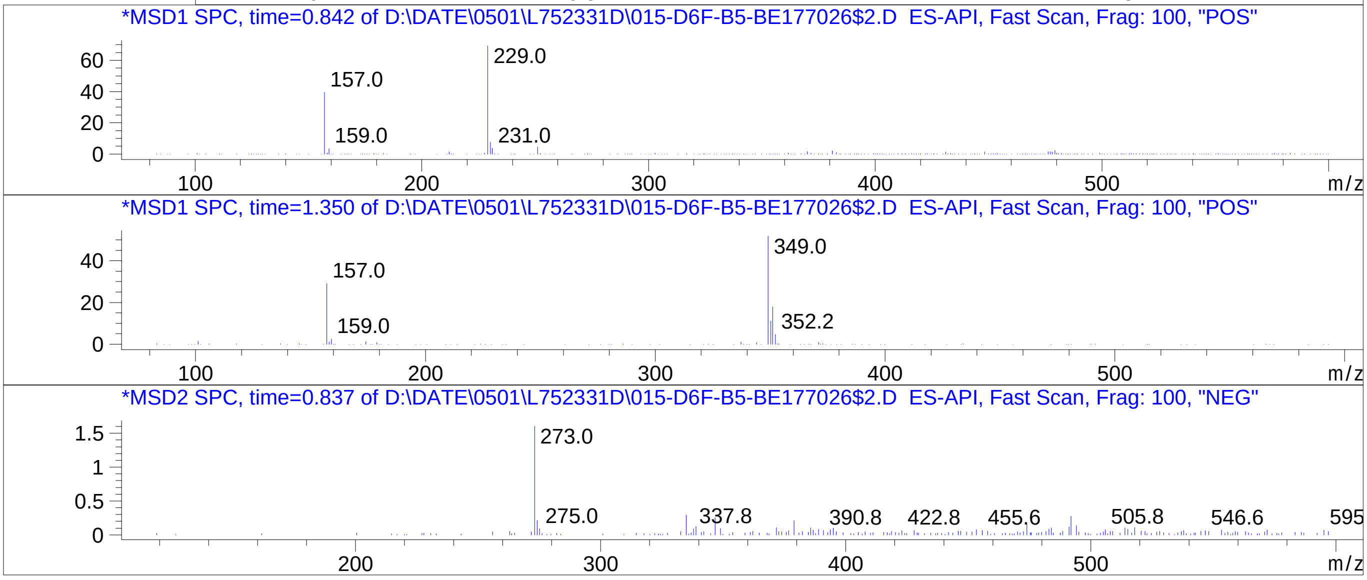


Inj.Date 5/1/2024 0
C:\Users\Public\Documents\ChemStation\1\Data\05_01\L752331D\SUPOR_30.M
