## Supplementary Data 13 for "SyntheMol-RL: a flexible reinforcement learning framework for designing novel and synthesizable antibiotics": Z2506480441.PDF

MaxPeak: 100.00%
Ret_Time: 0.709 min

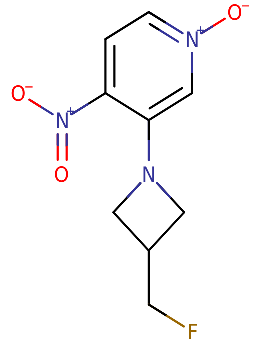


**Mol Wt** 227.19

**Exact Mass** 227.07

| # | Time | Area% |
|---|---|---|
| 1 | 0.709 | 100.00 |

# BE177068$11

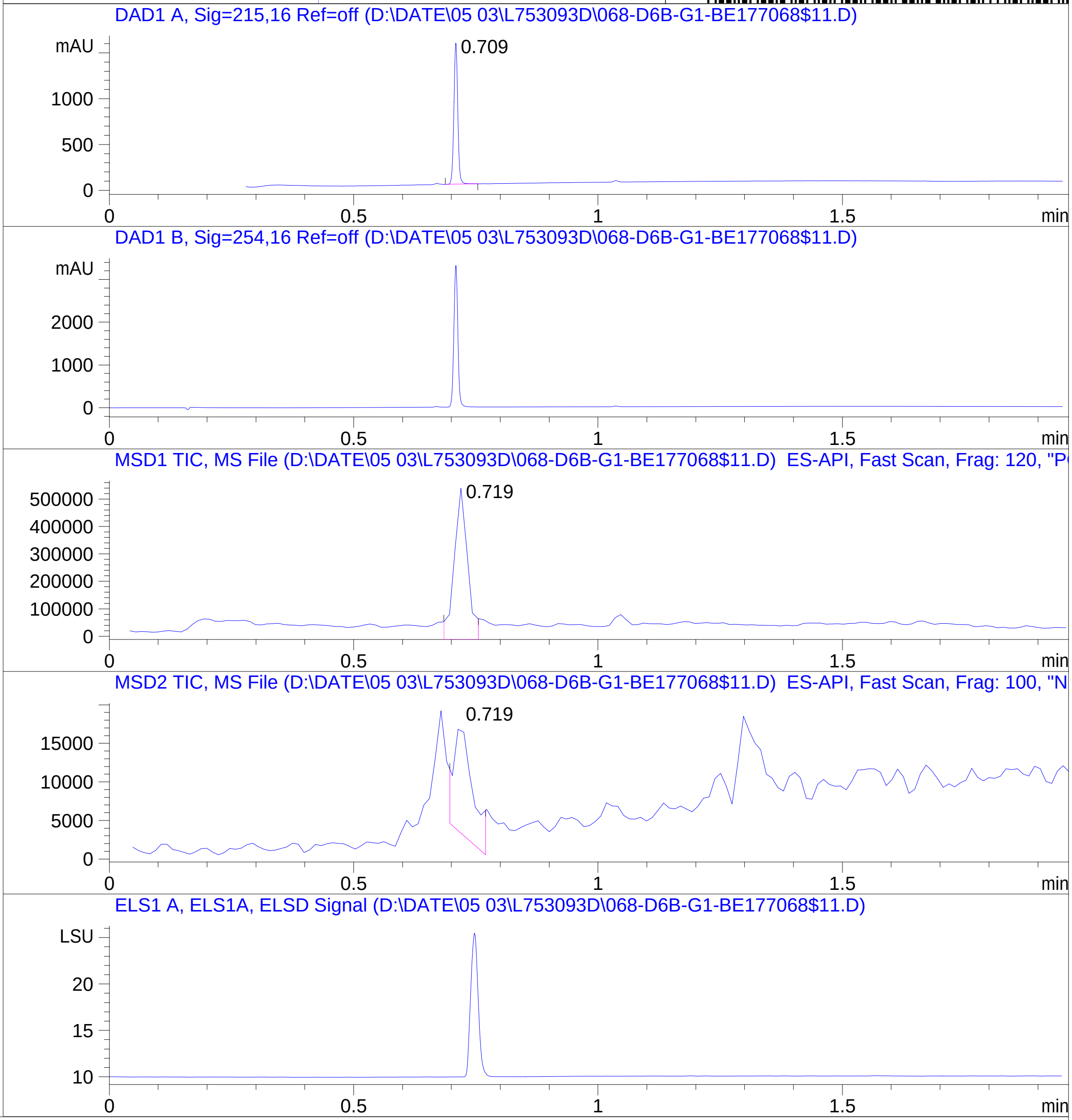


RT 0.719

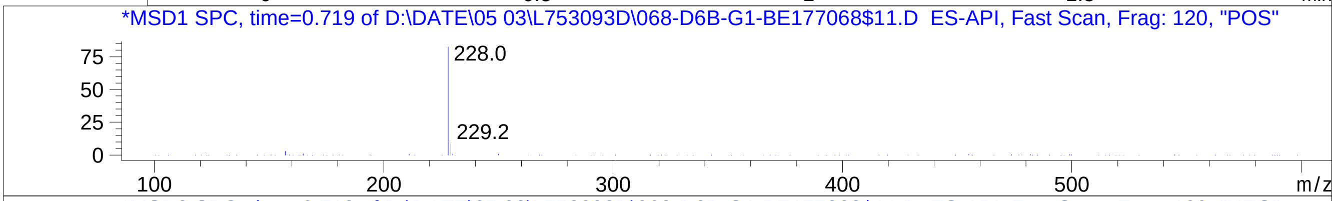


RT 0.719

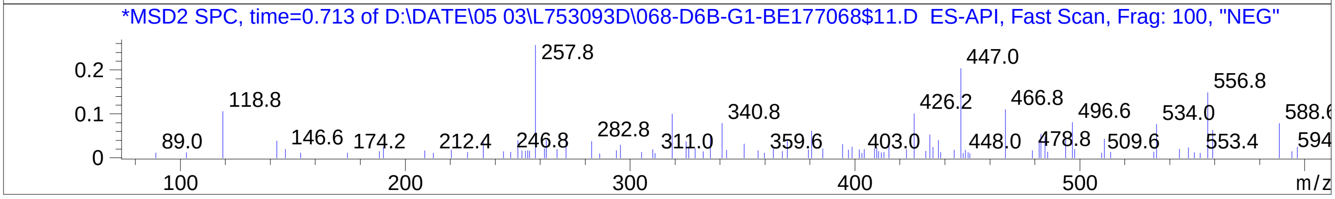


Inj.Date 5/3/2024 E 21
C:\Users\Public\Documents\ChemStation\1\Data\05_03\L753093D\SUPOR_96.M
