## Supplementary Data 13 for "SyntheMol-RL: a flexible reinforcement learning framework for designing novel and synthesizable antibiotics": Z2923649610.docx

CERTIFICATE of ANALYSIS

1. Identification

| Structure |  |
| --- | --- |
| Codes | Z2923649610 |
| Name | rac-1-[(3R,4R)-3-(hydroxymethyl)-4-methylpyrrolidin-1-yl]-2-[(5-sulfanyl-1,3,4-thiadiazol-2-yl)sulfanyl]ethan-1-one |
| Formula | C10H15N3O2S3 |
| Formula weight | 305.44 |
| CAS number |  |

2. Description

| Appearance | crystalline powder |
| --- | --- |
| Color | beige |
| Melting point, °C | N/A |
| Boiling point, °C | Not determined |

3. NMR spectra

| File name, *wmf | N/A |
| --- | --- |
| Identity | N/A |

4. LCMS data

| File name, *pdf | Z2923649610 |
| --- | --- |
| UV Area, % | 93.19 |

5. Comments

| Comments | No special comments. |
| --- | --- |

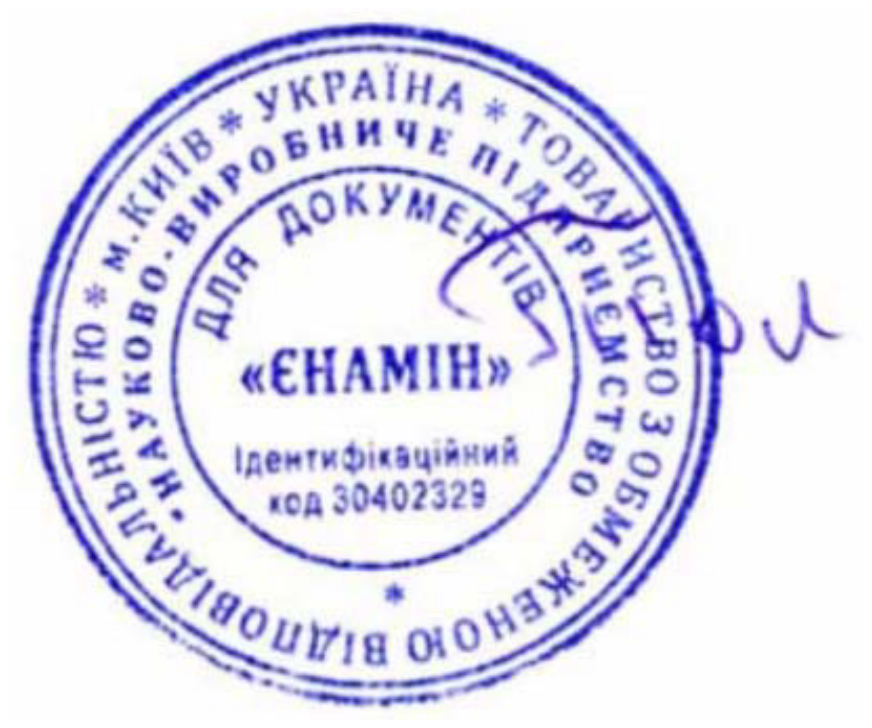

Sincerely yours,

Name: A. Konovets

Title: Head of QC department

Name of Department: QC department

Company name: Enamine Ltd.

MATERIAL SAFETY DATA

**Sheet**

**1. Product Information**

Product Name:	**rac-1-[(3R,4R)-3-(hydroxymethyl)-4-methylpyrrolidin-1-yl]-2-[(5-sulfanyl-1,3,4-thiadiazol-2-yl)sulfanyl]ethan-1-one**

Product Catalogue Number:	Z2923649610

CAS number:

Information of company: Enamine US Inc

1 Distribution Way

Monmouth Junction, NJ 08852

USA

**Experimental Product for Research&Development Use Only. Not for Drug, Household or Other Use**

**2. Composition/Information on Ingredients**

Product Name:	**rac-1-[(3R,4R)-3-(hydroxymethyl)-4-methylpyrrolidin-1-yl]-2-[(5-sulfanyl-1,3,4-thiadiazol-2-yl)sulfanyl]ethan-1-one**

Formula:	C10H15N3O2S3

Molecular Weight:	305.44 AMU

**3. Hazards Identification**

SPECIAL INDICATION OF HAZARDS TO HUMANS AND THE ENVIRONMENT

Harmful if swallowed or inhaled.

__________________________________________________________________

**4 - First Aid Measures**

__________________________________________________________________

AFTER INHALATION

If inhaled, remove to fresh air. If not breathing give artificial respiration. If breathing is difficult, give oxygen.

AFTER SKIN CONTACT

In case of skin contact, flush with copious amounts of water for at least 15 minutes. Remove contaminated clothing and shoes.

Call a physician.

AFTER EYE CONTACT

In case of contact with eyes, flush with copious amounts of water for at least 15 minutes. Assure adequate flushing by separating the eyelids with fingers. Call a physician.

AFTER INGESTION

If swallowed, wash out mouth with water provided person is conscious. Call a physician.

_______________________________________________________________________

**5 - Fire Fighting Measures**

______________________________________________________________________

EXTINGUISHING MEDIA

Suitable: Water spray. Carbon dioxide, dry chemical powder, or appropriate foam.

SPECIAL RISKS

Specific Hazard(s): Emits toxic fumes under fire conditions.

SPECIAL PROTECTIVE EQUIPMENT FOR FIREFIGHTERS

Wear self-contained breathing apparatus and protective clothing to prevent contact with skin and eyes.

________________________________________________________________________

**6 - Accidental Release Measures**

_________________________________________________________________________

PERSONAL PRECAUTION PROCEDURES TO BE FOLLOWED IN CASE OF LEAK OR SPILL

Evacuate area.

PROCEDURE(S) OF PERSONAL PRECAUTION(S)

Wear self-contained breathing apparatus, rubber boots, and heavy

rubber gloves.

METHODS FOR CLEANING UP

Wipe dry, place a rag in a bag and hold for waste disposal. Avoid fumes inhaling. Ventilate area and wash spill site after material pickup is complete.

_________________________________________________________________________________

**7 - Handling and Storage**

_________________________________________________________________________________

HANDLING

Directions for Safe Handling: Do not breathe vapor. Avoid contact with eyes, skin, and clothing. Avoid prolonged or repeated exposure.

STORAGE

Conditions of Storage: Keep tightly closed. Keep in room temperature.

Expire date: not available, reanalysis is required no more than once a year.

SPECIAL REQUIREMENTS: -

____________________________________________________________________________________

**8 - Exposure Controls / Personal Protection**

____________________________________________________________________________________

ENGINEERING CONTROLS

Safety shower and eye bath. Mechanical exhaust required.

___________________________________________________________________________________

**14 - Transport Information**

___________________________________________________________________________________

Non-hazard for transport. The above goods is/are non-corrosive, non-oxidizing, non-magnetic, non-toxic and not dangerous and it can be carried in any passenger aircraft.

___________________________________________________________________________________

**15 - Regulatory Information**

___________________________________________________________________________________

Safety Statements: Do not breathe dust. Avoid contact with skin and eyes.

________________________________________________________________________________

__

**16 - Other Information**

___________________________________________________________________________________

For R&D use only. Not for drug, household or other uses.

This is an experimental product whose properties are not fully evaluated yet. The information contained herein is based on the present state of our knowledge and therefore does not guarantee certain properties. Recipients of our product must take responsibility for observing existing laws and regulations.

___________________________________________________________________________________
