## Supplementary Data 13 for "SyntheMol-RL: a flexible reinforcement learning framework for designing novel and synthesizable antibiotics": Z3467229414.PDF

# BE177002$42

MaxPeak:  100.00%
Ret_Time: 1.050 min

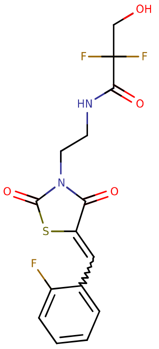


**Mol Wt** 374.34

**Exact Mass** 374.06

| # | Time | Area% |
|---|---|---|
| 1 | 1.050 | 100.00 |

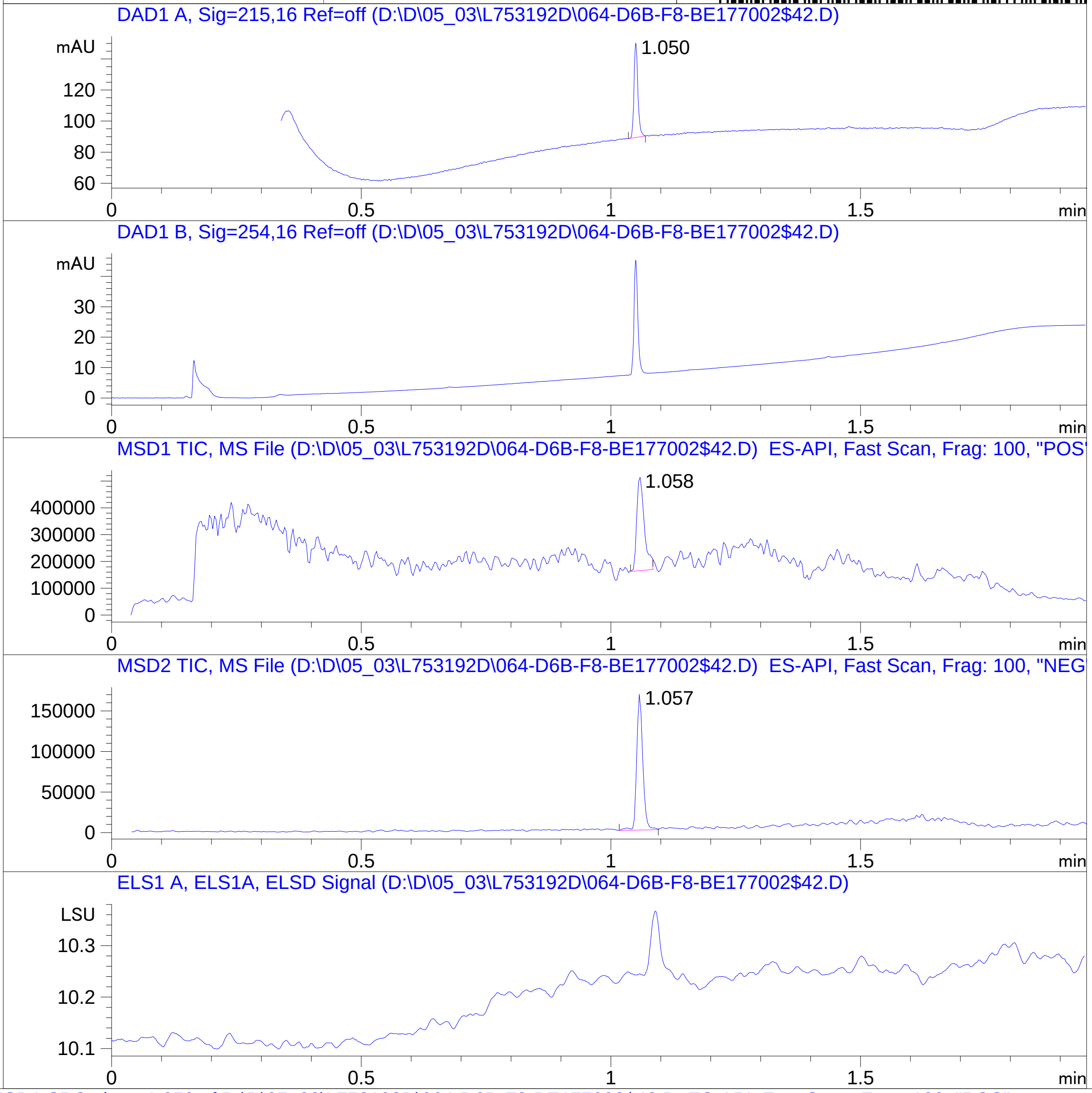


RT 1.058

RT 1.057

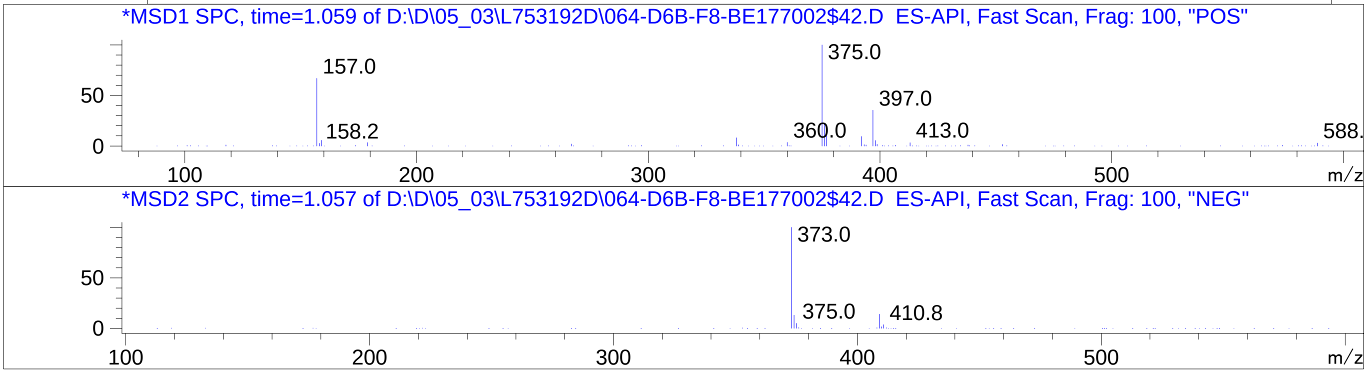


Inj.Date 5/3/2024 N
C:\Users\Public\Documents\ChemStation\1\Data\05_03\L753192D\SUPOR_30.M
