## Supplementary Data 13 for "SyntheMol-RL: a flexible reinforcement learning framework for designing novel and synthesizable antibiotics": Z3561876565.PDF

MaxPeak: 100.00%
Ret_Time: 0.993 min

# BE177069$1

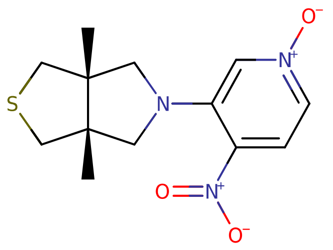


**Mol Wt** 295.36

**Exact Mass** 295.11

| # | Time | Area% |
|---|---|---|
| 1 | 0.993 | 100.00 |

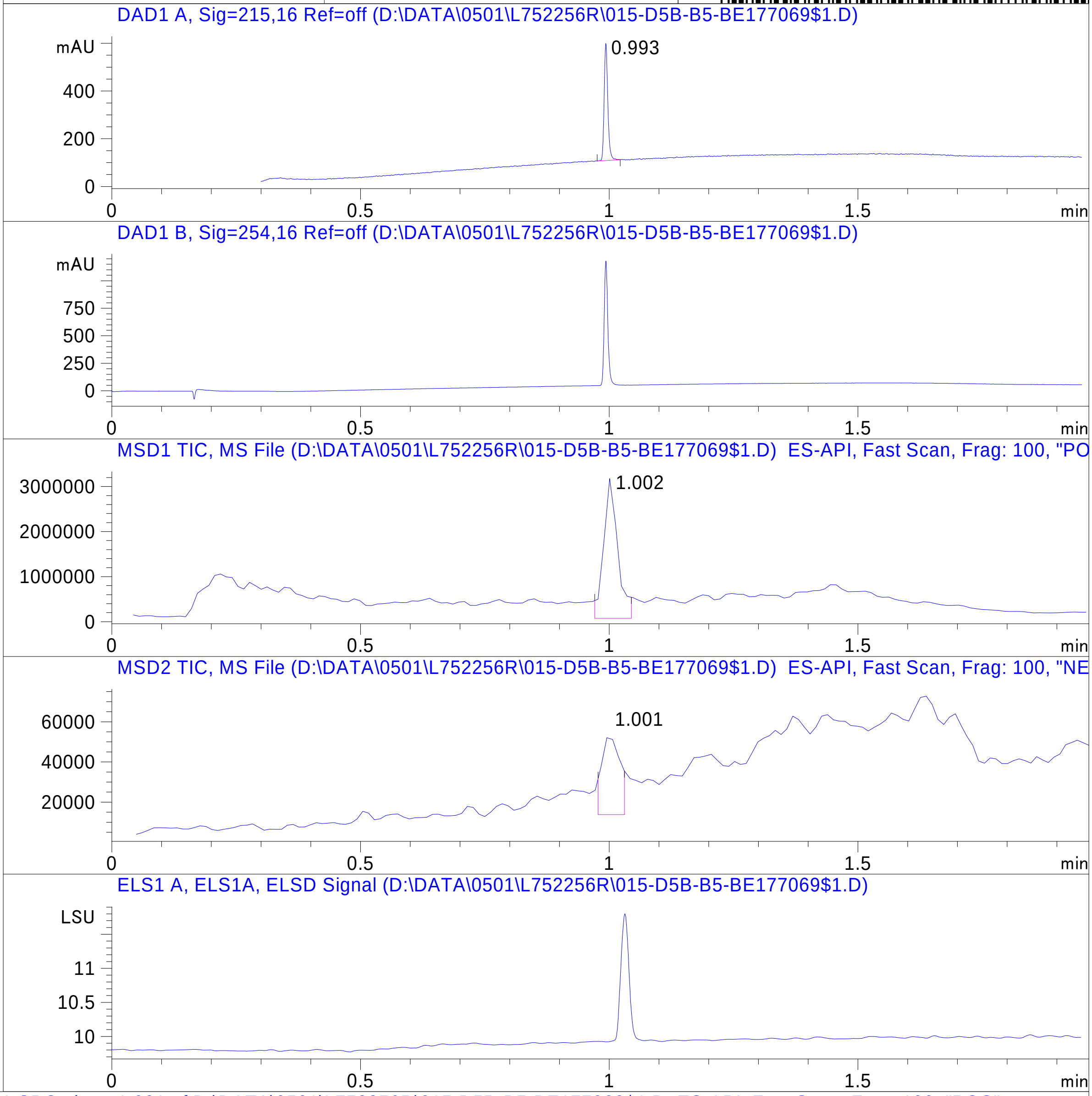


RT 1.002

RT 1.001

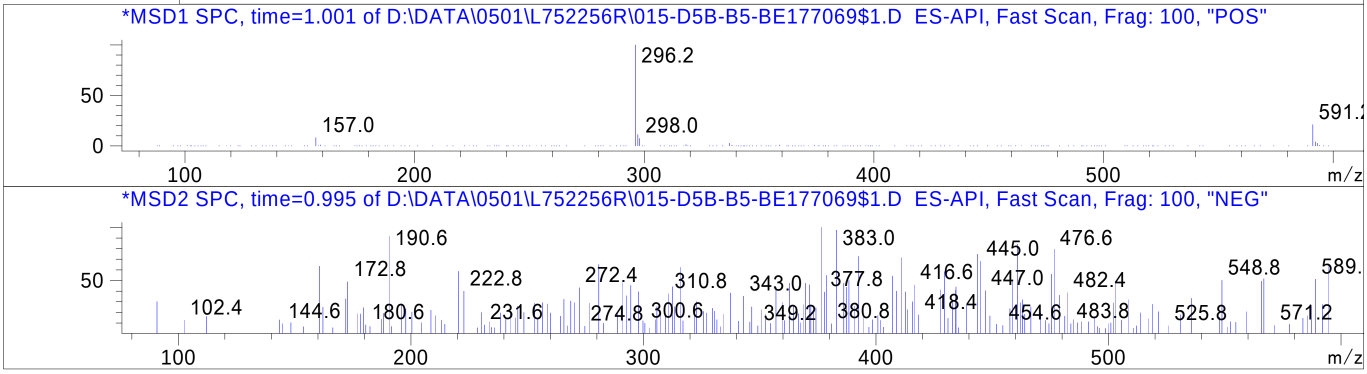


Inj.Date 5/1/2024 H 31

C:\Users\Public\Documents\ChemStation\1\Data\05_01\L752256R\SUPOR_30.M
