## Supplementary Data 13 for "SyntheMol-RL: a flexible reinforcement learning framework for designing novel and synthesizable antibiotics": Z3578472114.PDF

# BD252399$2

MaxPeak: 99.04%
Ret_Time: 1.216 min

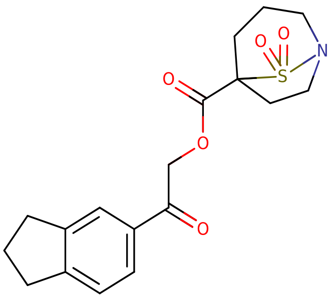


**Mol Wt** 363.43
**Exact Mass** 363.13

| # | Time | Area% |
|---|---|---|
| 1 | 1.216 | 99.04 |
| 2 | 1.300 | 0.96 |

DAD1 A, Sig=215,16 Ref=off (D:\DATE\02 29\L726103D\022-D5B-C3-BD252399$2.D)

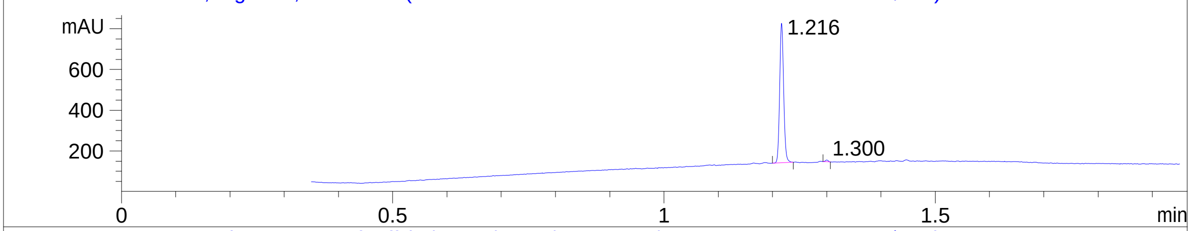


DAD1 B, Sig=254,16 Ref=off (D:\DATE\02 29\L726103D\022-D5B-C3-BD252399$2.D)

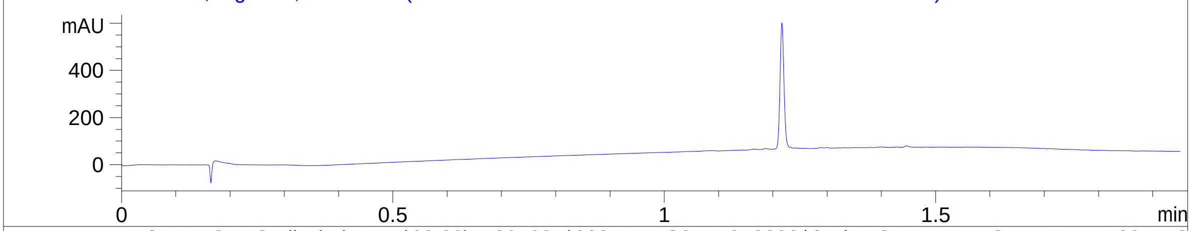


MSD1 TIC, MS File (D:\DATE\02 29\L726103D\022-D5B-C3-BD252399$2.D) ES-API, Fast Scan, Frag: 100, "PO

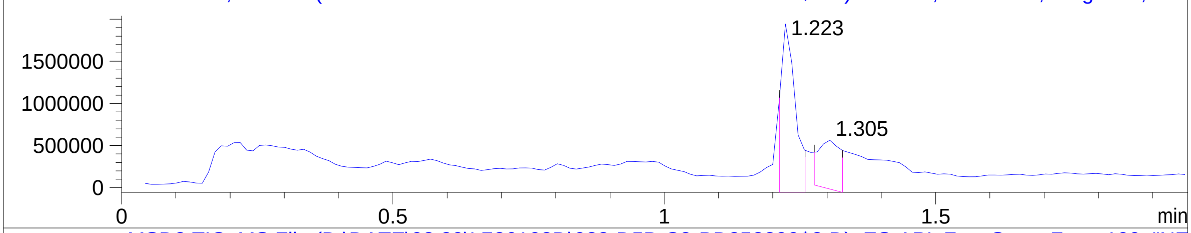


MSD2 TIC, MS File (D:\DATE\02 29\L726103D\022-D5B-C3-BD252399$2.D) ES-API, Fast Scan, Frag: 100, "NE

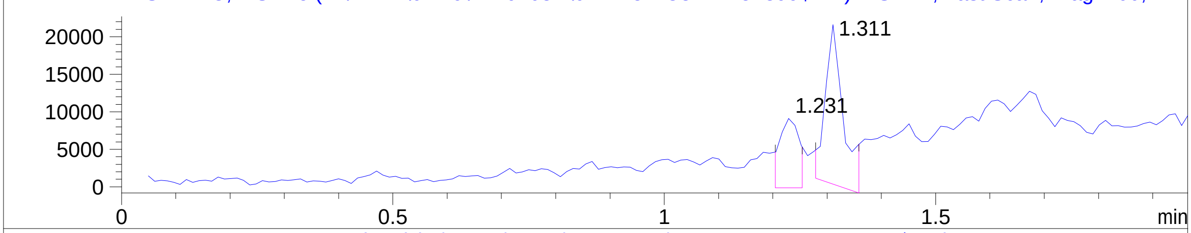


ELS1 A, ELS1A, ELSD Signal (D:\DATE\02 29\L726103D\022-D5B-C3-BD252399$2.D)

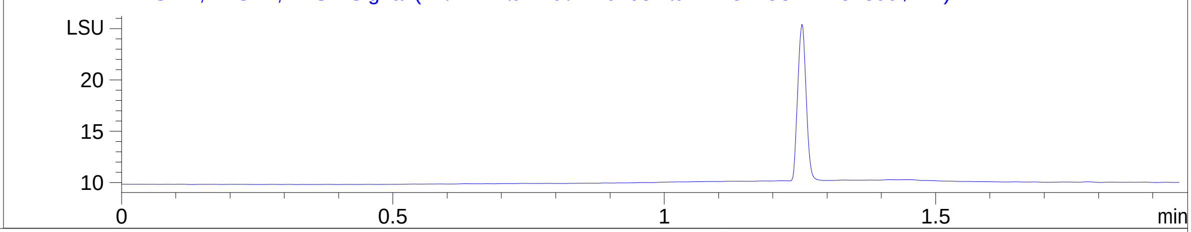


RT 1.223

*MSD1 SPC, time=1.223 of D:\DATE\02 29\L726103D\022-D5B-C3-BD252399$2.D ES-API, Fast Scan, Frag: 100, "POS"

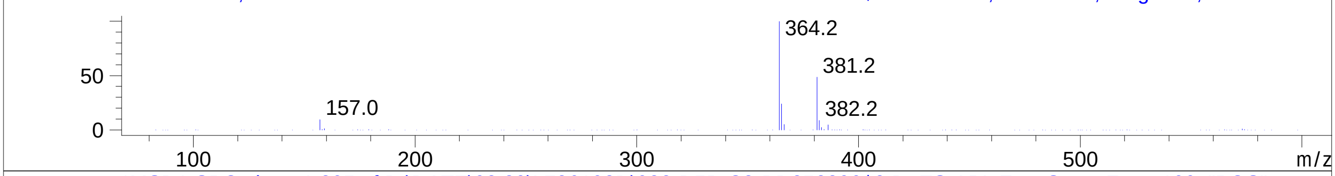


RT 1.305

*MSD1 SPC, time=1.305 of D:\DATE\02 29\L726103D\022-D5B-C3-BD252399$2.D ES-API, Fast Scan, Frag: 100, "POS"

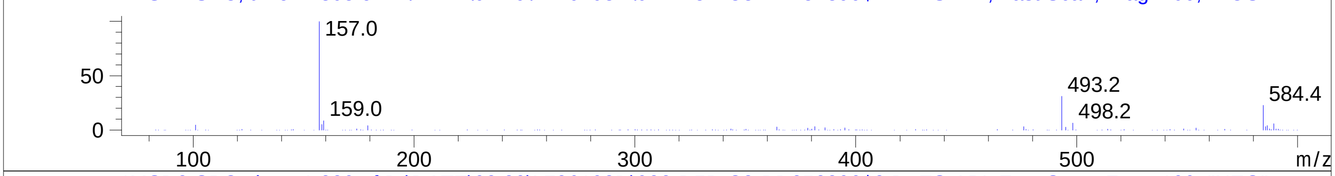


RT 1.231

*MSD2 SPC, time=1.229 of D:\DATE\02 29\L726103D\022-D5B-C3-BD252399$2.D ES-API, Fast Scan, Frag: 100, "NEG"

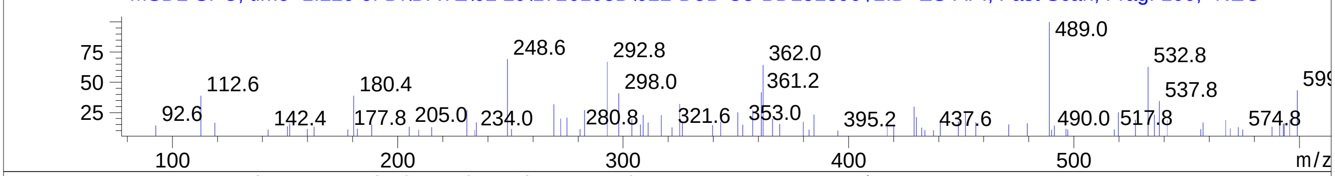


RT 1.311

*MSD2 SPC, time=1.311 of D:\DATE\02 29\L726103D\022-D5B-C3-BD252399$2.D ES-API, Fast Scan, Frag: 100, "NEG"

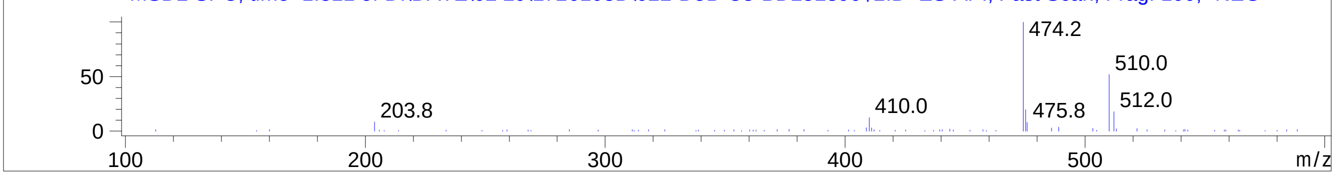


Inj.Date 2/29/2024 E 31
C:\Users\Public\Documents\ChemStation\1\Data\02_29\L726103D\SUPOR_30.M
