## Supplementary Data 13 for "SyntheMol-RL: a flexible reinforcement learning framework for designing novel and synthesizable antibiotics": Z3578475252.PDF

MaxPeak:  100.00%
Ret_Time: 0.873 min

# BD252452$1

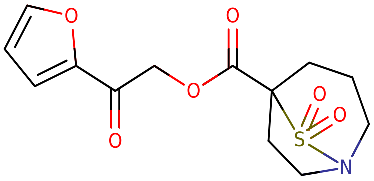

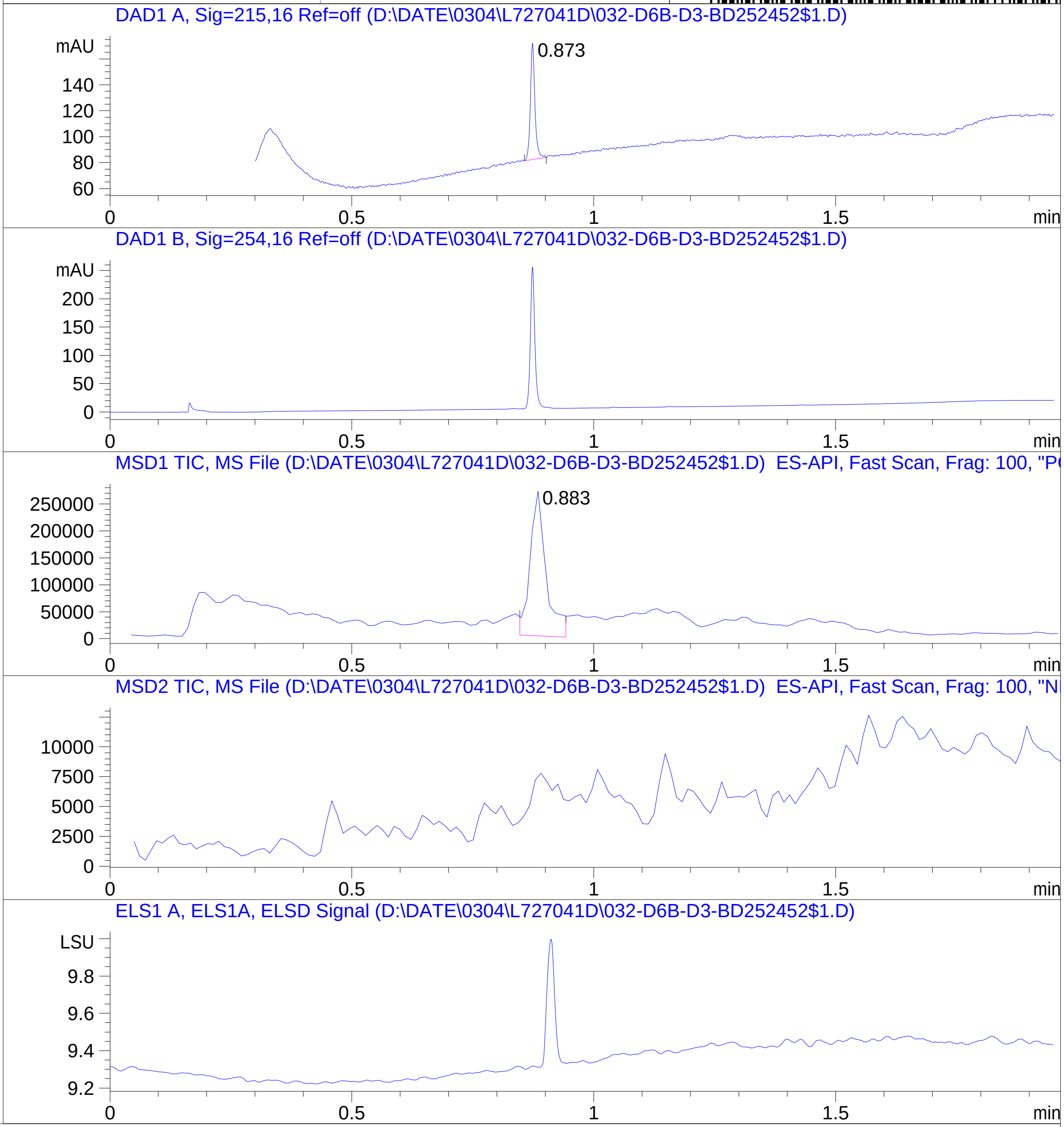


**Mol Wt** 313.33

**Exact Mass** 313.06

```
#  Time    Area%
-----------------
1  0.873  100.00
```

RT  0.883

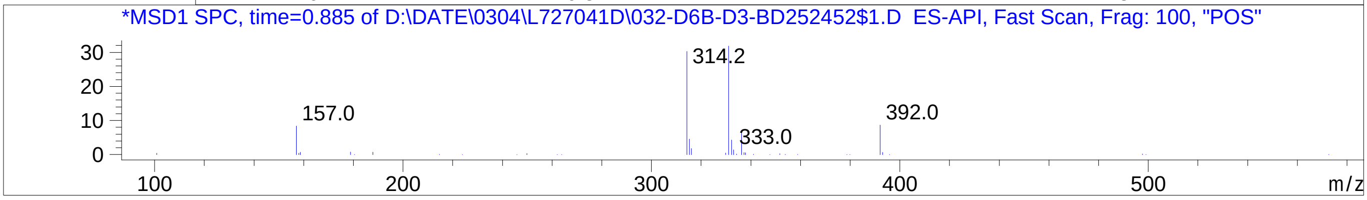


Inj.Date 3/4/2024    O
C:\Users\Public\Documents\ChemStation\1\Data\03_04\L727041D\SUPOR_30.M
