## Supplementary Data 13 for "SyntheMol-RL: a flexible reinforcement learning framework for designing novel and synthesizable antibiotics": Z4309747056.PDF

MaxPeak: 93.31%
Ret_Time: 1.065 min

# BD252455$A

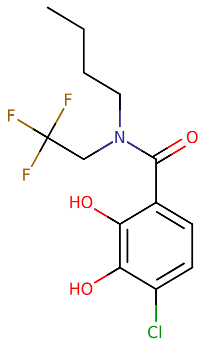


**Mol Wt** 325.71

**Exact Mass** 325.09

| # | Time | Area% |
|---|---|---|
| 1 | 0.707 | 1.13 |
| 2 | 0.913 | 1.21 |
| 3 | 1.065 | 93.31 |
| 4 | 1.199 | 4.35 |

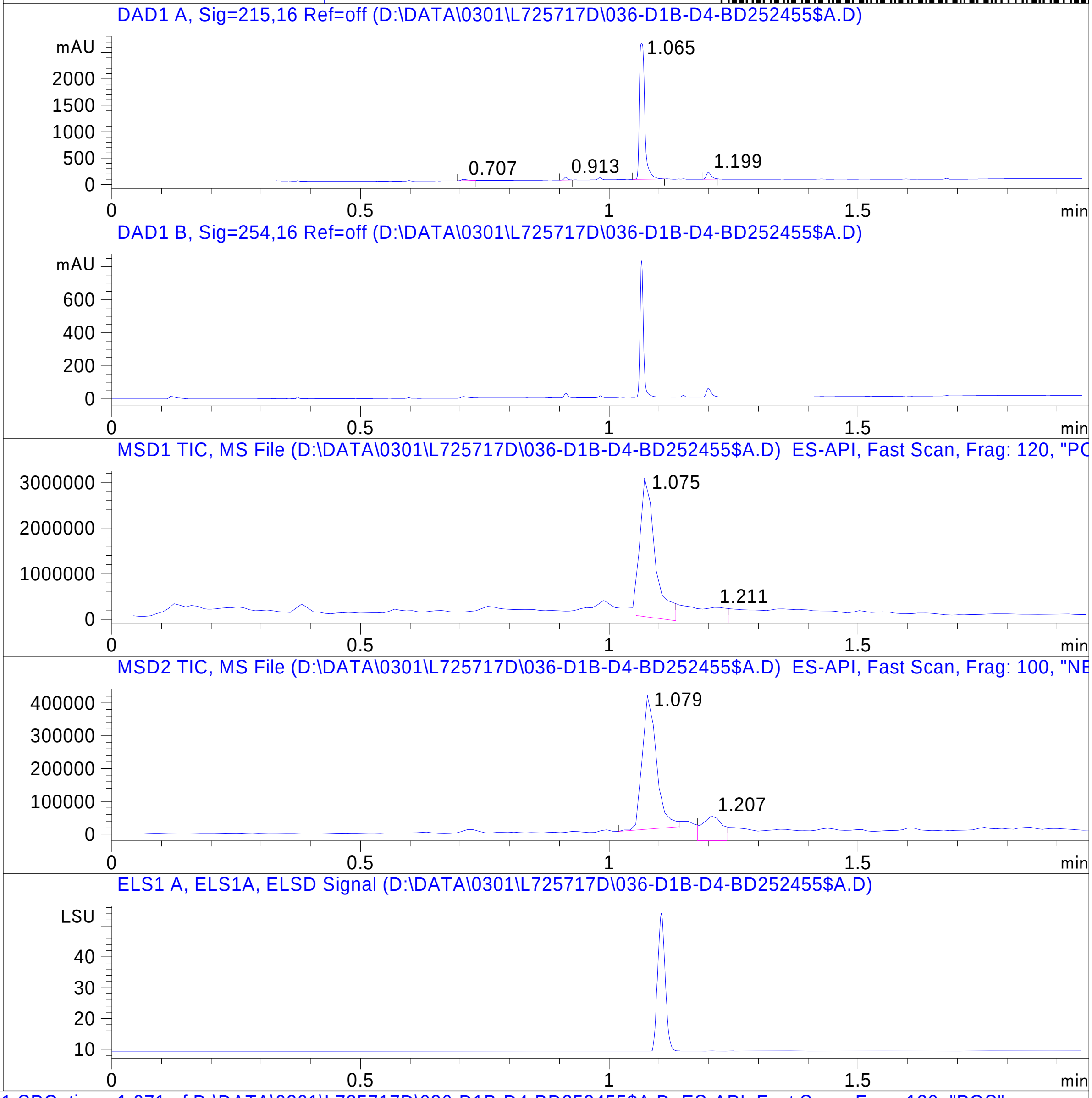


RT 1.075

RT 1.211

RT 1.079

RT 1.207

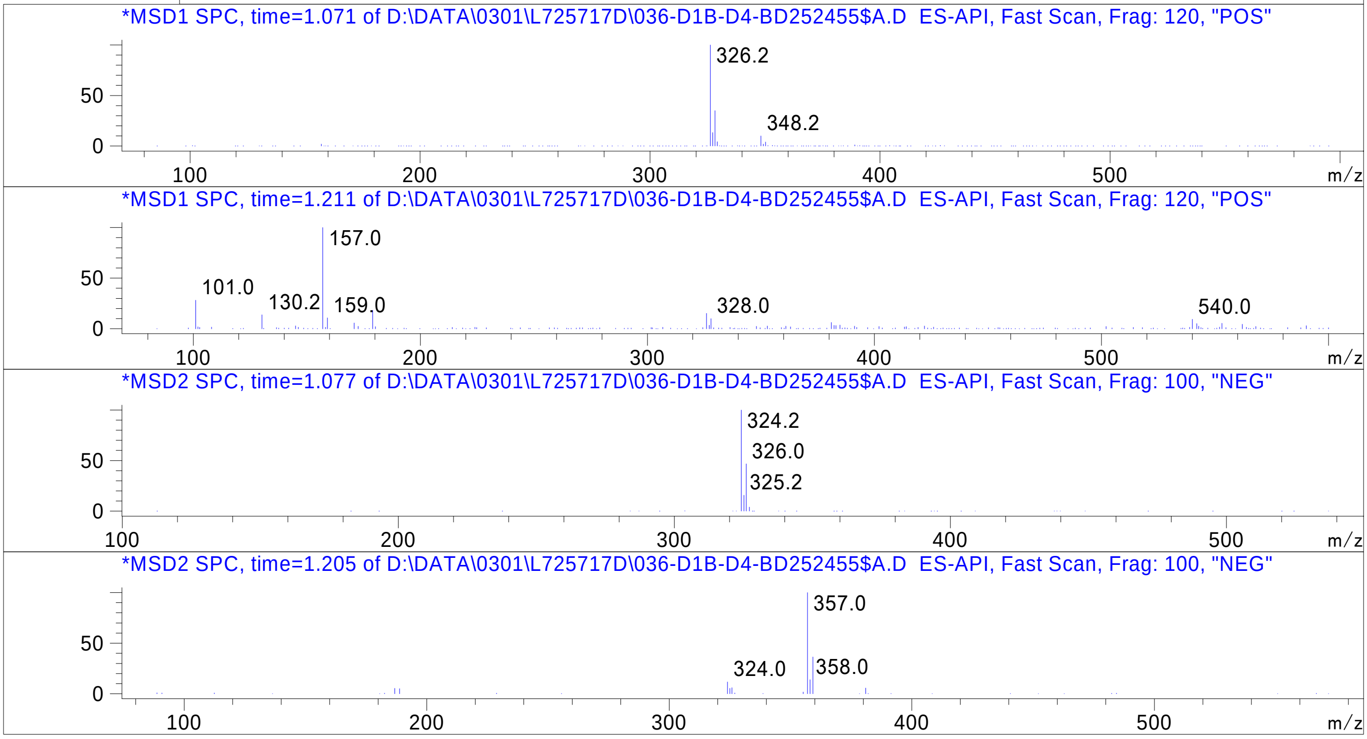


Inj.Date 2/29/2024 H

C:\Users\Public\Documents\ChemStation\1\Data\02_29\L725717D\SUPOR_96.M
