## Supplementary Data 13 for "SyntheMol-RL: a flexible reinforcement learning framework for designing novel and synthesizable antibiotics": Z5082764588.PDF

MaxPeak: 97.39%
Ret_Time: 0.803 min

# BE224663$9

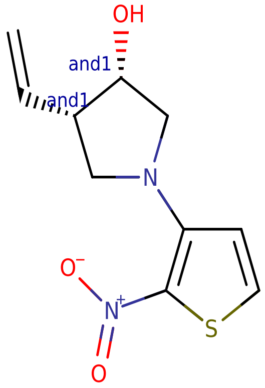


**Mol Wt** 240.28

**Exact Mass** 240.06

| # | Time | Area% |
|---|---|---|
| 1 | 0.803 | 97.39 |
| 2 | 0.847 | 2.61 |

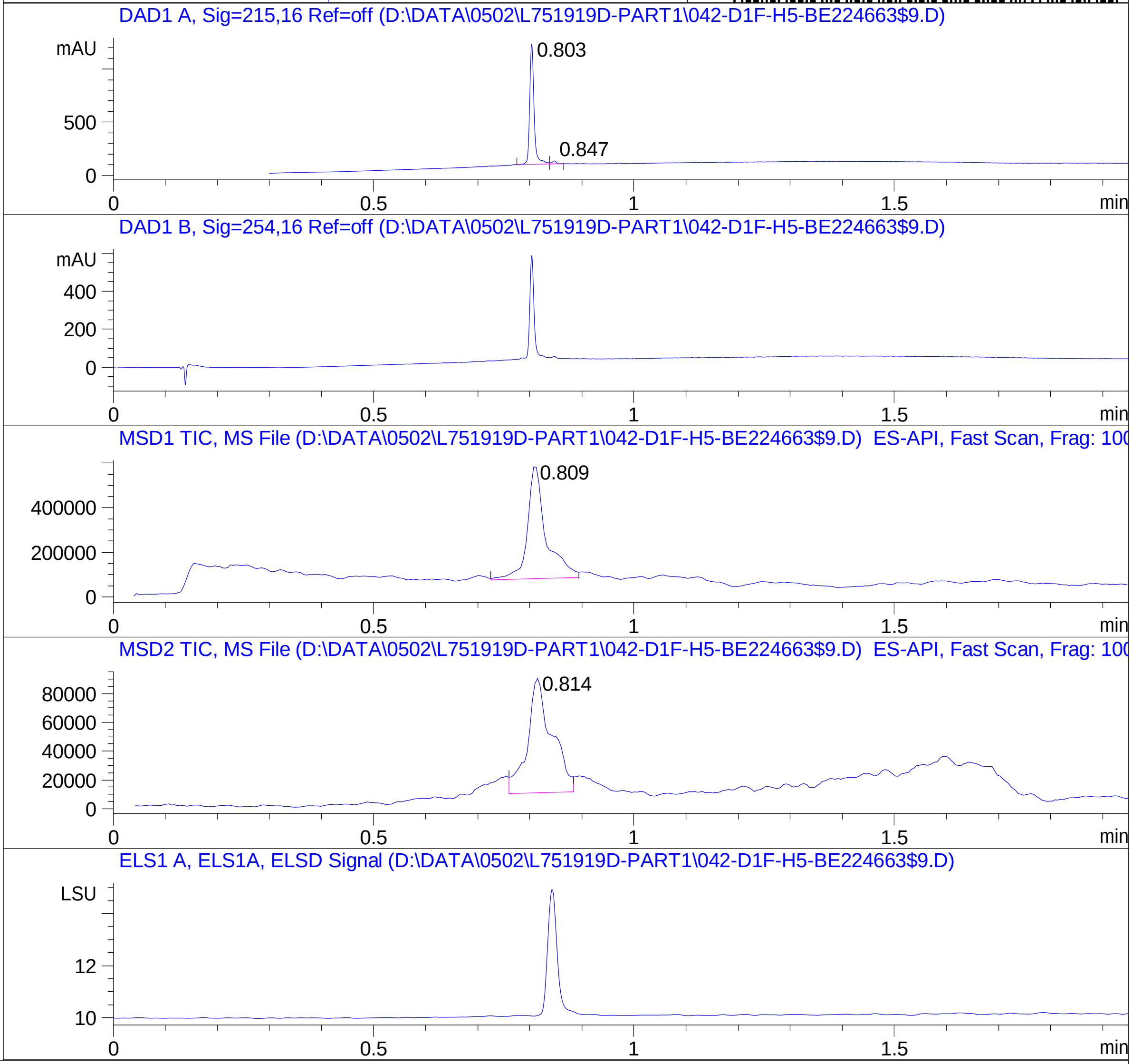


RT 0.809

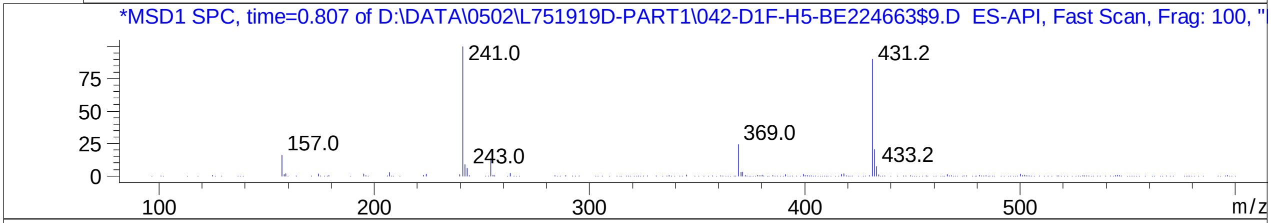


RT 0.814

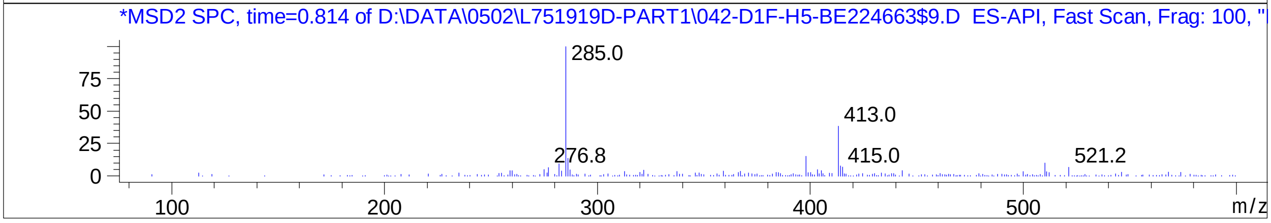


Inj.Date 5/1/2024 LB

C:\Users\Public\Documents\ChemStation\1\Data\05_01\1751919D\SUPOR_96.M
