## Supplementary Data 13 for "SyntheMol-RL: a flexible reinforcement learning framework for designing novel and synthesizable antibiotics": Z5401174482.PDF

MaxPeak: 96.48%
Ret_Time: 1.154 min

# BE177071$1

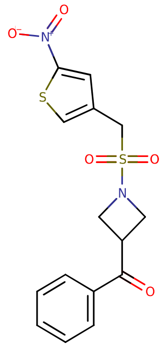


**Mol Wt** 366.41

**Exact Mass** 366.03

| # | Time | Area% |
|---|---|---|
| 1 | 1.124 | 3.52 |
| 2 | 1.154 | 96.48 |

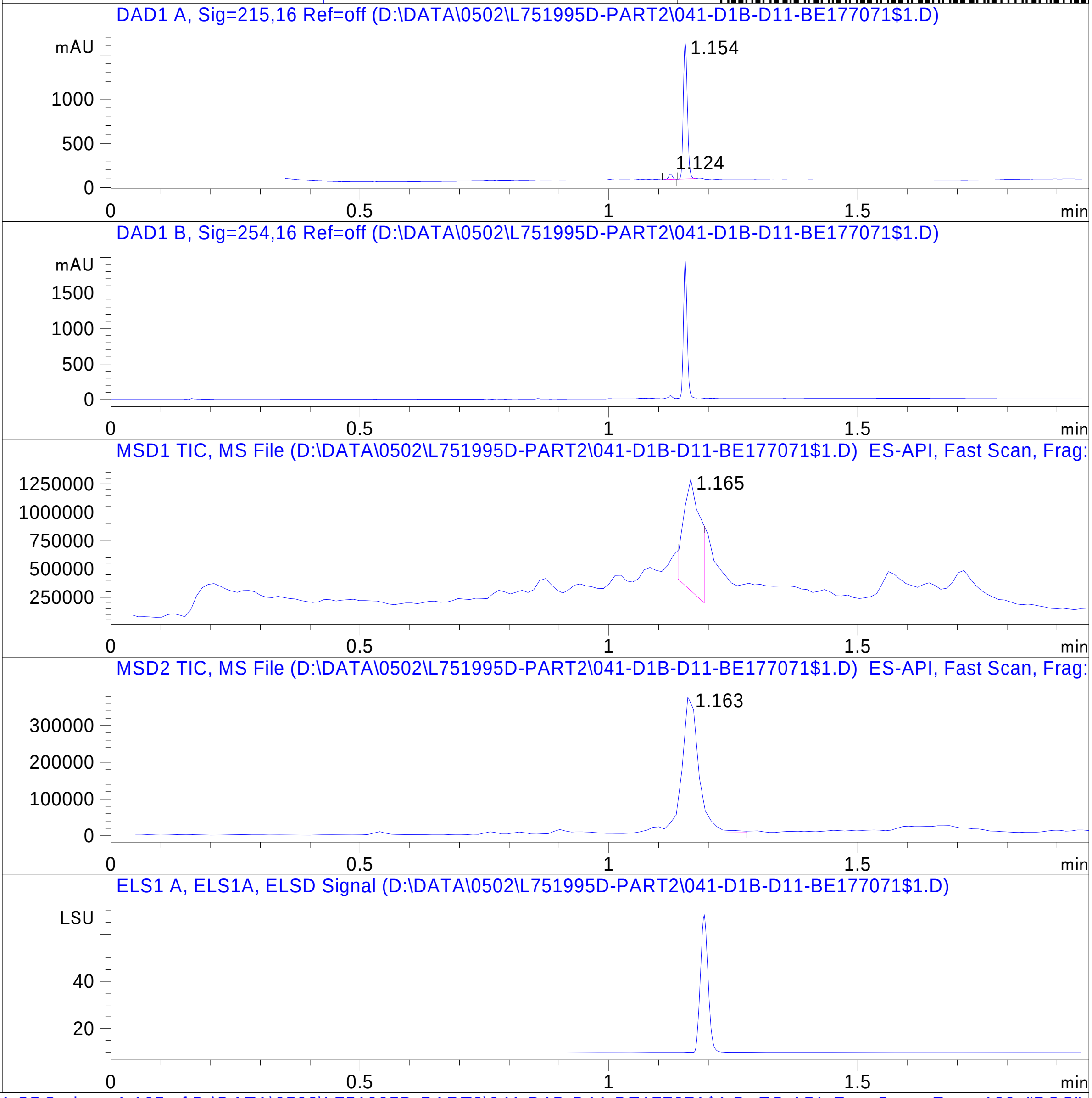


RT 1.165

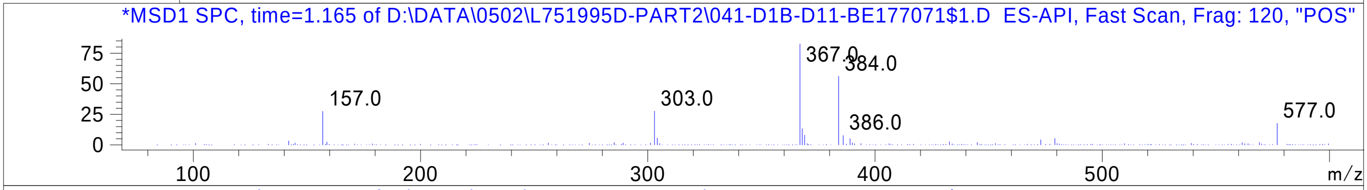


RT 1.163

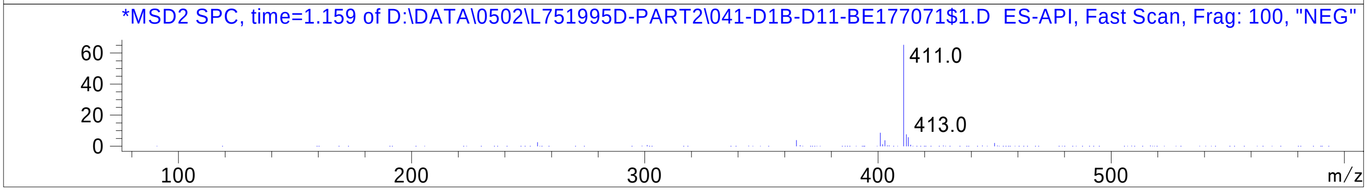


Inj.Date 5/1/2024 H

C:\Users\Public\Documents\ChemStation\1\Data\05_01\2751995D\SUPOR_96.M
