## Supplementary Data 13 for "SyntheMol-RL: a flexible reinforcement learning framework for designing novel and synthesizable antibiotics": Z5887221497.PDF

# BE177016$5

MaxPeak: 100.00%
Ret_Time: 1.135 min

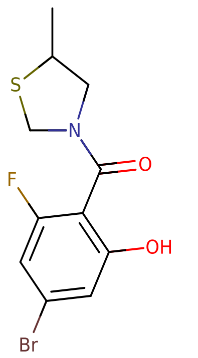


**Mol Wt** 320.18

**Exact Mass** 320.98

| # | Time | Area% |
|---|---|---|
| 1 | 1.135 | 100.00 |

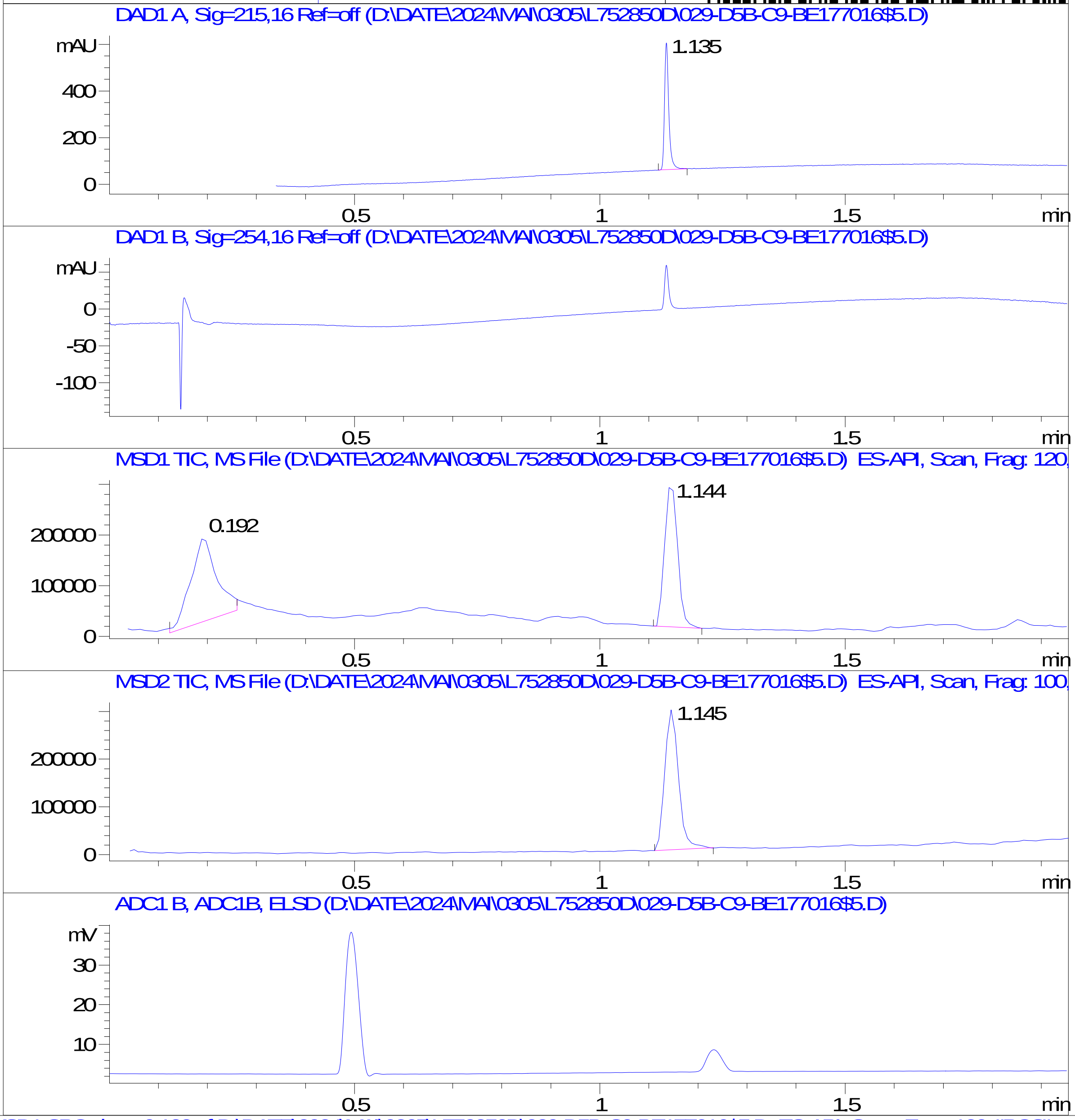


RT 0.192

RT 1.144

RT 1.145

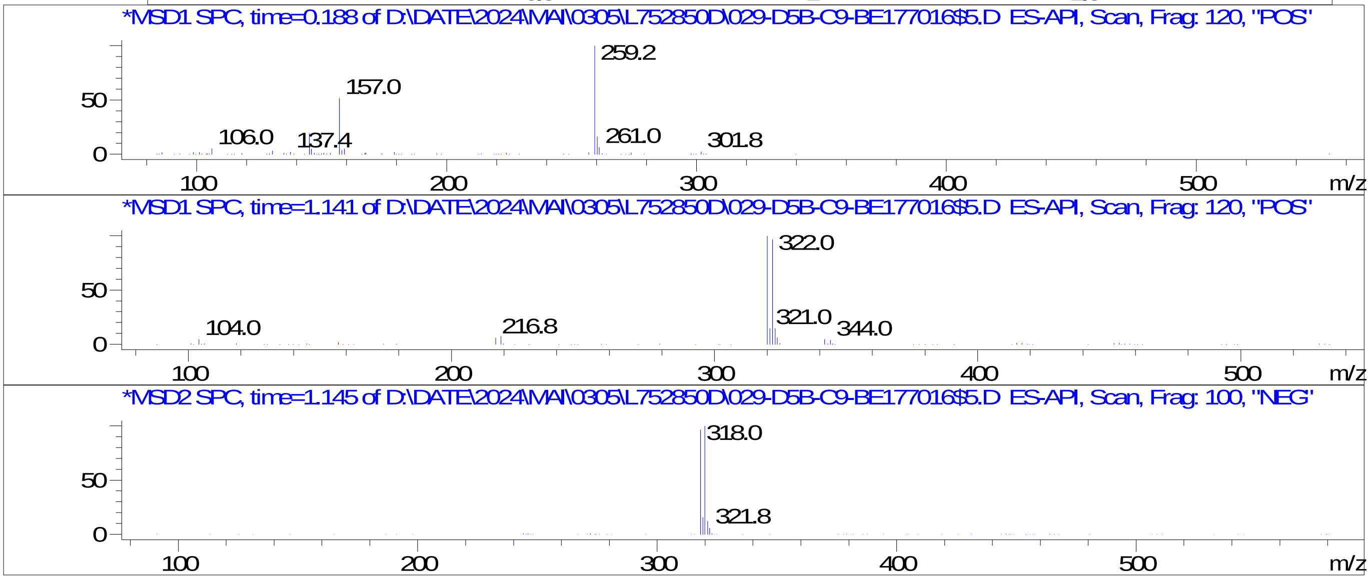


Inj.Date 5/2/2024 AN 
C:\Users\Public\Documents\ChemStation\1\Data\05_02\L752850D\SUPOR_30.M
