## Supplementary Data 13 for "SyntheMol-RL: a flexible reinforcement learning framework for designing novel and synthesizable antibiotics": Z5887248823.PDF

MaxPeak: 99.46%
Ret_Time: 1.156 min

# BD252513$1

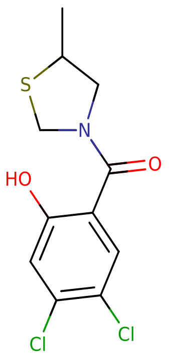


**Mol Wt** 292.18

**Exact Mass** 291

| # | Time | Area% |
|---|---|---|
| 1 | 0.967 | 0.54 |
| 2 | 1.156 | 99.46 |

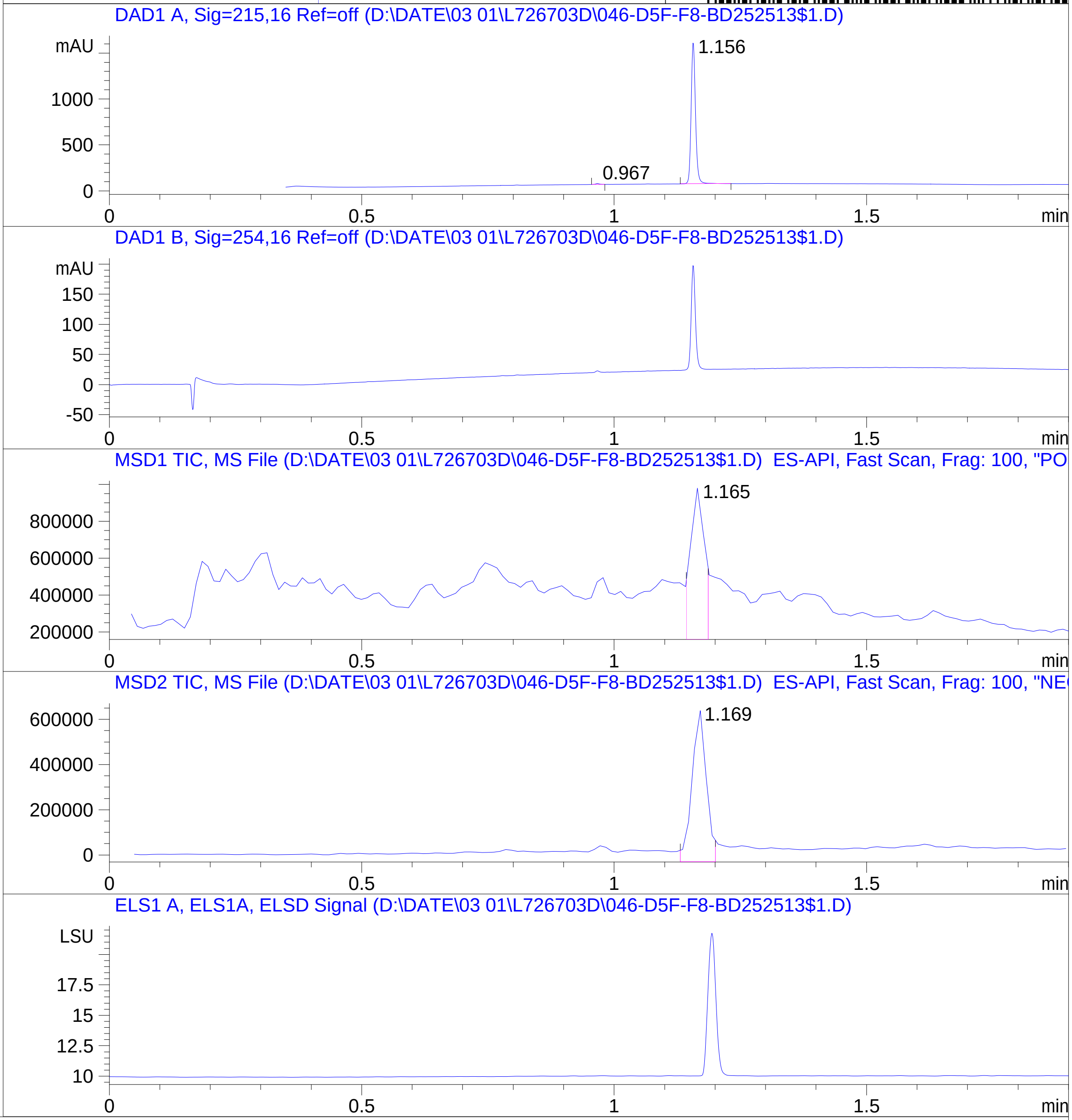


RT 1.165

RT 1.169

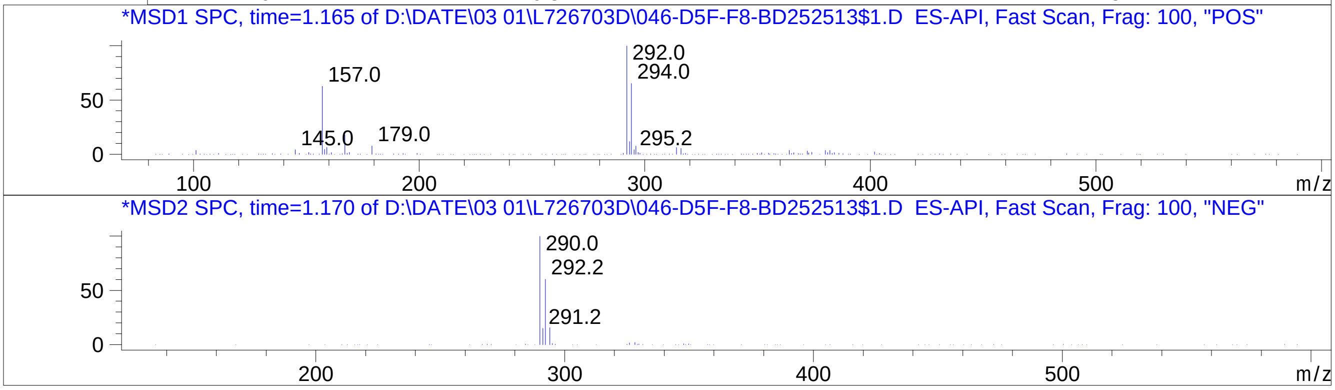


Inj.Date 3/1/2024 E
C:\Users\Public\Documents\ChemStation\1\Data\03_01\L726703D\SUPOR_30.M
