## Supplementary Data 13 for "SyntheMol-RL: a flexible reinforcement learning framework for designing novel and synthesizable antibiotics": Z8031540398.PDF

MaxPeak: 100.00%
Ret_Time: 1.053 min

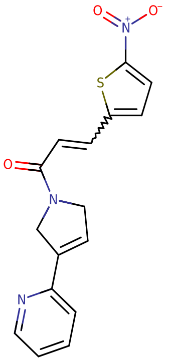


# T8998986

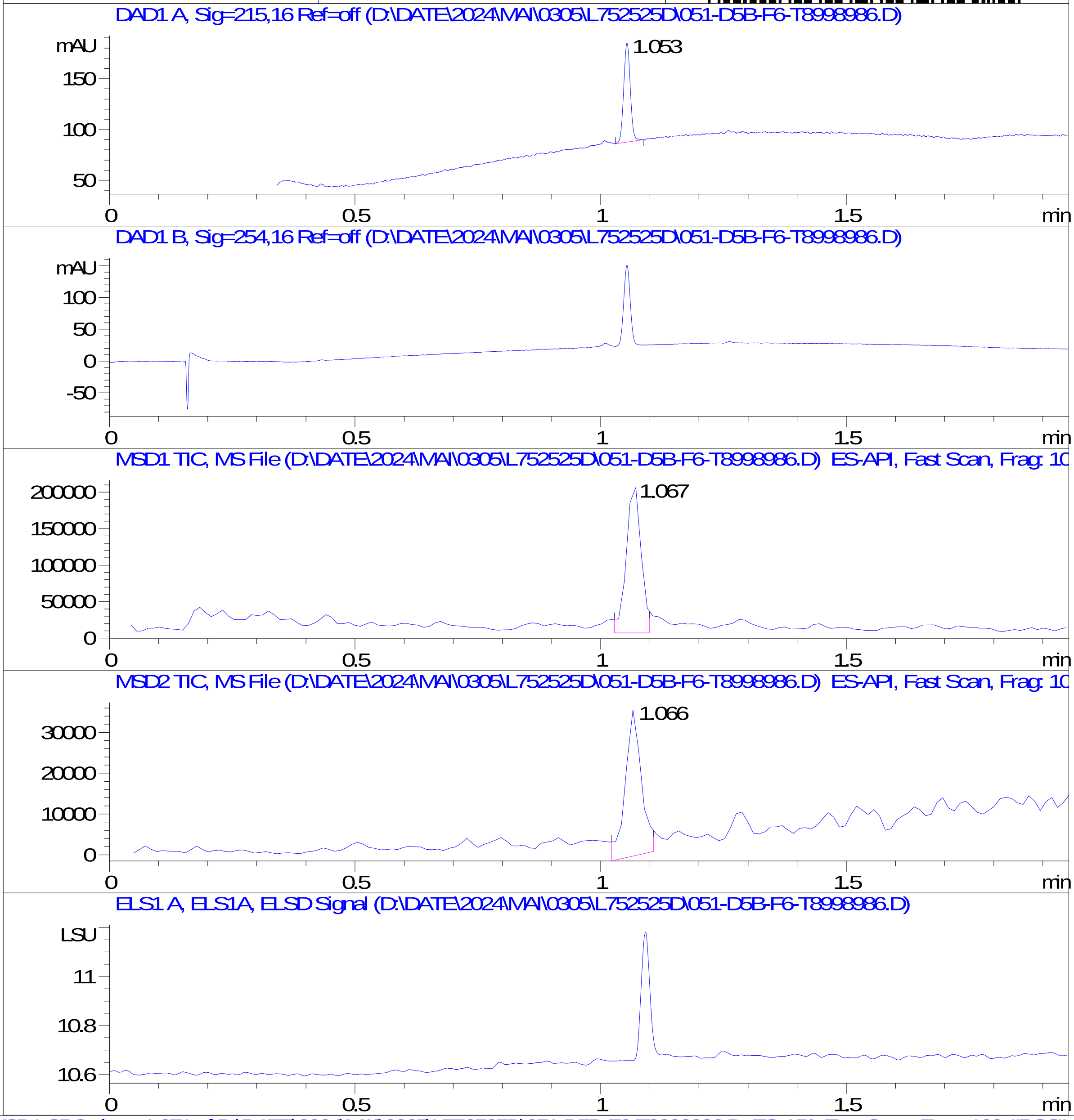


**Mol Wt** 327.36

**Exact Mass** 327.07

```
#  Time   Area%
-----------------
1  1.053  100.00
```

RT 1.067

RT 1.066

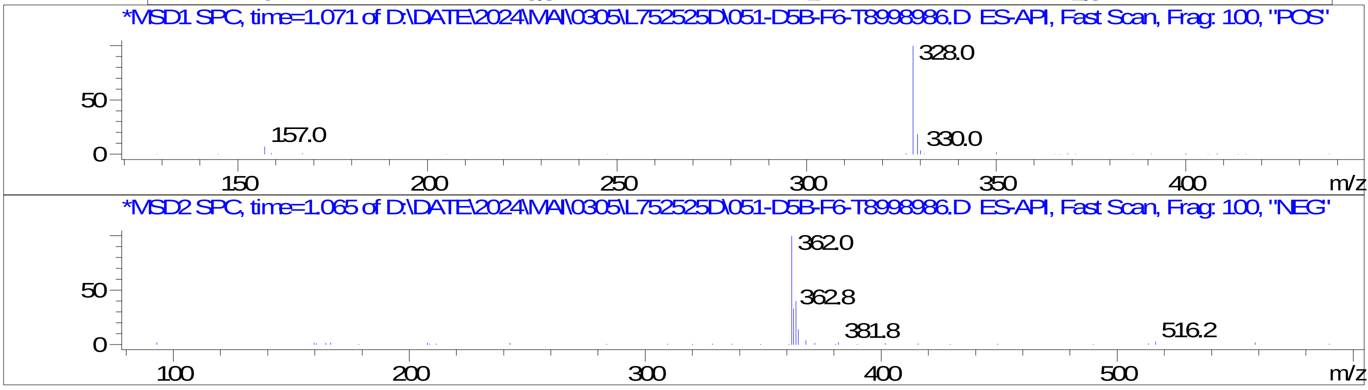


Inj.Date 5/2/2024 AN -23-
C:\Users\Public\Documents\ChemStation\1\Data\05_02\L752525D\SUPOR_30.M
