## Supplementary Data 13 for "SyntheMol-RL: a flexible reinforcement learning framework for designing novel and synthesizable antibiotics": Z8921960434.PDF

# BD252785$3

MaxPeak: 100.00%
Ret_Time: 0.478 min

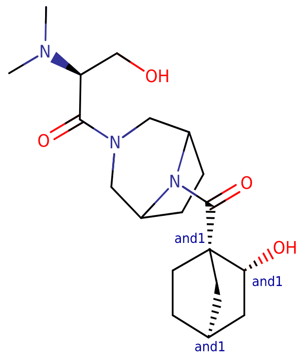


**Mol Wt** 365.47

**Exact Mass** 365.27

| # | Time | Area% |
| --- | --- | --- |
| 1 | 0.478 | 100.00 |

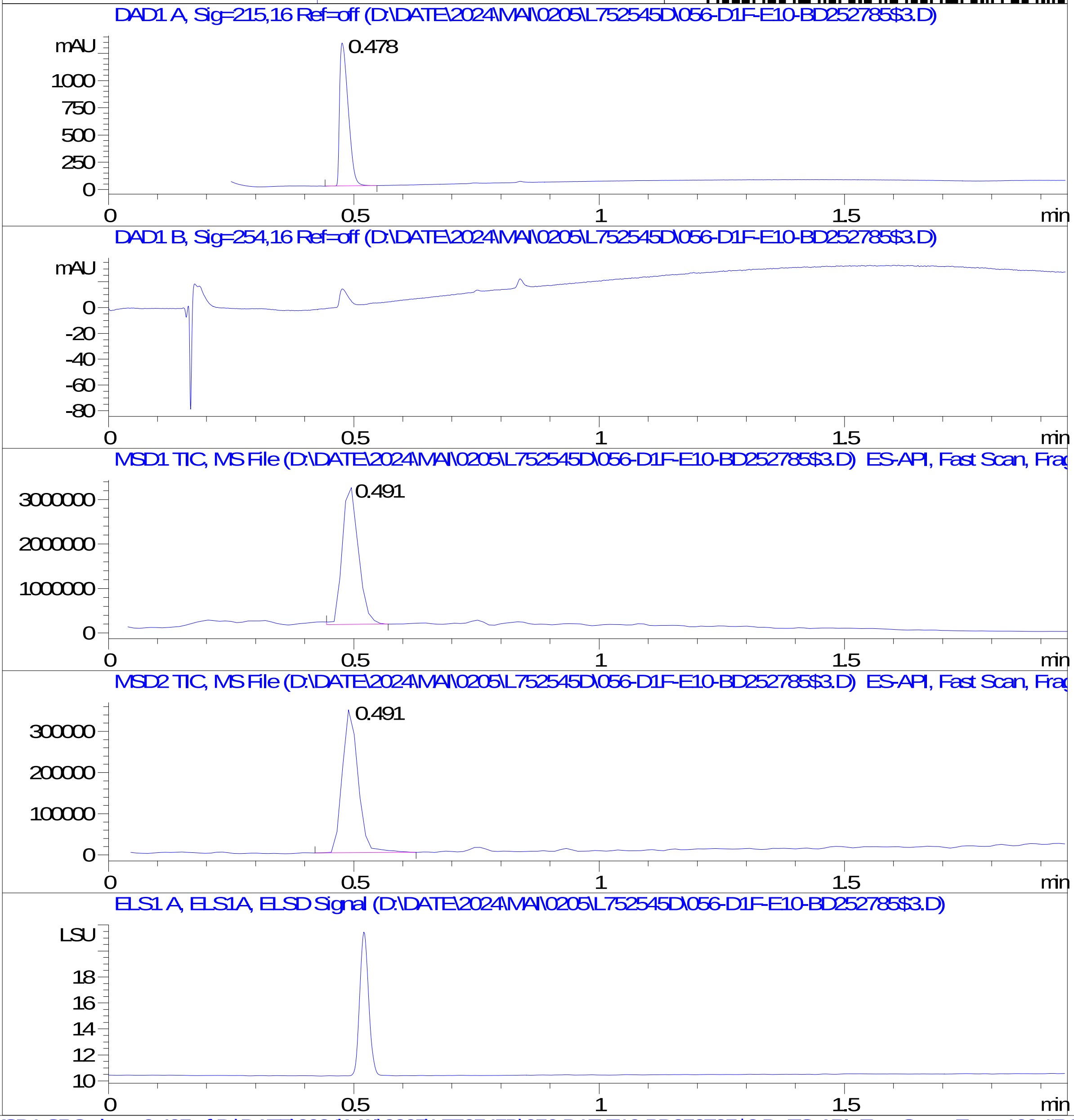


RT 0.491

RT 0.491

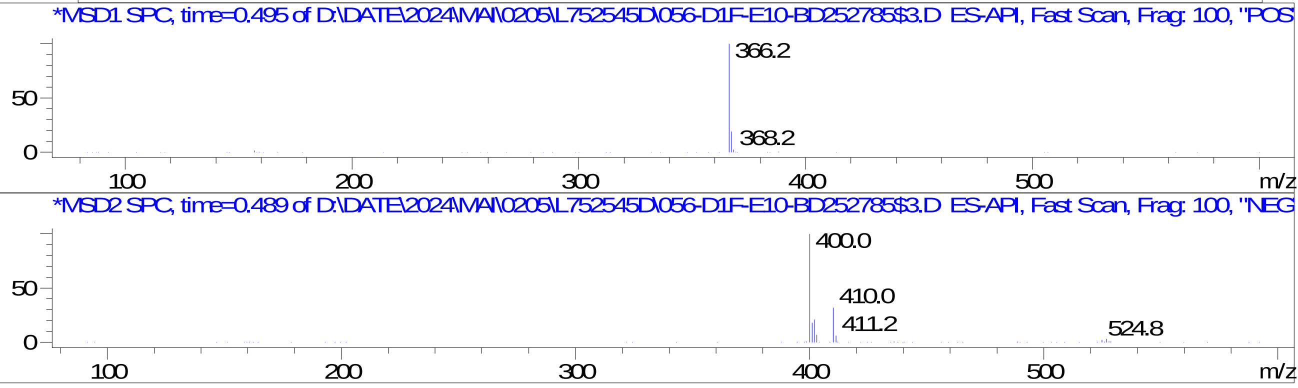


Inj.Date 5/2/2024 AN
C:\Users\Public\Documents\ChemStation\1\Data\05_02\L752545D\SUPOR_96.M
