## Supplementary Data 13 for "SyntheMol-RL: a flexible reinforcement learning framework for designing novel and synthesizable antibiotics": Z8921960447.PDF

MaxPeak: 98.02%
Ret_Time: 0.607 min

# BD252507$2

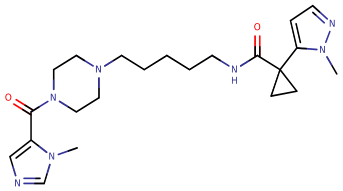


**Mol Wt** 427.54

**Exact Mass** 427.31

| # | Time | Area% |
|---|---|---|
| 1 | 0.607 | 98.02 |
| 2 | 1.282 | 1.98 |

DAD1 A, Sig=215,16 Ref=off (D:\DATE\0502\L752587D\066-D1B-H1-BD252507$2.D)

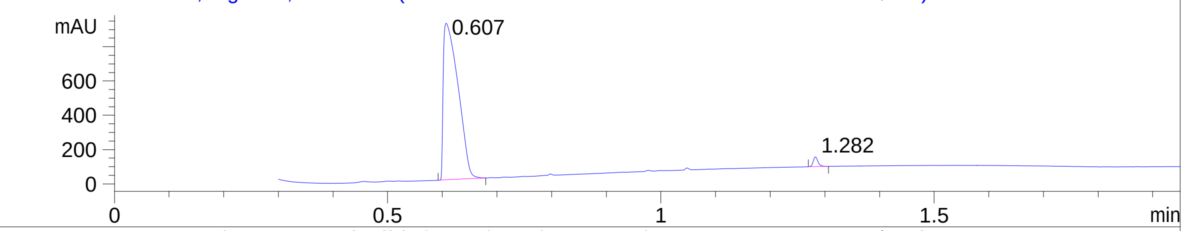


DAD1 B, Sig=254,16 Ref=off (D:\DATE\0502\L752587D\066-D1B-H1-BD252507$2.D)

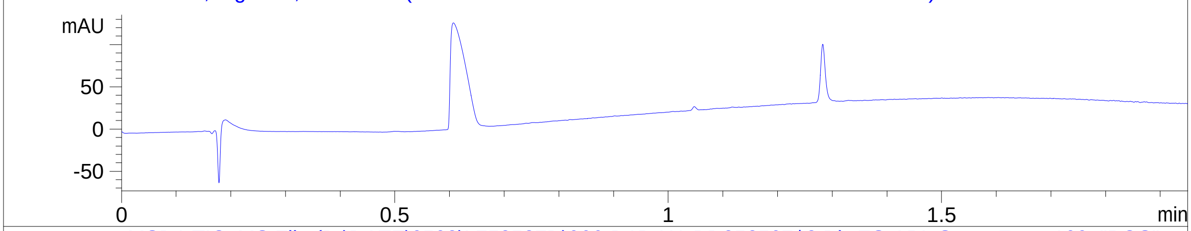


MSD1 TIC, MS File (D:\DATE\0502\L752587D\066-D1B-H1-BD252507$2.D) ES-API, Scan, Frag: 100, "POS"

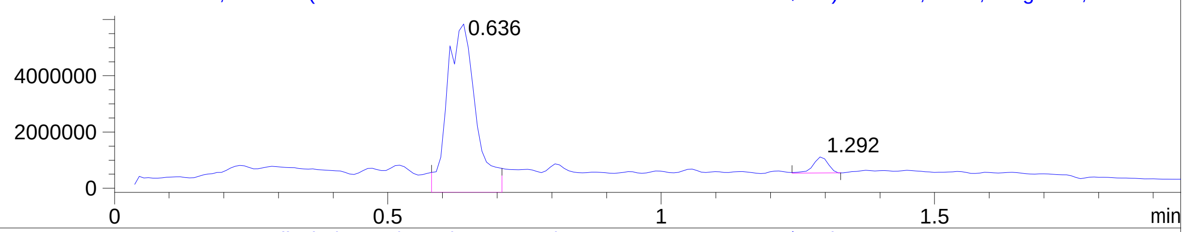


MSD2 TIC, MS File (D:\DATE\0502\L752587D\066-D1B-H1-BD252507$2.D) ES-API, Scan, Frag: 100, "NEG"

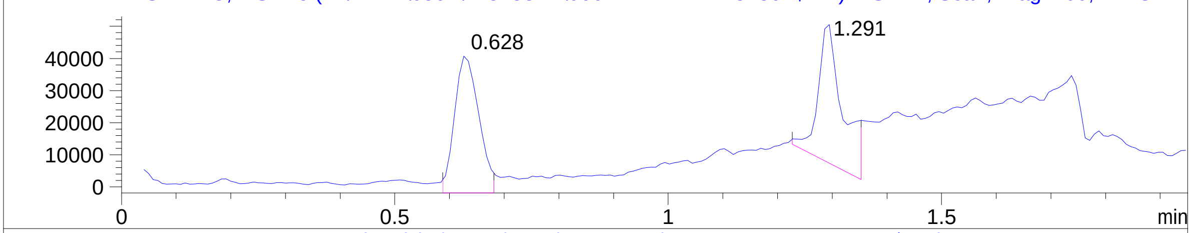


ELS1 A, ELS1A, ELSD Signal (D:\DATE\0502\L752587D\066-D1B-H1-BD252507$2.D)

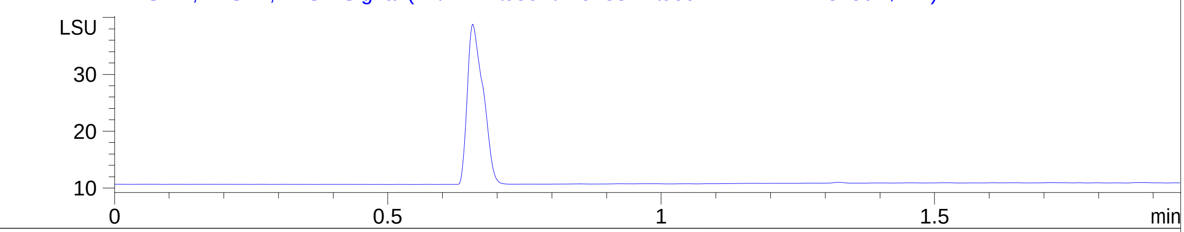


*MSD1 SPC, time=0.638 of D:\DATE\0502\L752587D\066-D1B-H1-BD252507$2.D ES-API, Scan, Frag: 100, "POS"

RT 0.636

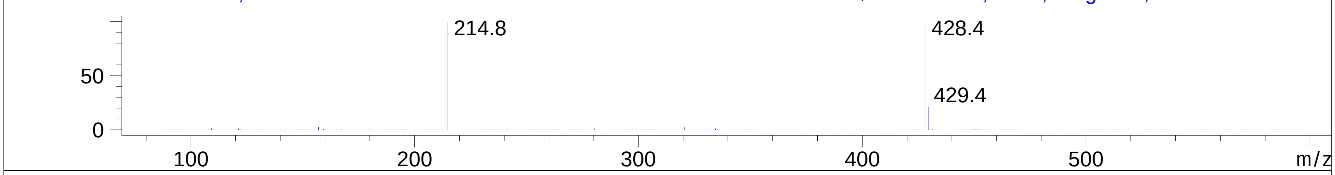


*MSD1 SPC, time=1.290 of D:\DATE\0502\L752587D\066-D1B-H1-BD252507$2.D ES-API, Scan, Frag: 100, "POS"

RT 1.292

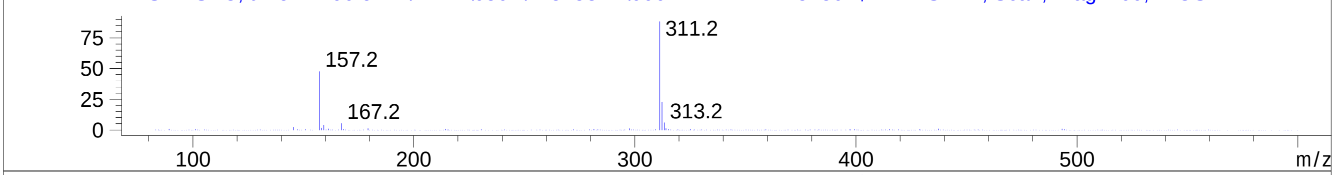


*MSD2 SPC, time=0.626 of D:\DATE\0502\L752587D\066-D1B-H1-BD252507$2.D ES-API, Scan, Frag: 100, "NEG"

RT 0.628

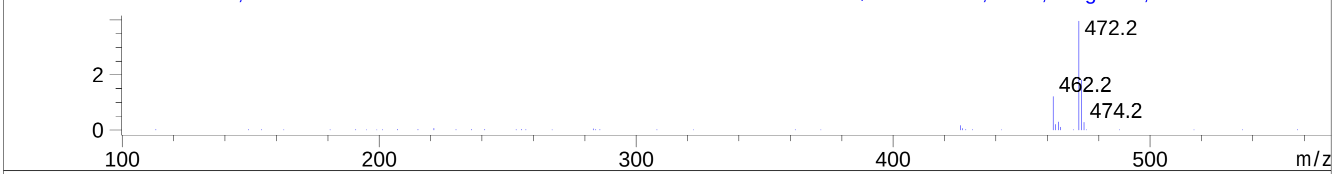


*MSD2 SPC, time=1.294 of D:\DATE\0502\L752587D\066-D1B-H1-BD252507$2.D ES-API, Scan, Frag: 100, "NEG"

RT 1.291

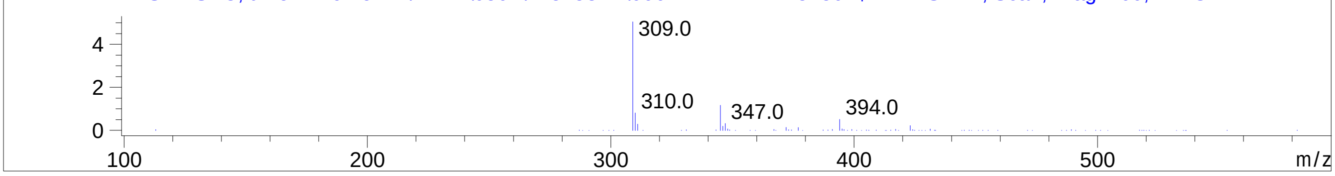
