## Supplementary Data 13 for "SyntheMol-RL: a flexible reinforcement learning framework for designing novel and synthesizable antibiotics": Z8921960455.PDF

# BD252787$2

MaxPeak: 100.00%
Ret_Time: 0.884 min

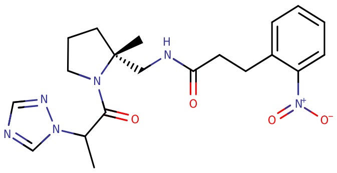


**Mol Wt** 414.46

**Exact Mass** 414.22

| # | Time | Area% |
|---|---|---|
| 1 | 0.884 | 100.00 |

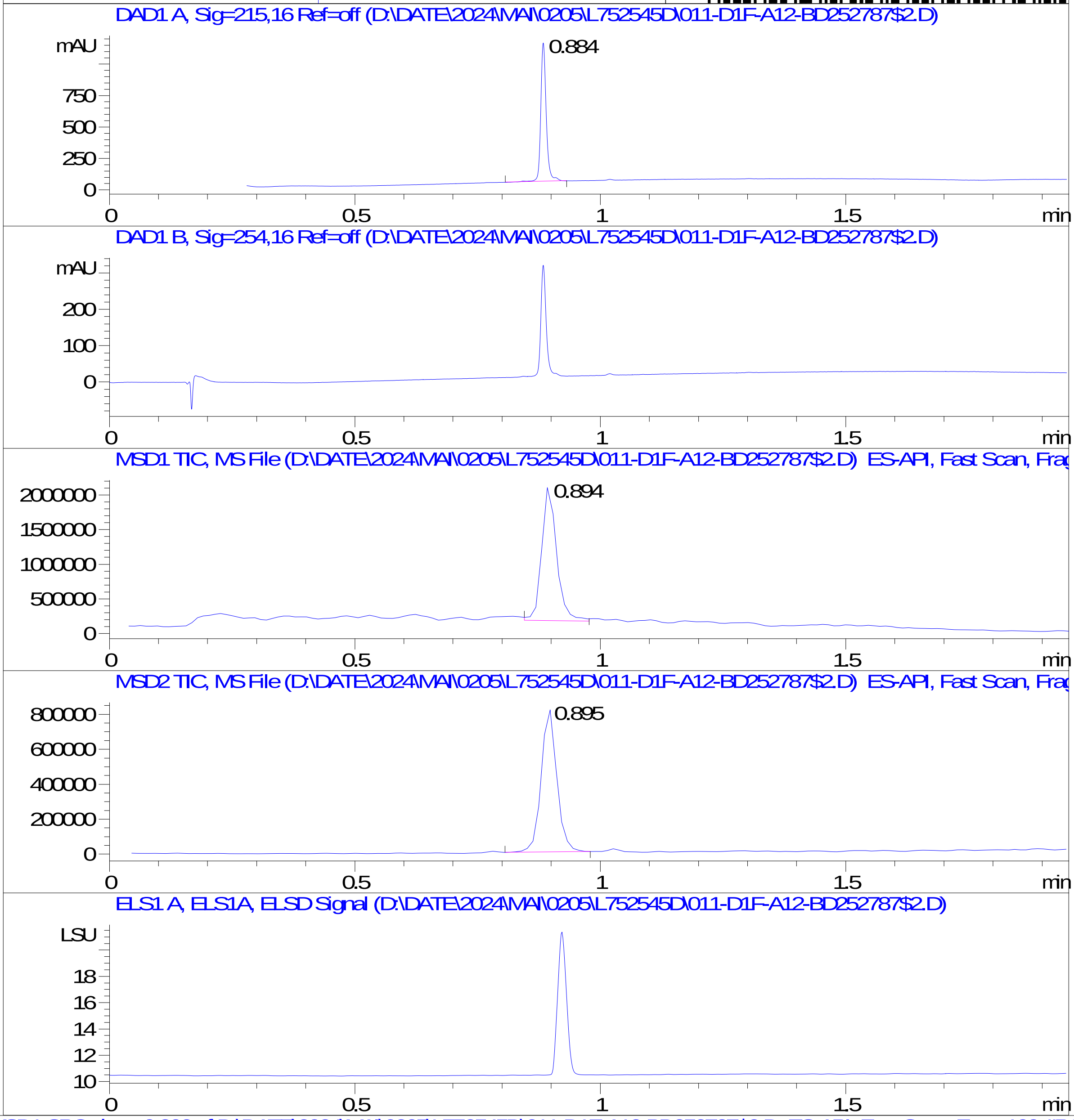


RT 0.894

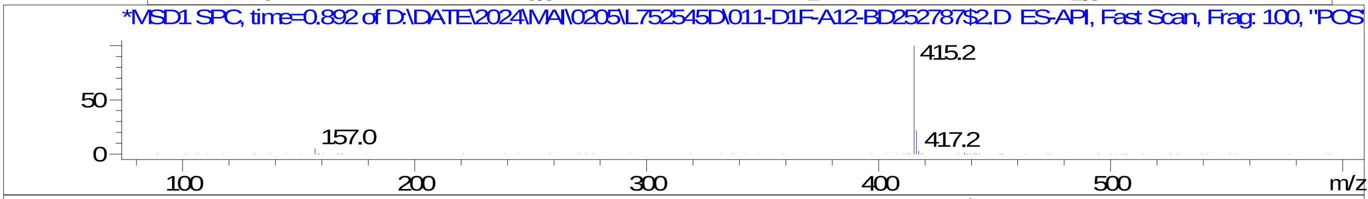


RT 0.895

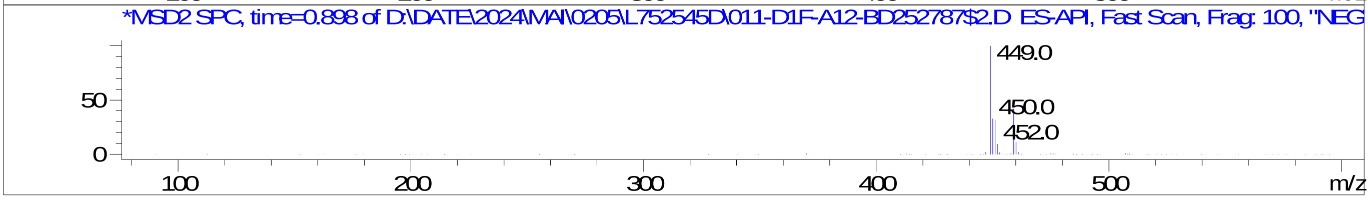


Inj.Date 5/2/2024 AN
C:\Users\Public\Documents\ChemStation\1\Data\05_02\L752545D\SUPOR_96.M
