## Supplementary Data 13 for "SyntheMol-RL: a flexible reinforcement learning framework for designing novel and synthesizable antibiotics": Z8921960482.PDF

BD252523$3

MaxPeak: 100.00%
Ret_Time: 1.145 min

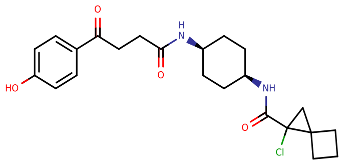


Mol Wt 432.94

Exact Mass 432.22

| # | Time | Area% |
|---|---|---|
| 1 | 1.145 | 100.00 |

DAD1 A, Sig=215,16 Ref=off (D:\DATE\03 01\L726419D\SAMPL000017.D)

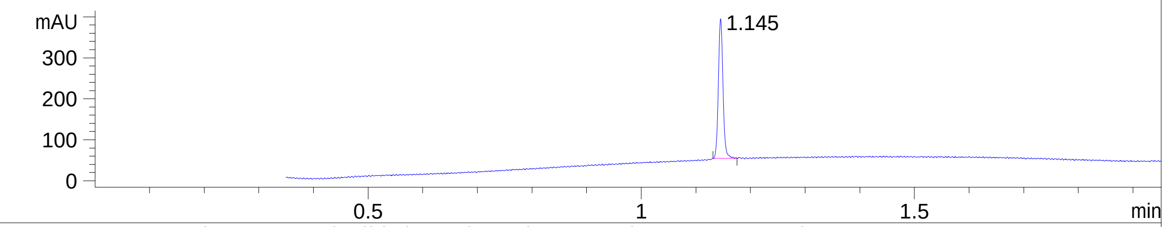


DAD1 B, Sig=254,16 Ref=off (D:\DATE\03 01\L726419D\SAMPL000017.D)

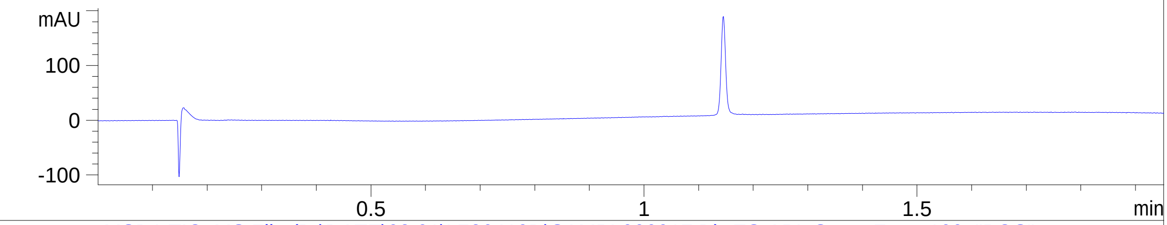


MSD1 TIC, MS File (D:\DATE\03 01\L726419D\SAMPL000017.D) ES-API, Scan, Frag: 100, "POS"

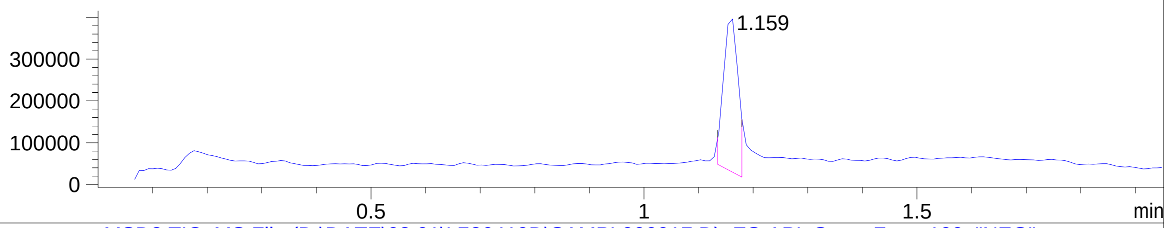


MSD2 TIC, MS File (D:\DATE\03 01\L726419D\SAMPL000017.D) ES-API, Scan, Frag: 100, "NEG"

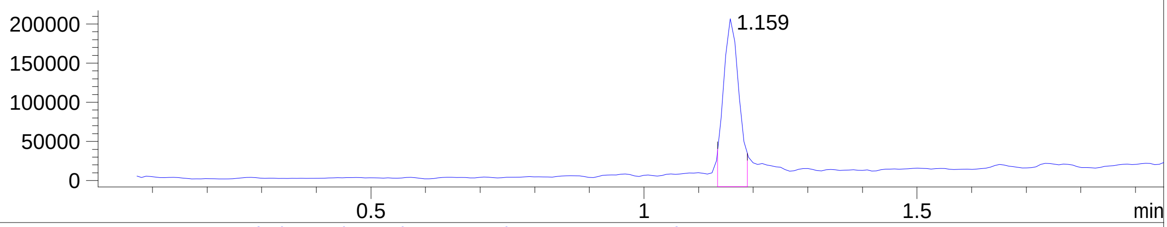


ADC1 A, ELSD (D:\DATE\03 01\L726419D\SAMPL000017.D)

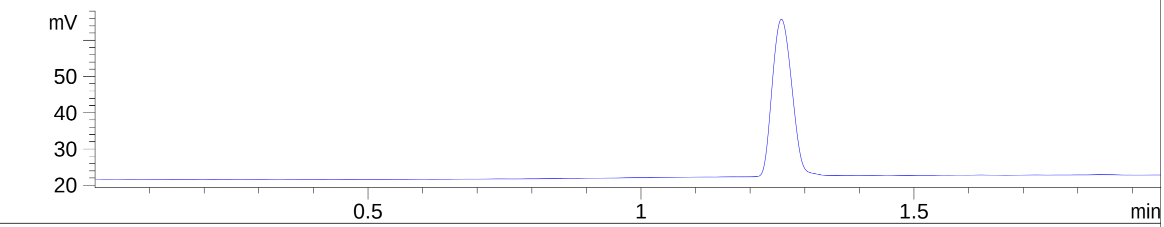


RT 1.159

*MSD1 SPC, time=1.162 of D:\DATE\03 01\L726419D\SAMPL000017.D ES-API, Scan, Frag: 100, "POS"

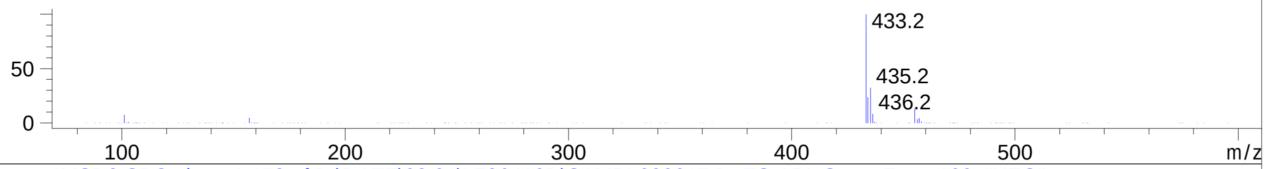


RT 1.159

*MSD2 SPC, time=1.158 of D:\DATE\03 01\L726419D\SAMPL000017.D ES-API, Scan, Frag: 100, "NEG"

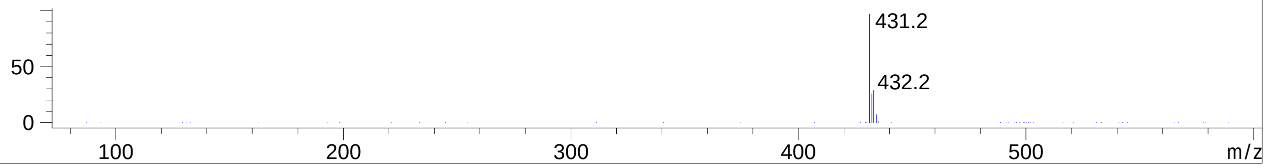
