## Supplementary Data 13 for "SyntheMol-RL: a flexible reinforcement learning framework for designing novel and synthesizable antibiotics": Z8921960485.PDF

MaxPeak:  95.30%
Ret_Time: 1.070 min

# BD252746$1

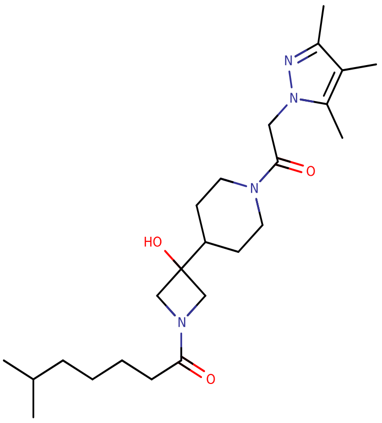


**Mol Wt** 432.6

**Exact Mass** 432.37

| # | Time | Area% |
|---|---|---|
| 1 | 0.845 | 4.70 |
| 2 | 1.070 | 95.30 |

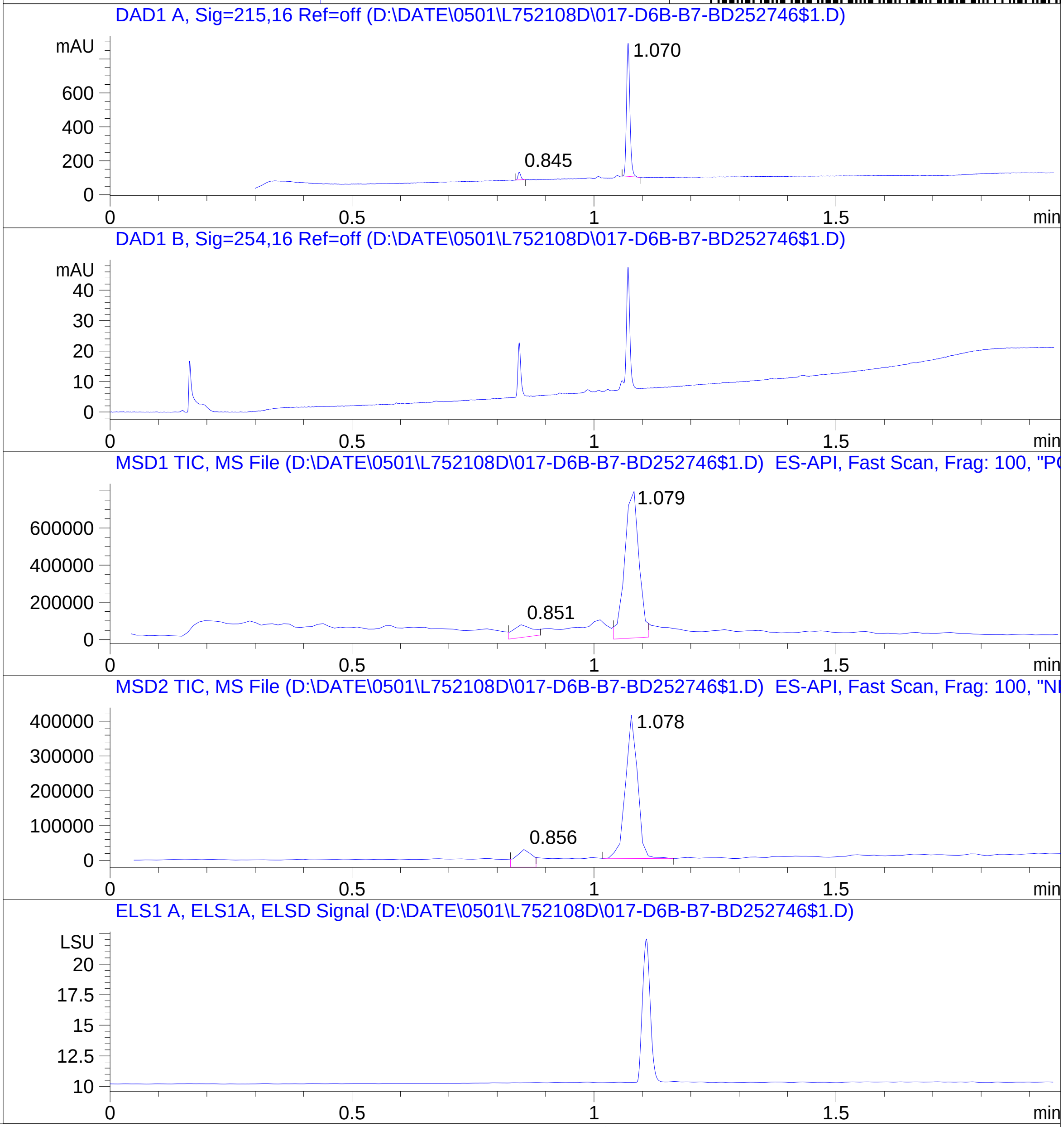


RT 0.851

RT 1.079

RT 0.856

RT 1.078

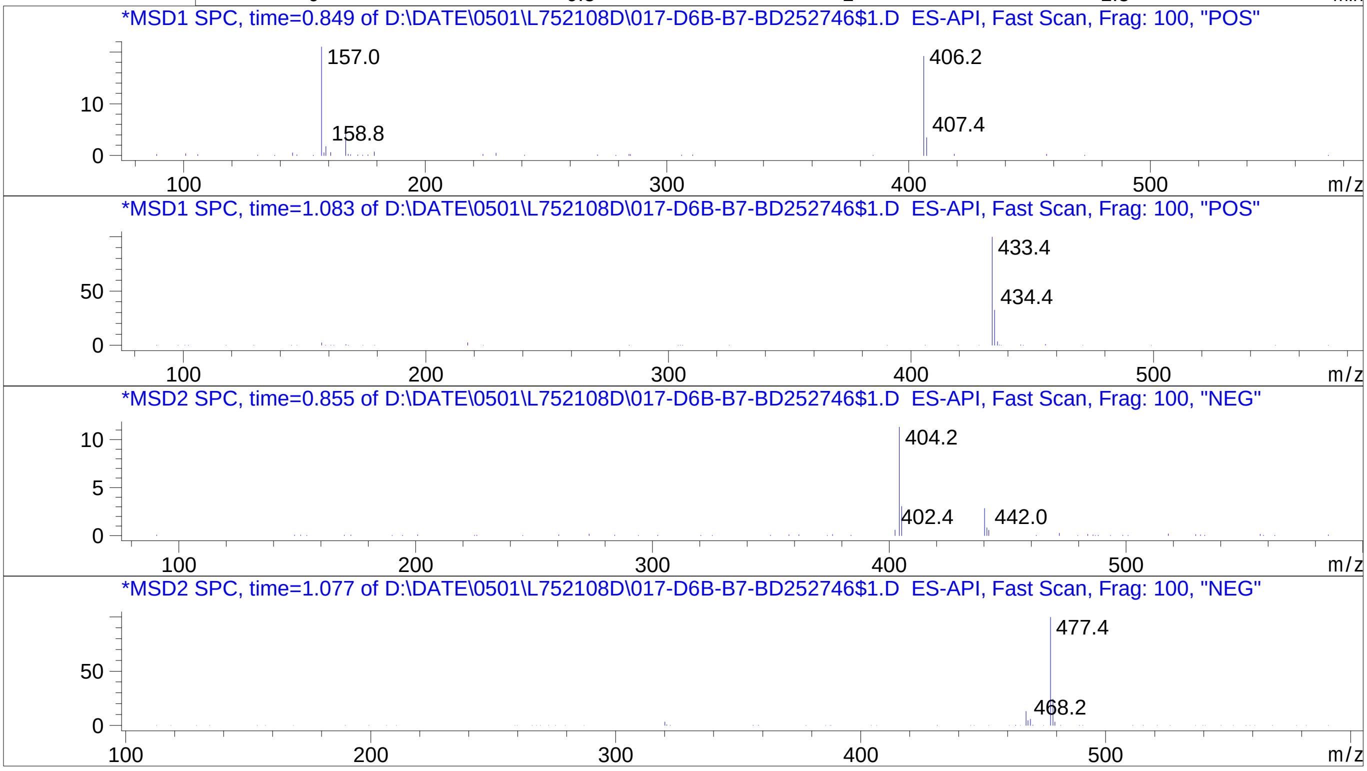


Inj.Date 5/1/2024 0 32
C:\Users\Public\Documents\ChemStation\1\Data\05_01\L752108D\SUPOR_30.M
