## Supplementary Data 13 for "SyntheMol-RL: a flexible reinforcement learning framework for designing novel and synthesizable antibiotics": Z8921960492.PDF

MaxPeak: 48.76%
Ret_Time: 0.986 min

# BD252788$2

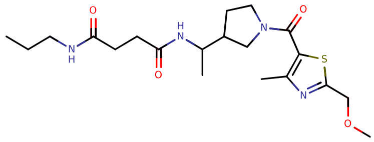

**Mol Wt** 424.56

**Exact Mass** 424.25

| # | Time | Area% |
|---|---|---|
| 1 | 0.974 | 47.86 |
| 2 | 0.986 | 48.76 |
| 3 | 1.021 | 3.39 |

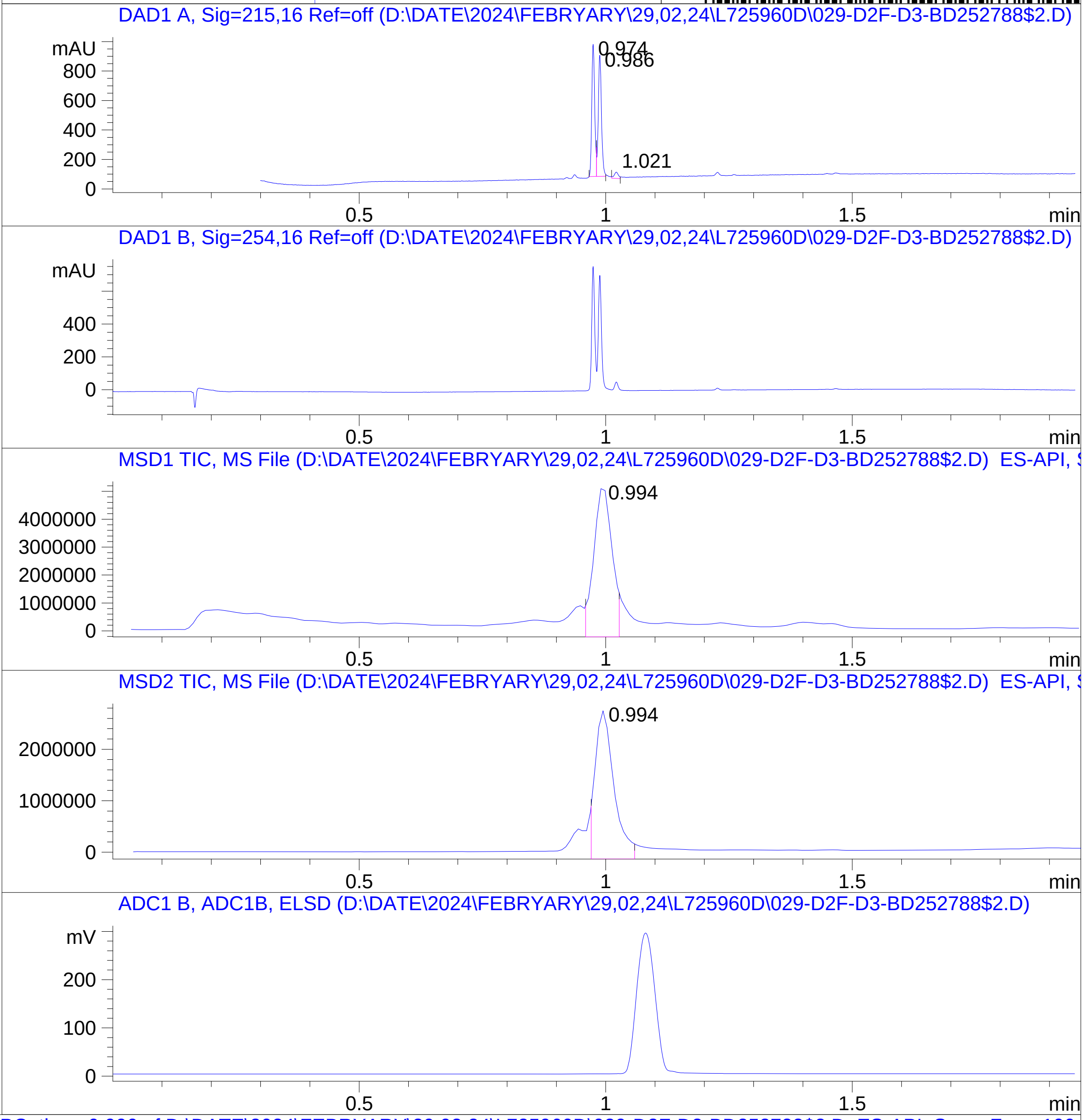


RT 0.994

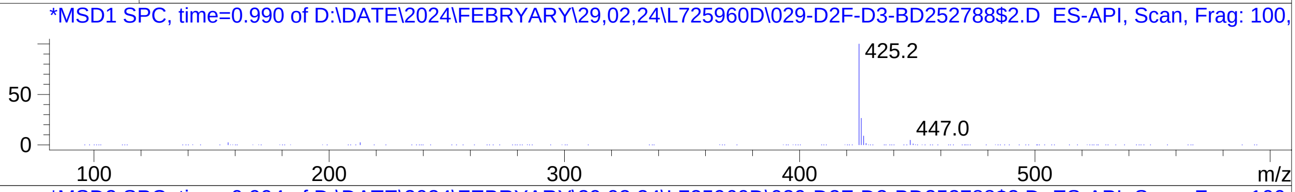


RT 0.994

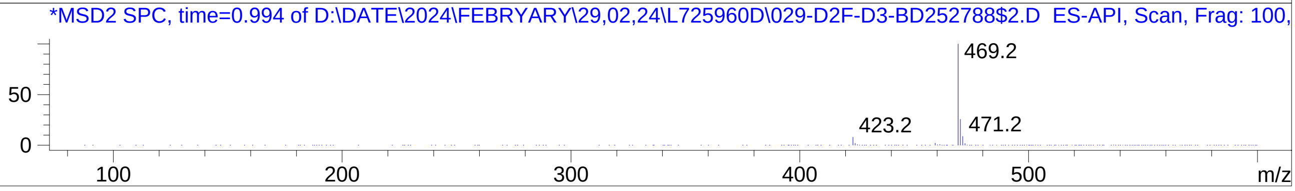


Inj.Date 2/29/2024 T -26- ->
C:\Users\Public\Documents\ChemStation\1\Data\02_29\L725960D\SUPOR_96.M
