## Supplementary Data 13 for "SyntheMol-RL: a flexible reinforcement learning framework for designing novel and synthesizable antibiotics": Z8921960494.PDF

MaxPeak: 95.19%
Ret_Time: 1.154 min

# BD252508$12

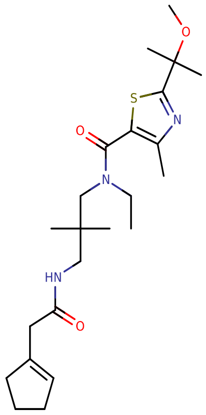

**Mol Wt** 435.62

**Exact Mass** 435.31

| # | Time | Area% |
|---|---|---|
| 1 | 0.875 | 3.27 |
| 2 | 1.068 | 1.54 |
| 3 | 1.154 | 95.19 |

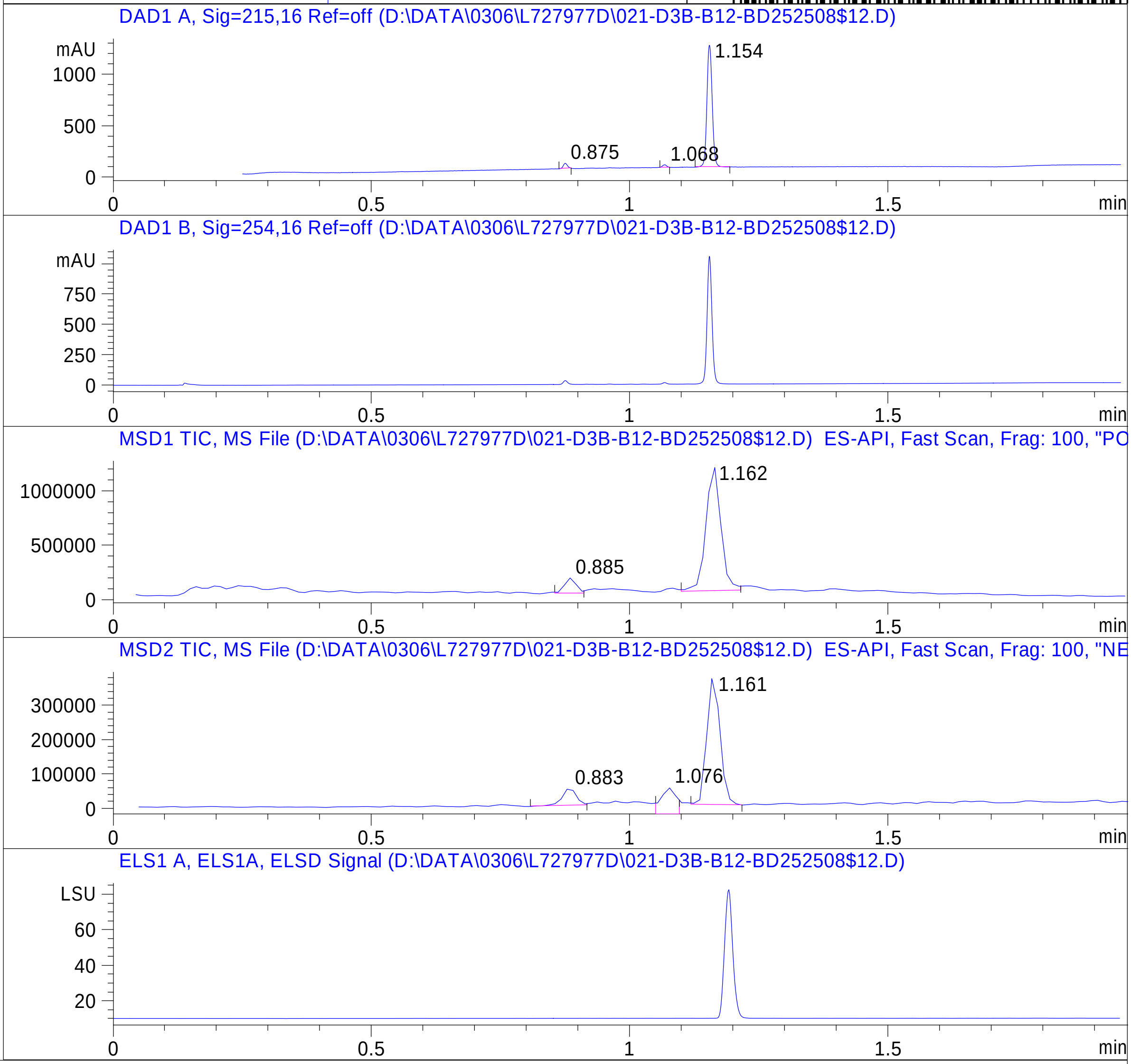


RT 0.885

RT 1.162

RT 0.883

RT 1.076

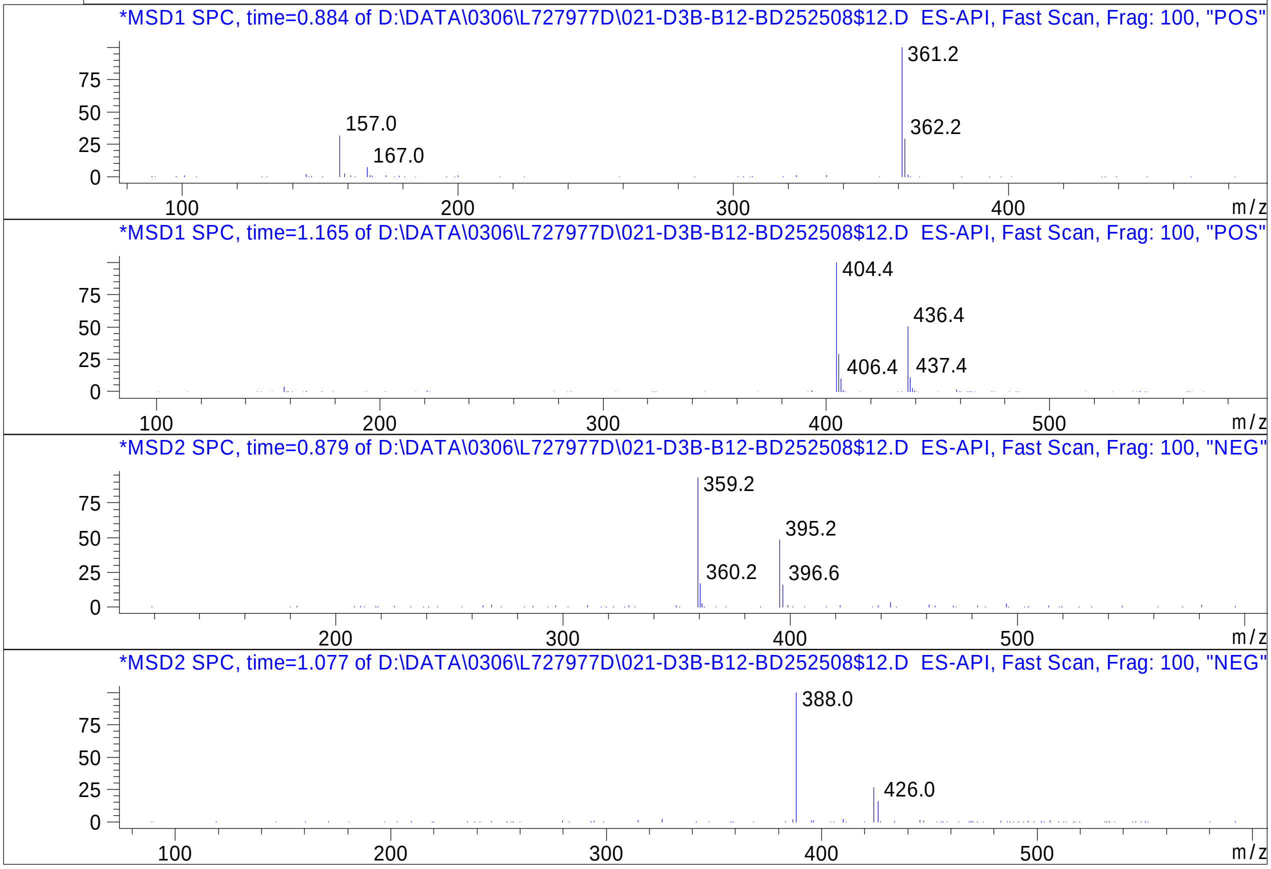


Inj.Date 3/6/2024 LB

C:\Users\Public\Documents\ChemStation\1\Data\03_05\L727977D\SUPOR_96.M

RT 1.161

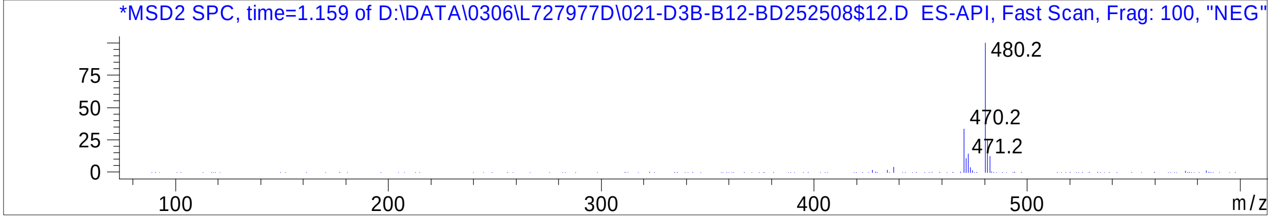


Inj.Date 3/6/2024 LB

C:\Users\Public\Documents\ChemStation\1\Data\03_05\L727977D\SUPOR_96.M
