## Supplementary Data 13 for "SyntheMol-RL: a flexible reinforcement learning framework for designing novel and synthesizable antibiotics": Z8921960511.PDF

MaxPeak: 96.10%
Ret_Time: 0.942 min

# BD252789$1

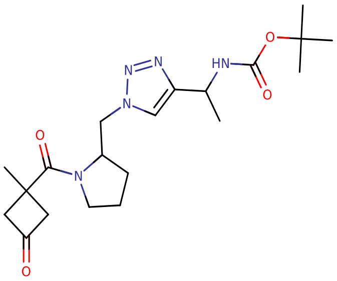

**Mol Wt** 405.49

**Exact Mass** 405.27

| # | Time | Area% |
|---|---|---|
| 1 | 0.853 | 3.90 |
| 2 | 0.942 | 96.10 |

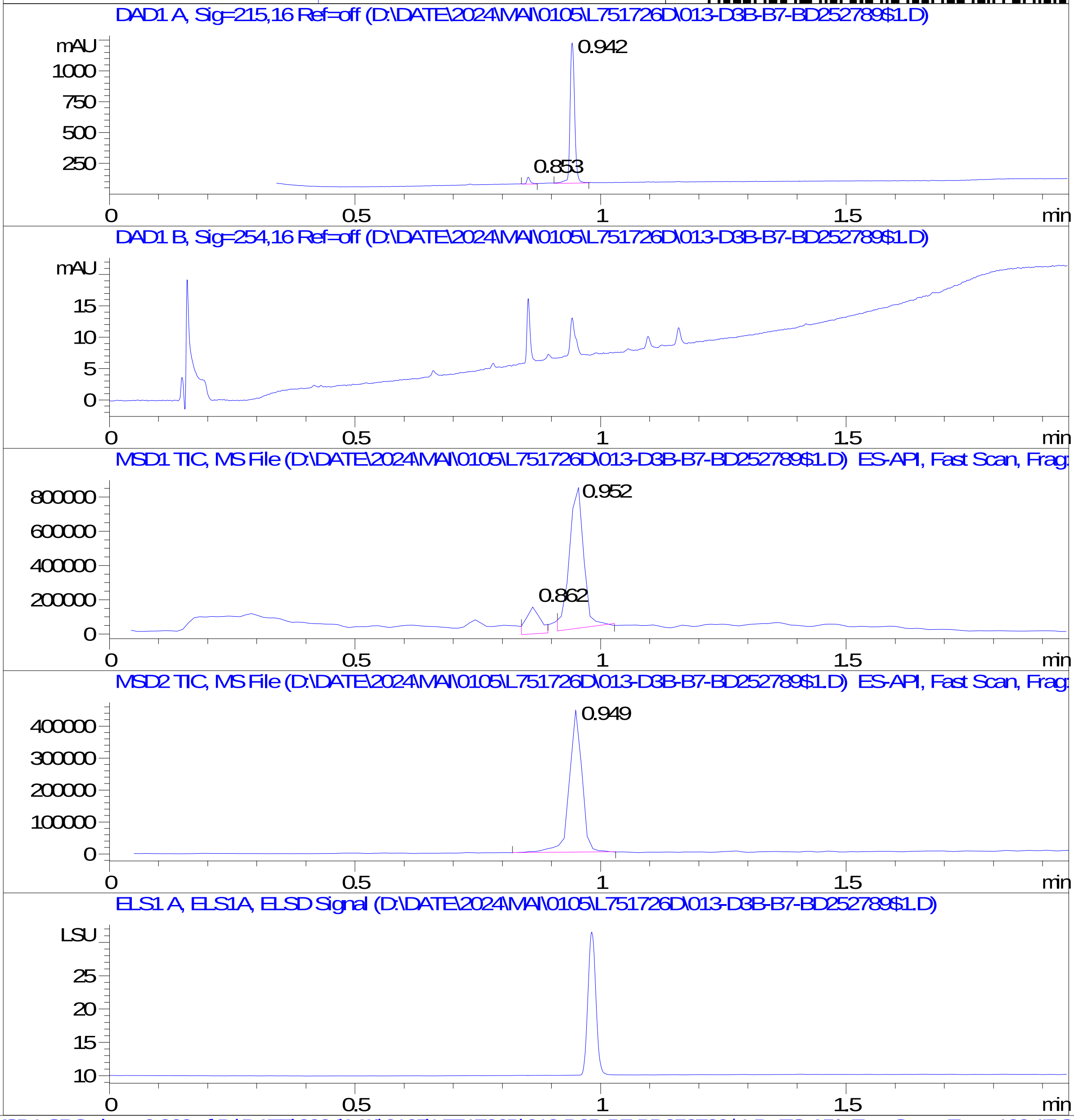


RT 0.862

RT 0.952

RT 0.949

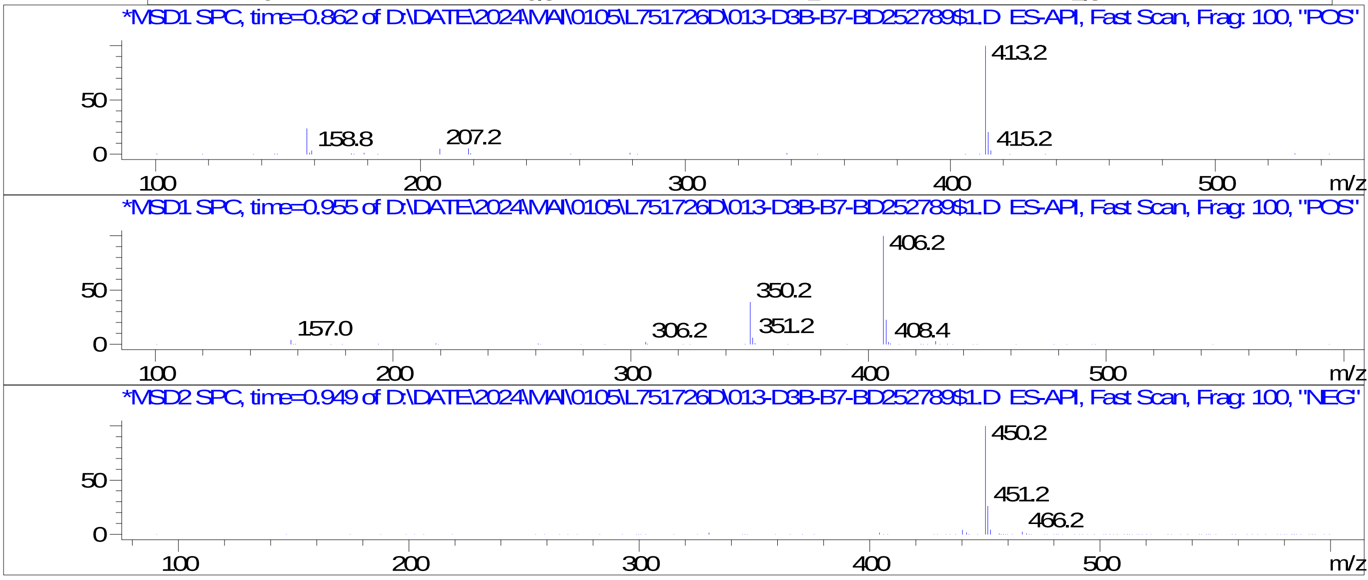


Inj.Date 4/30/2024 AN 34
C:\Users\Public\Documents\ChemStation\1\Data\04_30\L751726D\SUPOR_96.M
