## Supplementary Data 13 for "SyntheMol-RL: a flexible reinforcement learning framework for designing novel and synthesizable antibiotics": Z8921960594.PDF

MaxPeak: 100.00%
Ret_Time: 1.167 min

# BD252891$3

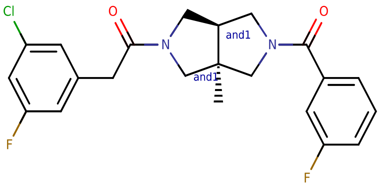


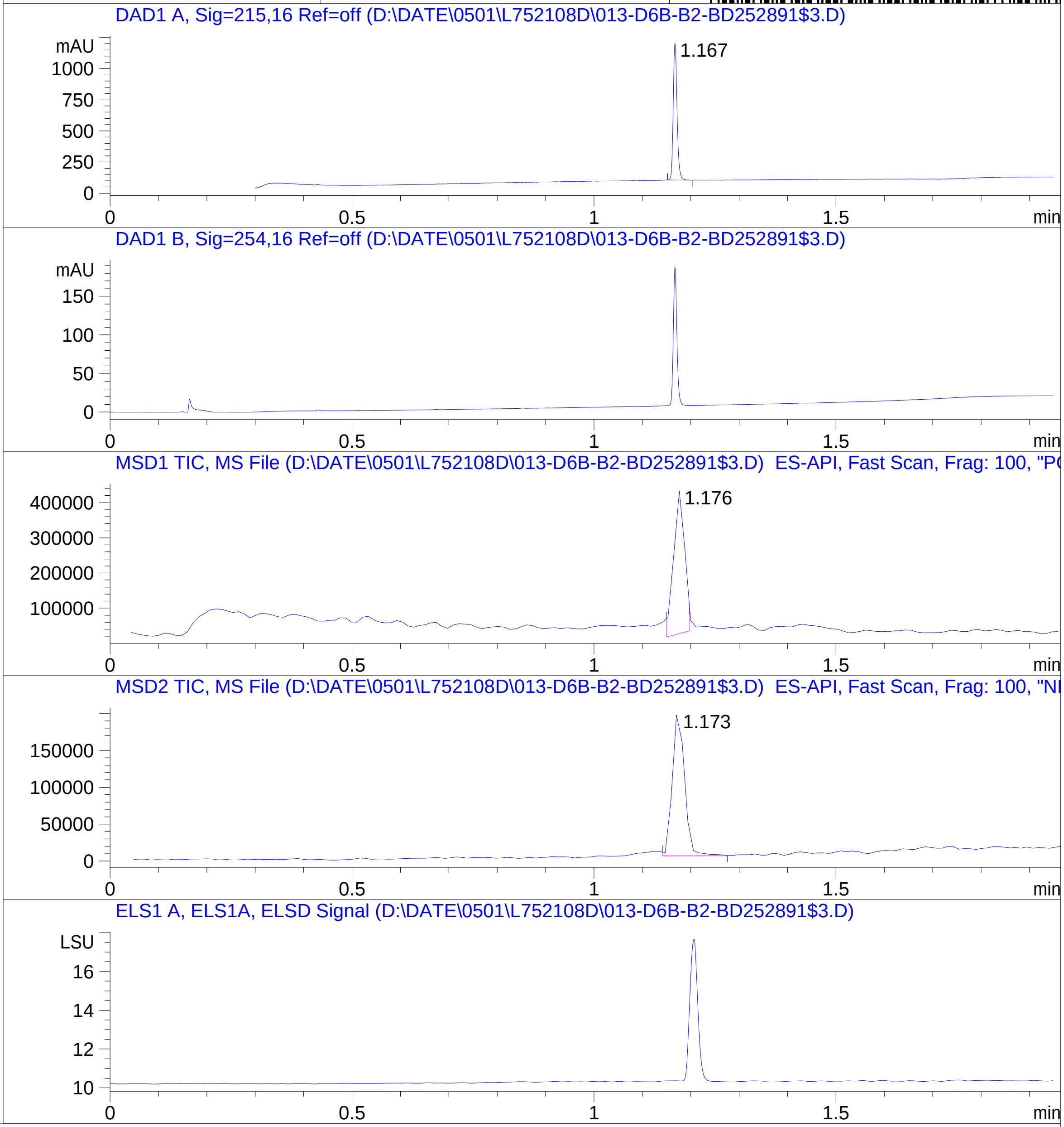


**Mol Wt** 418.86

**Exact Mass** 418.16

| # | Time | Area% |
|---|---|---|
| 1 | 1.167 | 100.00 |

RT 1.176

RT 1.173

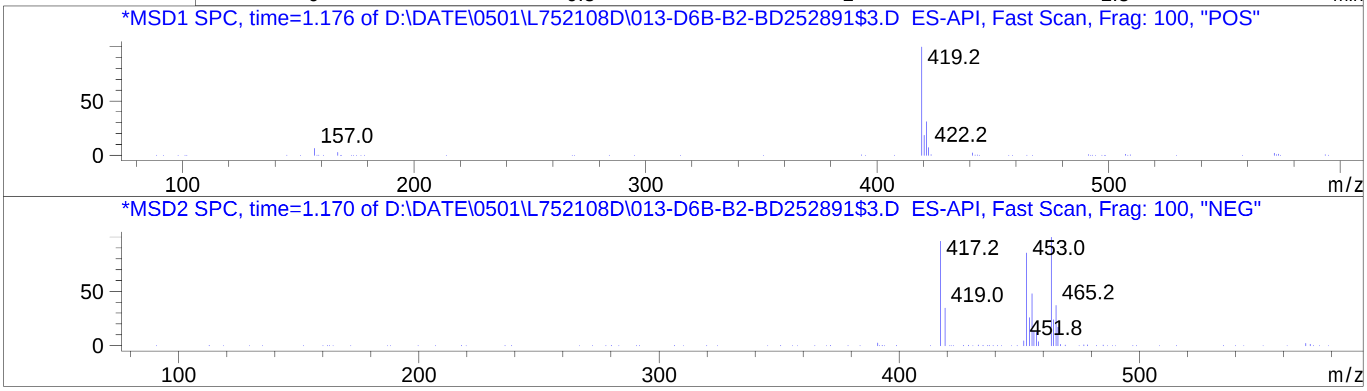


Inj.Date 5/1/2024 0 32
C:\Users\Public\Documents\ChemStation\1\Data\05_01\L752108D\SUPOR_30.M
