## Supplementary Data 13 for "SyntheMol-RL: a flexible reinforcement learning framework for designing novel and synthesizable antibiotics": Z8921960624.PDF

MaxPeak: 98.86%
Ret_Time: 1.019 min

# BD252752$2

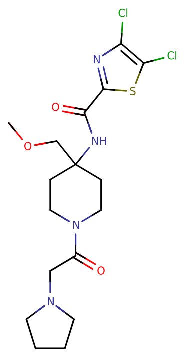


**Mol Wt** 435.37

**Exact Mass** 434.12

| # | Time | Area% |
|---|---|---|
| 1 | 1.019 | 98.86 |
| 2 | 1.072 | 1.14 |

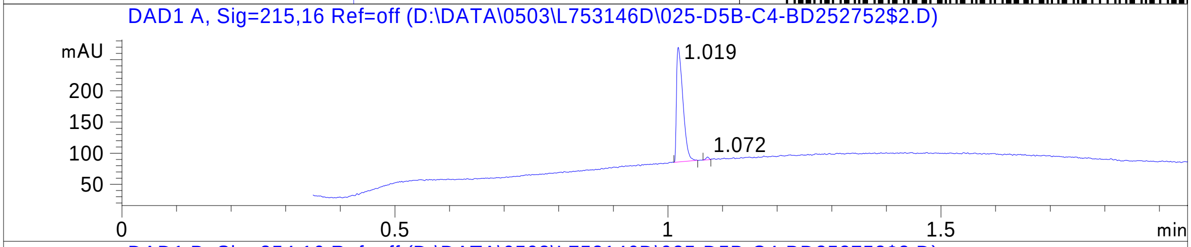


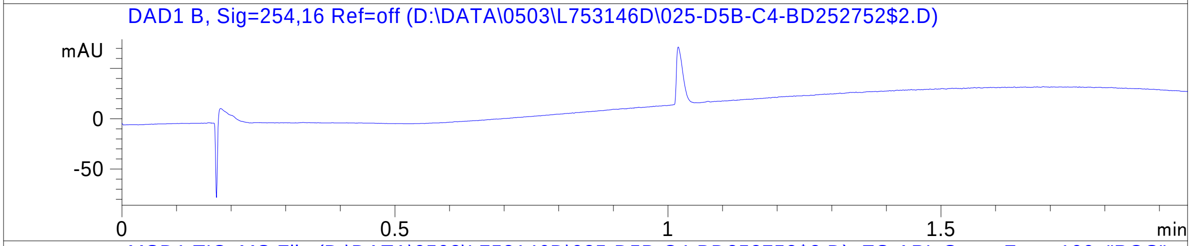


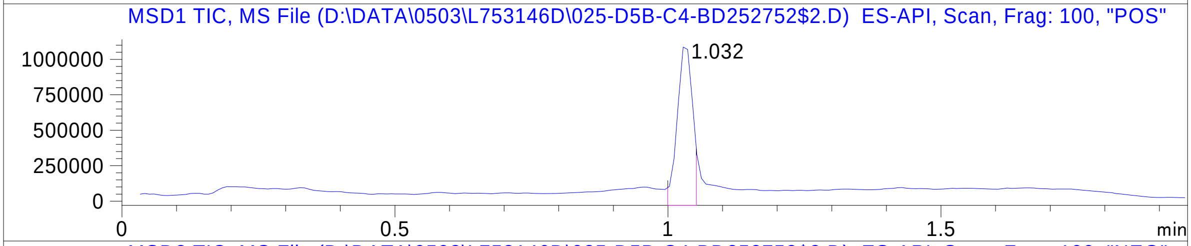


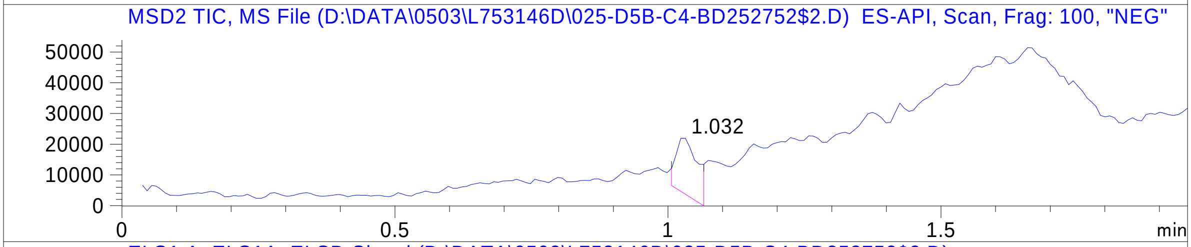


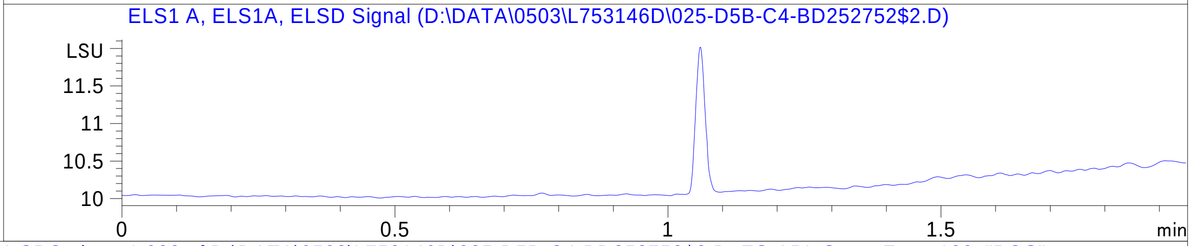


RT 1.032

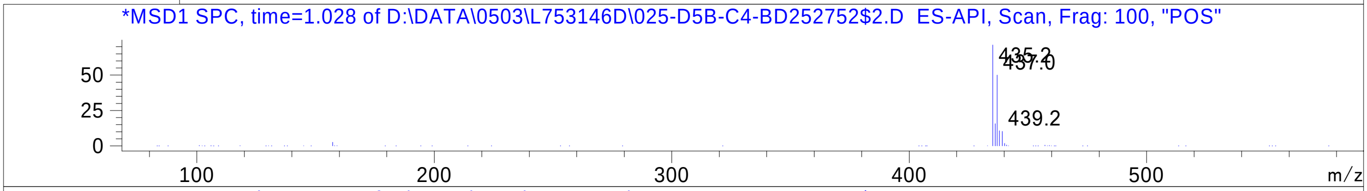


RT 1.032

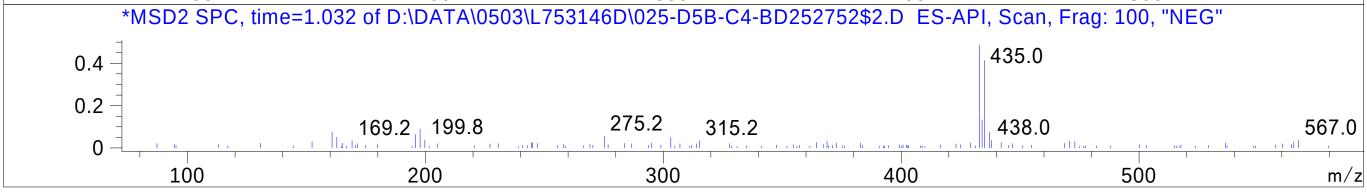


Inj.Date 5/3/2024 H 
C:\Users\Public\Documents\ChemStation\1\Data\05_03\L753146D\SUPOR_30.M
