## Supplementary Data 13 for "SyntheMol-RL: a flexible reinforcement learning framework for designing novel and synthesizable antibiotics": Z8921960672.PDF

# BD252967$3

MaxPeak:  68.75%
Ret_Time: 0.602 min

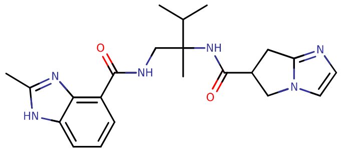

**Mol Wt** 408.5

**Exact Mass** 408.26

| # | Time | Area% |
|---|---|---|
| 1 | 0.490 | 4.97 |
| 2 | 0.602 | 68.75 |
| 3 | 0.646 | 26.27 |

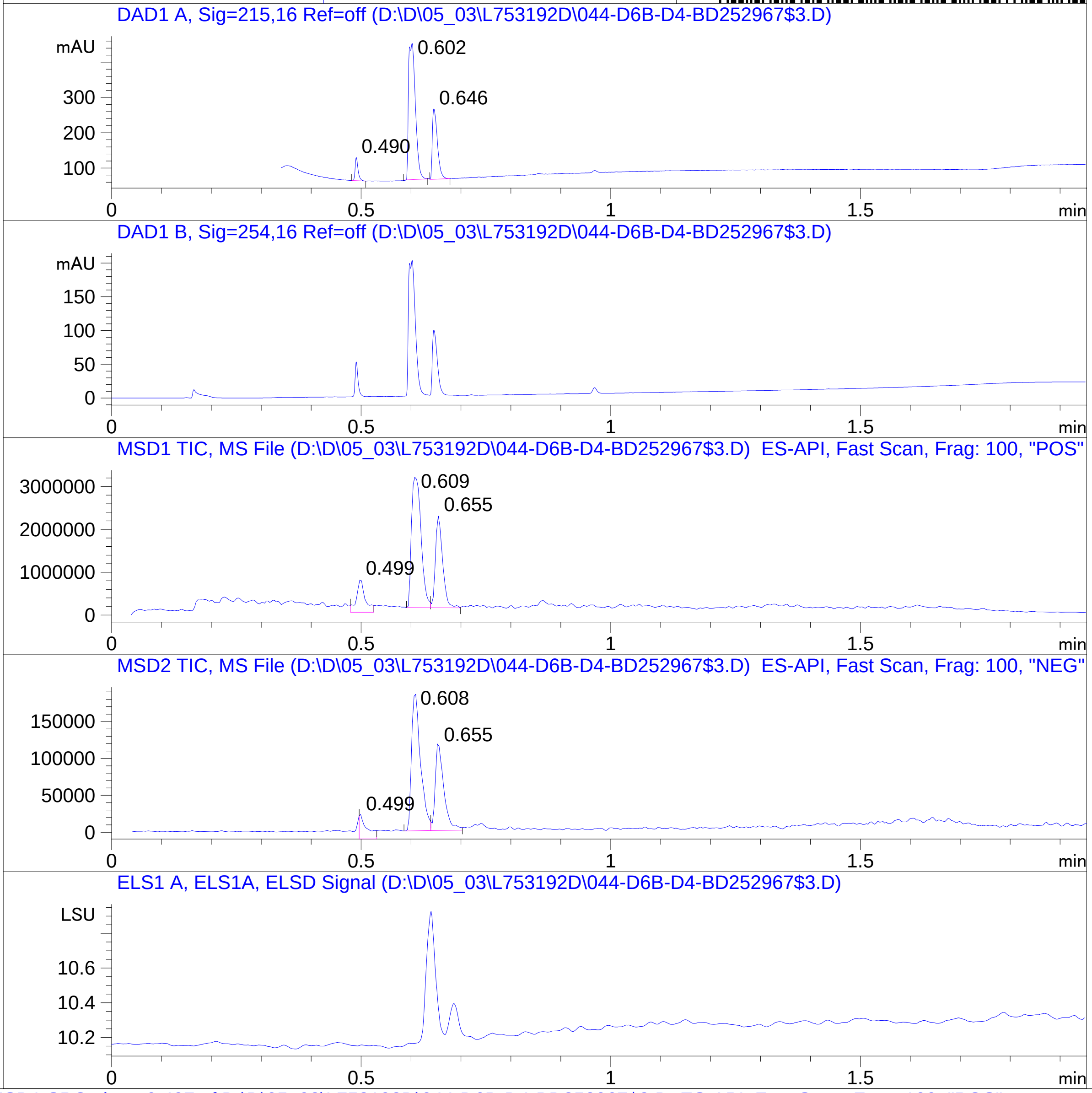


RT 0.499

RT 0.609

RT 0.655

RT 0.499

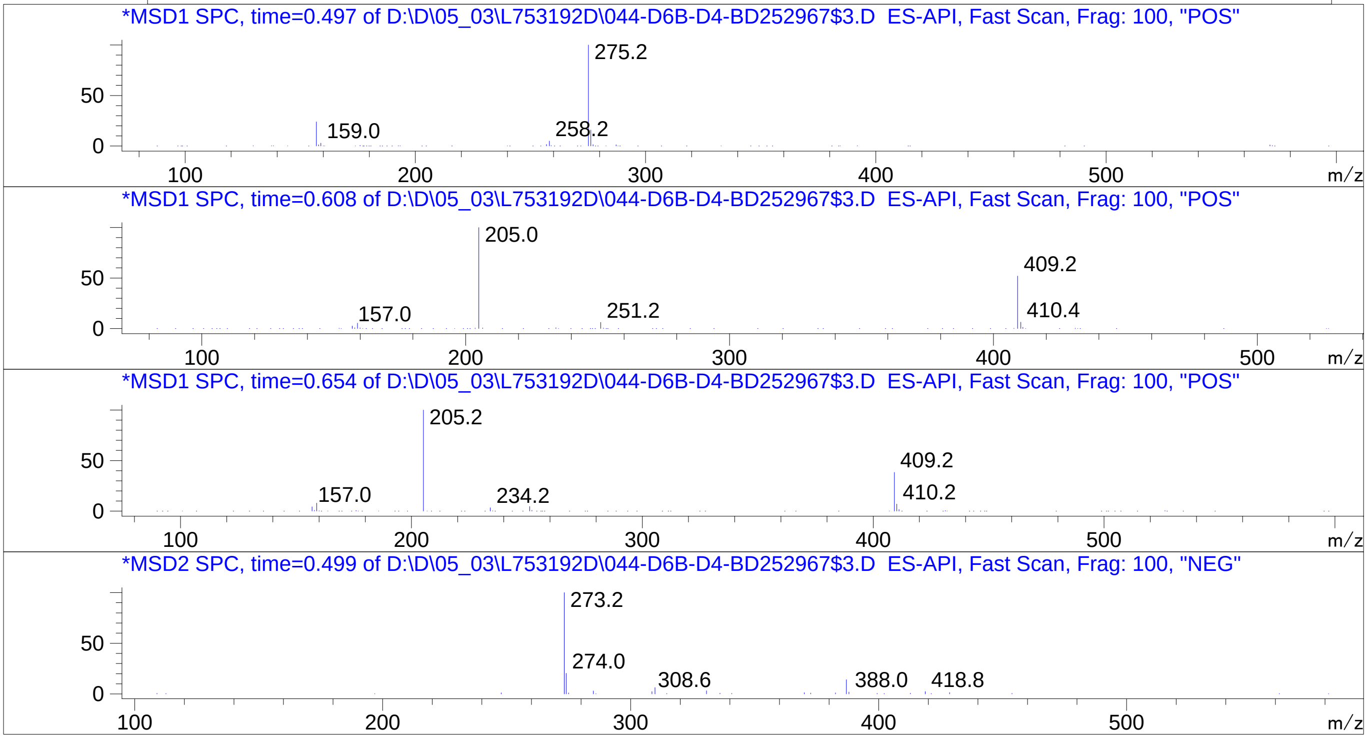


Inj.Date 5/3/2024 N
C:\Users\Public\Documents\ChemStation\1\Data\05_03\L753192D\SUPOR_30.M

RT 0.608

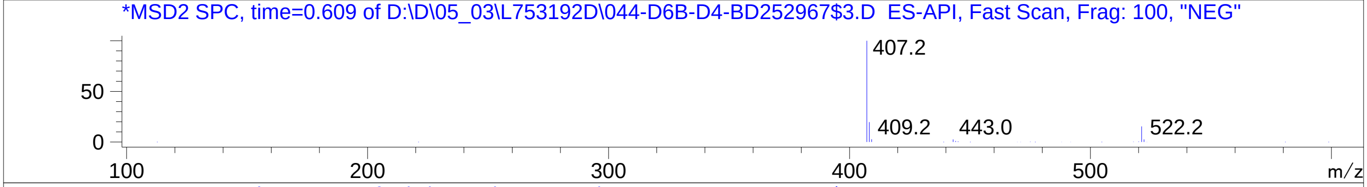


RT 0.655

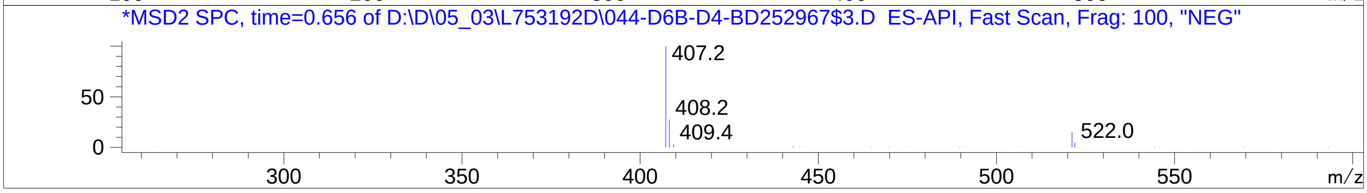


Inj.Date 5/3/2024 N
C:\Users\Public\Documents\ChemStation\1\Data\05_03\L753192D\SUPOR_30.M
