## Supplementary Data 13 for "SyntheMol-RL: a flexible reinforcement learning framework for designing novel and synthesizable antibiotics": Z8921960690.PDF

MaxPeak:  98.60%
Ret_Time: 0.926 min

# BD252969$1

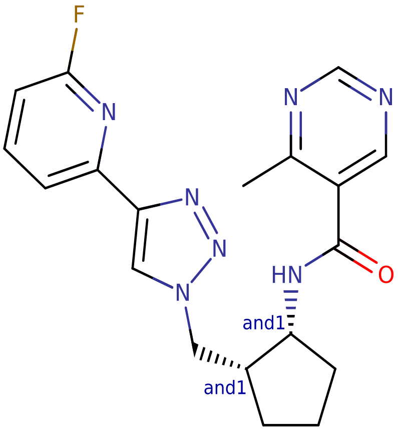


**Mol Wt** 381.41

**Exact Mass** 381.19

| # | Time | Area% |
|---|---|---|
| 1 | 0.926 | 98.60 |
| 2 | 1.157 | 1.40 |

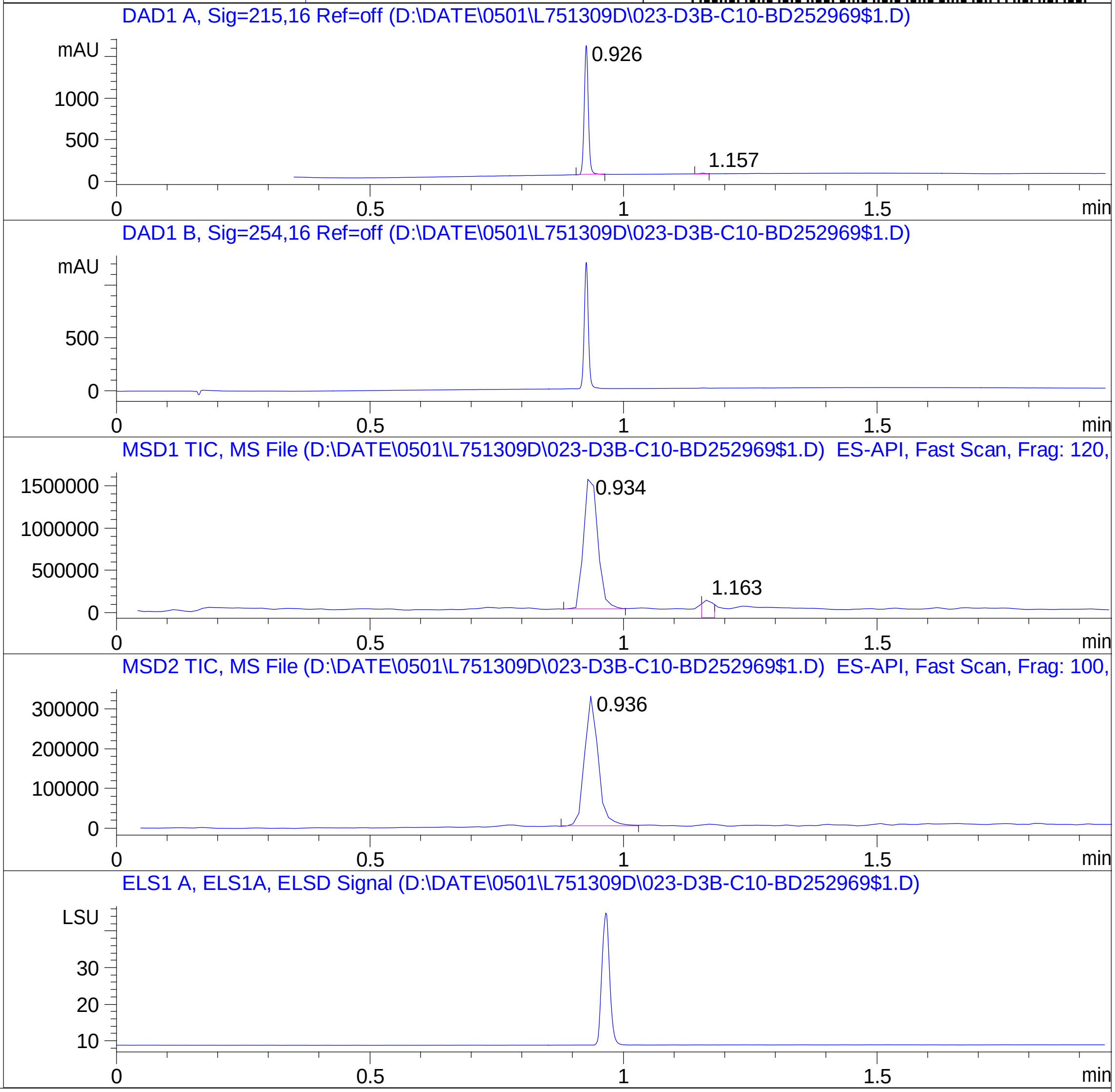


RT 0.934

RT 1.163

RT 0.936

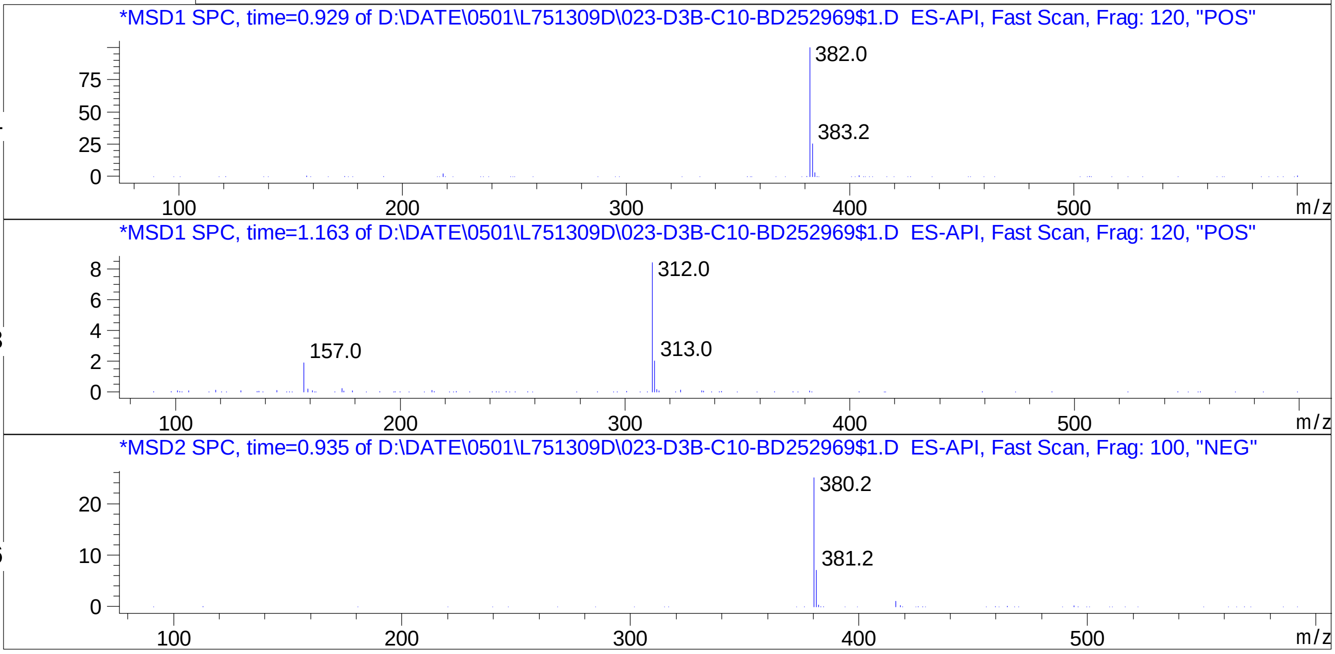


Inj.Date 5/1/2024 CH 21

C:\Users\Public\Documents\ChemStation\1\Data\04_30\L751309D\SUPOR_96.M
