## Supplementary Data 13 for "SyntheMol-RL: a flexible reinforcement learning framework for designing novel and synthesizable antibiotics": Z8921960719.PDF

# BD252822$1

MaxPeak: 96.26%
Ret_Time: 1.092 min

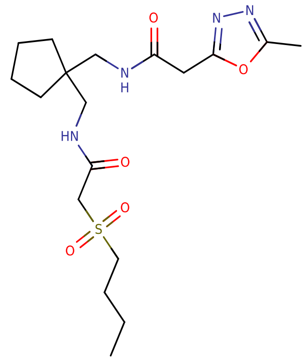


**Mol Wt** 414.52

**Exact Mass** 414.22

| # | Time | Area% |
|---|---|---|
| 1 | 1.092 | 96.26 |
| 2 | 1.151 | 3.74 |

DAD1 A, Sig=215,16 Ref=off (D:\DATE\2024\MAI\0305\L752562D\SAMPL000047.D)

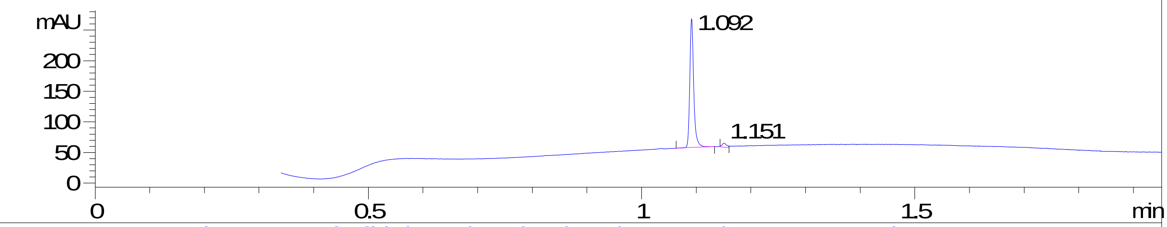


DAD1 B, Sig=254,16 Ref=off (D:\DATE\2024\MAI\0305\L752562D\SAMPL000047.D)

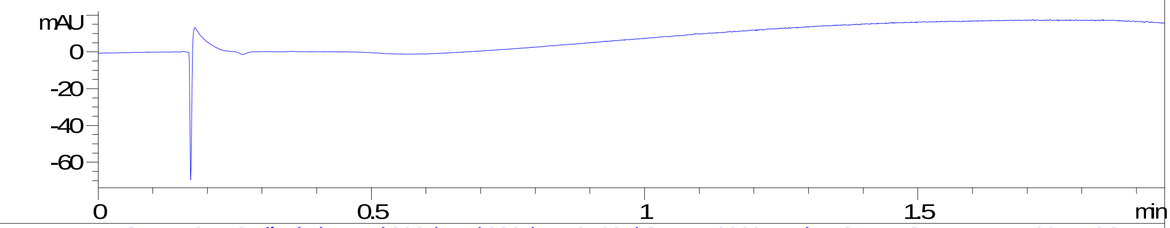


MSD1 TIC, MS File (D:\DATE\2024\MAI\0305\L752562D\SAMPL000047.D) ES-API, Scan, Frag: 100, "POS"

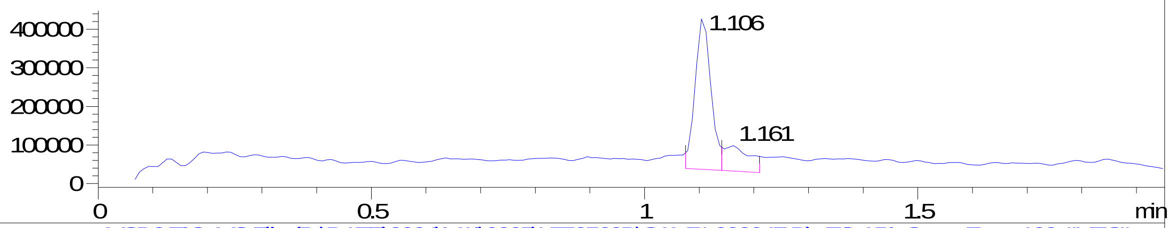


MSD2 TIC, MS File (D:\DATE\2024\MAI\0305\L752562D\SAMPL000047.D) ES-API, Scan, Frag: 100, "NEG"

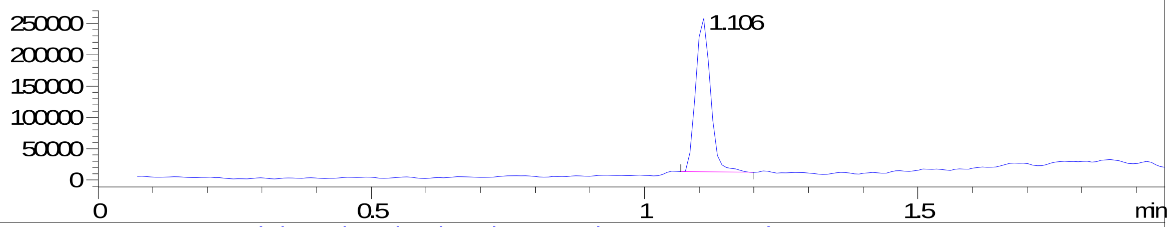


ADC1 A, ELSD (D:\DATE\2024\MAI\0305\L752562D\SAMPL000047.D)

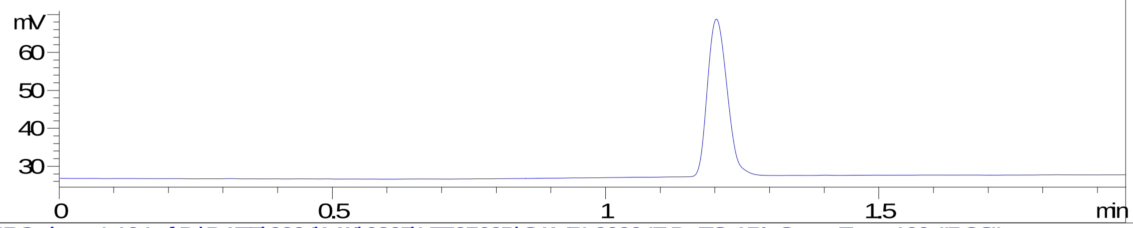


RT 1.106

*MSD1 SPC, time=1.104 of D:\DATE\2024\MAI\0305\L752562D\SAMPL000047.D ES-API, Scan, Frag: 100, "POS"

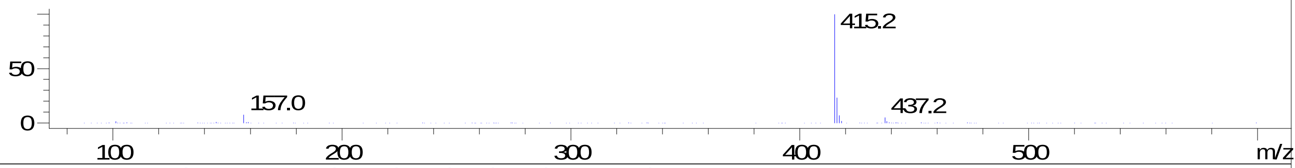


RT 1.161

*MSD1 SPC, time=1.162 of D:\DATE\2024\MAI\0305\L752562D\SAMPL000047.D ES-API, Scan, Frag: 100, "POS"

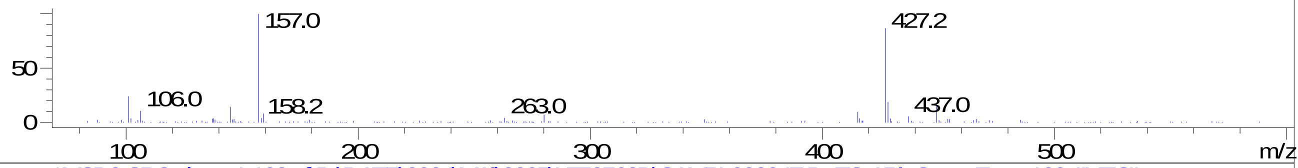


RT 1.106

*MSD2 SPC, time=1.108 of D:\DATE\2024\MAI\0305\L752562D\SAMPL000047.D ES-API, Scan, Frag: 100, "NEG"

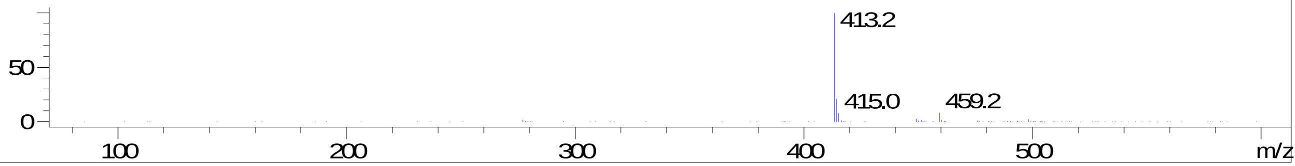
