## Supplementary Data 13 for "SyntheMol-RL: a flexible reinforcement learning framework for designing novel and synthesizable antibiotics": Z8921960720.PDF

# BD252936$2

MaxPeak: 97.88%
Ret_Time: 1.073 min

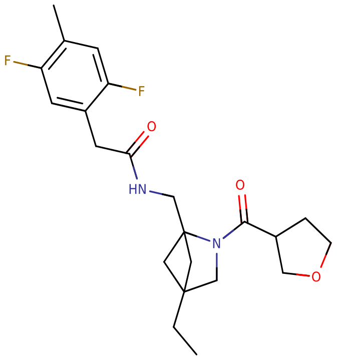


**Mol Wt** 406.47

**Exact Mass** 406.25

| # | Time | Area% |
|---|---|---|
| 1 | 1.041 | 2.12 |
| 2 | 1.073 | 97.88 |

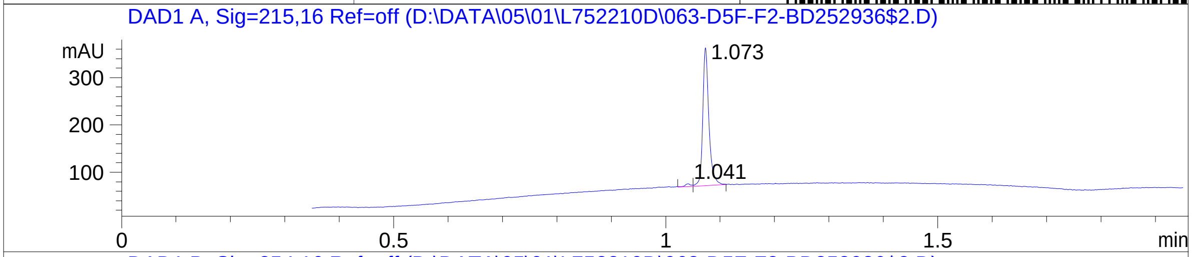


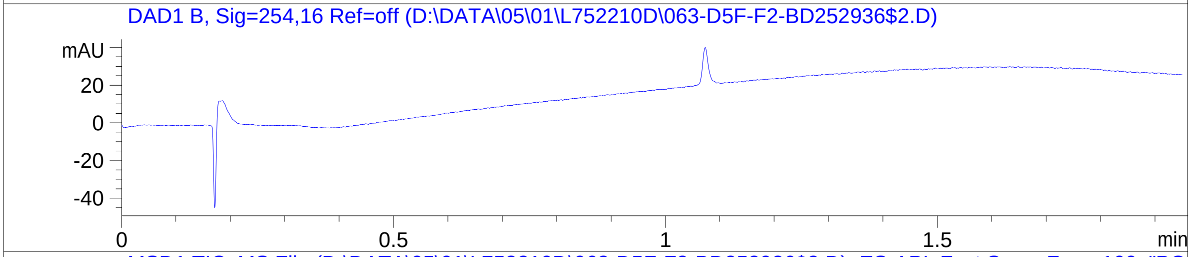


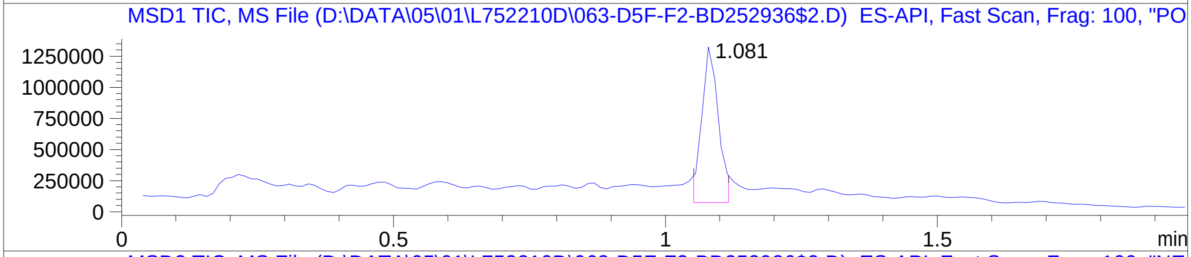


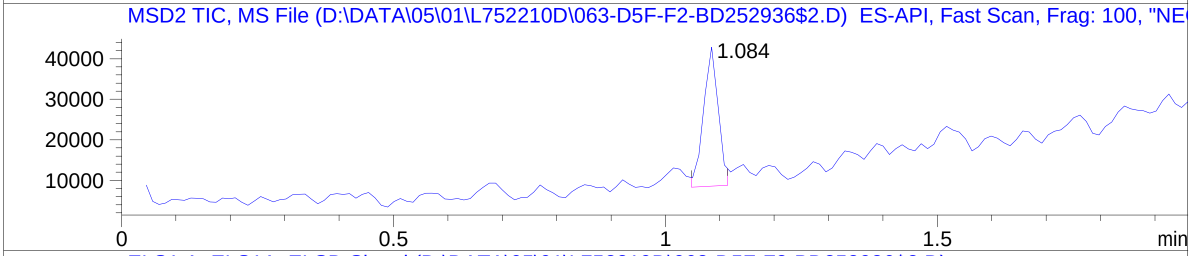


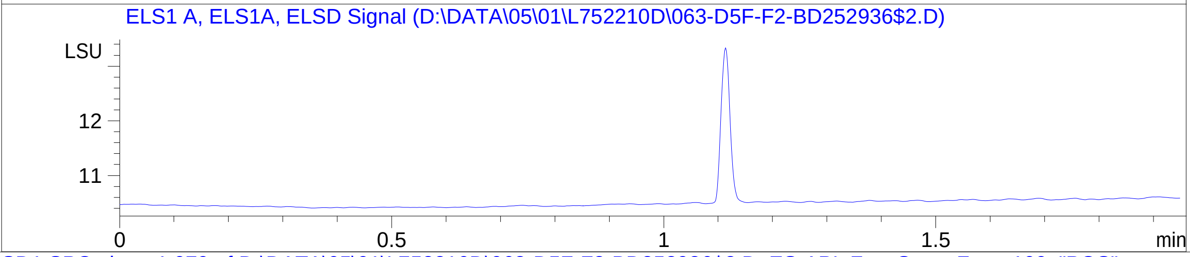


RT 1.081

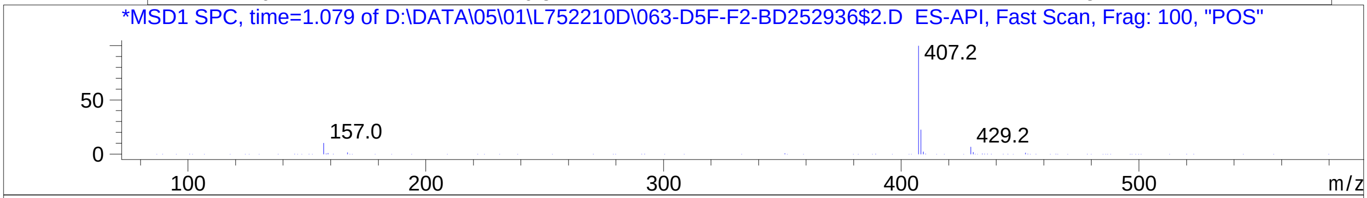


RT 1.084

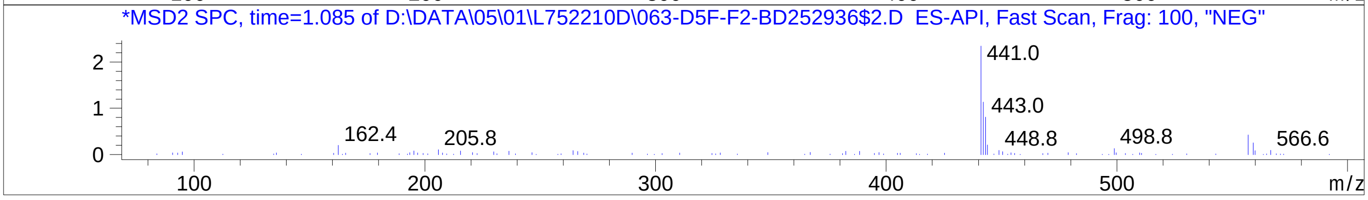


Inj.Date 5/1/2024 CH
C:\Users\Public\Documents\ChemStation\1\Data\05_01\L752210D\SUPOR_30.M
