## Supplementary Data 13 for "SyntheMol-RL: a flexible reinforcement learning framework for designing novel and synthesizable antibiotics": Z8921960722.PDF

# BD252972$1

MaxPeak: 100.00%
Ret_Time: 1.039 min

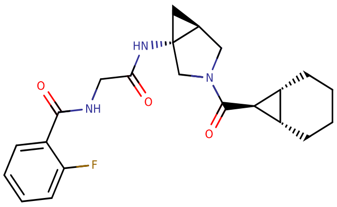

**Mol Wt** 399.46

**Exact Mass** 399.23

| # | Time | Area% |
|---|---|---|
| 1 | 1.039 | 100.00 |

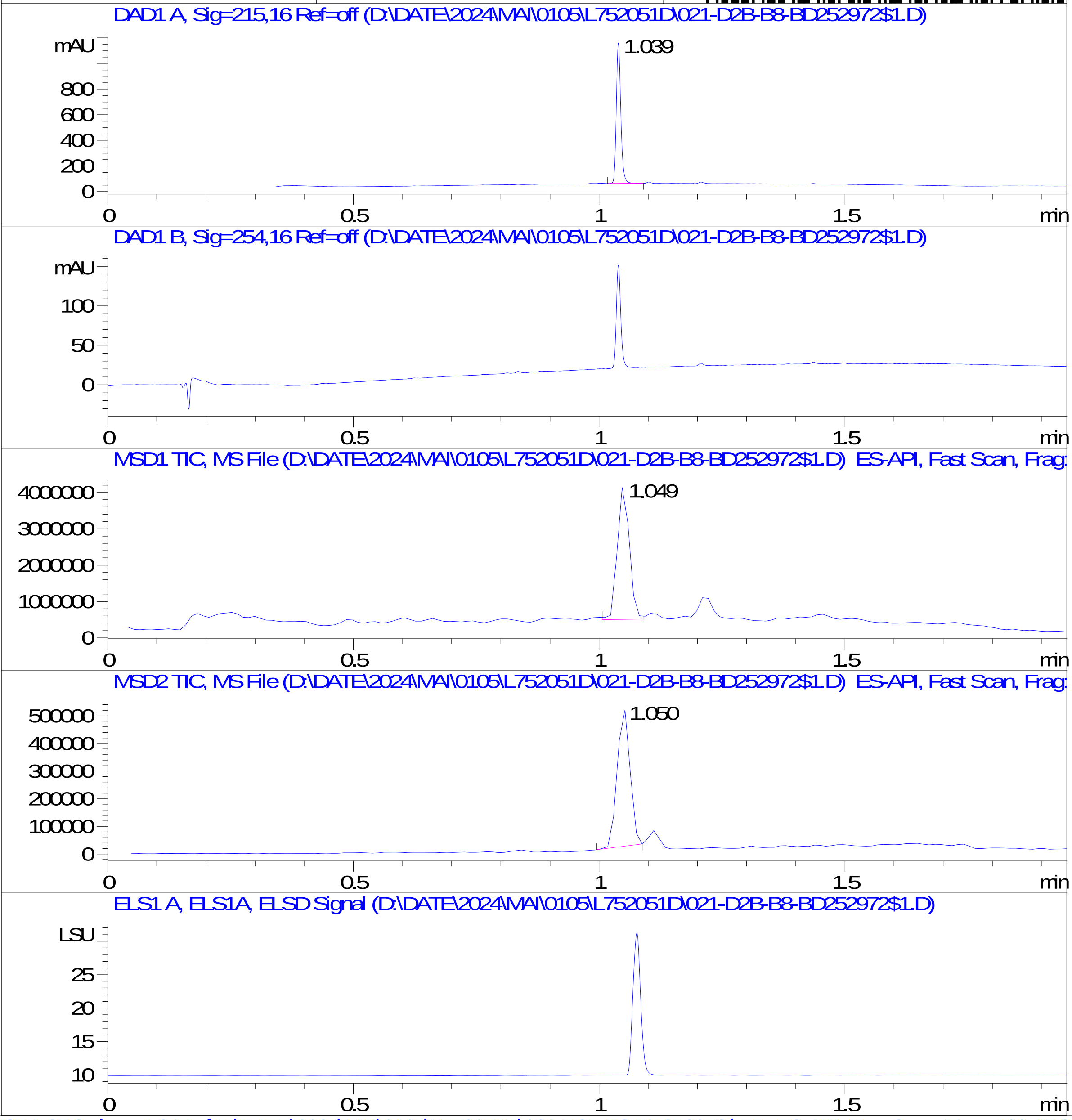


RT 1.049

RT 1.050

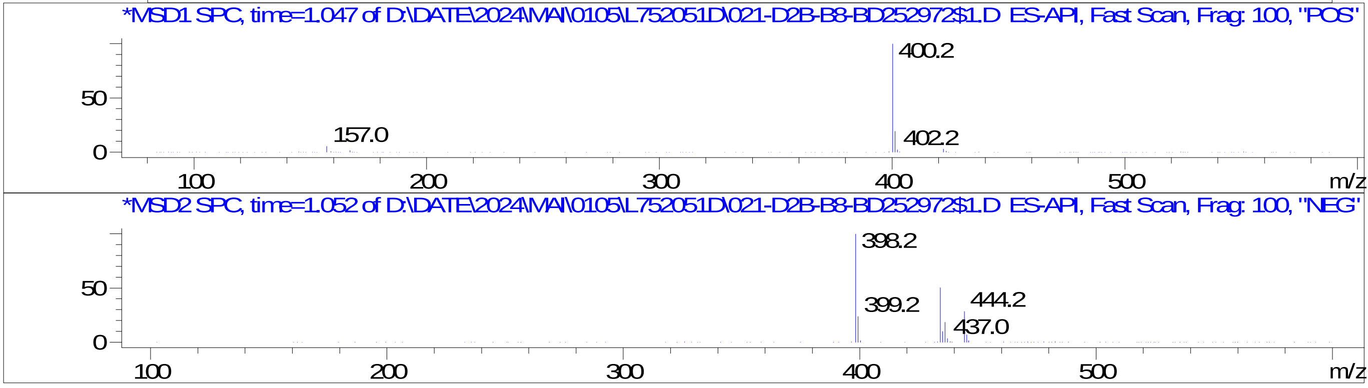


Inj.Date 5/1/2024 AN 20
C:\Users\Public\Documents\ChemStation\1\Data\05_01\L752051D\SUPOR_96.M
