## Supplementary Data 13 for "SyntheMol-RL: a flexible reinforcement learning framework for designing novel and synthesizable antibiotics": Z8921960726.PDF

MaxPeak: 57.46%
Ret_Time: 1.036 min

# BD252824$2

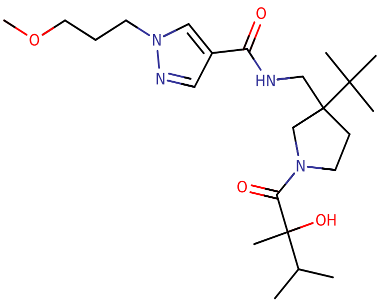


**Mol Wt** 436.59

**Exact Mass** 436.36

| # | Time | Area% |
|---|---|---|
| 1 | 1.029 | 41.41 |
| 2 | 1.036 | 57.46 |
| 3 | 1.066 | 1.12 |

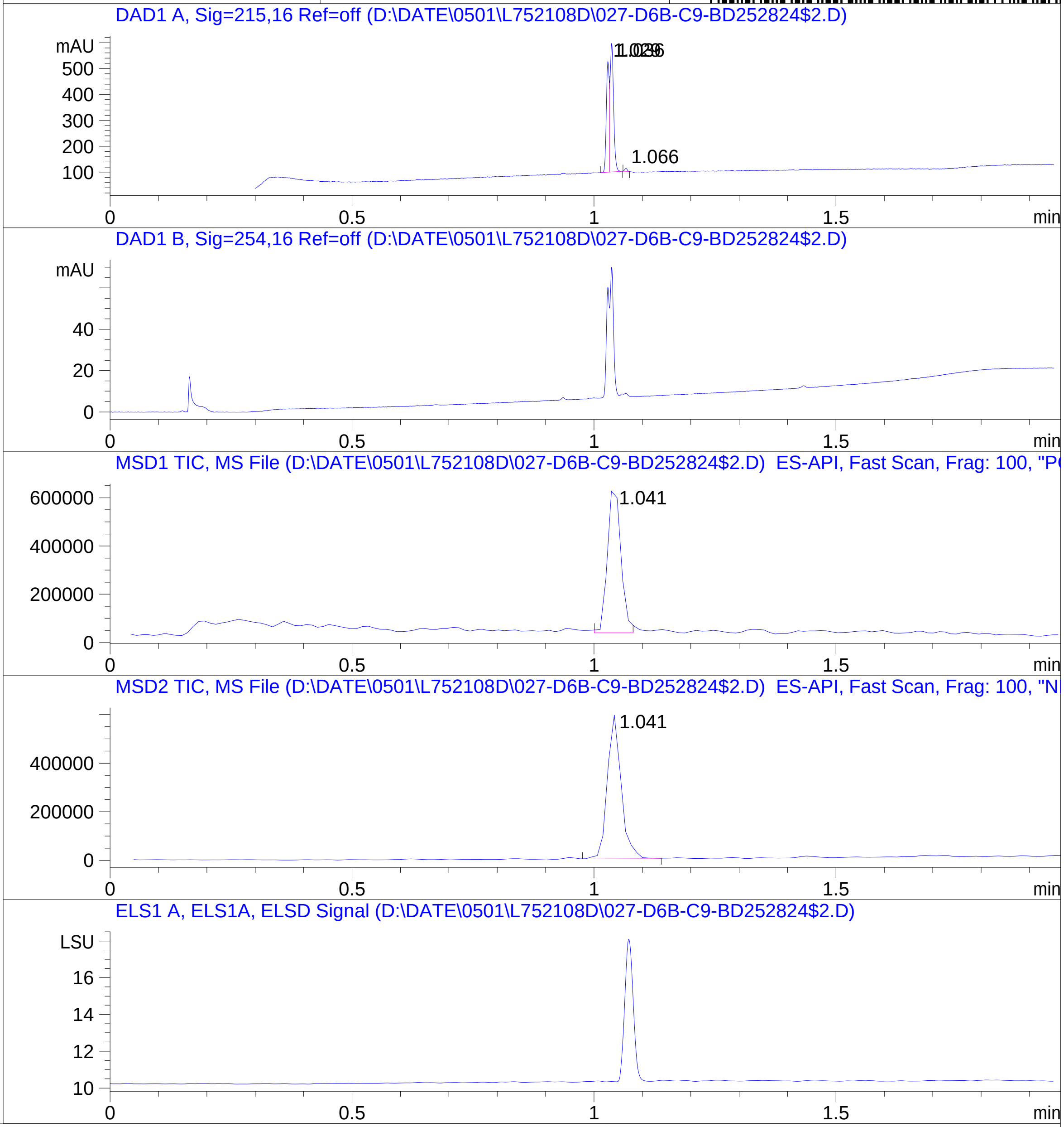


RT 1.041

RT 1.041

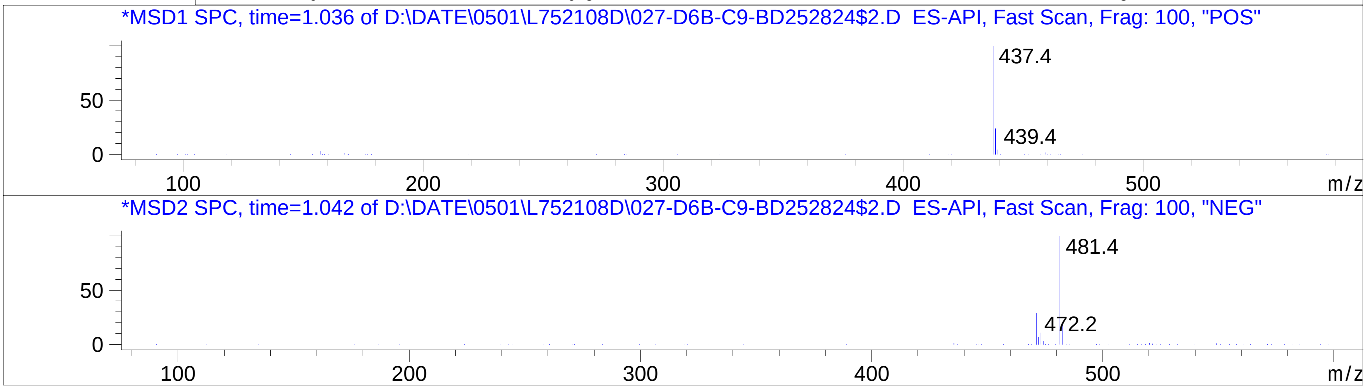


Inj.Date 5/1/2024 0 32
C:\Users\Public\Documents\ChemStation\1\Data\05_01\L752108D\SUPOR_30.M
