## Supplementary Data 13 for "SyntheMol-RL: a flexible reinforcement learning framework for designing novel and synthesizable antibiotics": Z8921960815.PDF

# BD252759$11

MaxPeak: 100.00%
Ret_Time: 0.502 min

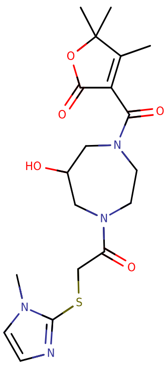

**Mol Wt** 422.5
**Exact Mass** 422.18

| # | Time | Area% |
|---|---|---|
| 1 | 0.502 | 100.00 |

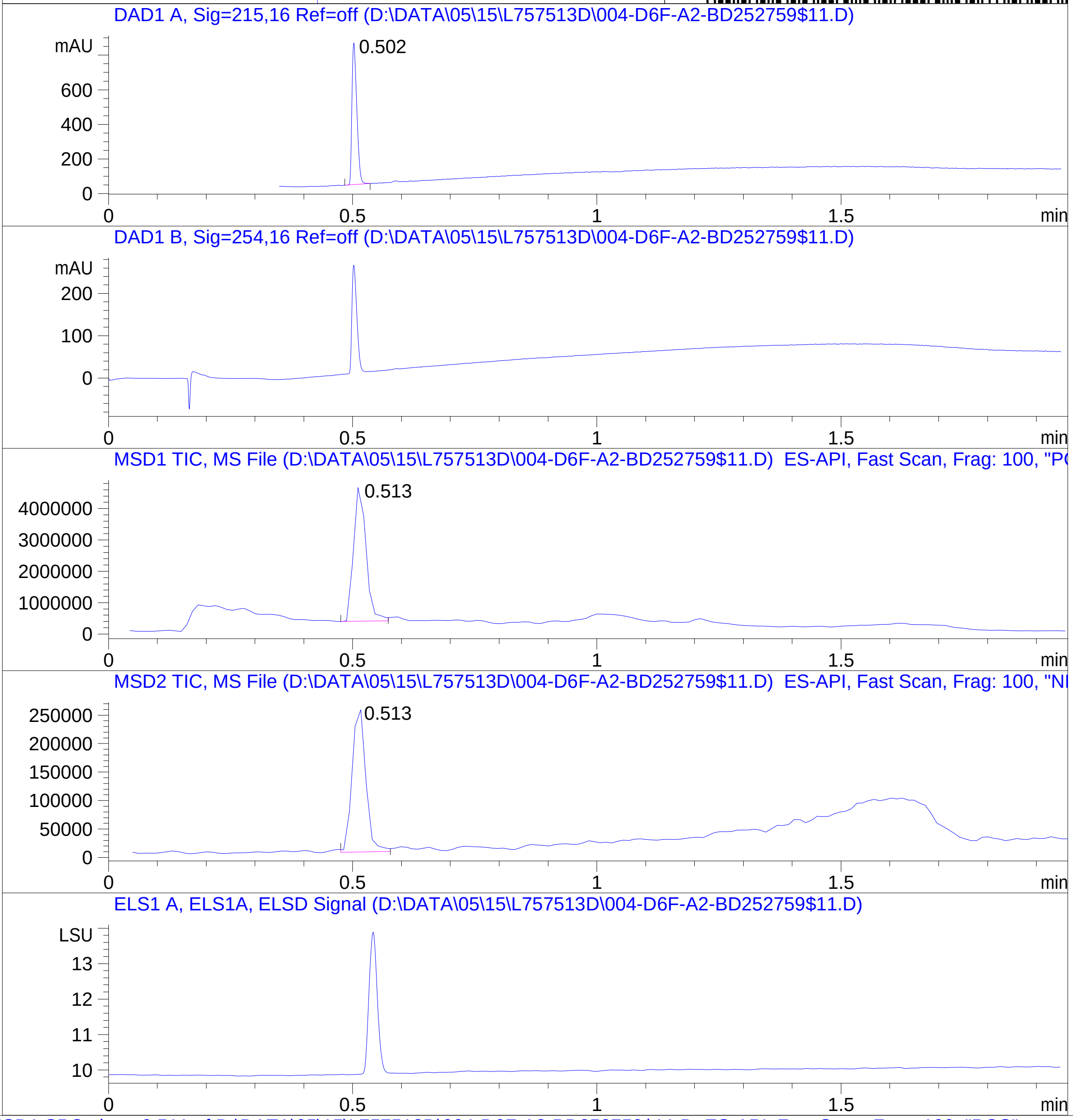


RT 0.513

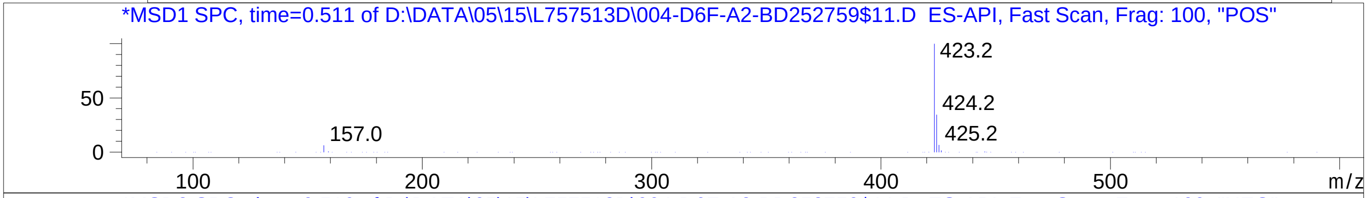


RT 0.513

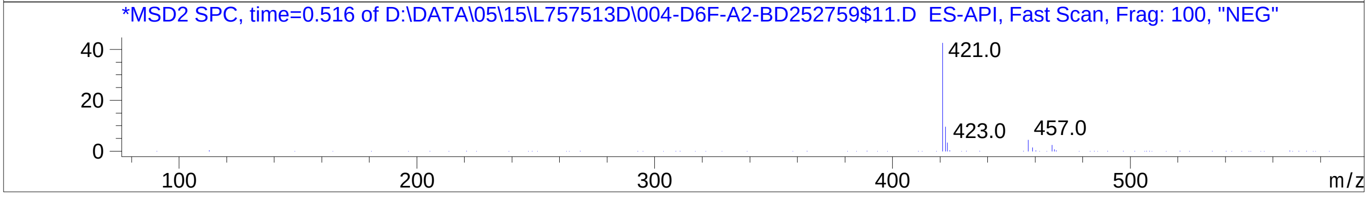


Inj.Date 5/15/2024 CH 31
C:\Users\Public\Documents\ChemStation\1\Data\05_15\L757513D\SUPOR_30.M
