## Supplementary figures and images for "SyntheMol-RL: a flexible reinforcement learning framework for designing novel and synthesizable antibiotics"

### Z1481198550.PDF

MaxPeak: 98.83%
Ret_Time: 0.906 min

# BE177025$1

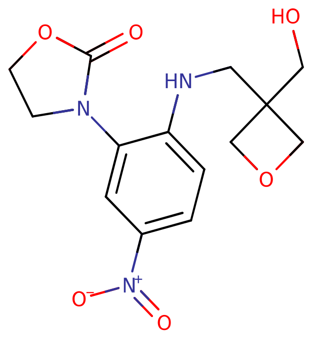


**Mol Wt** 323.3

**Exact Mass** 323.11

| # | Time | Area% |
|---|---|---|
| 1 | 0.846 | 1.17 |
| 2 | 0.906 | 98.83 |

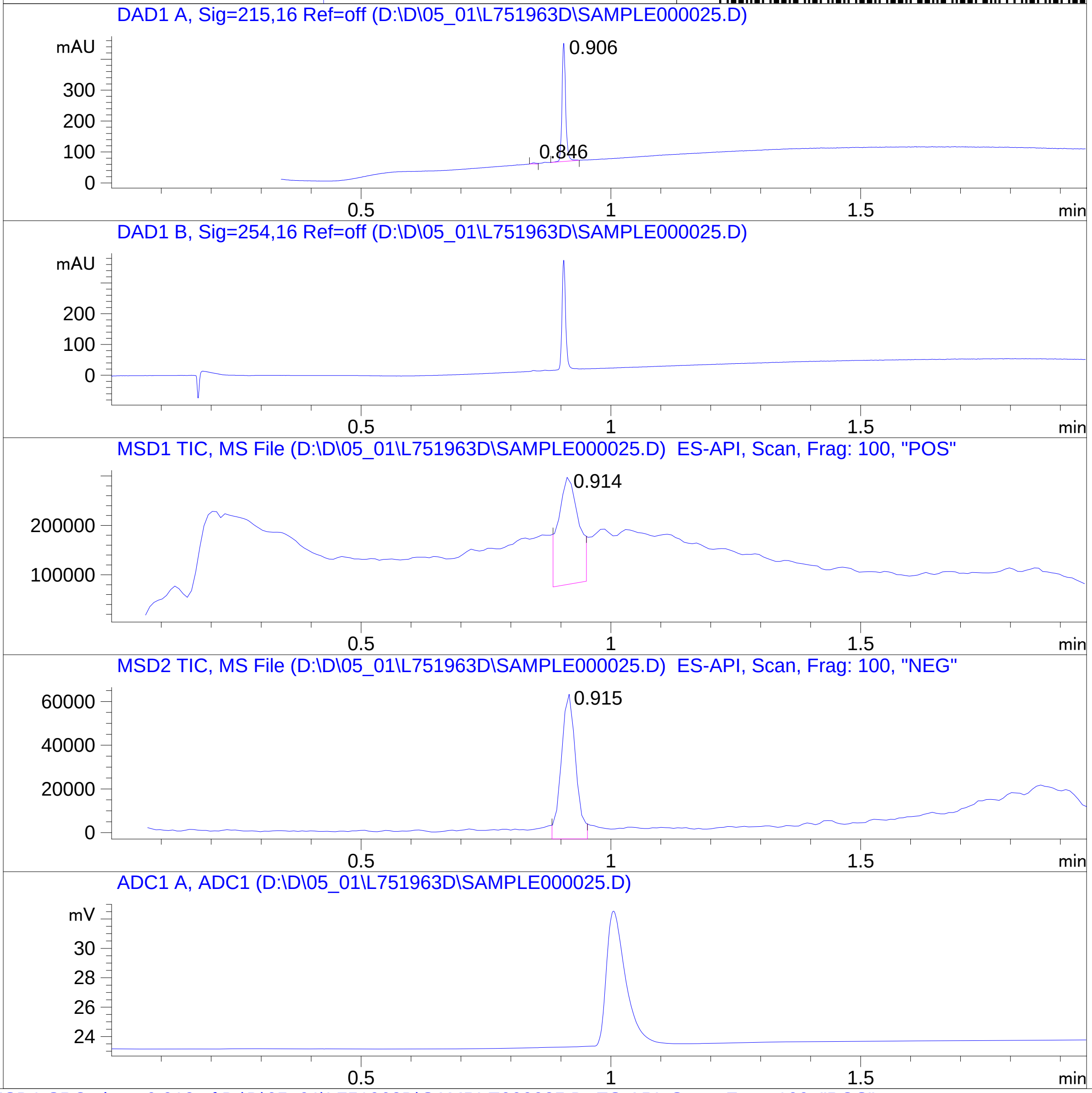


RT 0.914

RT 0.915

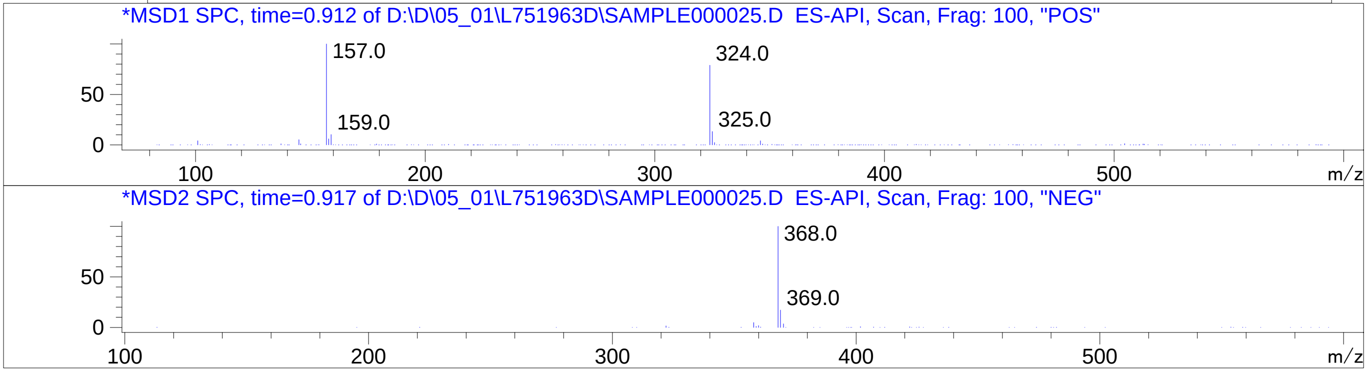


Inj.Date 5/1/2024 N
C:\CHEM32\1\METHODS\SUPOR_30.M

### Z1488448345.PDF

MaxPeak: 100.00%
Ret_Time: 0.987 min

# BE177080$9

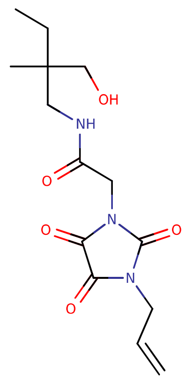


**Mol Wt** 311.33

**Exact Mass** 311.16

| # | Time | Area% |
|---|---|---|
| 1 | 0.987 | 100.00 |

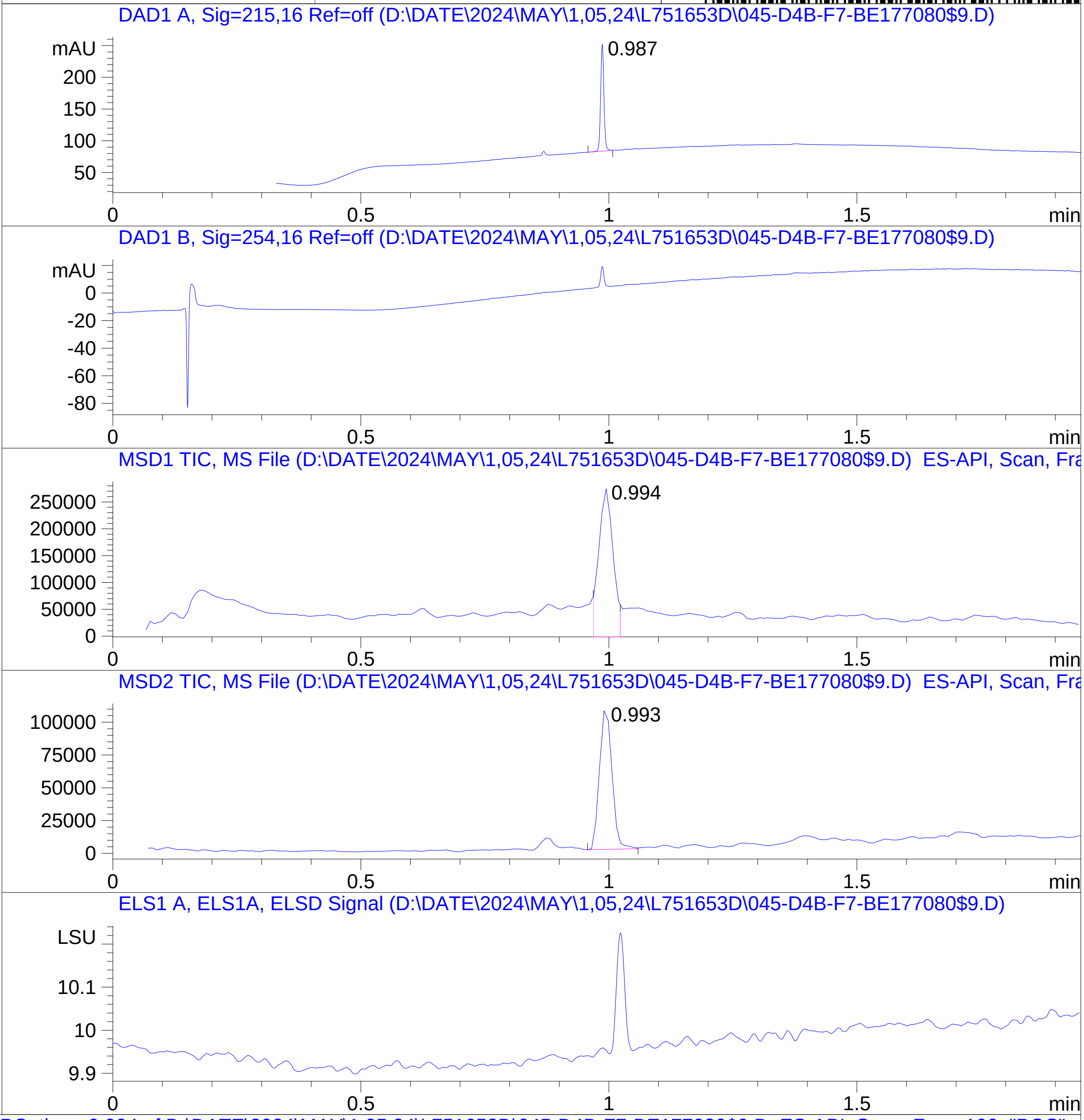


RT 0.994

RT 0.993

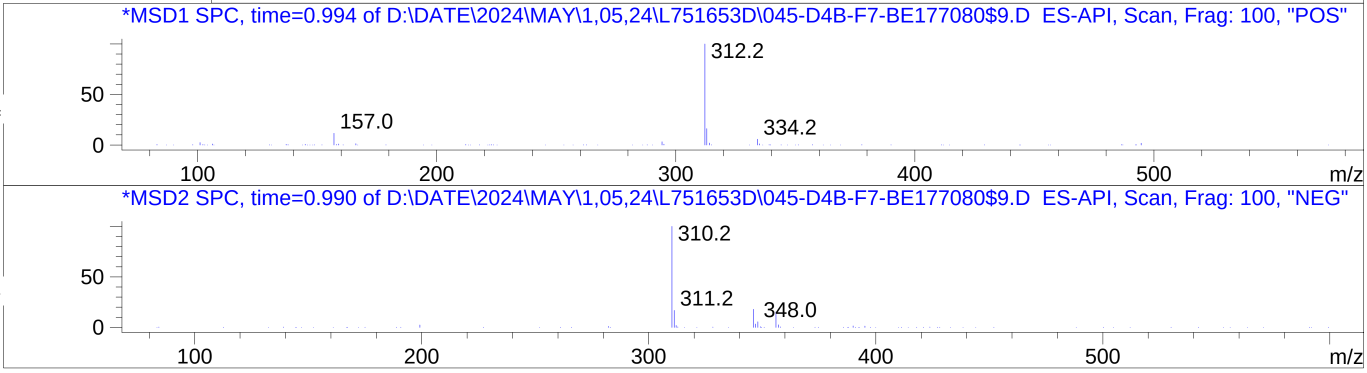


Inj.Date 5/1/2024 T  ->
C:\Chem32\1\Data\04_30\L751653D\SUPOR_30.M

### Z1545576111.PDF

MaxPeak: 100.00%
Ret_Time: 1.151 min

BE177057$1

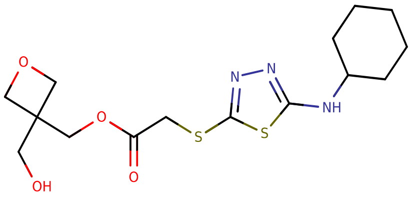


**Mol Wt** 373.49

**Exact Mass** 373.13

| # | Time | Area% |
|---|---|---|
| 1 | 1.151 | 100.00 |

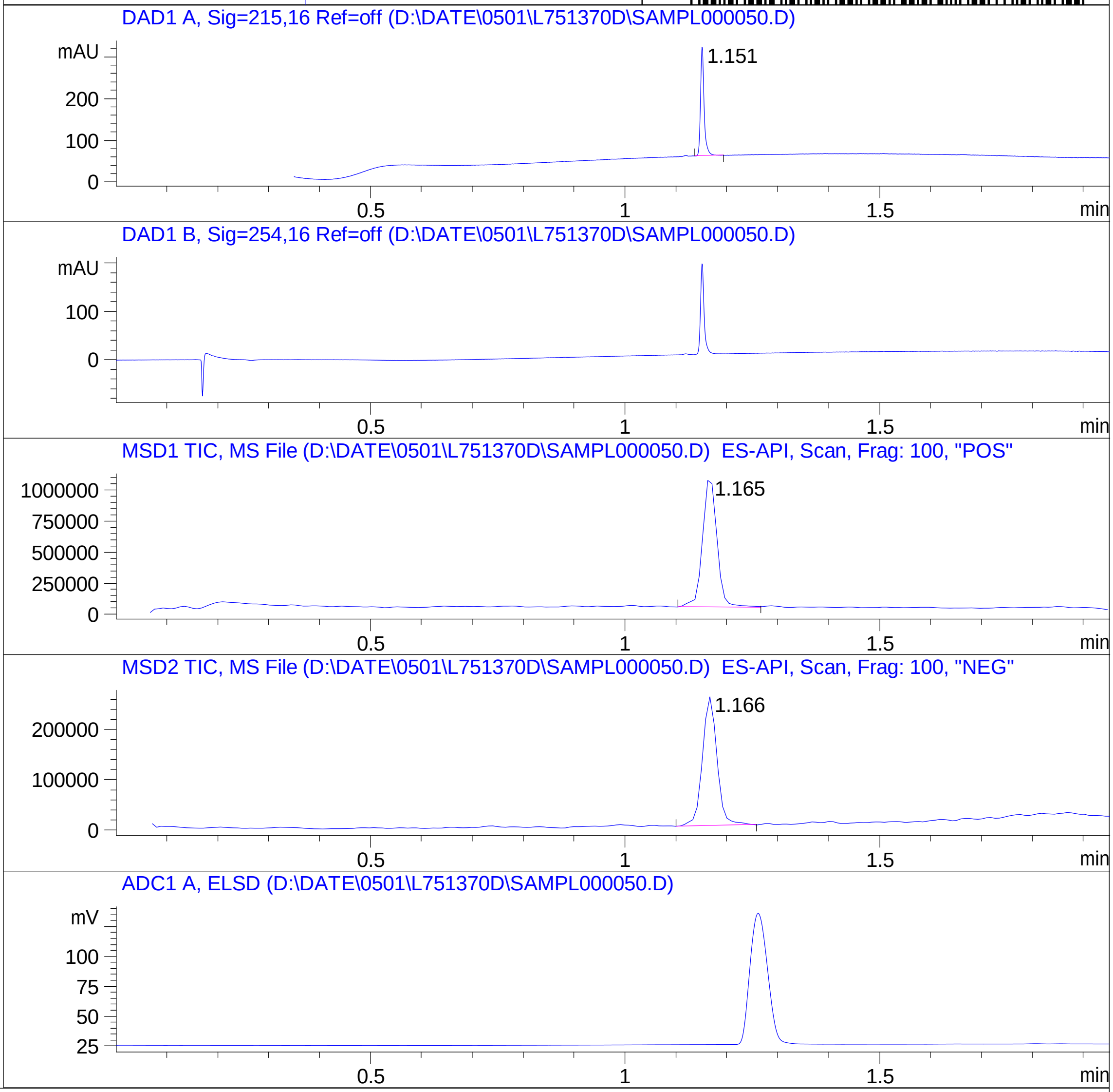


RT 1.165

RT 1.166

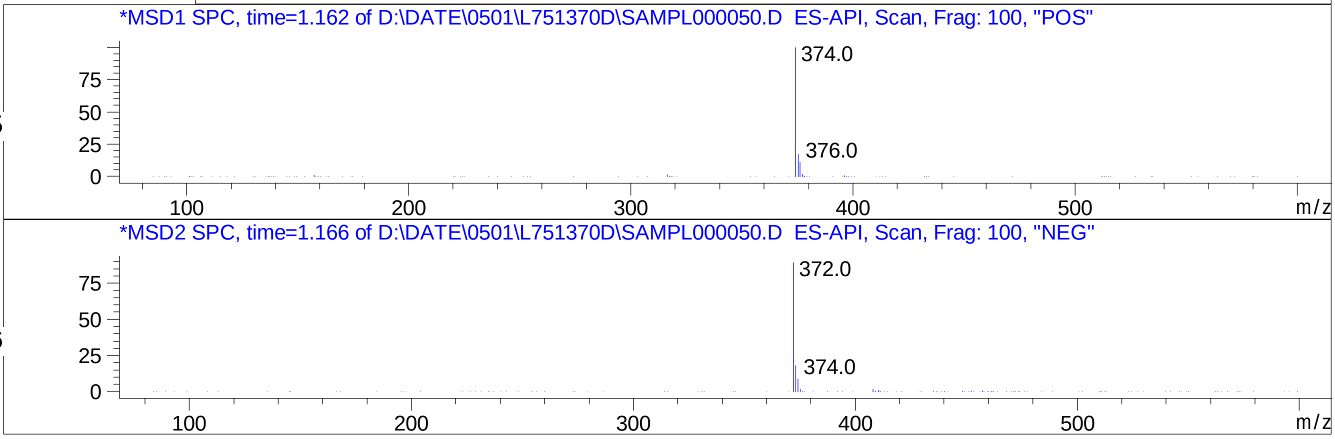


Inj.Date 4/30/2024 CH -6-

C:\CHEM32\1\METHODS\SUPOR_30.M

### Z1593017843.PDF

# BE177058$2

MaxPeak:  100.00%
Ret_Time: 0.952 min

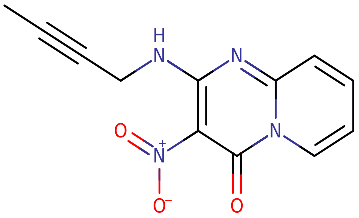


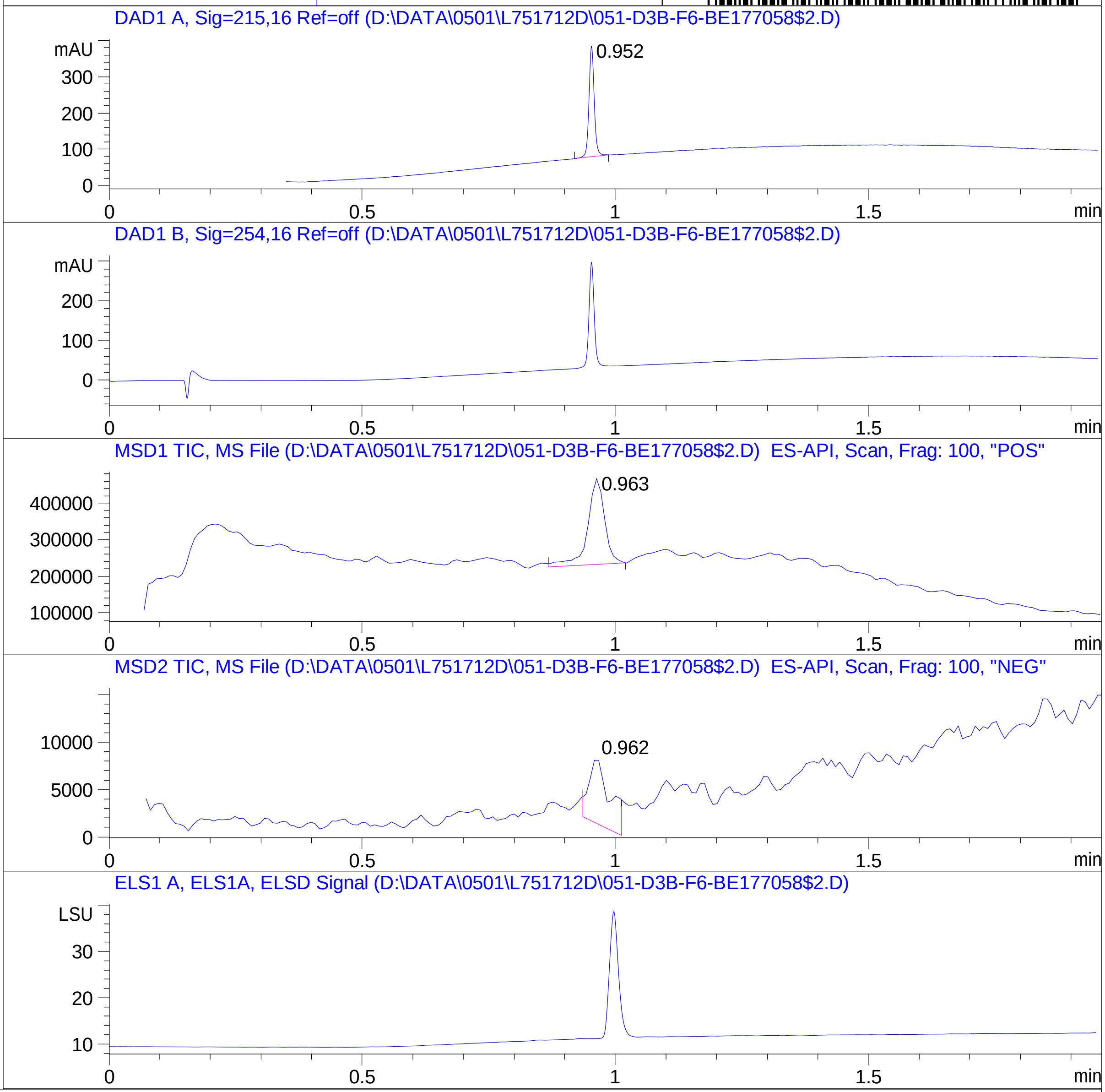


**Mol Wt** 258.23

**Exact Mass** 258.07

| # | Time | Area% |
|---|---|---|
| 1 | 0.952 | 100.00 |

RT 0.963

RT 0.962

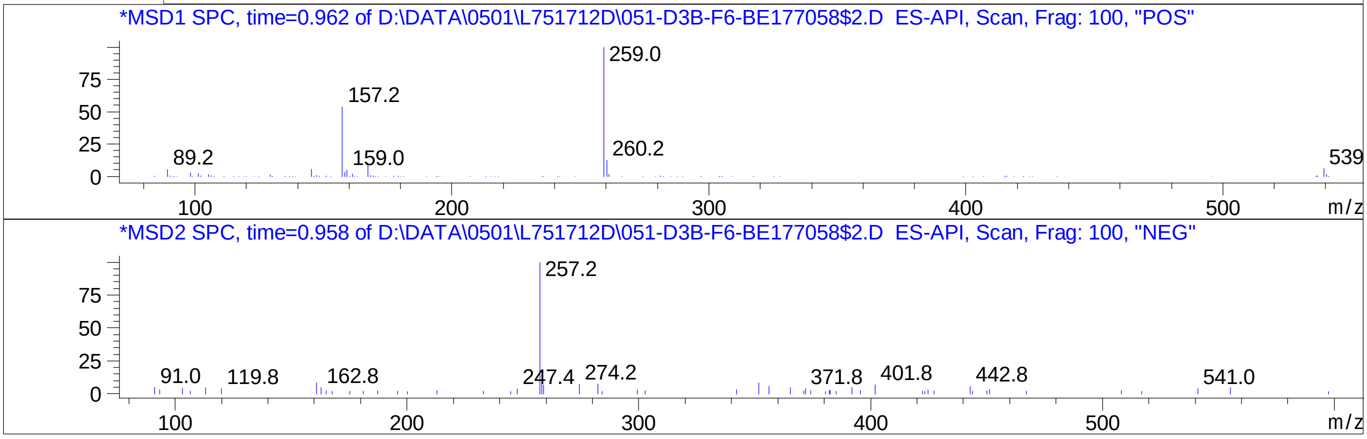


Inj.Date 5/1/2024 VB 13

C:\Chem32\1\Data\04_30\L751712D\SUPOR_30.M

### Z1801673820.PDF

# BE177060$4

MaxPeak: 100.00%
Ret_Time: 0.974 min

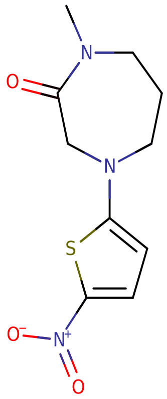


**Mol Wt** 255.29

**Exact Mass** 255.07

| # | Time | Area% |
|---|---|---|
| 1 | 0.974 | 100.00 |

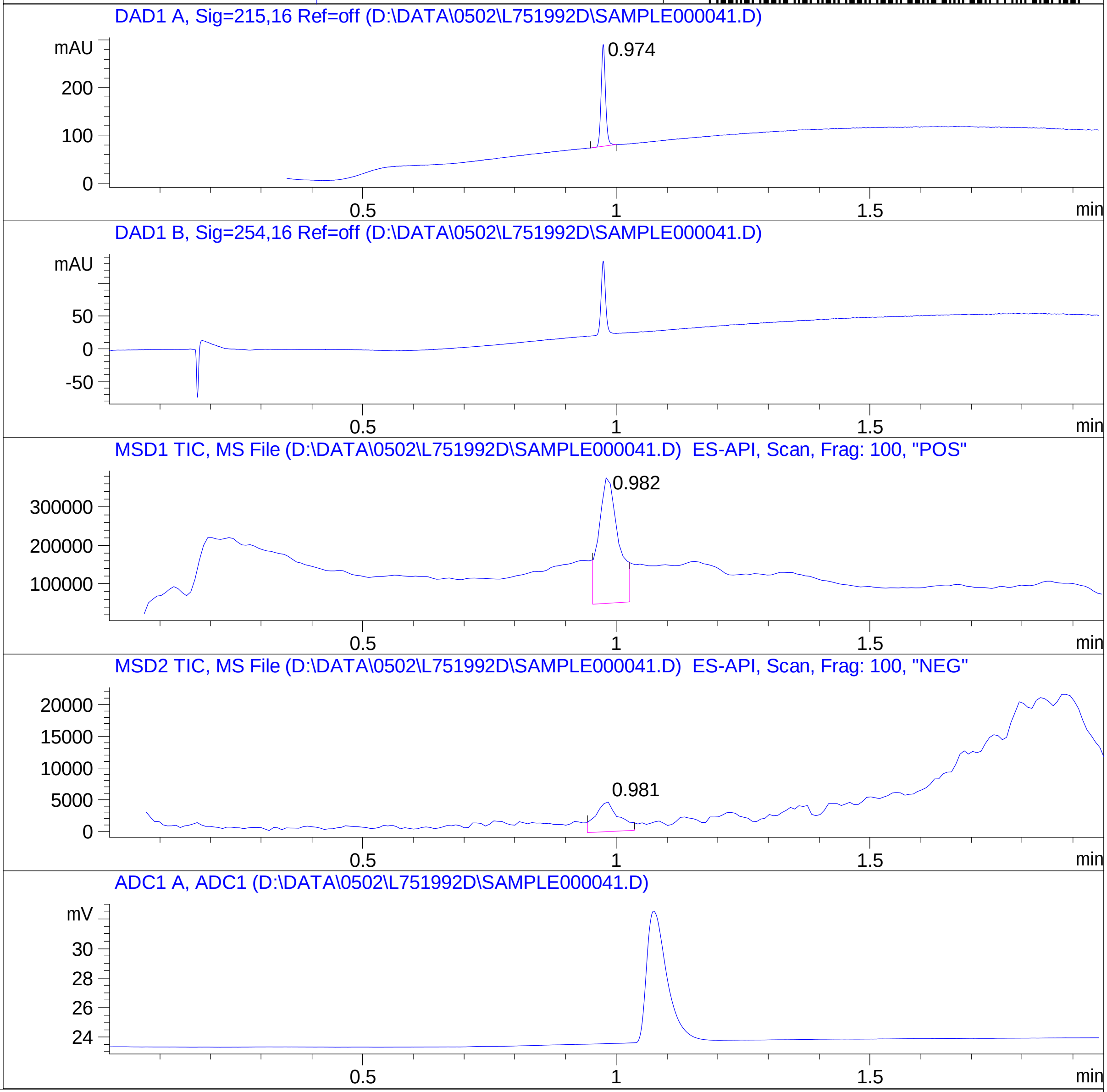


RT 0.982

RT 0.981

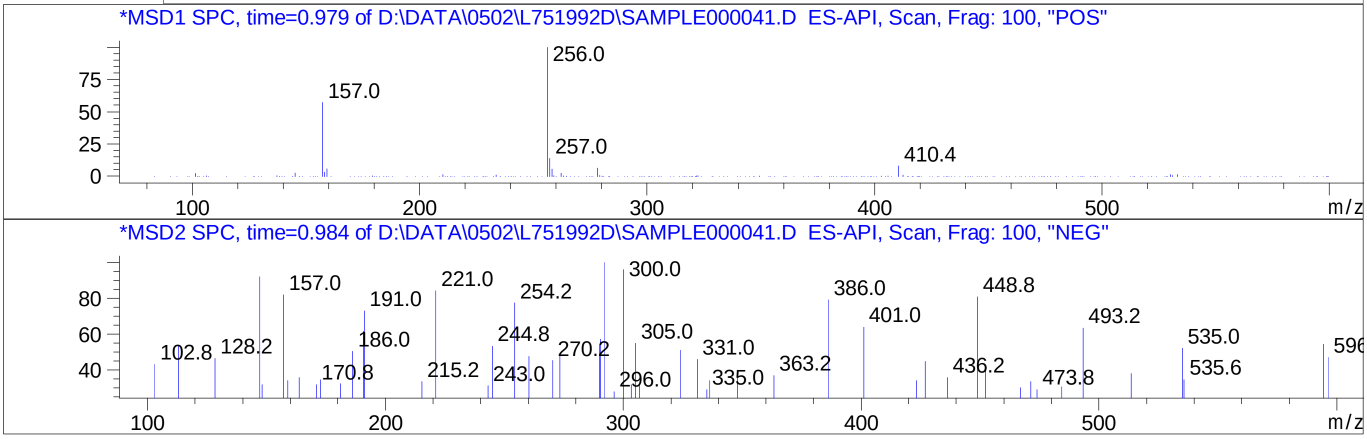

### Z2017198490.PDF

MaxPeak: 100.00%
Ret_Time: 0.969 min

# BE177021$1

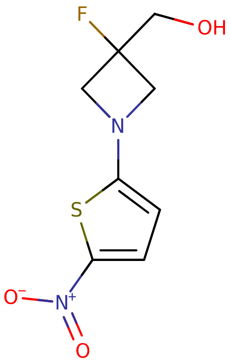


**Mol Wt** 232.23

**Exact Mass** 232.03

| # | Time | Area% |
| --- | --- | --- |
| 1 | 0.969 | 100.00 |

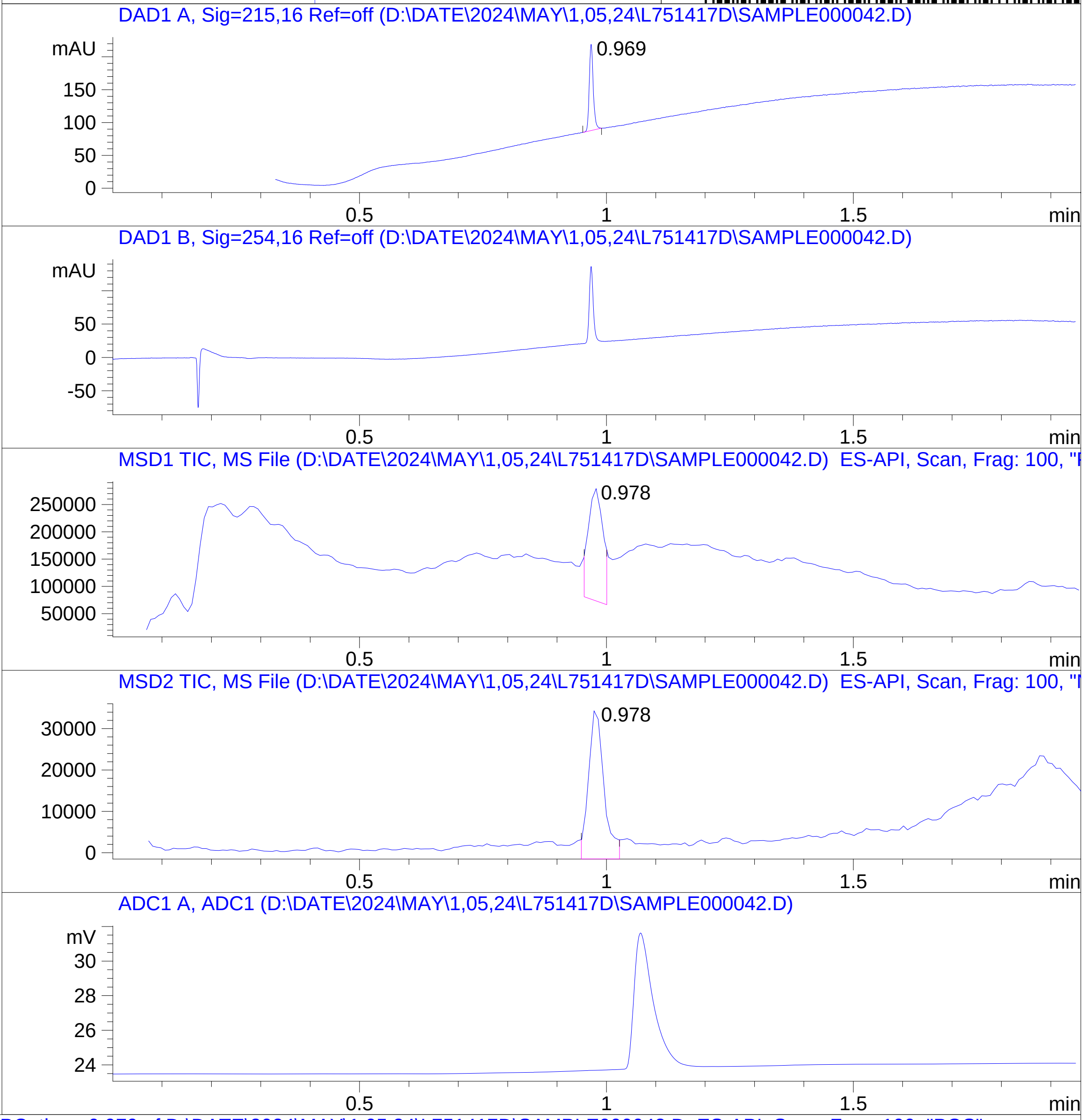


RT 0.978

RT 0.978

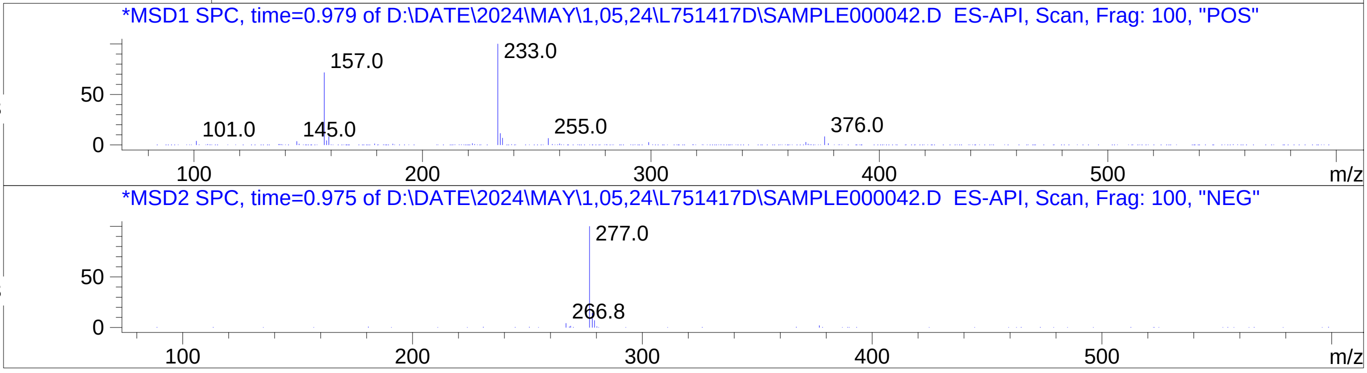

### Z2088917026.PDF

# BD252372$2

MaxPeak: 90.55%
Ret_Time: 0.929 min

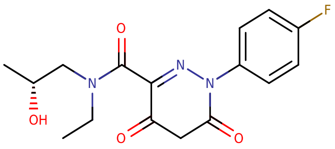


**Mol Wt** 335.33

**Exact Mass** 335.14

| # | Time | Area% |
|---|---|---|
| 1 | 0.765 | 1.15 |
| 2 | 0.929 | 90.55 |
| 3 | 0.974 | 7.05 |
| 4 | 1.147 | 1.26 |

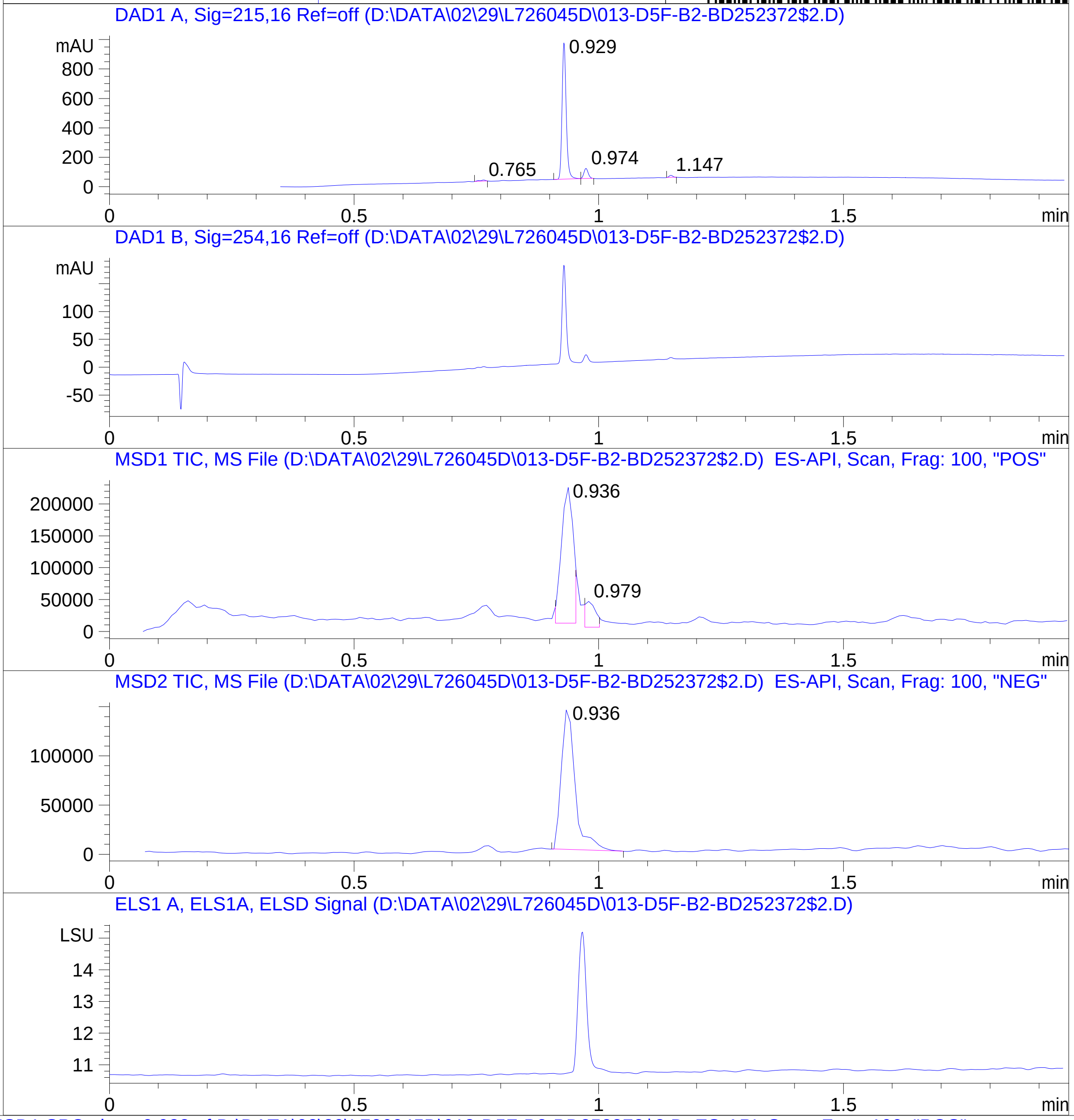


RT 0.936

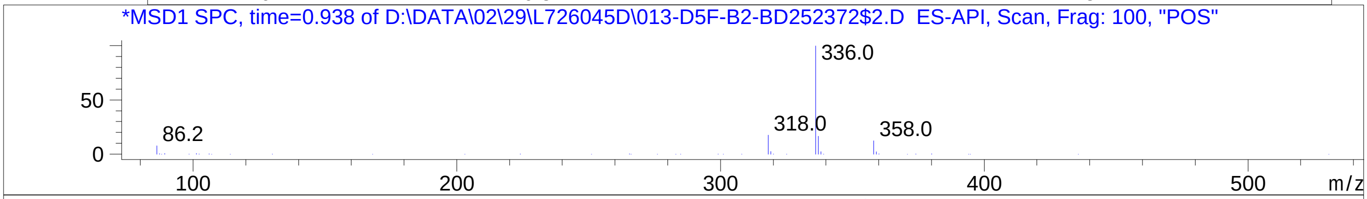


RT 0.979

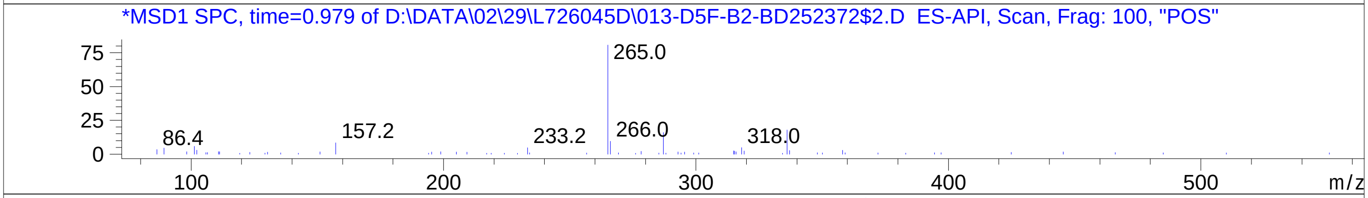


RT 0.936

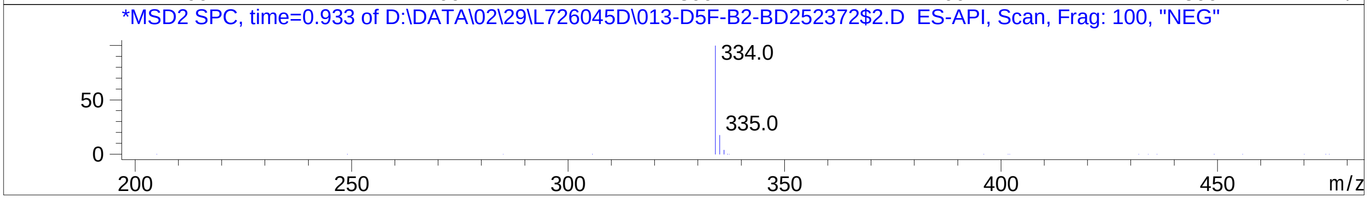


Inj.Date 2/29/2024 CH -16-
C:\Chem32\1\Data\02_29\L726045D\SUPOR_30.M

### Z2088938699.PDF

MaxPeak: 100.00%
Ret_Time: 0.670 min

# BE177042$2

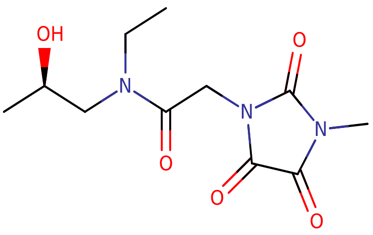


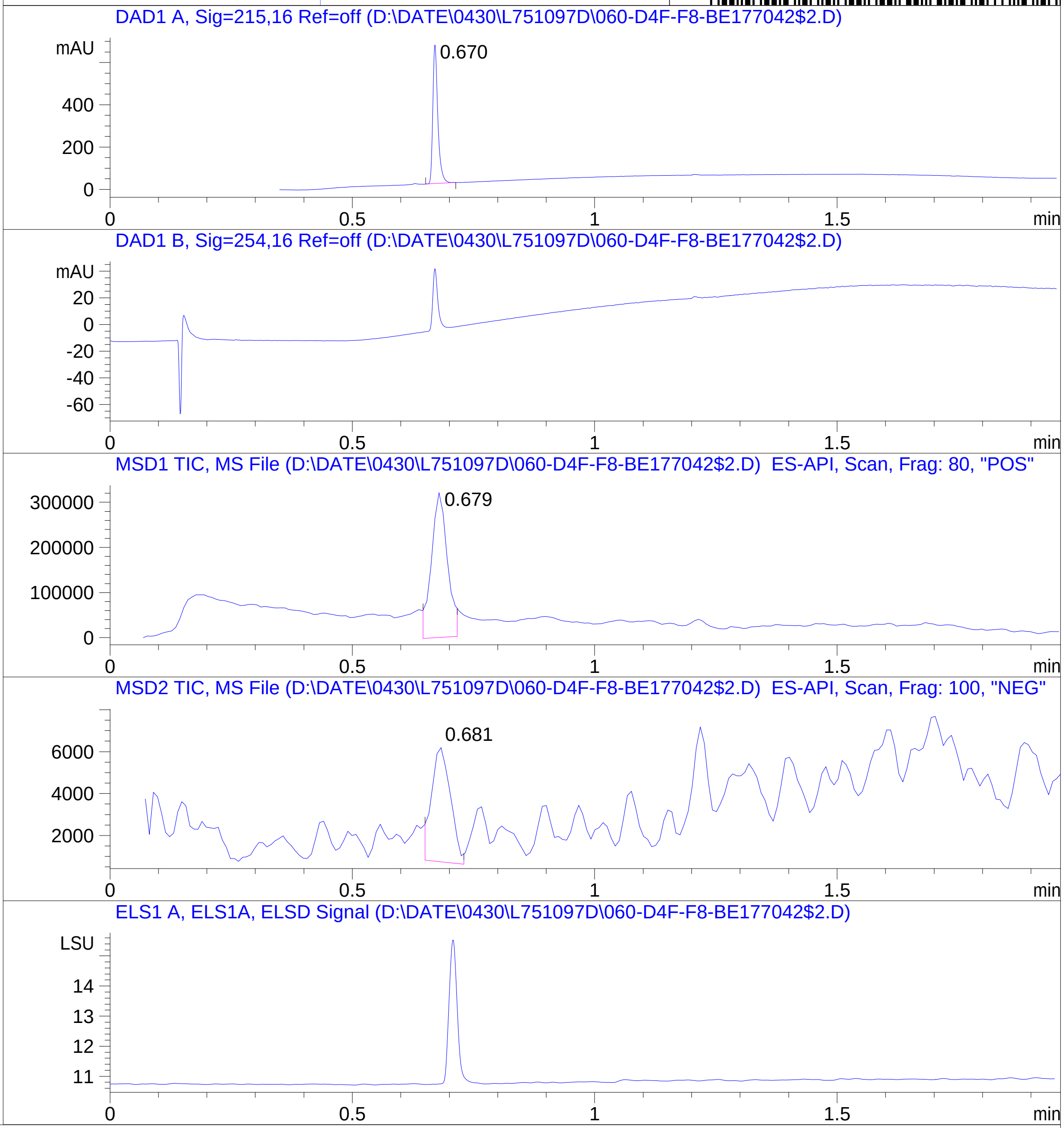


**Mol Wt** 271.27

**Exact Mass** 271.12

```
#  Time   Area%
-----------------
1  0.670  100.00
```

RT 0.679

RT 0.681

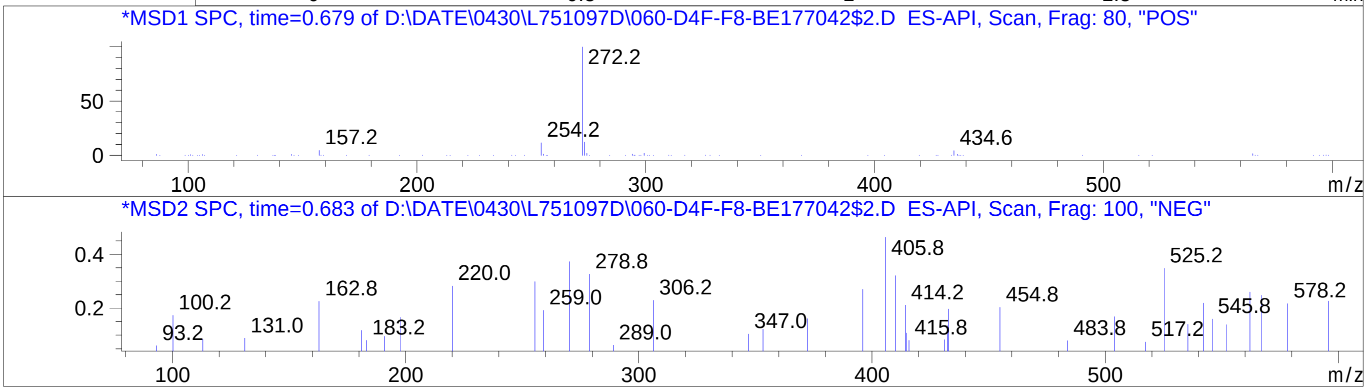

### Z2217869832.PDF

MaxPeak: 95.37%
Ret_Time: 0.899 min

# BE177055$3

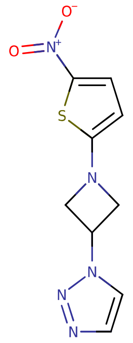


**Mol Wt** 251.26

**Exact Mass** 251.04

| # | Time | Area% |
|---|---|---|
| 1 | 0.899 | 95.37 |
| 2 | 0.920 | 1.72 |
| 3 | 1.179 | 1.20 |
| 4 | 1.197 | 1.72 |

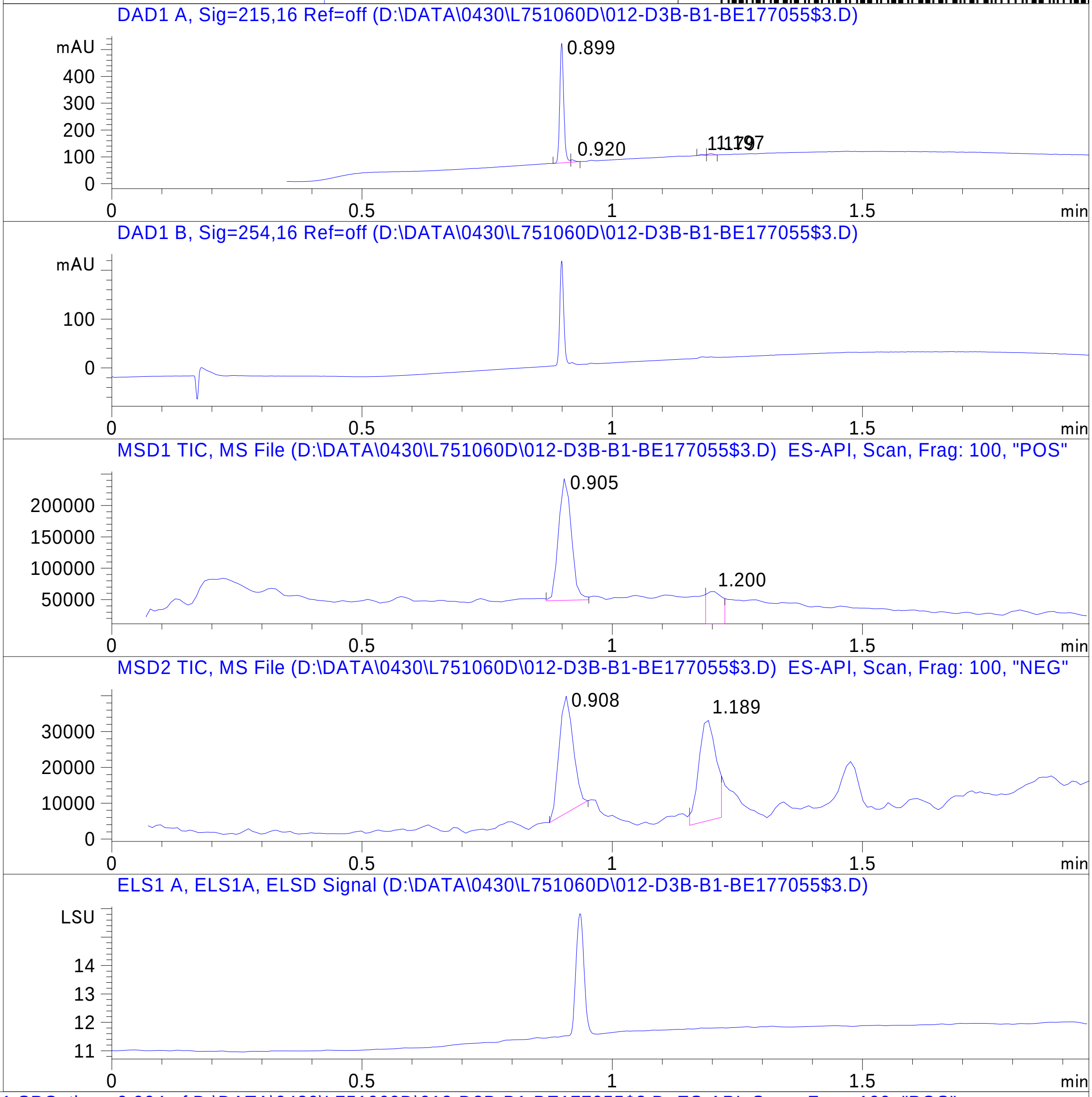


RT 0.905

RT 1.200

RT 0.908

RT 1.189

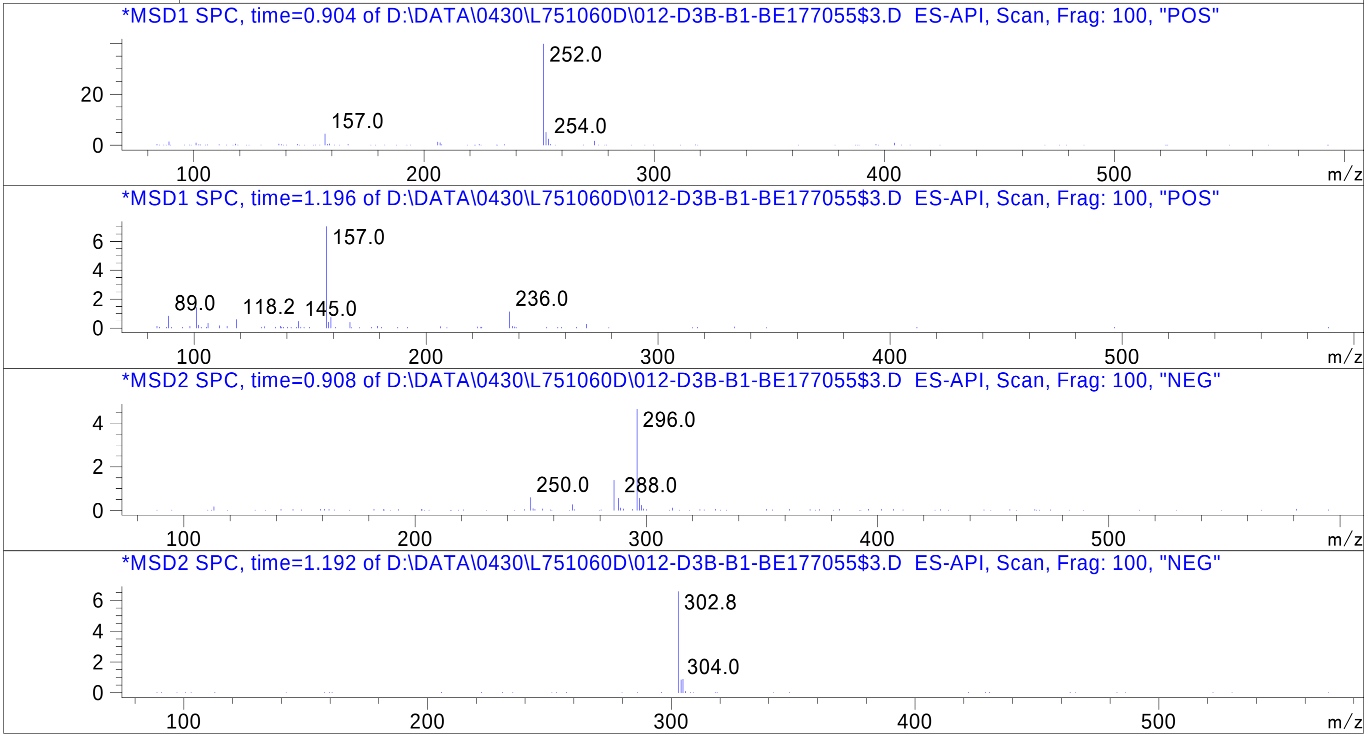


Inj.Date 4/29/2024 H -15-

C:\Chem32\1\Data\04_29\L751060D\SUPOR_30.M

### Z2274502603.PDF

MaxPeak: 100.00%
Ret_Time: 1.057 min

# BE177067$1

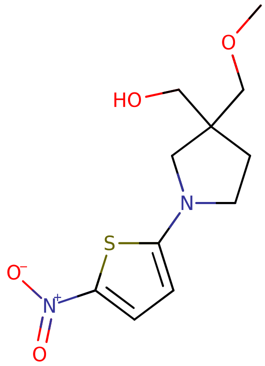


**Mol Wt** 272.32

**Exact Mass** 272.09

| # | Time | Area% |
|---|---|---|
| 1 | 1.057 | 100.00 |

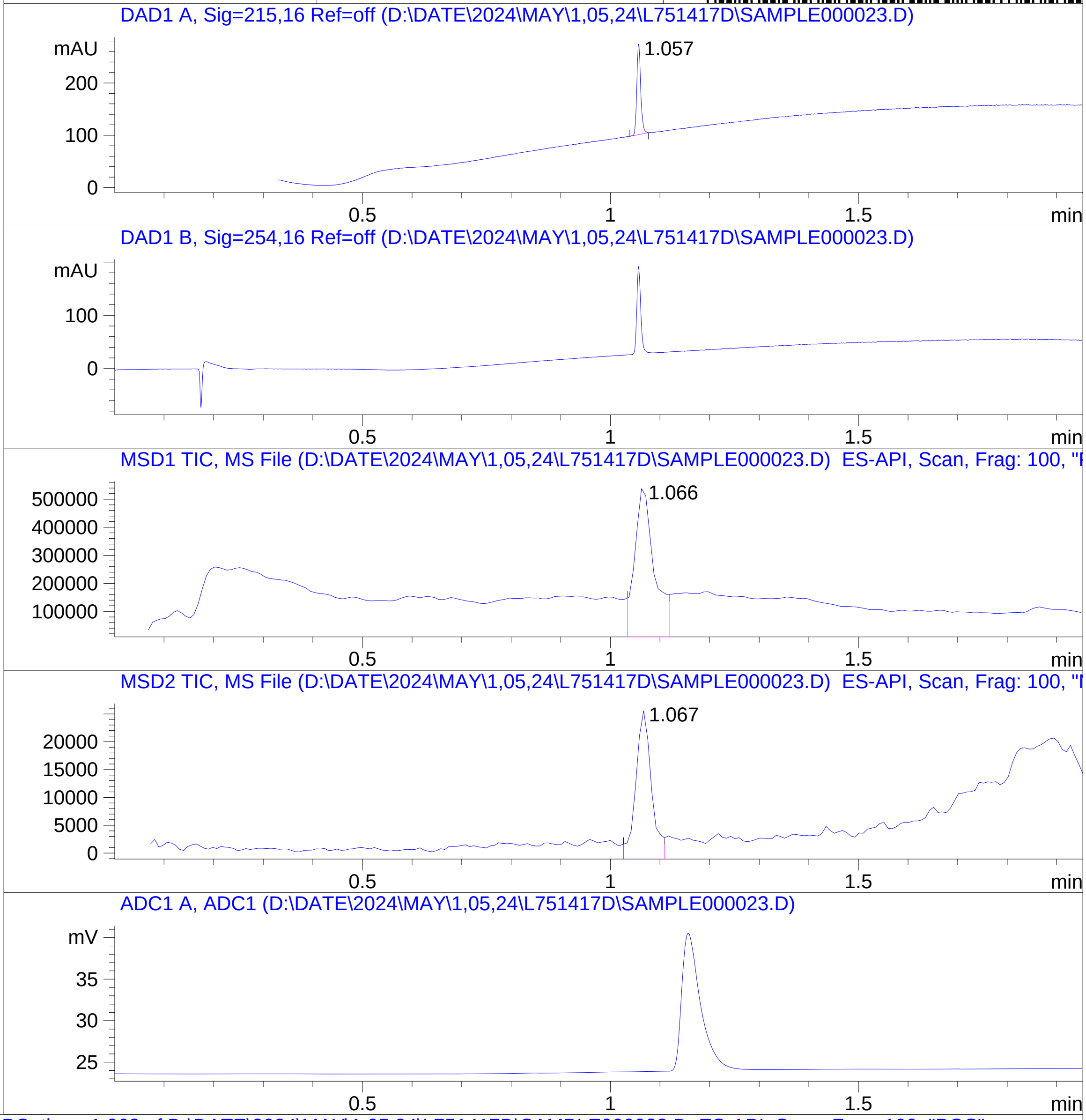


RT 1.066

RT 1.067

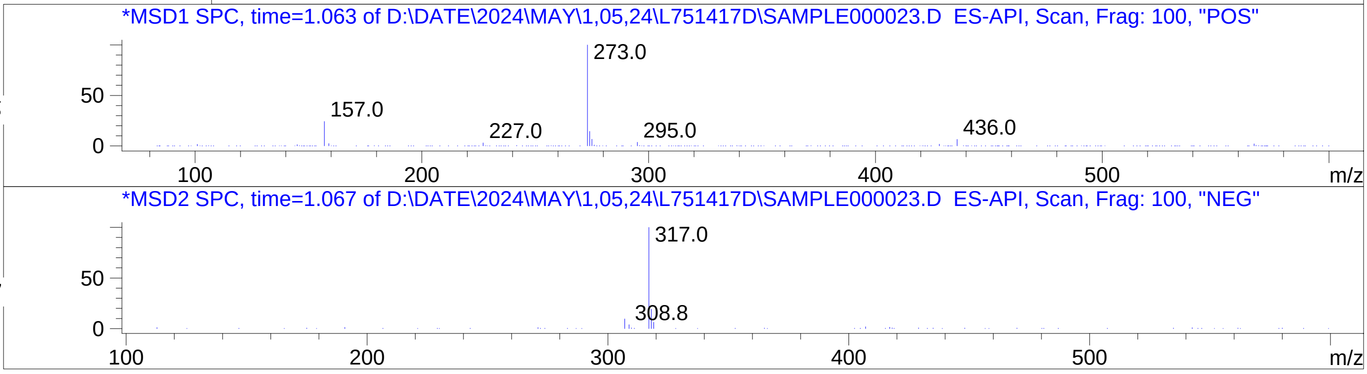

### Z2408300473.PDF

MaxPeak: 100.00%
Ret_Time: 1.082 min

# BE177001$3

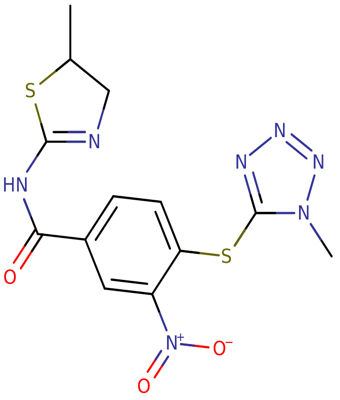


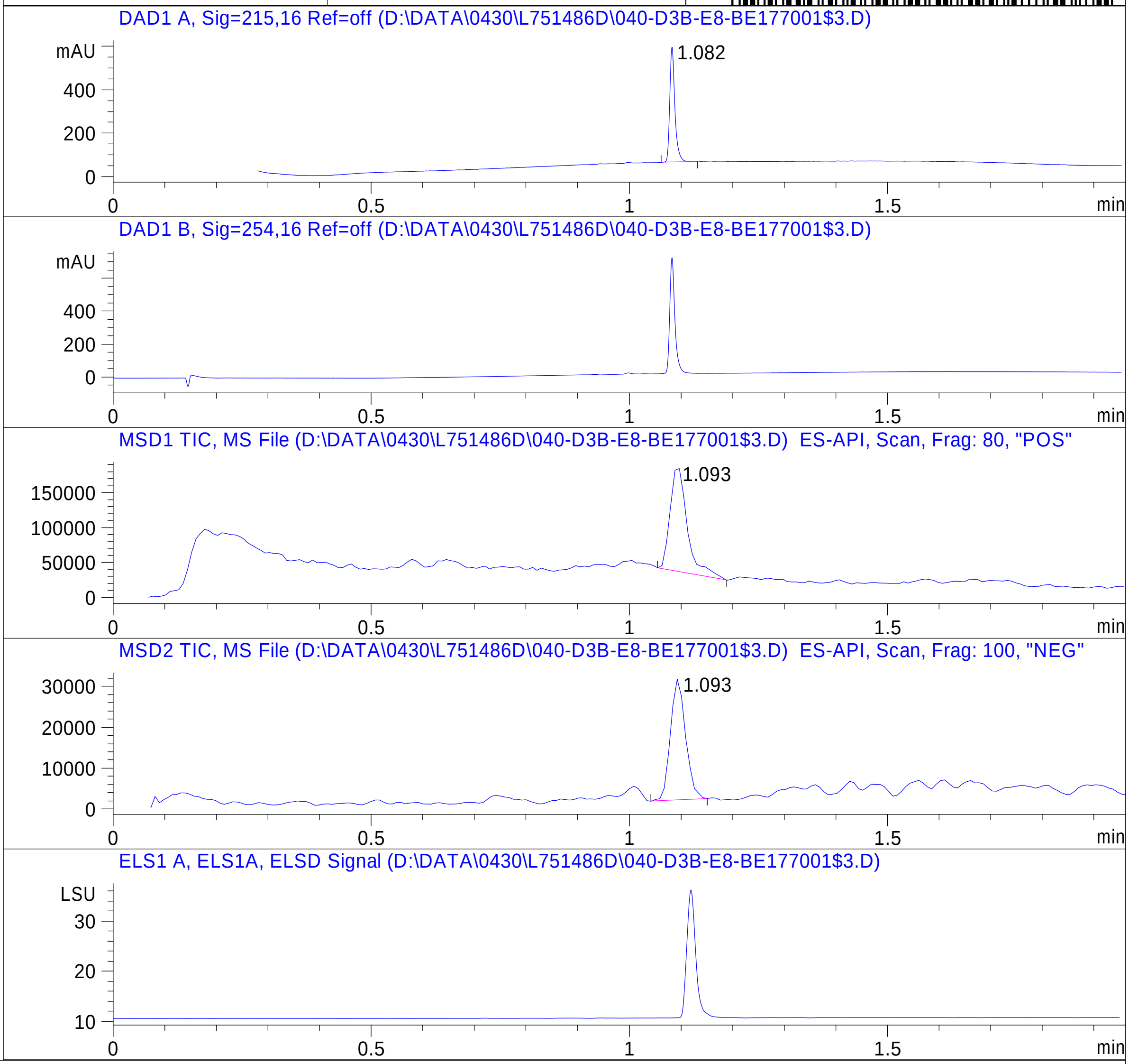


**Mol Wt** 379.42

**Exact Mass** 379.04

| # | Time | Area% |
|---|---|---|
| 1 | 1.082 | 100.00 |

RT 1.093

RT 1.093

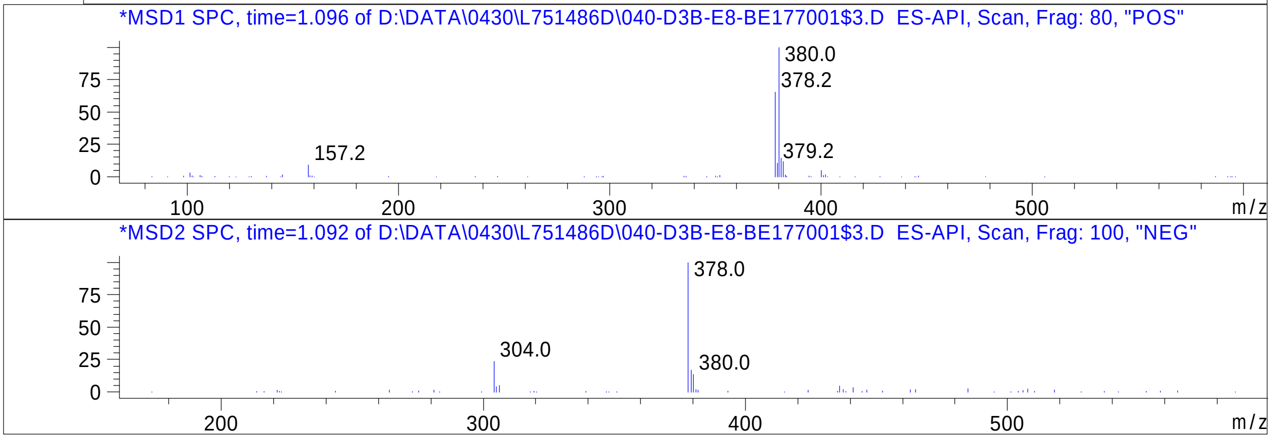

### Z2506338958.PDF

MaxPeak: 100.00%
Ret_Time: 0.786 min

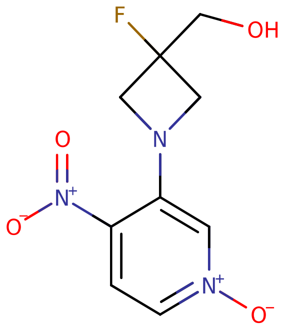


**Mol Wt** 243.19
**Exact Mass** 243.06

| # | Time | Area% |
|---|---|---|
| 1 | 0.786 | 100.00 |

# BE177081$2

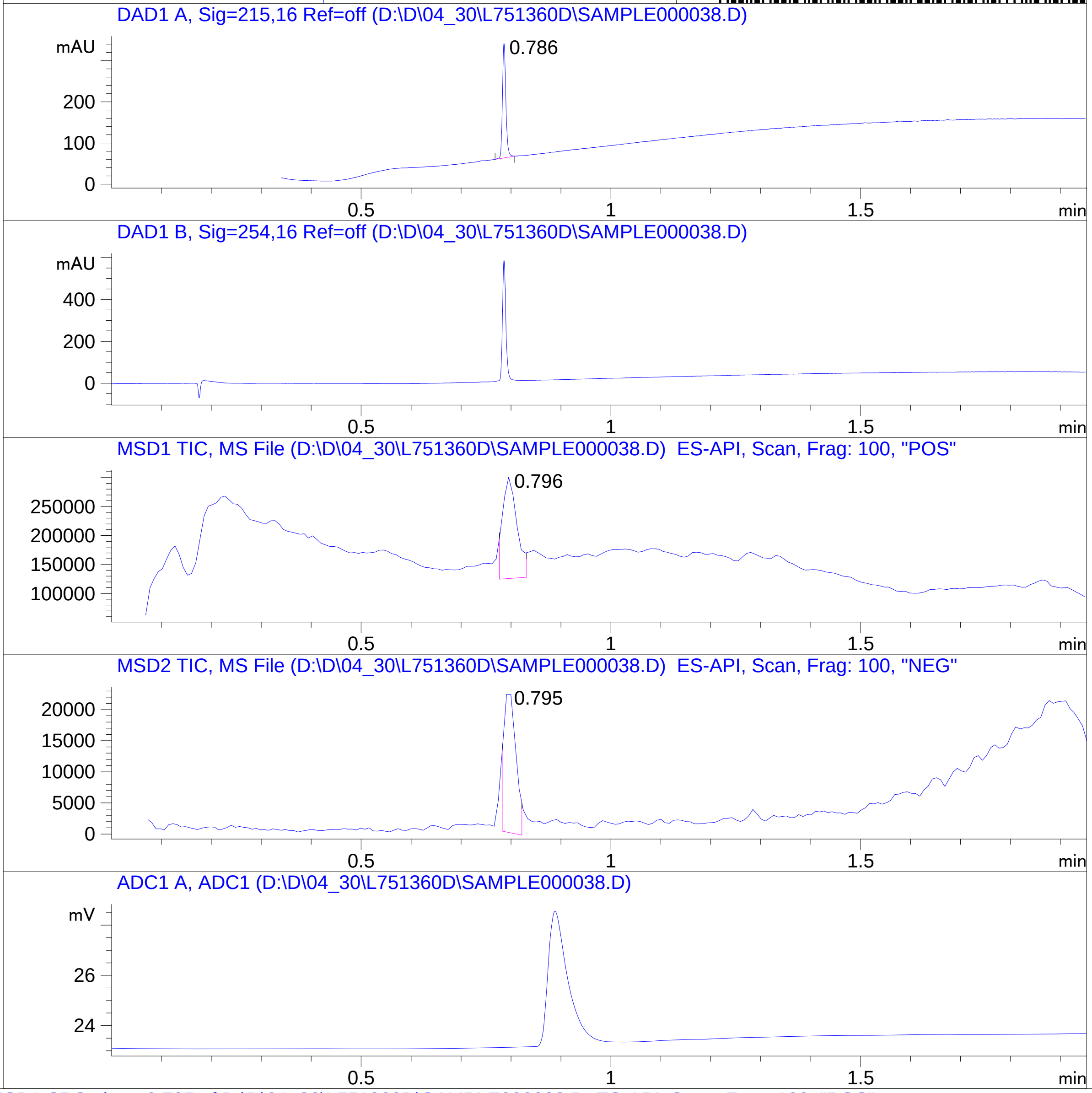


RT 0.796

RT 0.795

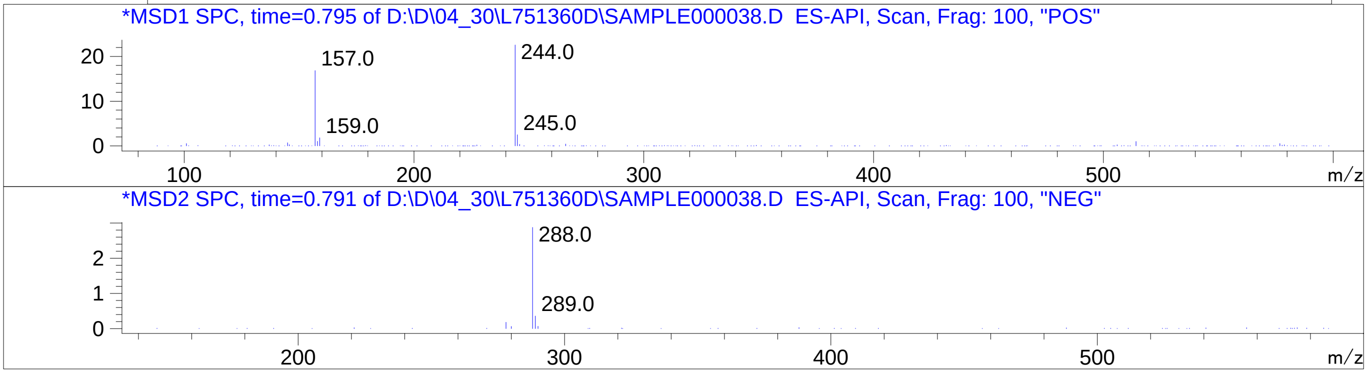


Inj.Date 4/30/2024 N
C:\CHEM32\1\METHODS\SUPOR_30.M

### Z2506437728.PDF

MaxPeak: 100.00%
Ret_Time: 1.075 min

# BE177011$2

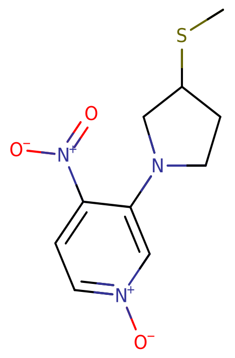


**Mol Wt** 255.29

**Exact Mass** 255.07

| # | Time | Area% |
|---|---|---|
| 1 | 1.075 | 100.00 |

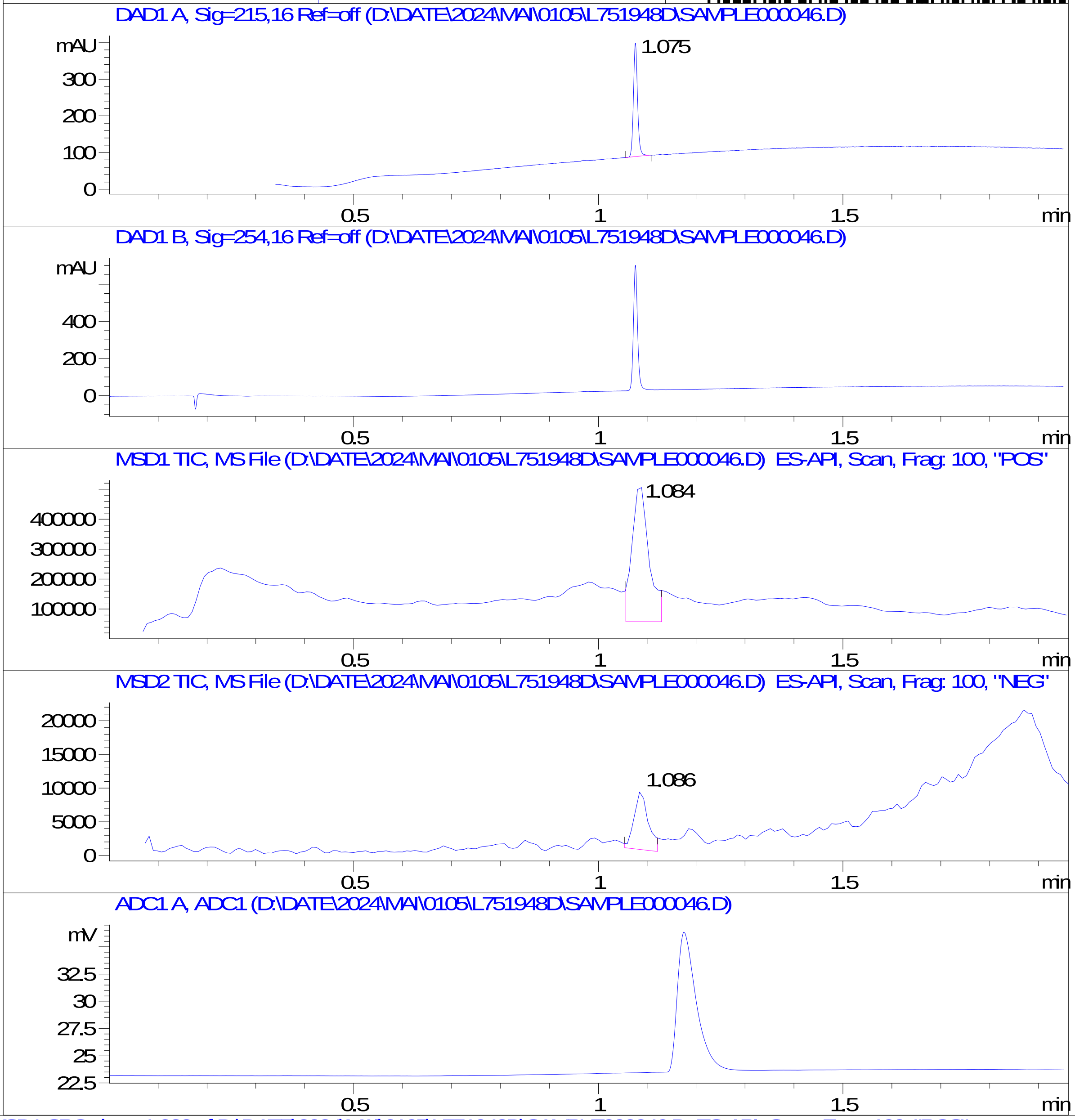


RT 1.084

RT 1.086

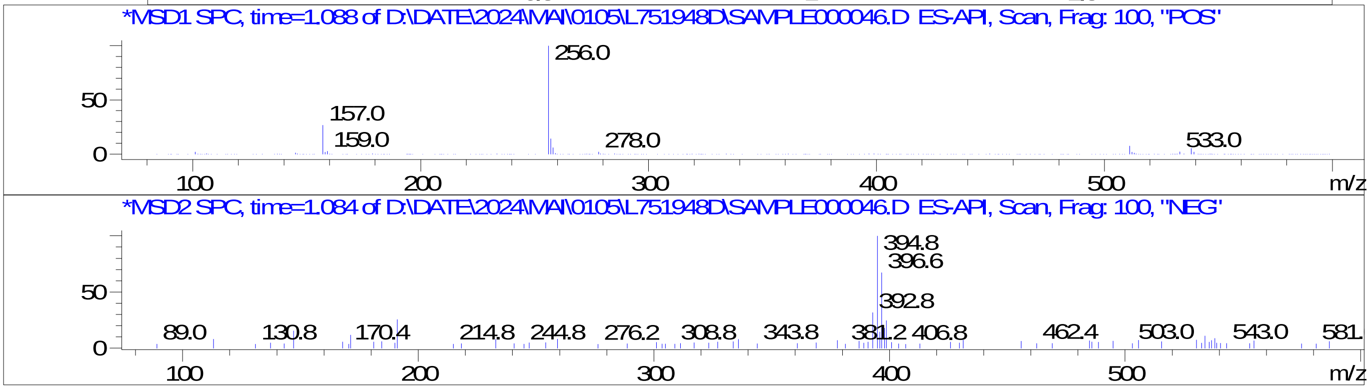

### Z2615025764.PDF

# T8886413

MaxPeak: 100.00%
Ret_Time: 0.891 min

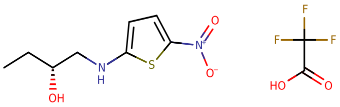

**Mol Wt** 330.28
**Exact Mass** 216.06

| # | Time | Area% |
|---|---|---|
| 1 | 0.891 | 100.00 |

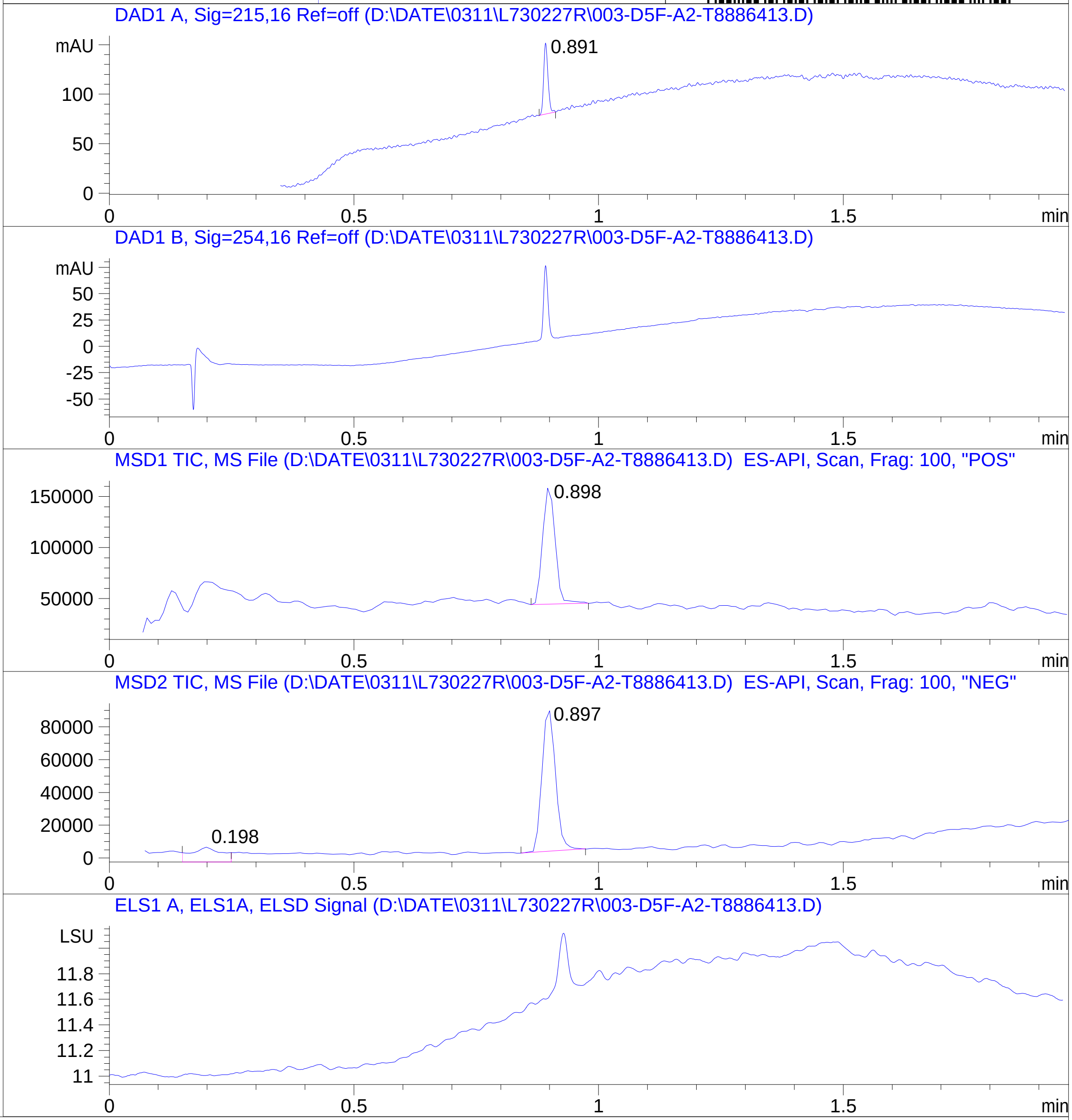


RT 0.898

RT 0.198

RT 0.897

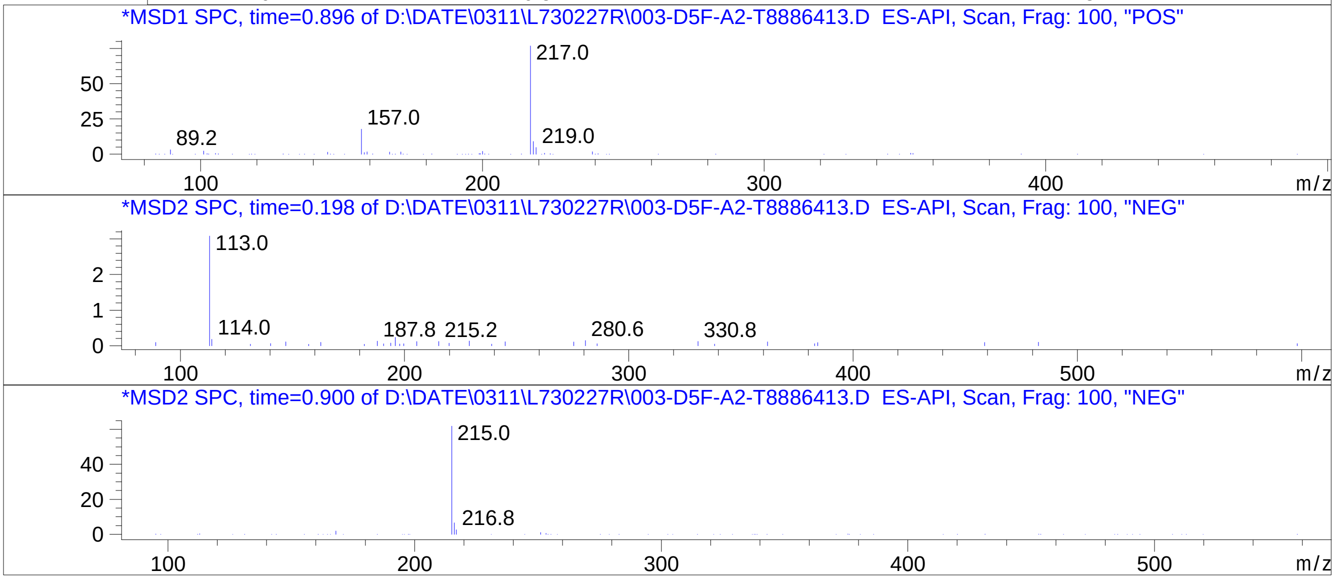

### Z2615977885.PDF

MaxPeak: 100.00%
Ret_Time: 1.013 min

# BE333773$1

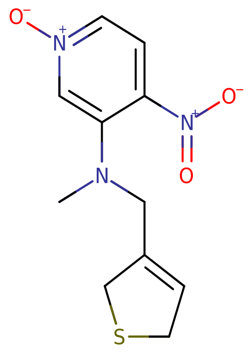


**Mol Wt** 267.3

**Exact Mass** 267.07

| # | Time | Area% |
|---|---|---|
| 1 | 1.013 | 100.00 |

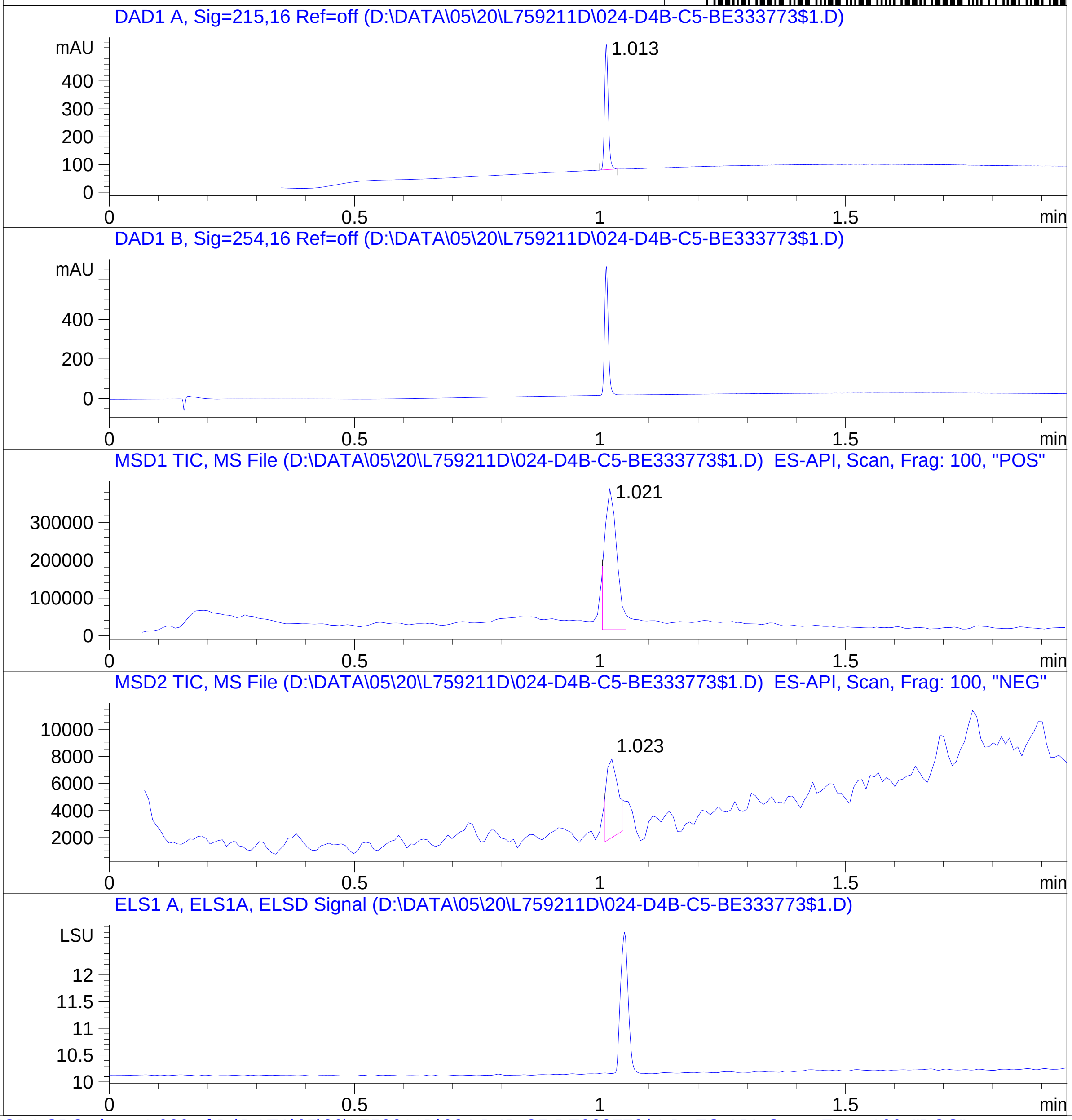


RT 1.021

RT 1.023

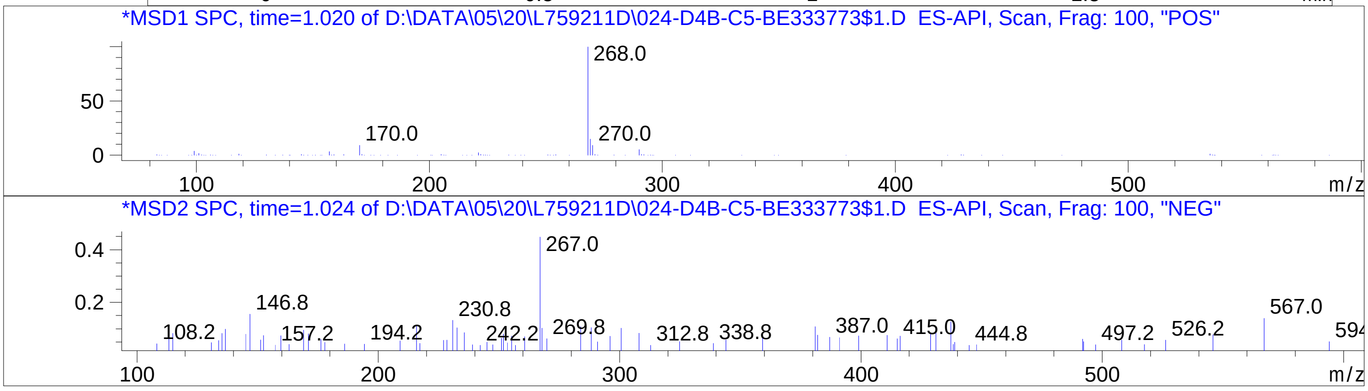


Inj.Date 5/20/2024 CH -18-

C:\Chem32\1\Data\05_20\L759211D\SUPOR_30.M

### Z2701366582.PDF

MaxPeak:  100.00%
Ret_Time: 1.154 min

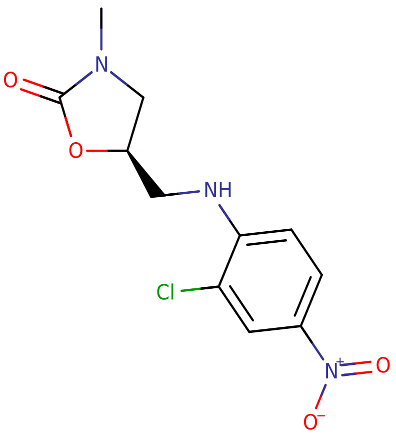


**Mol Wt** 285.68

**Exact Mass** 285.05

| # | Time | Area% |
|---|---|---|
| 1 | 1.154 | 100.00 |

# BE177083$3

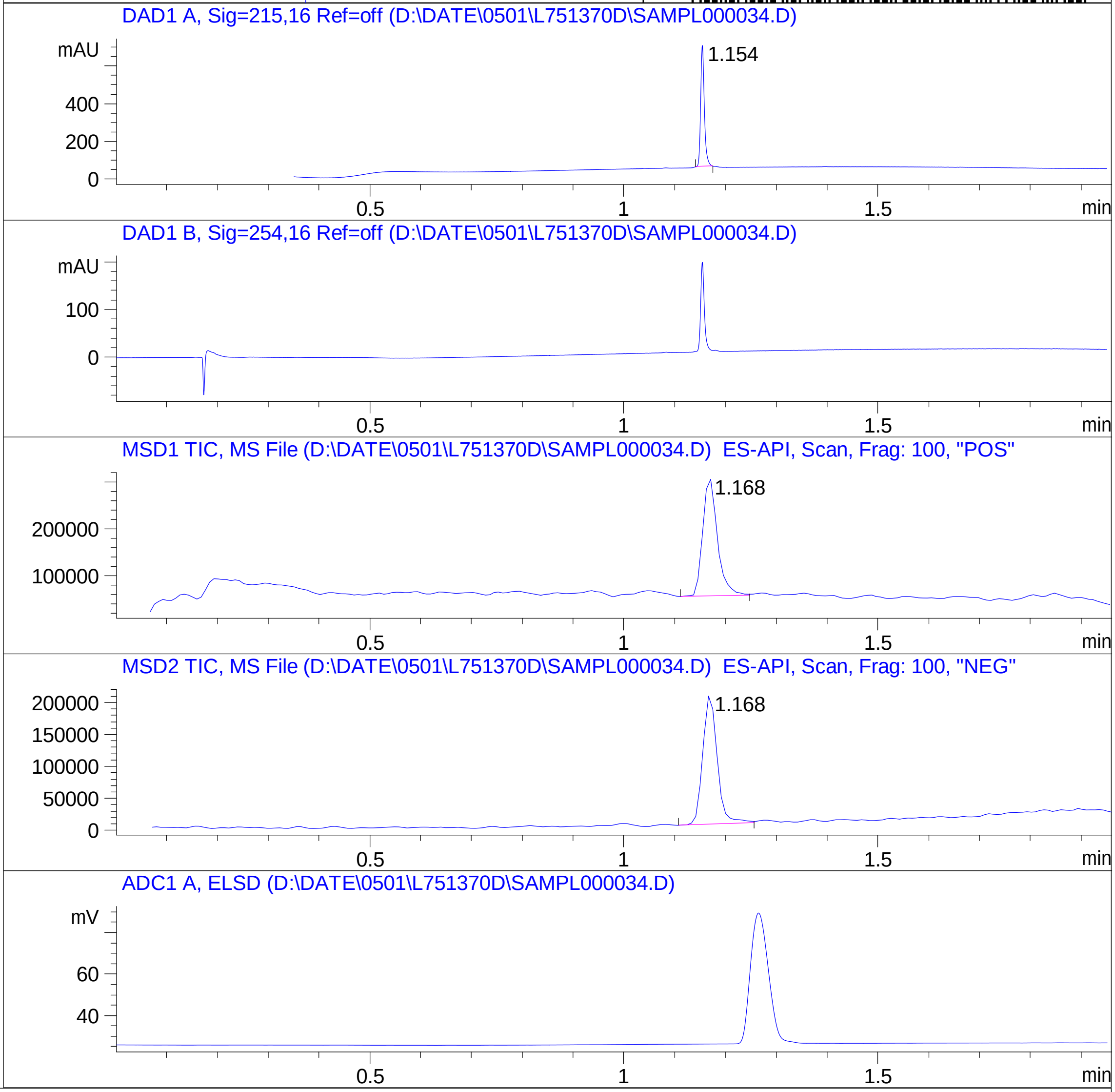


RT 1.168

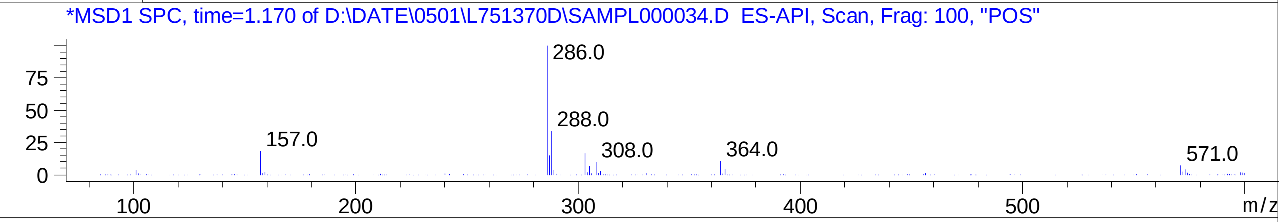


RT 1.168

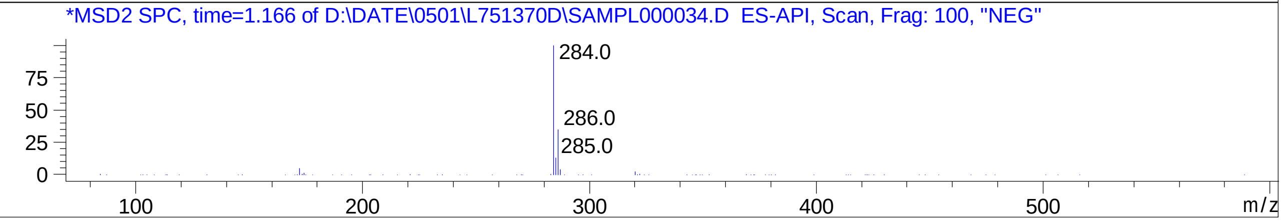

### Z2861572762.PDF

MaxPeak: 100.00%
Ret_Time: 0.894 min

# BE224652$2

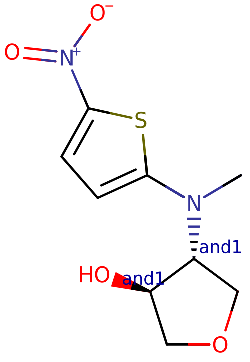


**Mol Wt** 244.27

**Exact Mass** 244.05

| # | Time | Area% |
|---|---|---|
| 1 | 0.894 | 100.00 |

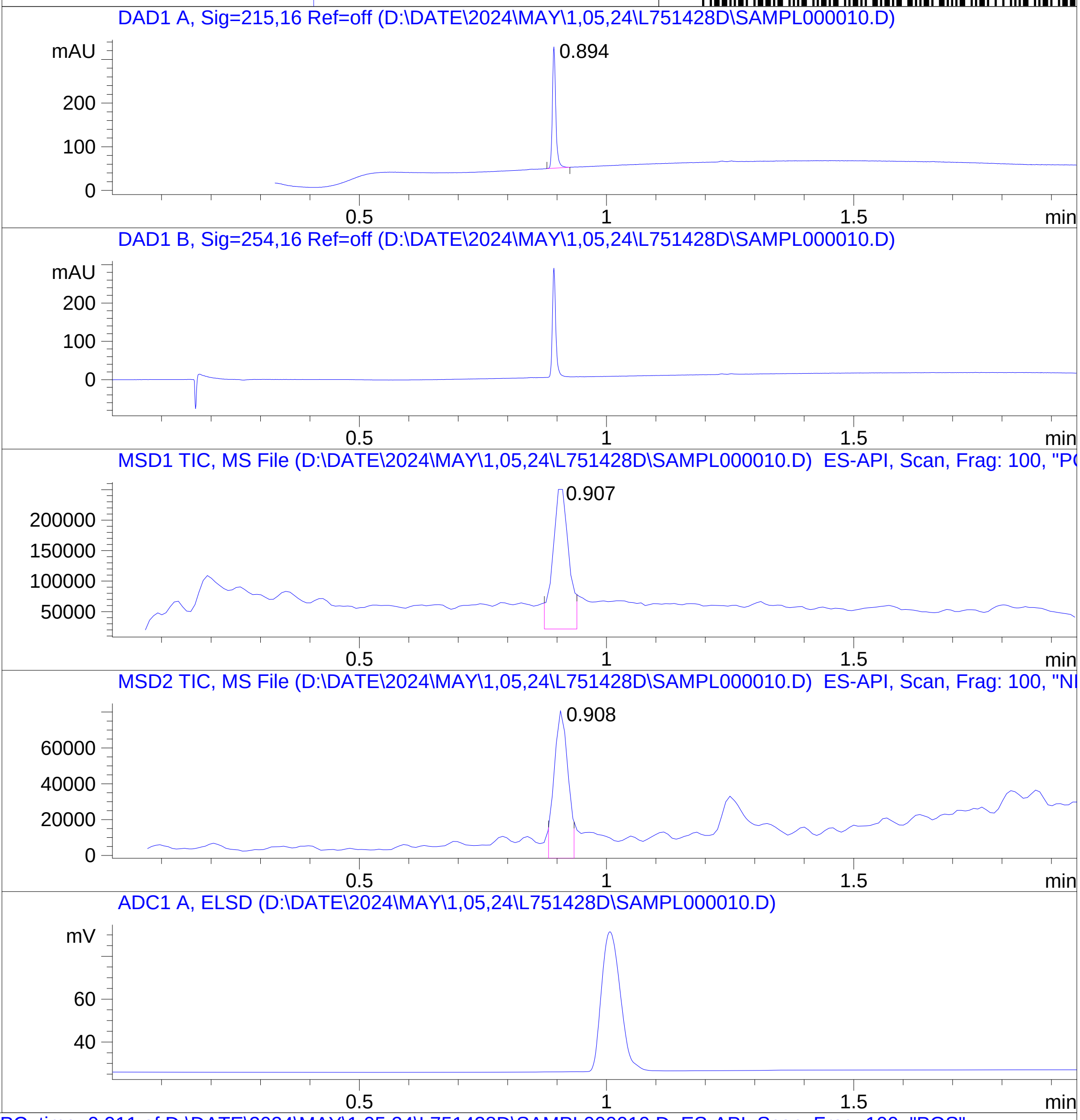


RT 0.907

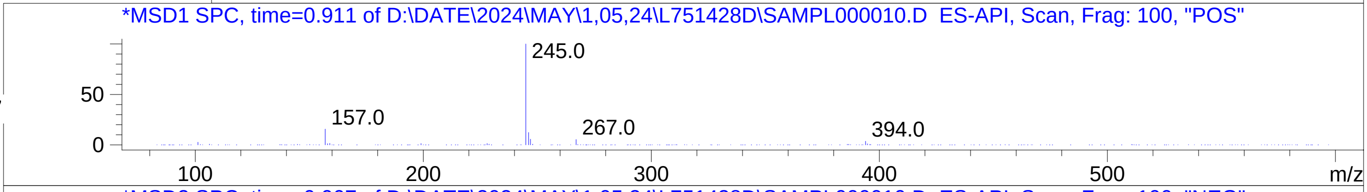


RT 0.908

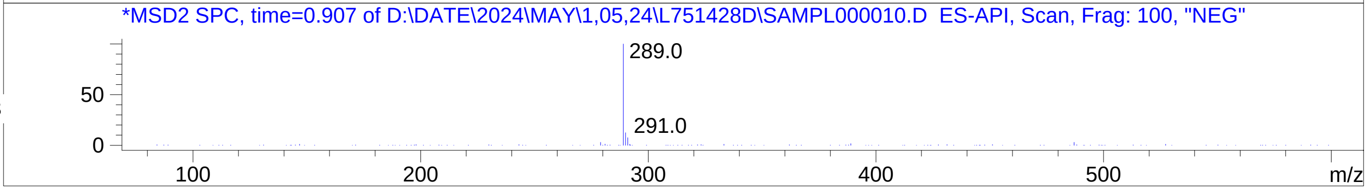

### Z2864250236.PDF

MaxPeak: 100.00%
Ret_Time: 0.957 min

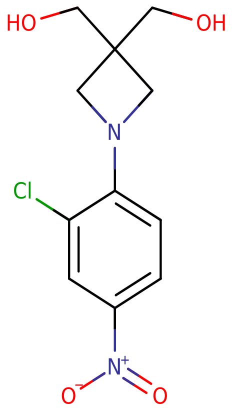

**Mol Wt** 272.68

**Exact Mass** 272.06

| # | Time | Area% |
| --- | --- | --- |
| 1 | 0.957 | 100.00 |

# BD252376$3

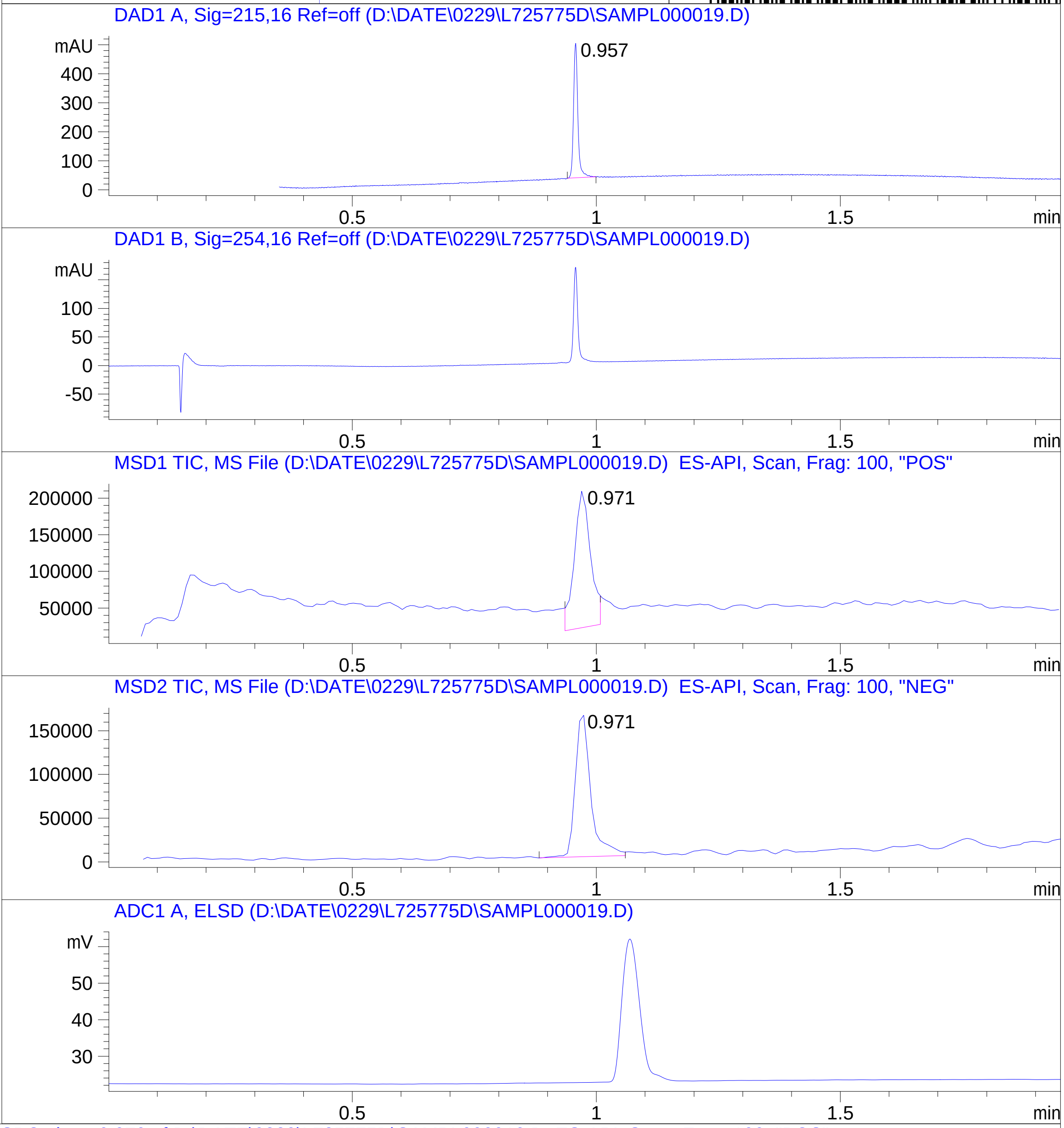


RT 0.971

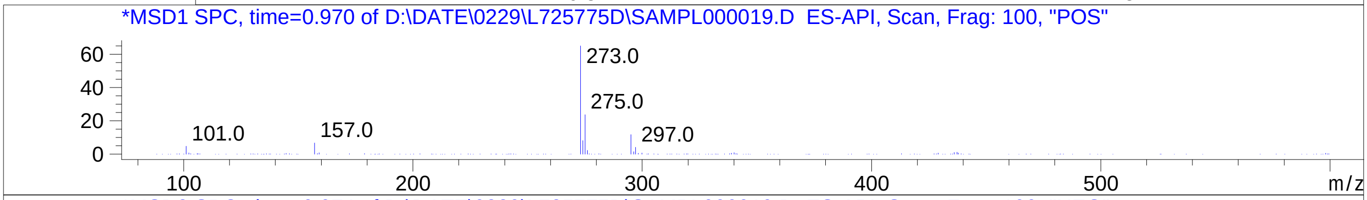


RT 0.971

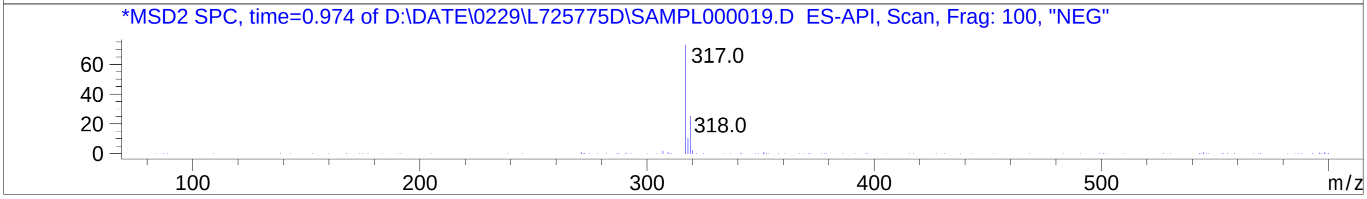

### Z2909101279.PDF

MaxPeak: 100.00%
Ret_Time: 1.122 min

# BE177012$5

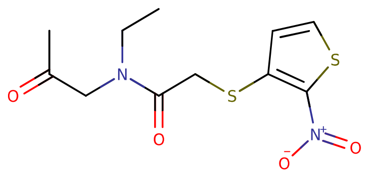

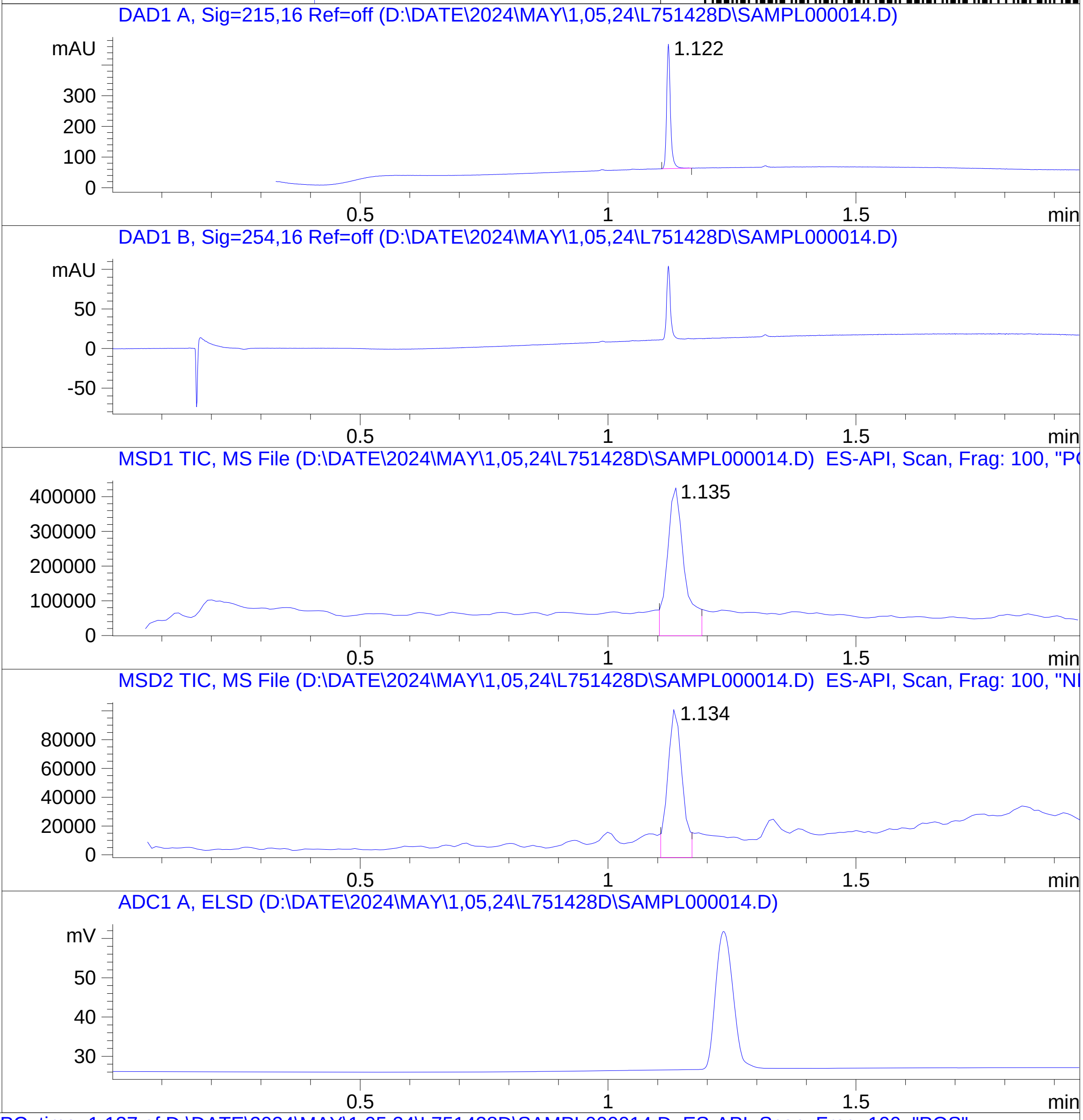


**Mol Wt** 302.37
**Exact Mass** 302.04

| # | Time | Area% |
|---|---|---|
| 1 | 1.122 | 100.00 |

RT 1.135

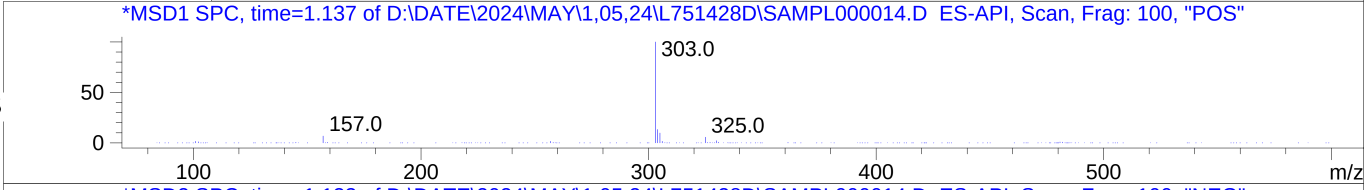


RT 1.134

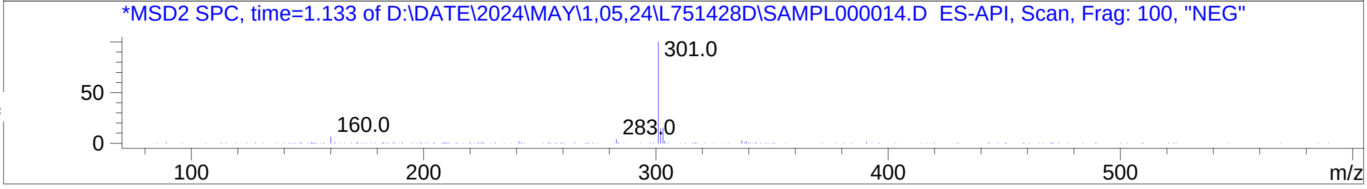

### Z2923649610.PDF

# BE224657$1

MaxPeak: 93.19%
Ret_Time: 0.935 min

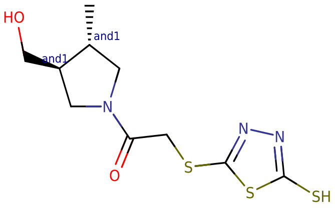


**Mol Wt** 305.44

**Exact Mass** 305.04

| # | Time | Area% |
|---|---|---|
| 1 | 0.935 | 93.19 |
| 2 | 0.971 | 4.06 |
| 3 | 1.521 | 2.75 |

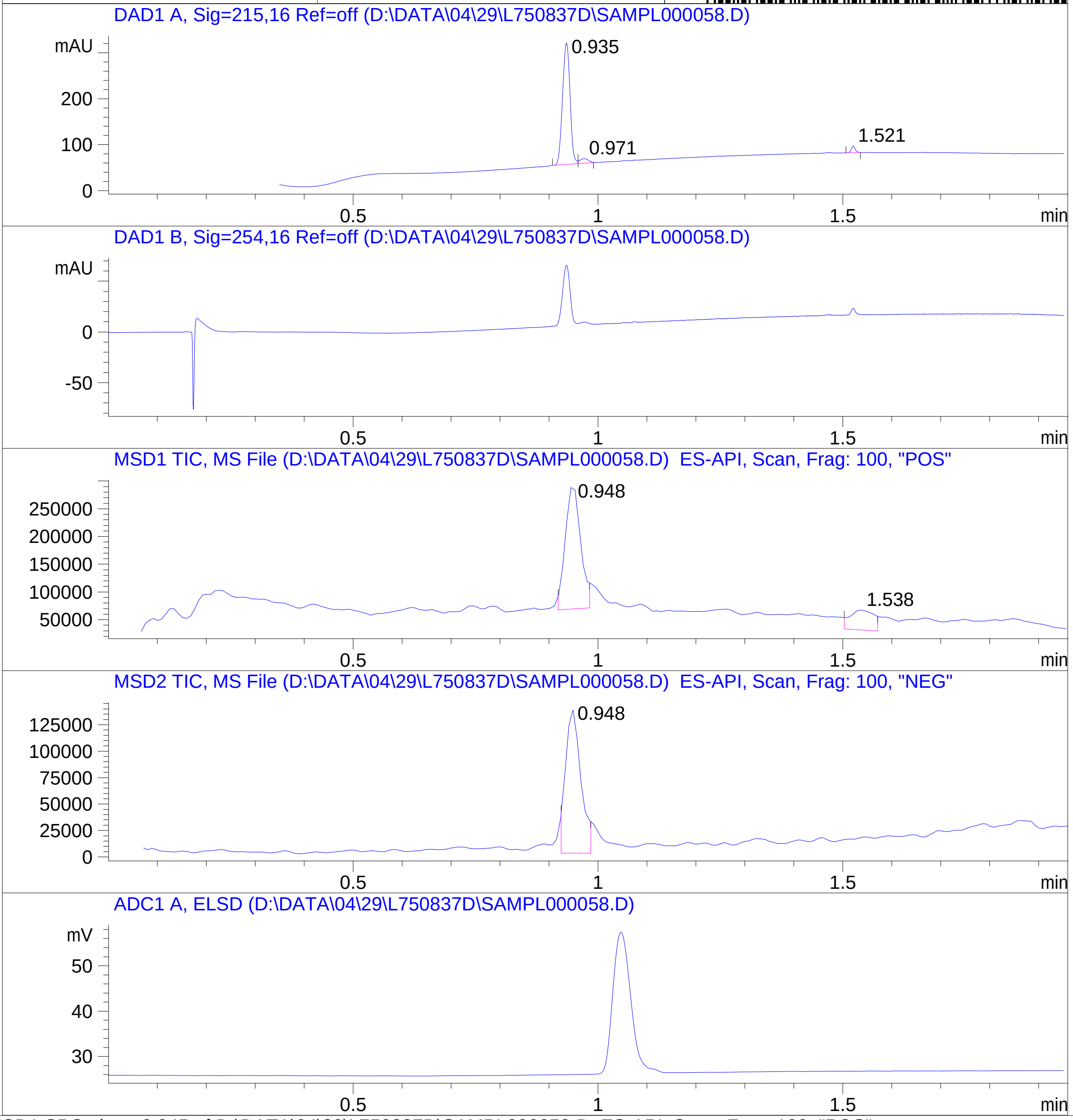


RT 0.948

RT 1.538

RT 0.948

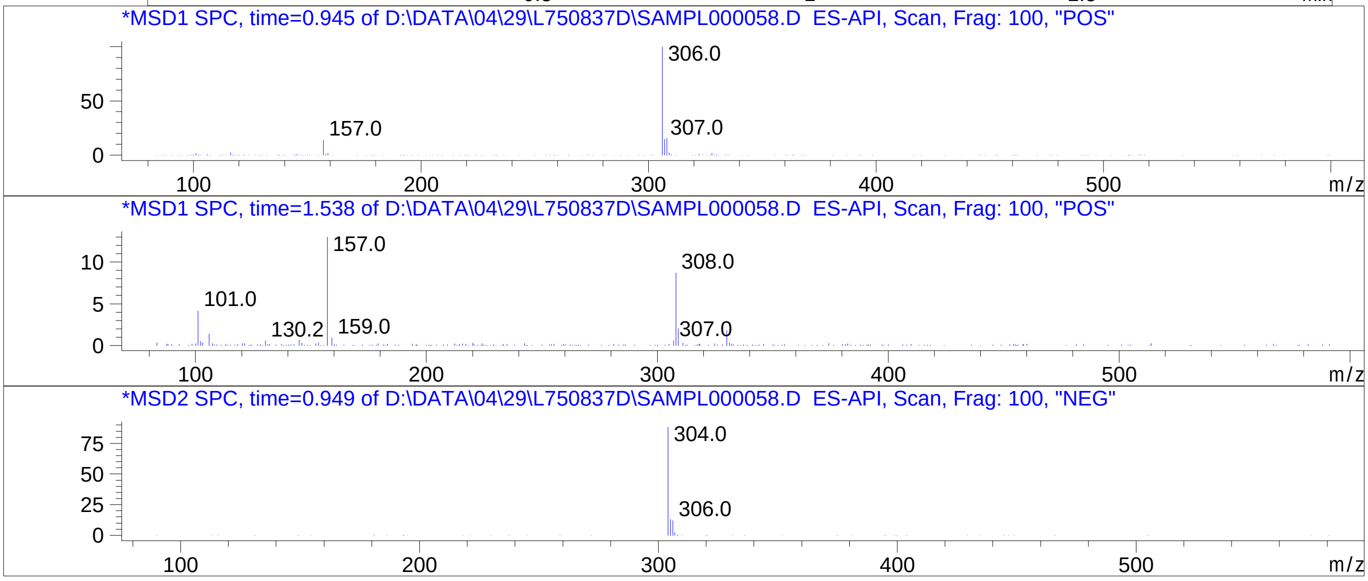


Inj.Date 4/29/2024 CH -6-
C:\CHEM32\1\METHODS\SUPOR_30.M

### Z3080120329.PDF

# BE224653$2

MaxPeak: 100.00%
Ret_Time: 0.983 min

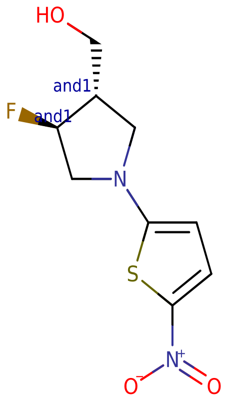


**Mol Wt** 246.26

**Exact Mass** 246.05

| # | Time | Area% |
| --- | --- | --- |
| 1 | 0.983 | 100.00 |

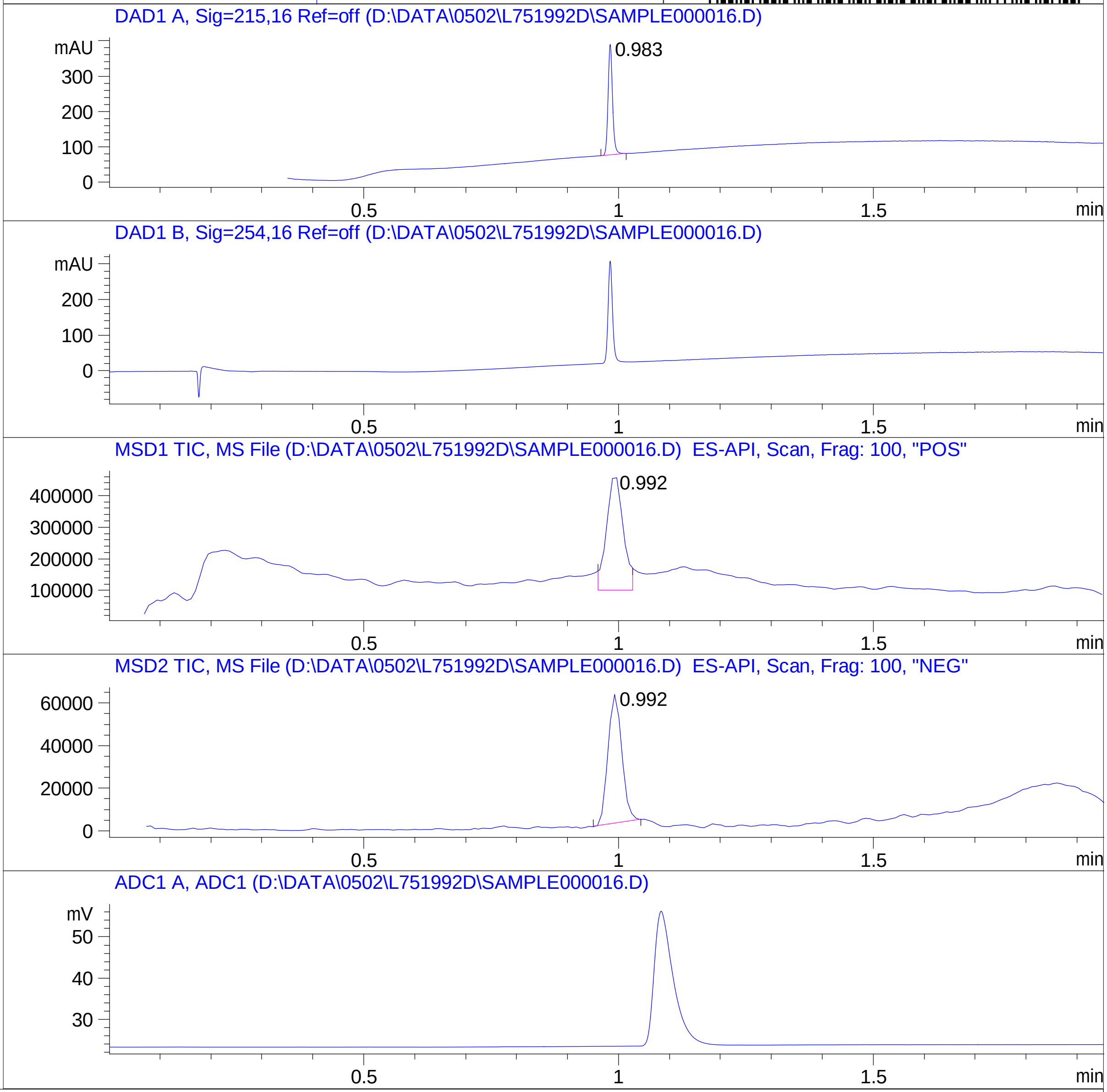


RT 0.992

RT 0.992

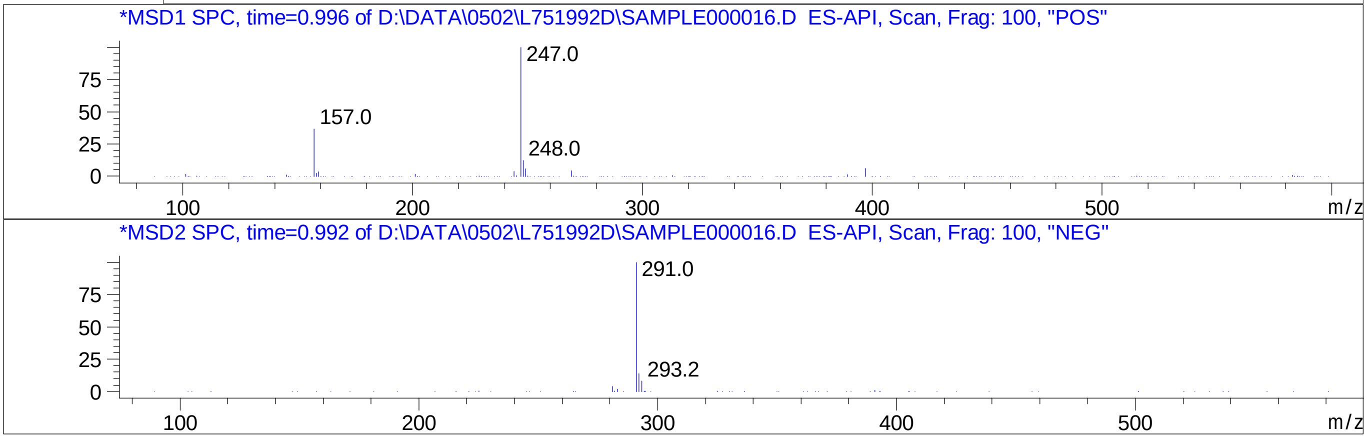


Inj.Date 5/1/2024 VB 

C:\CHEM32\1\METHODS\SUPOR_30.M

### Z3395690423.PDF

MaxPeak: 90.95%
Ret_Time: 0.781 min

# BE177084$7

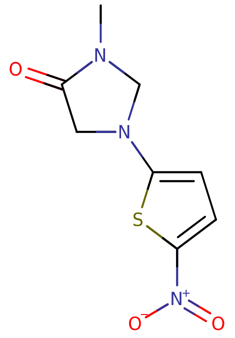


**Mol Wt** 227.24

**Exact Mass** 227.03

| # | Time | Area% |
|---|---|---|
| 1 | 0.781 | 90.95 |
| 2 | 1.197 | 2.93 |
| 3 | 1.289 | 6.12 |

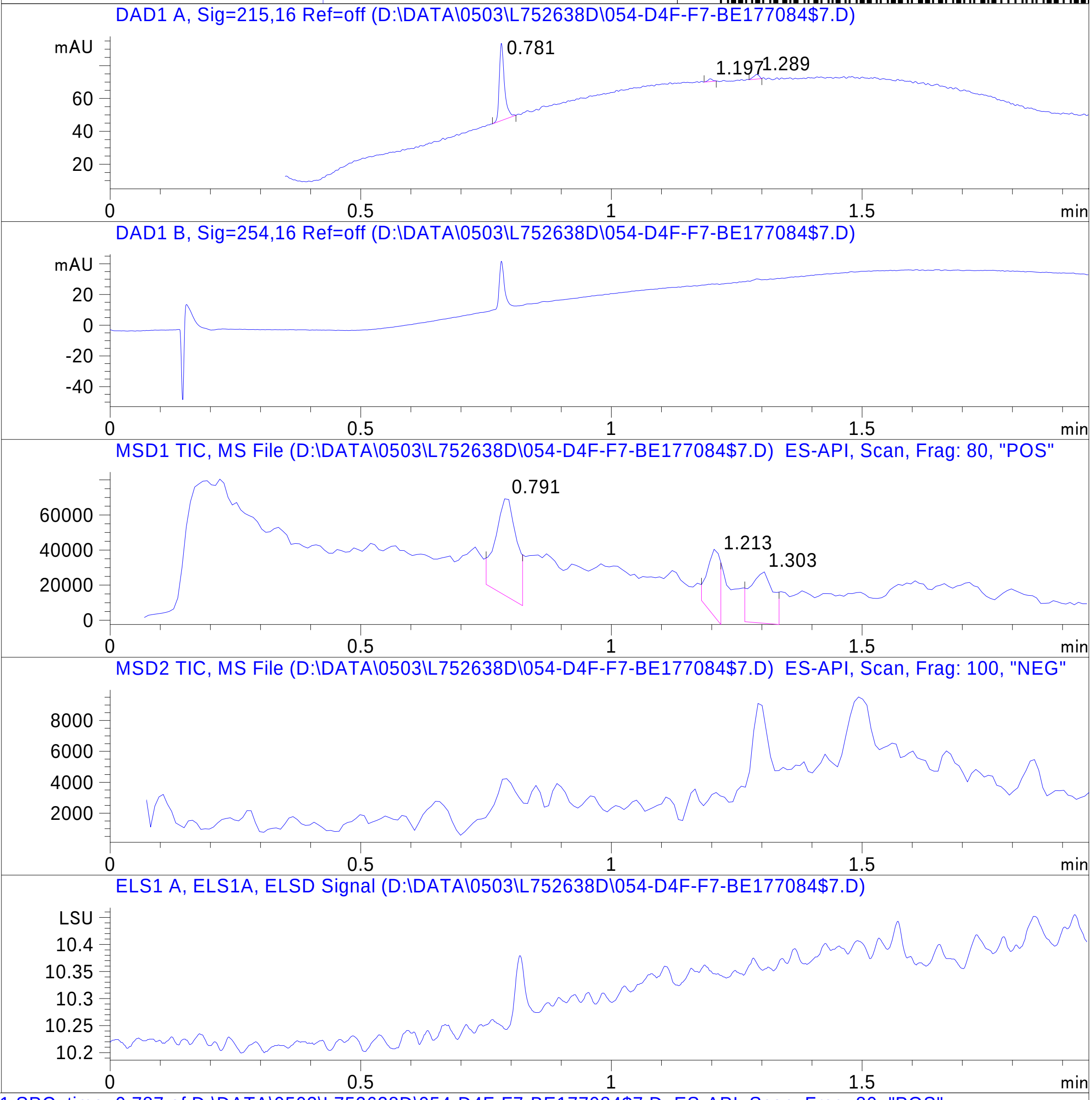


RT 0.791

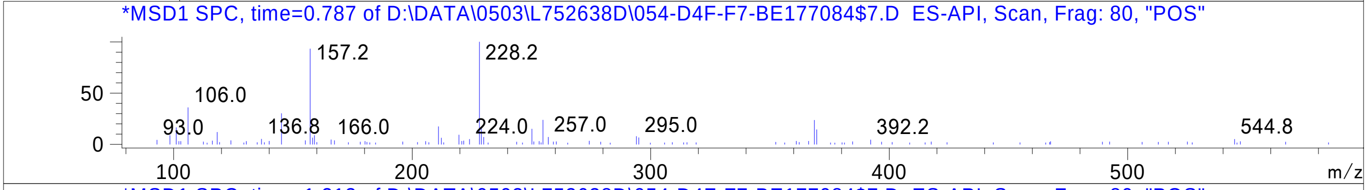


RT 1.213

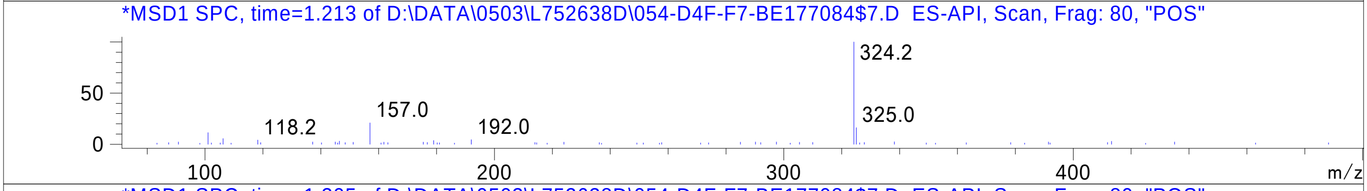


RT 1.303

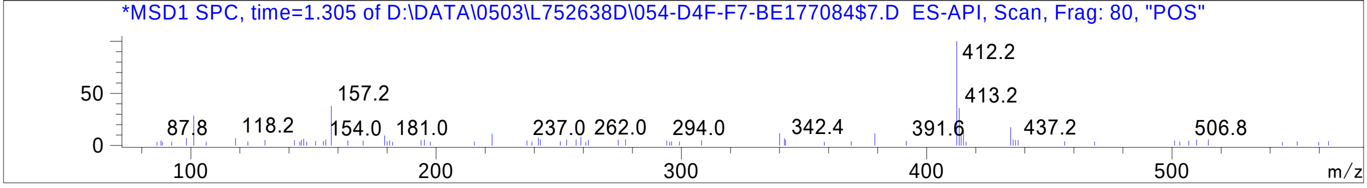

### Z3605614073.PDF

MaxPeak: 100.00%
Ret_Time: 1.122 min

# BE177035$2

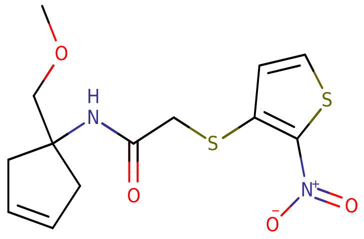


**Mol Wt** 328.41

**Exact Mass** 328.06

| # | Time | Area% |
|---|---|---|
| 1 | 1.122 | 100.00 |

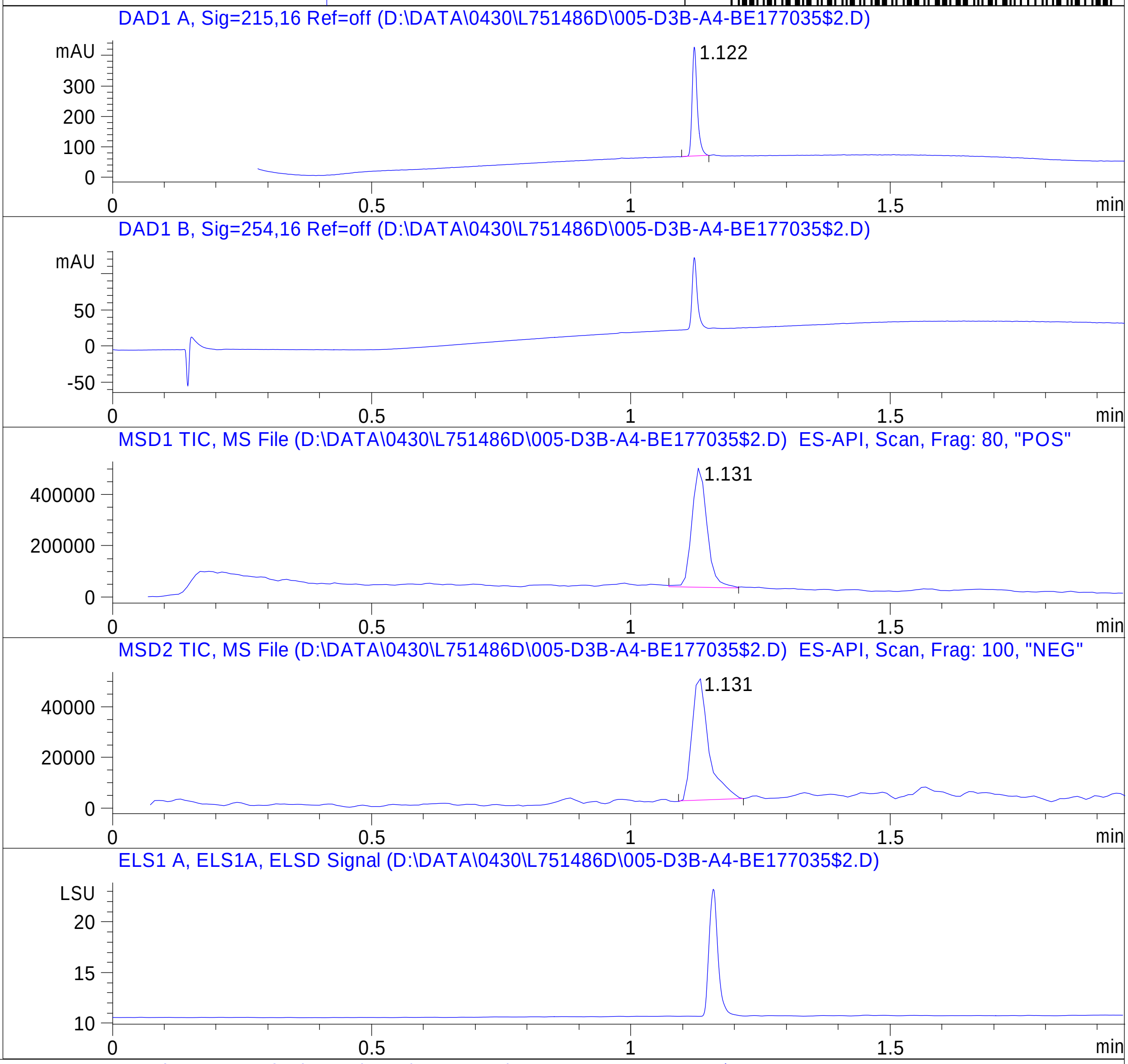


RT 1.131

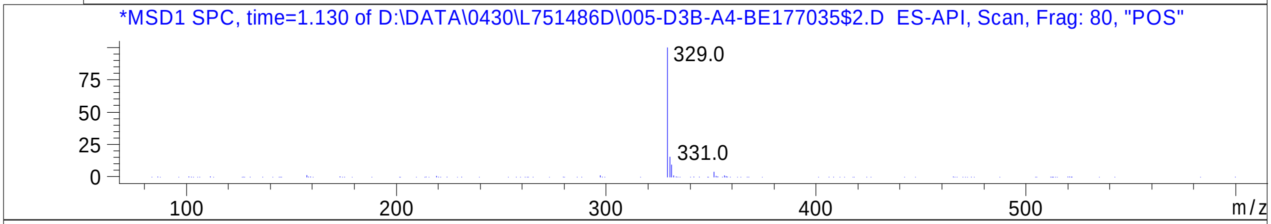


RT 1.131

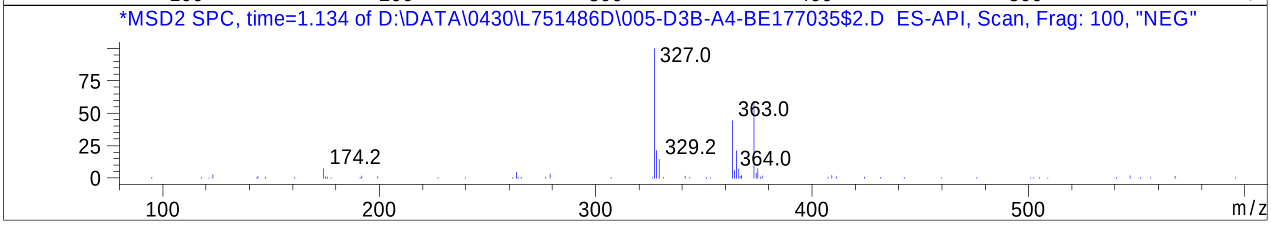


Inj.Date 4/30/2024 LB



C:\Chem32\1\Data\04_30\L751486D\SUPOR_30.M

### Z3633671332.PDF

MaxPeak:  98.38%
Ret_Time: 0.954 min

# BE224661$3

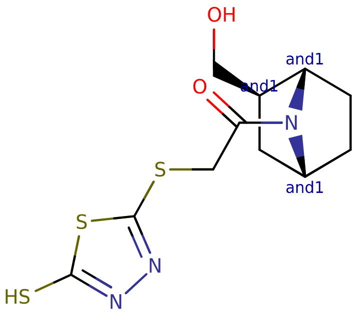


**Mol Wt** 317.45

**Exact Mass** 317.04

| # | Time | Area% |
|---|---|---|
| 1 | 0.954 | 98.38 |
| 2 | 1.003 | 1.62 |

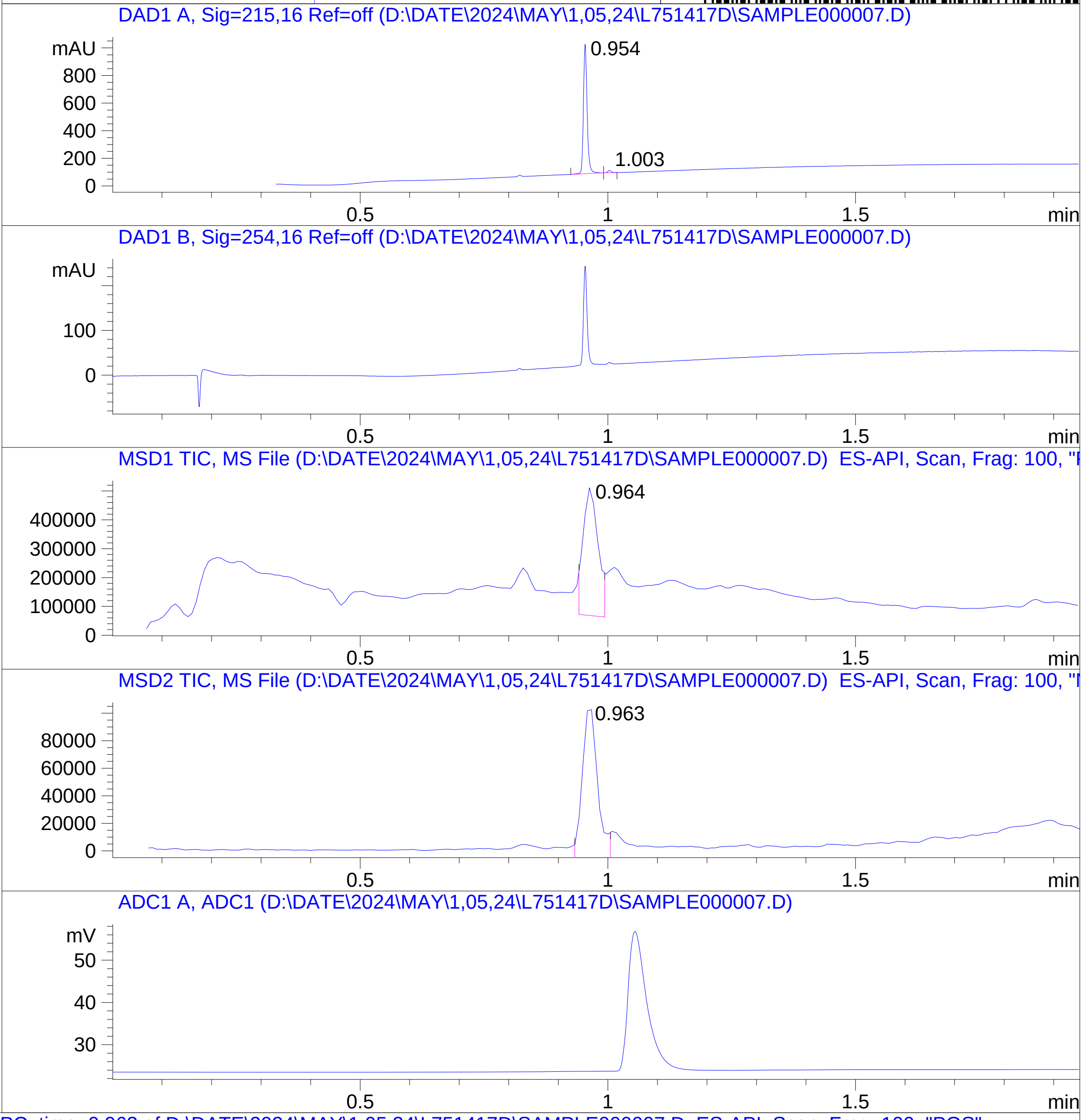


RT 0.964

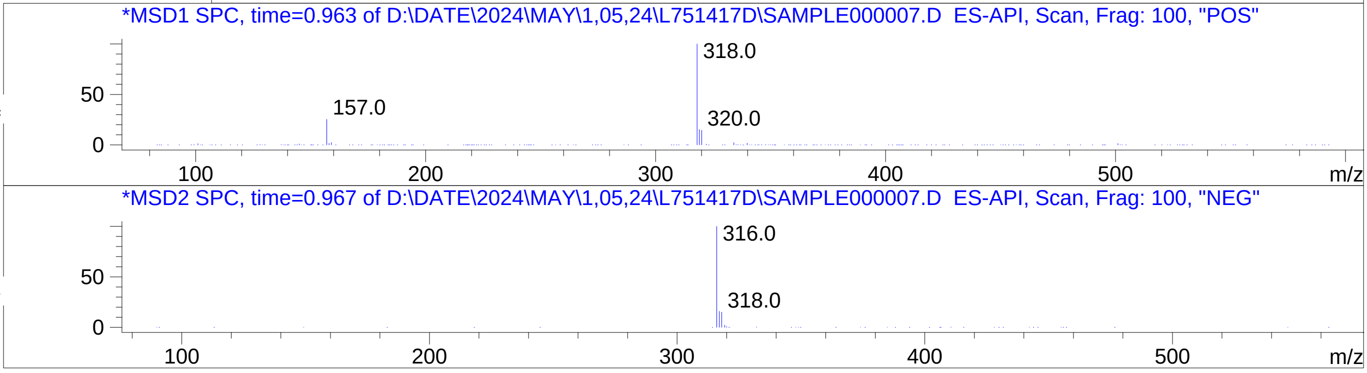


RT 0.963

Inj.Date 4/30/2024 T - 4 - ->
C:\CHEM32\1\METHODS\SUPOR_30.M

### Z3649980129.PDF

MaxPeak: 95.35%
Ret_Time: 0.635 min

# BD252392$2

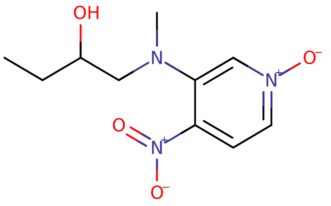


**Mol Wt** 241.24

**Exact Mass** 241.11

| # | Time | Area% |
|---|---|---|
| 1 | 0.489 | 2.17 |
| 2 | 0.576 | 2.49 |
| 3 | 0.635 | 95.35 |

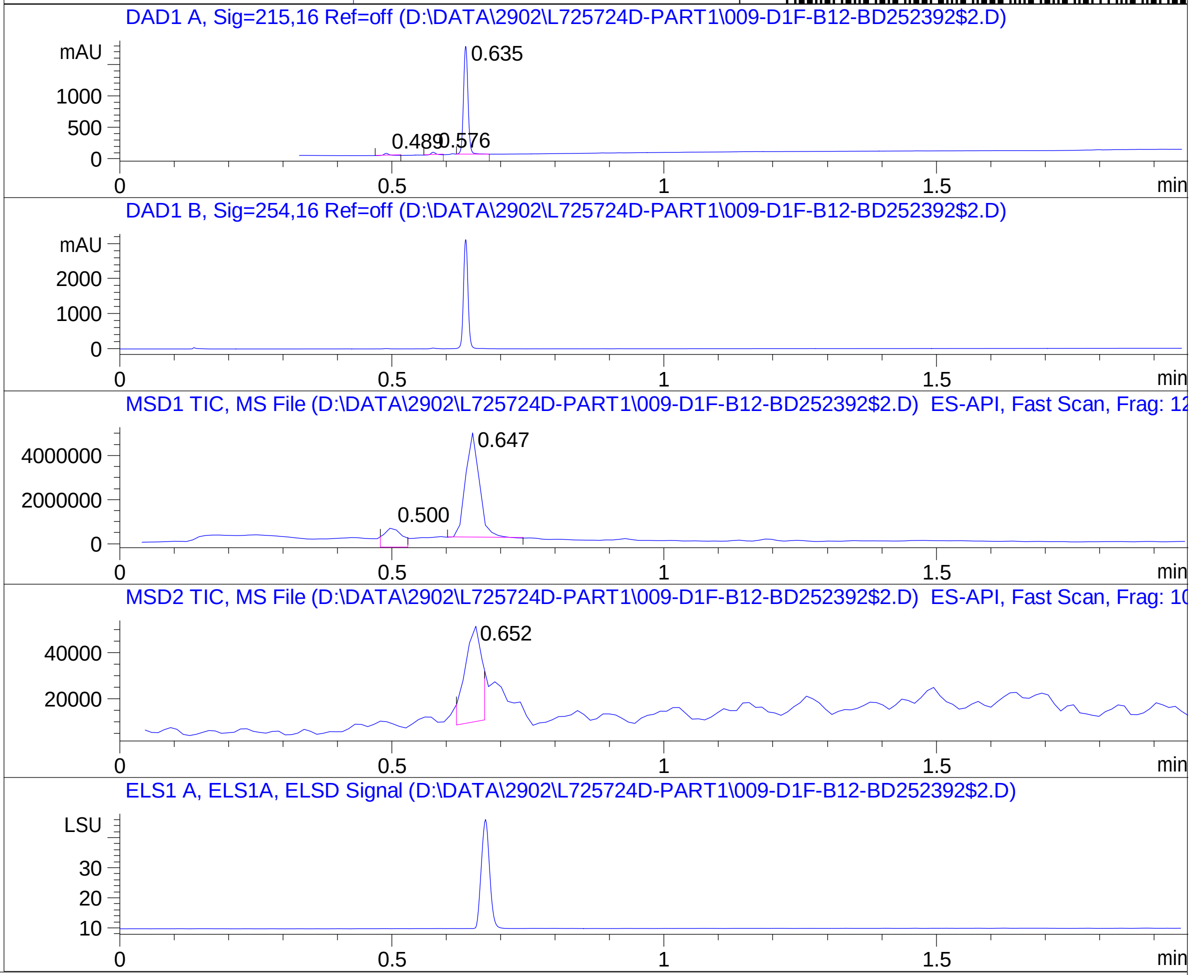


RT 0.500

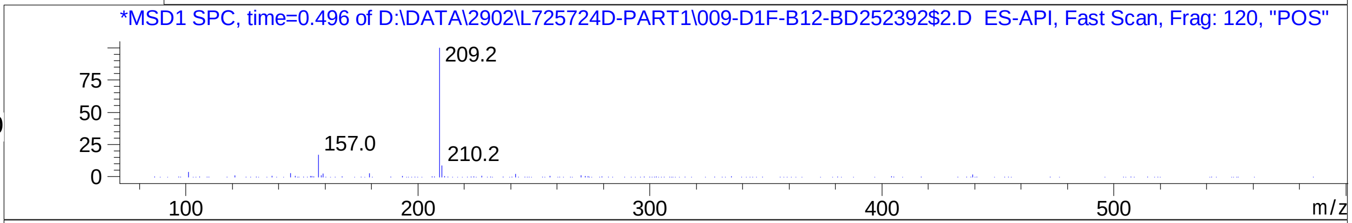


RT 0.647

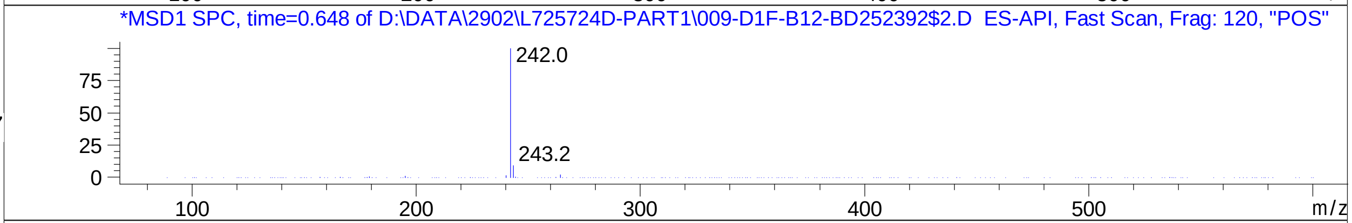


RT 0.652

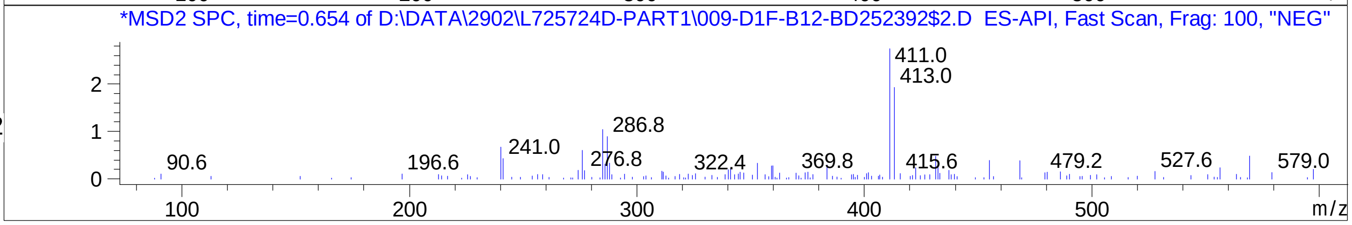

### Z3832034072.PDF

# BE177070$1

MaxPeak: 100.00%
Ret_Time: 0.851 min

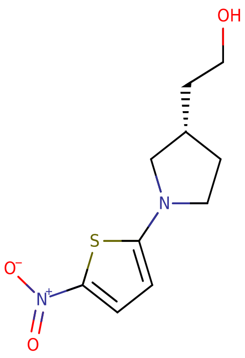


**Mol Wt** 242.3

**Exact Mass** 242.08

| # | Time | Area% |
|---|---|---|
| 1 | 0.851 | 100.00 |

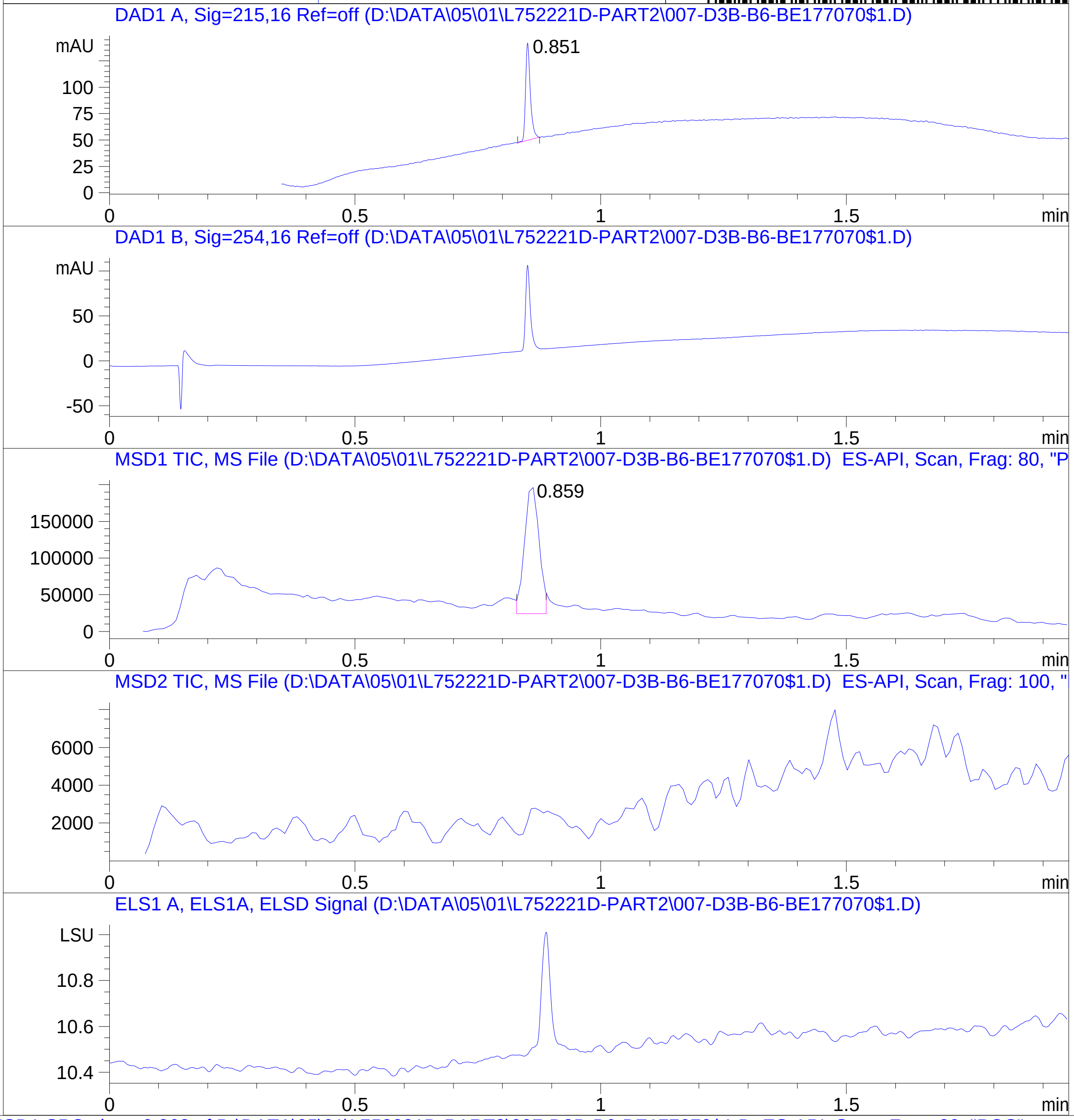


RT 0.859

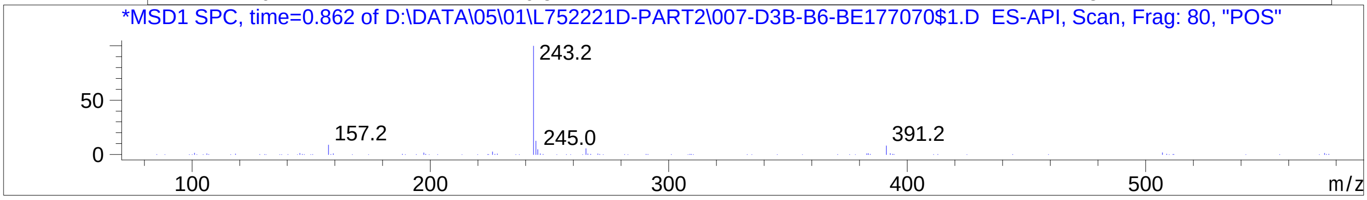


Inj.Date 5/1/2024 CH -16-
C:\Chem32\1\Data\05_01\2752221D\SUPOR_30.M

### Z4002664779.PDF

# BE177004$1

MaxPeak: 90.13%
Ret_Time: 1.070 min

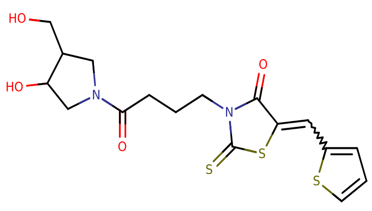


**Mol Wt** 412.55

**Exact Mass** 412.07

| # | Time | Area% |
|---|---|---|
| 1 | 1.070 | 90.13 |
| 2 | 1.158 | 9.87 |

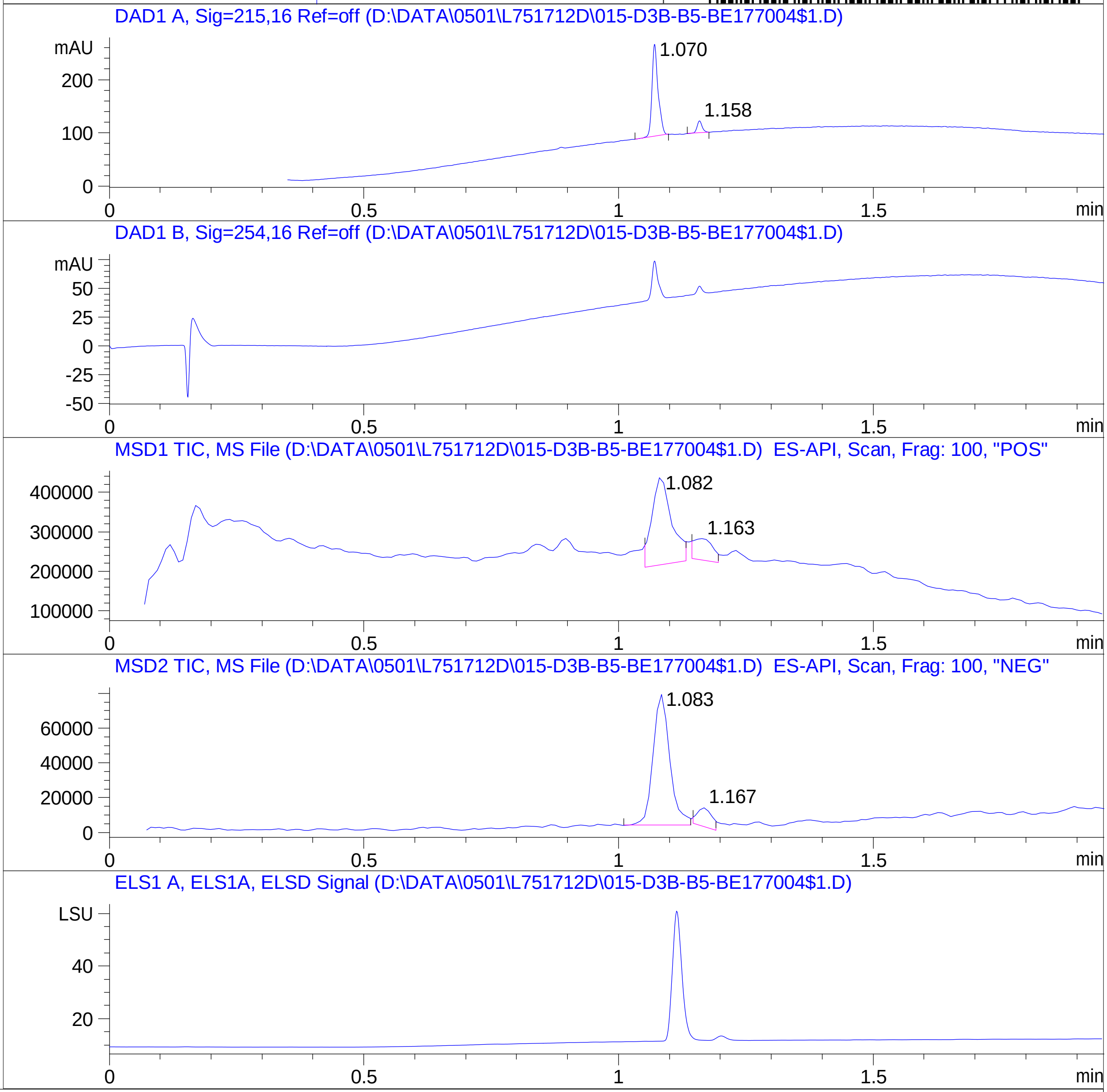


RT 1.082

RT 1.163

RT 1.083

RT 1.167

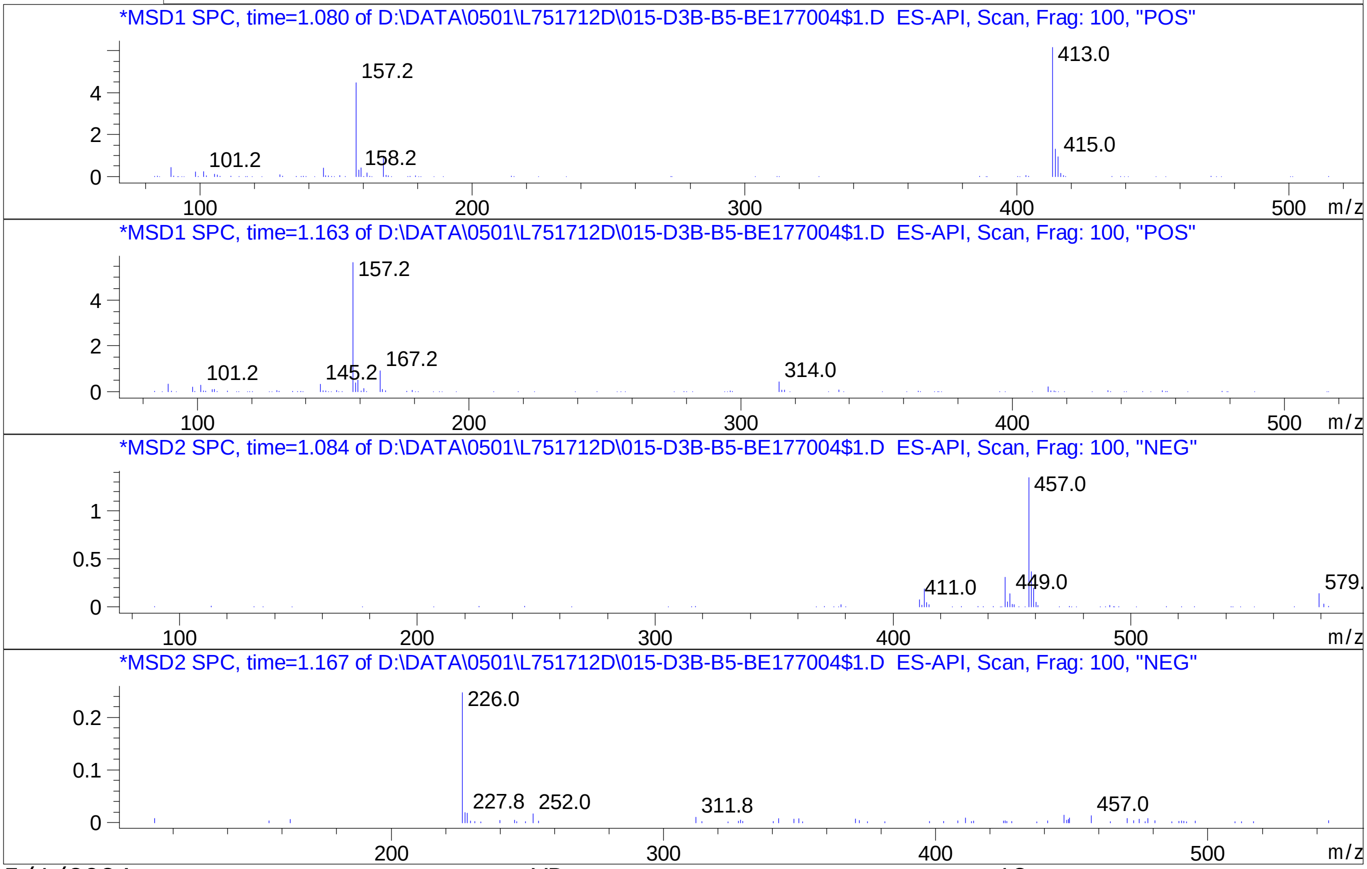


Inj.Date 5/1/2024 VB 13

C:\Chem32\1\Data\04_30\L751712D\SUPOR_30.M

### Z4137781945.PDF

MaxPeak: 100.00%
Ret_Time: 1.466 min

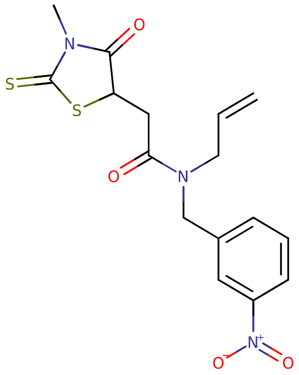

**Mol Wt** 379.45

**Exact Mass** 379.07

| # | Time | Area% |
|---|---|---|
| 1 | 1.466 | 100.00 |

BE177061$3

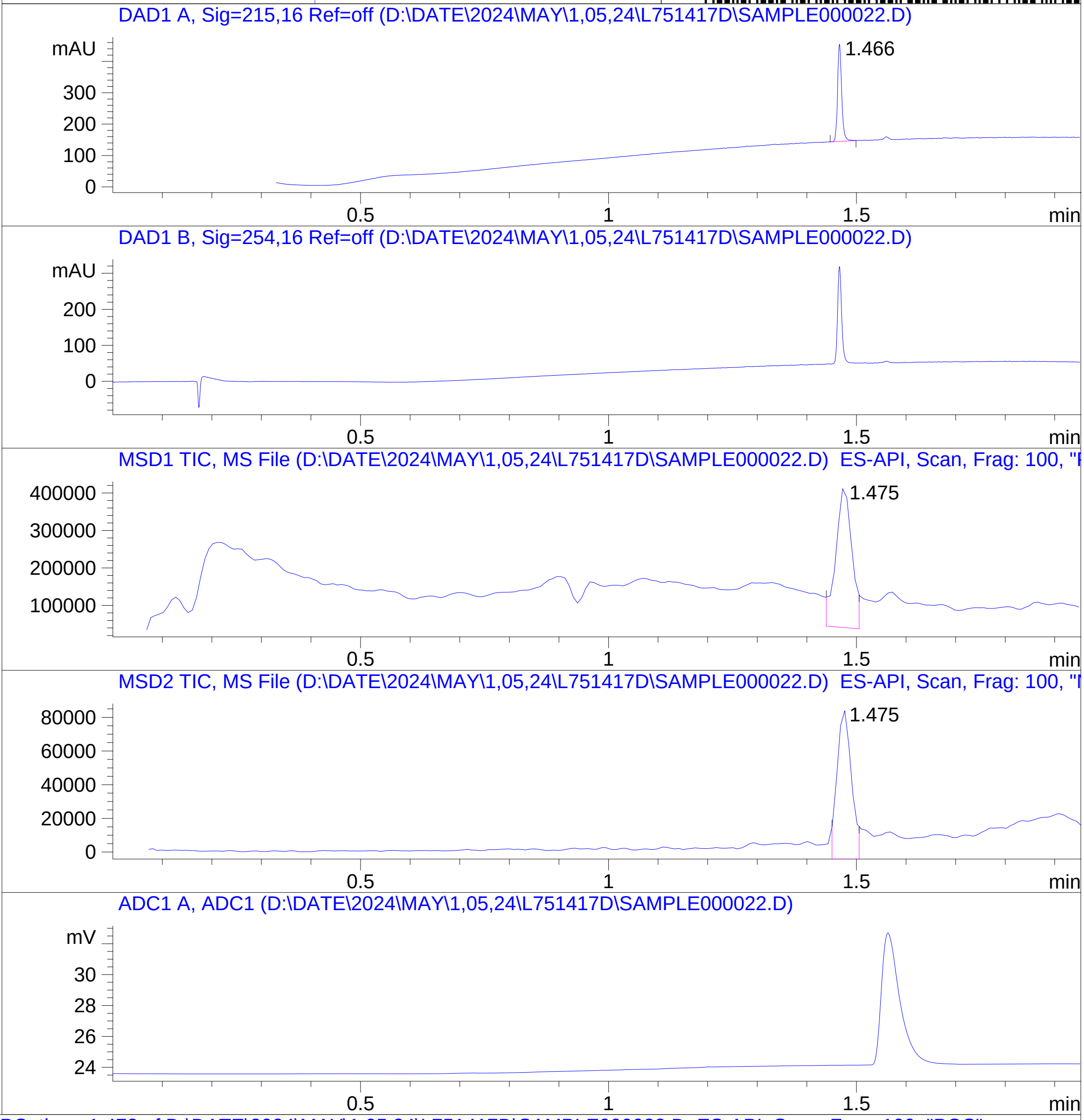


RT 1.475

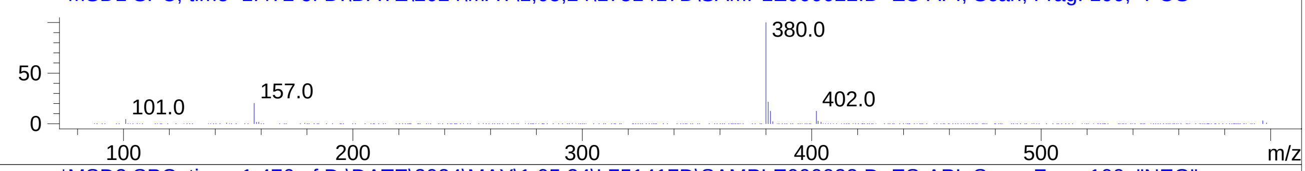


RT 1.475

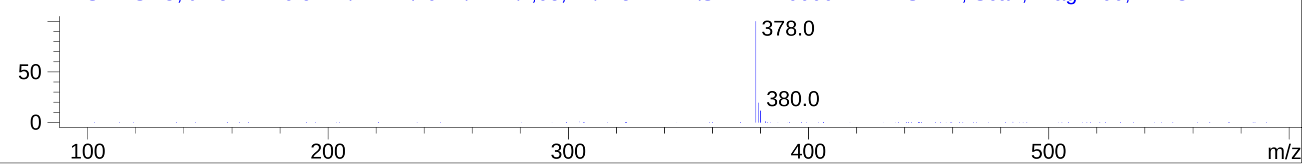

### Z4468759270.PDF

MaxPeak: 100.00%
Ret_Time: 1.436 min

# BE177013$2

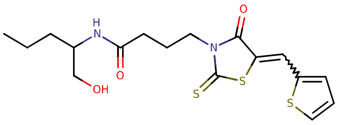

**Mol Wt** 398.56

**Exact Mass** 398.1

| # | Time | Area% |
|---|---|---|
| 1 | 1.436 | 100.00 |

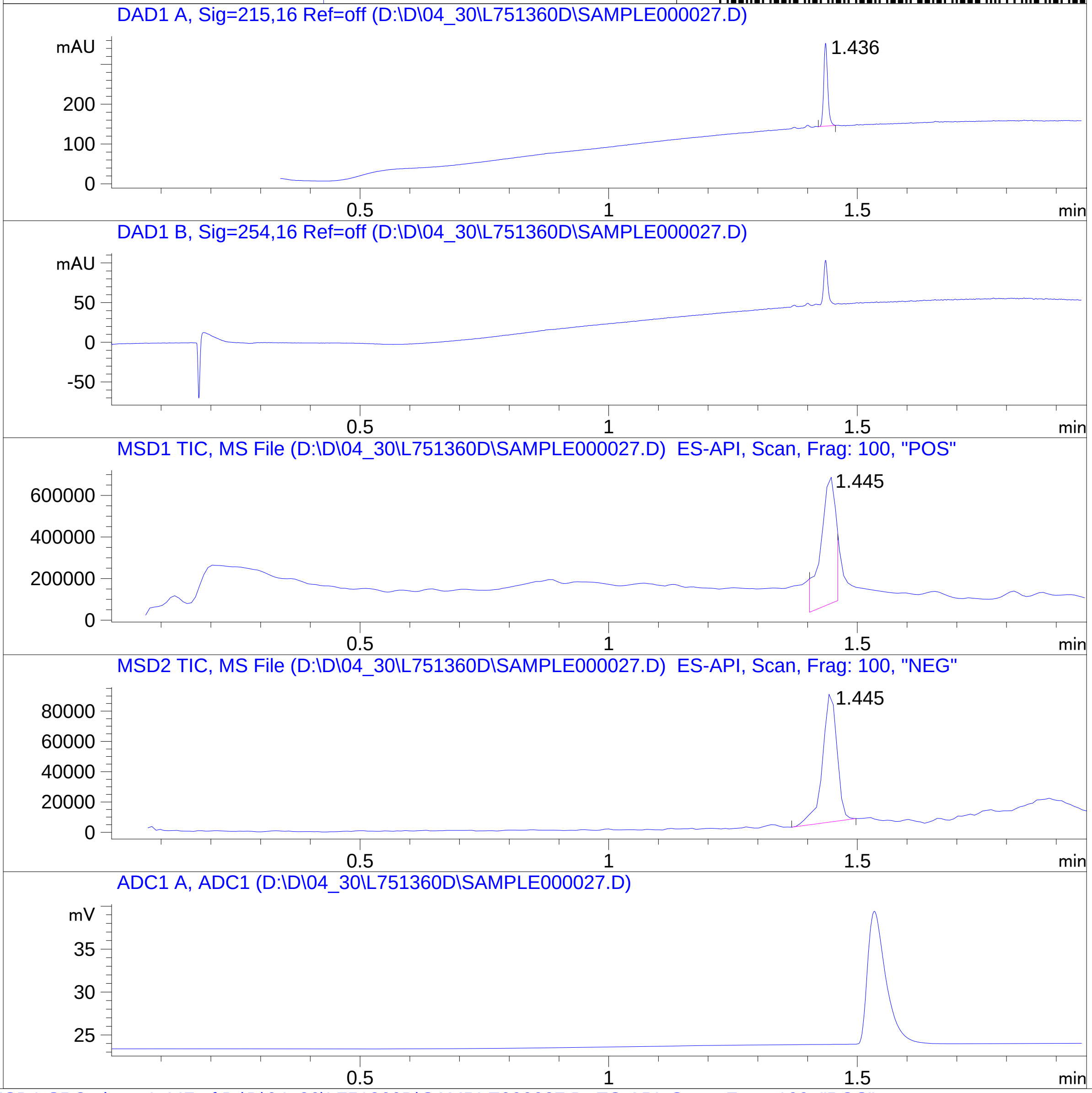


RT 1.445

RT 1.445

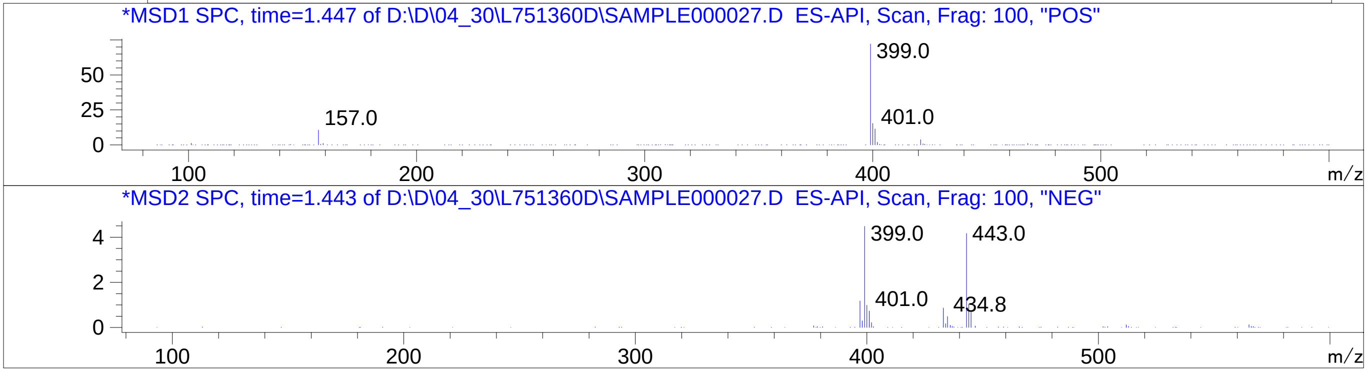

### Z5138684202.PDF

BE177062$2

MaxPeak: 100.00%
Ret_Time: 1.090 min

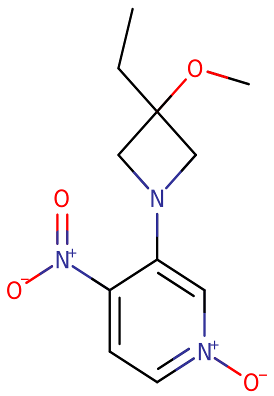

**Mol Wt** 253.26

**Exact Mass** 253.11

| # | Time | Area% |
|---|---|---|
| 1 | 1.090 | 100.00 |

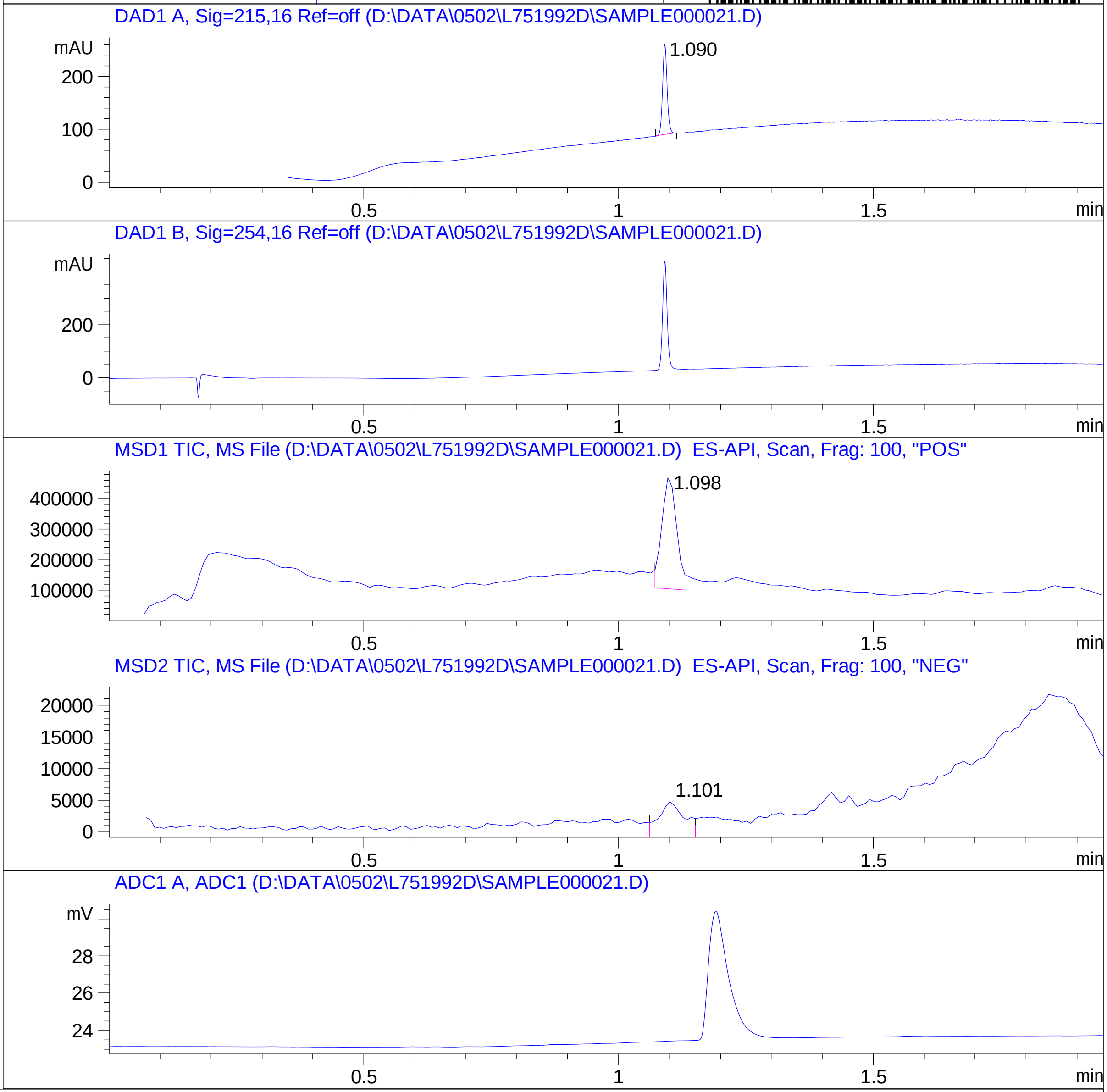


RT 1.098

RT 1.101

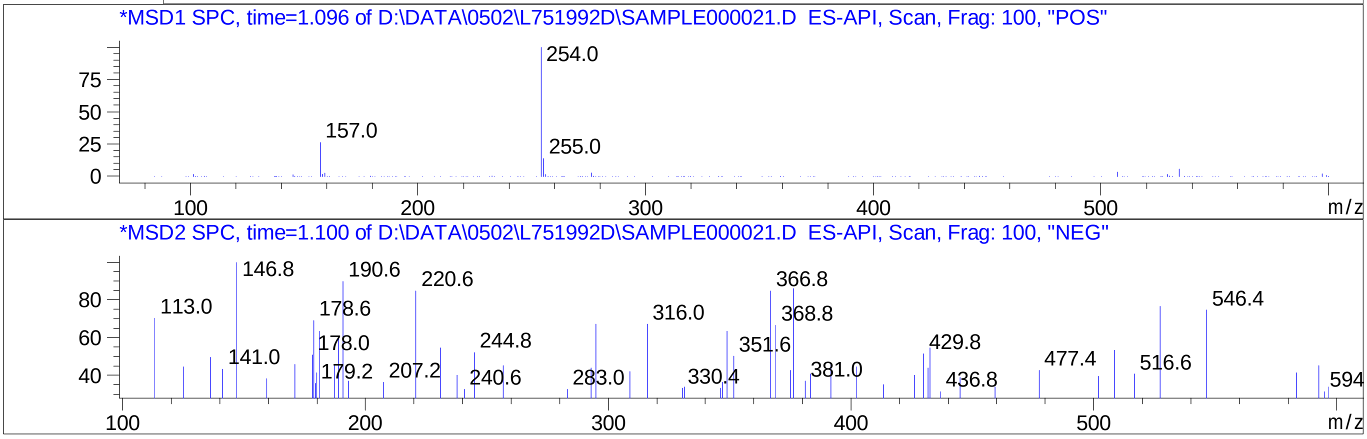


Inj.Date 5/2/2024 VB 

C:\CHEM32\1\METHODS\SUPOR_30.M

### Z5318592211.PDF

# BE177014$2

MaxPeak: 100.00%
Ret_Time: 0.470 min

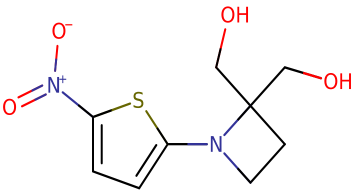


**Mol Wt** 244.27

**Exact Mass** 244.05

| # | Time | Area% |
|---|---|---|
| 1 | 0.470 | 100.00 |

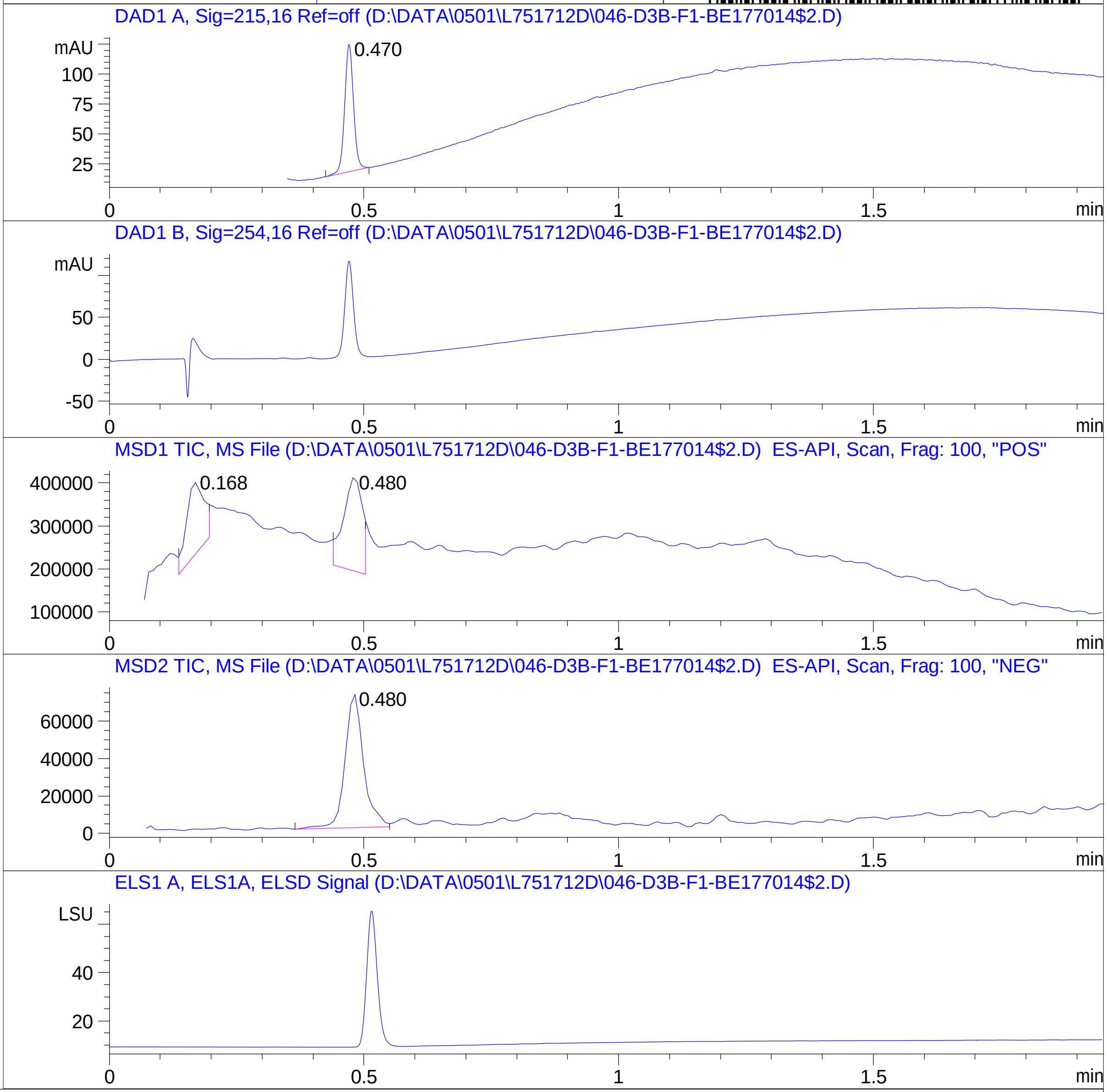


RT 0.168

RT 0.480

RT 0.480

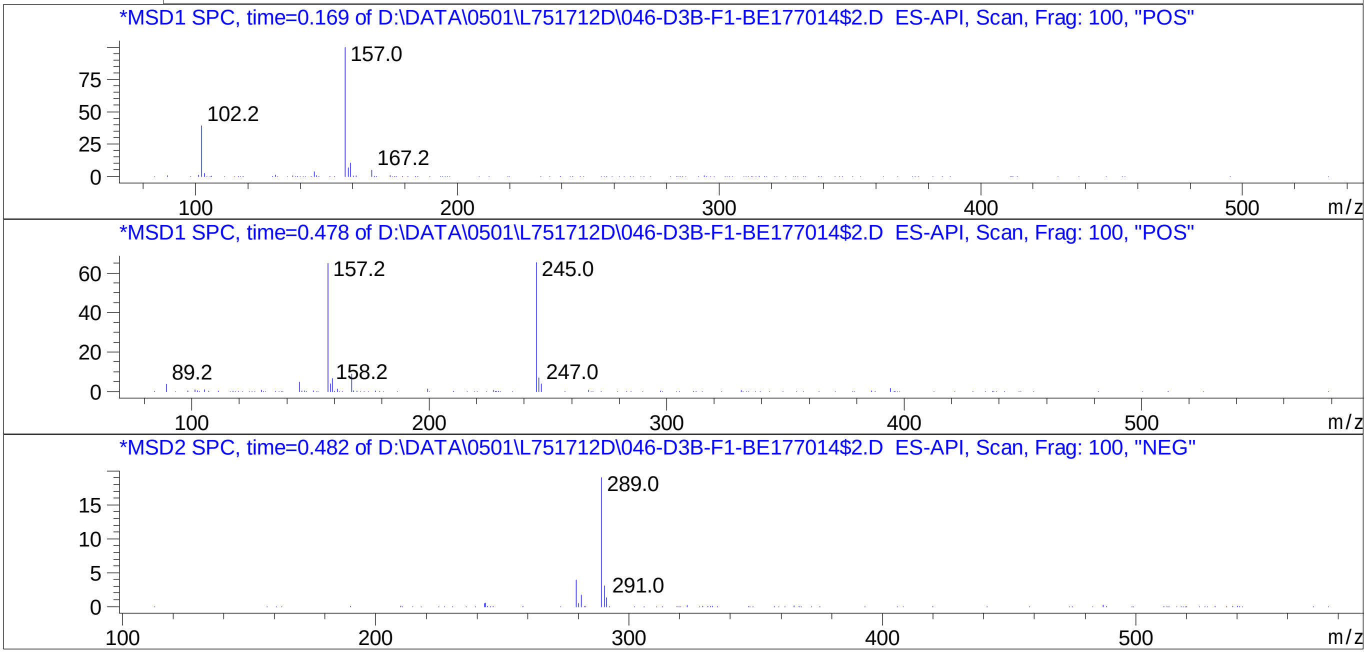


Inj.Date 5/1/2024 VB 13
C:\Chem32\1\Data\04_30\L751712D\SUPOR_30.M

### Z5318592387.PDF

MaxPeak: 100.00%
Ret_Time: 1.045 min

# BE177005$8

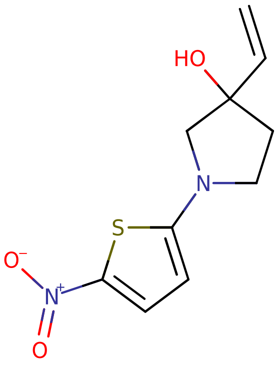


**Mol Wt** 240.28

**Exact Mass** 240.06

| # | Time | Area% |
|---|---|---|
| 1 | 1.045 | 100.00 |

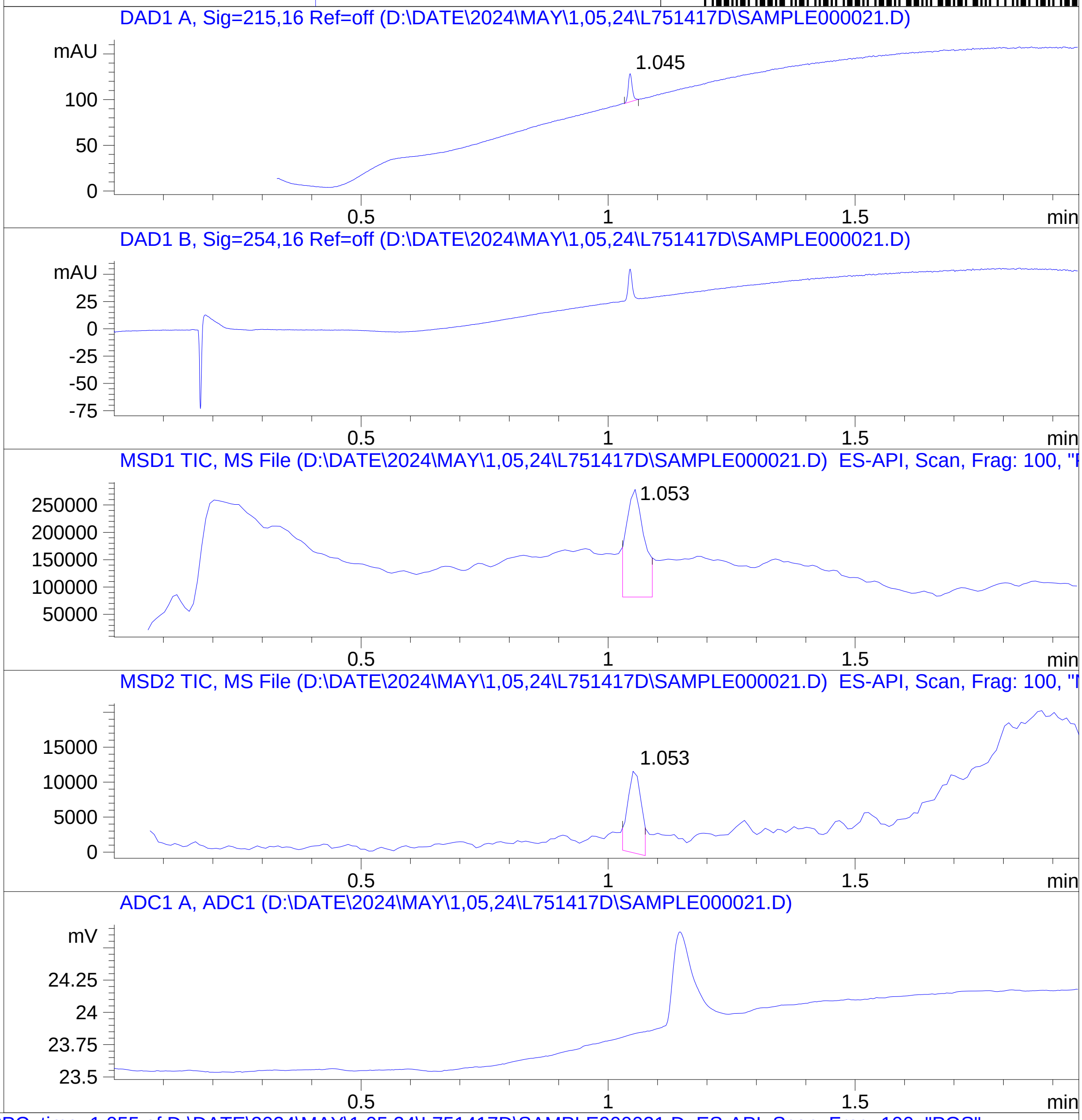


RT 1.053

RT 1.053

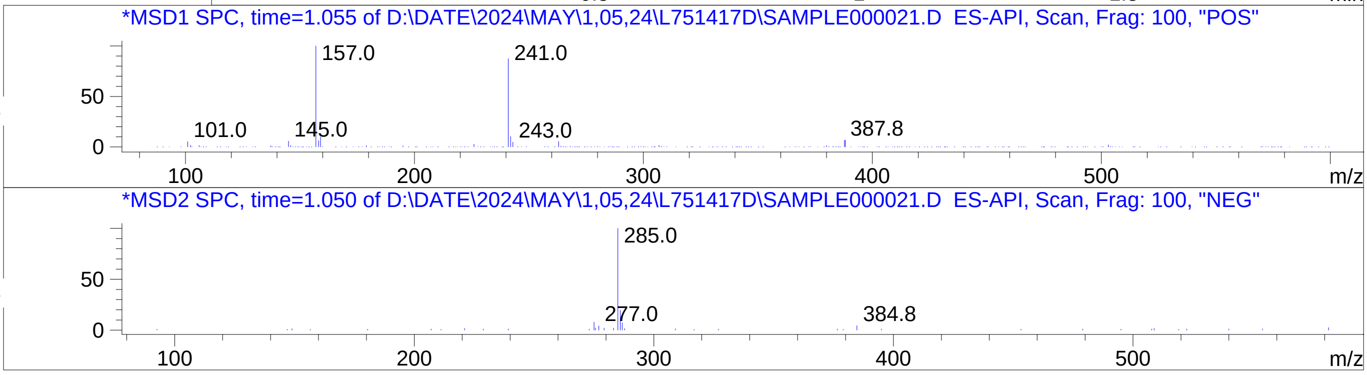

### Z5340501662.PDF

MaxPeak: 92.87%
Ret_Time: 1.292 min

BE177044$2

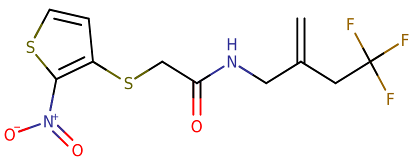


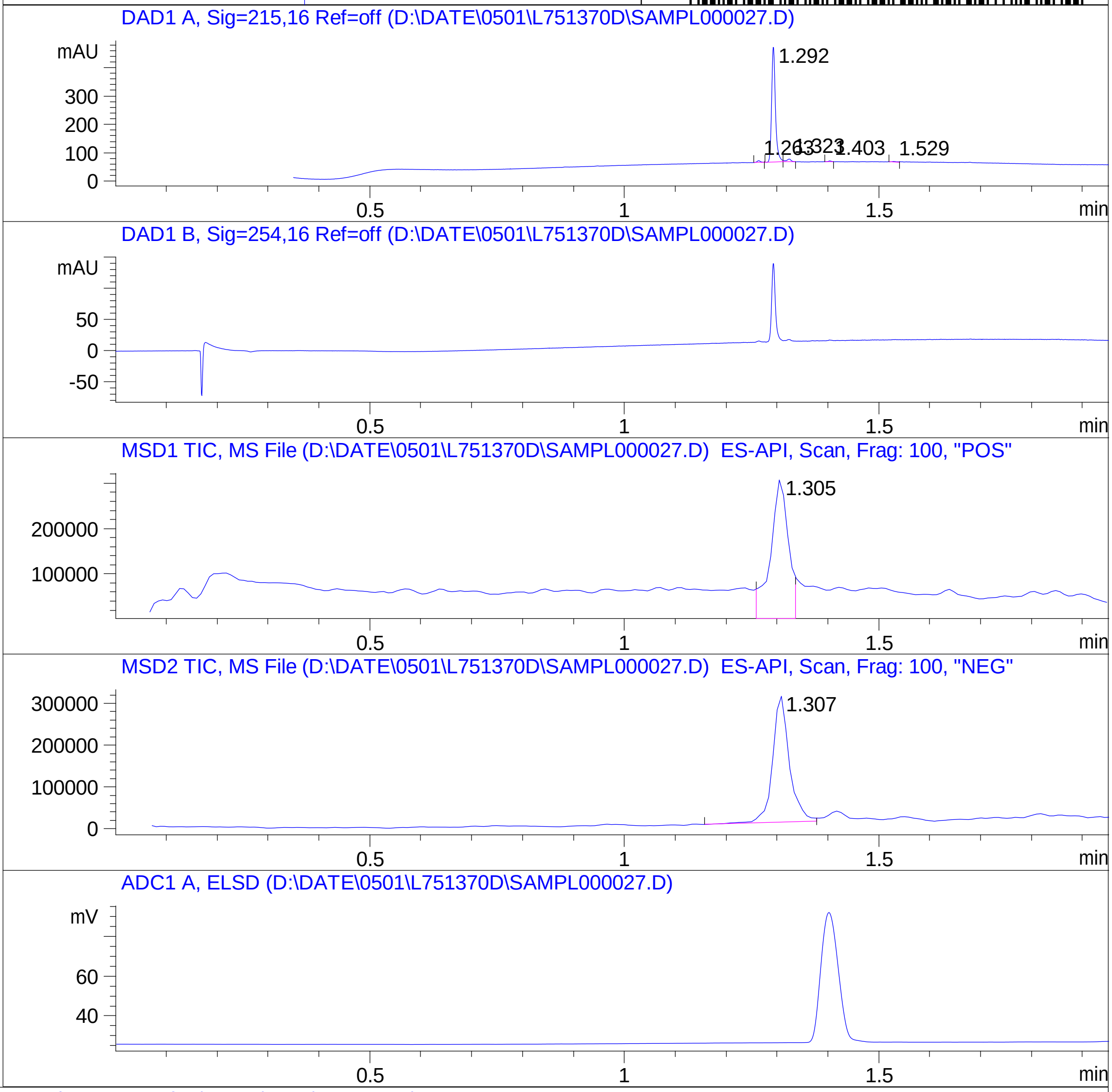


**Mol Wt** 340.34

**Exact Mass** 340.02

| # | Time | Area% |
|---|---|---|
| 1 | 1.263 | 1.48 |
| 2 | 1.292 | 92.87 |
| 3 | 1.323 | 3.37 |
| 4 | 1.403 | 1.03 |
| 5 | 1.529 | 1.24 |

RT 1.305

RT 1.307

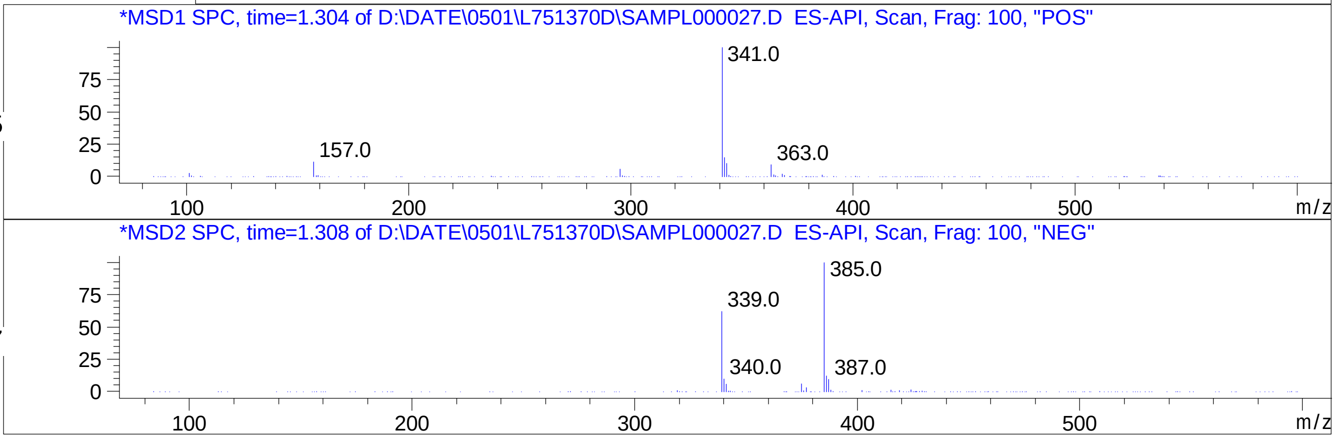


Inj.Date 4/30/2024 CH -6-

C:\CHEM32\1\METHODS\SUPOR_30.M

### Z5423194706.PDF

MaxPeak: 97.16%
Ret_Time: 0.940 min

# BE177086$2

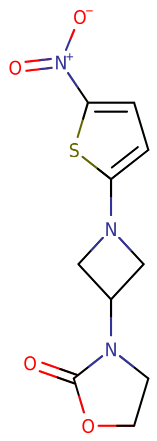


**Mol Wt** 269.28

**Exact Mass** 269.04

| # | Time | Area% |
|---|---|---|
| 1 | 0.940 | 97.16 |
| 2 | 0.983 | 1.08 |
| 3 | 1.235 | 1.76 |

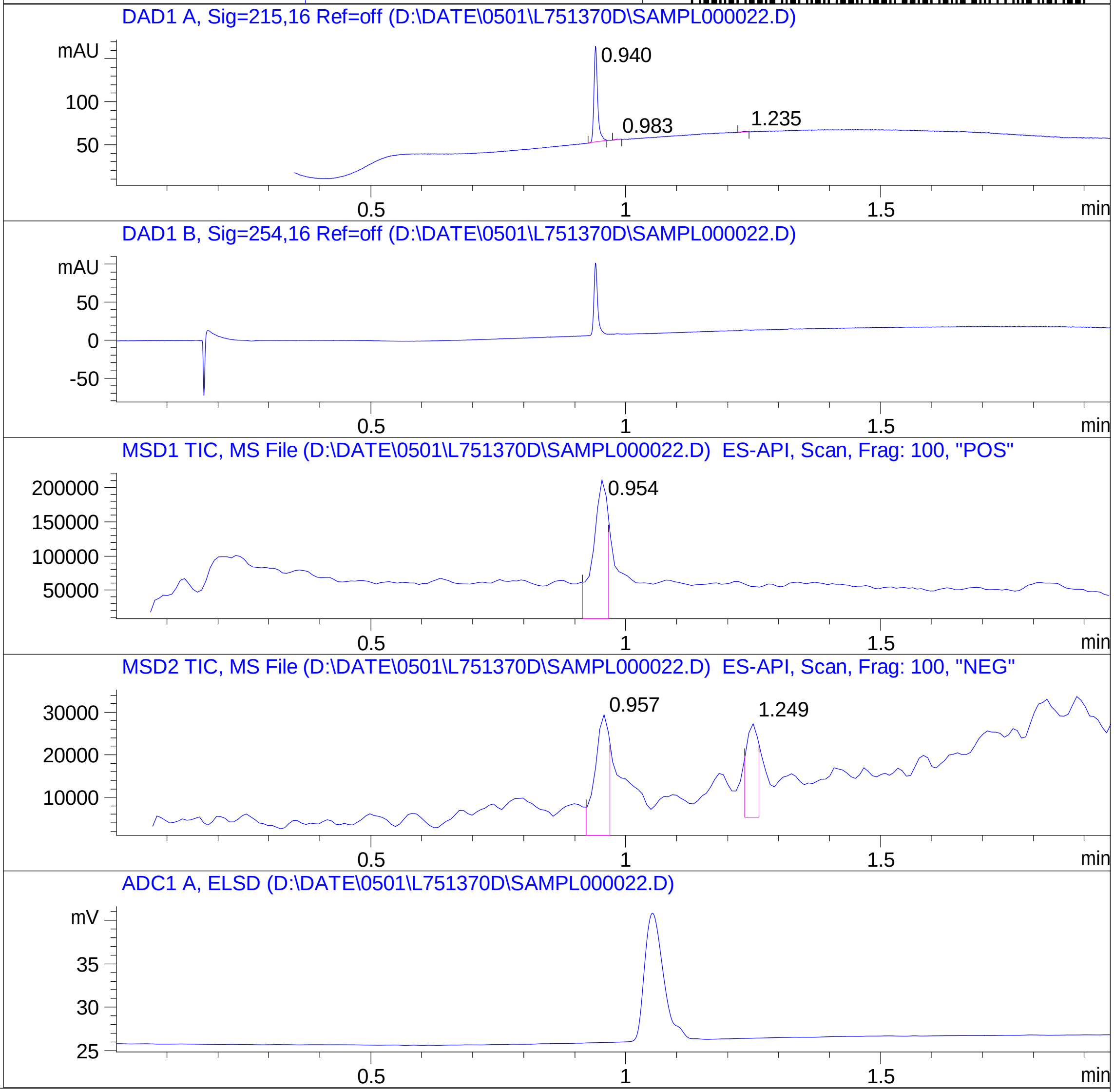


RT 0.954

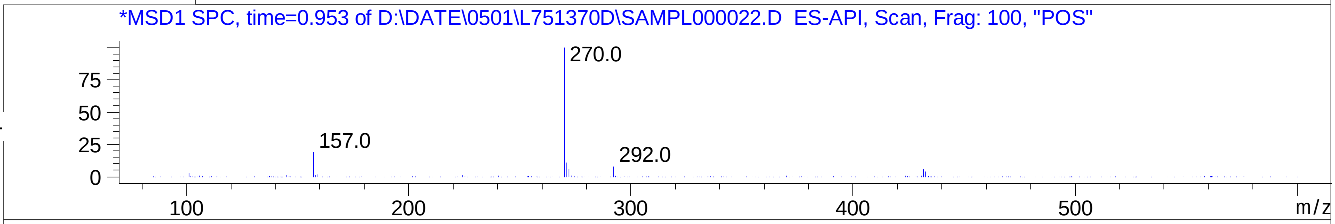


RT 0.957

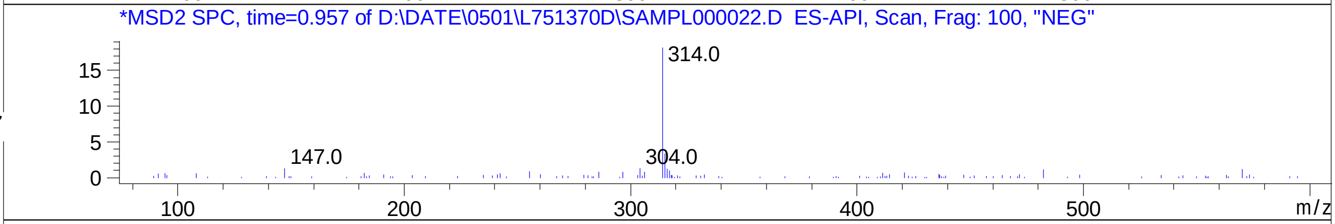


RT 1.249

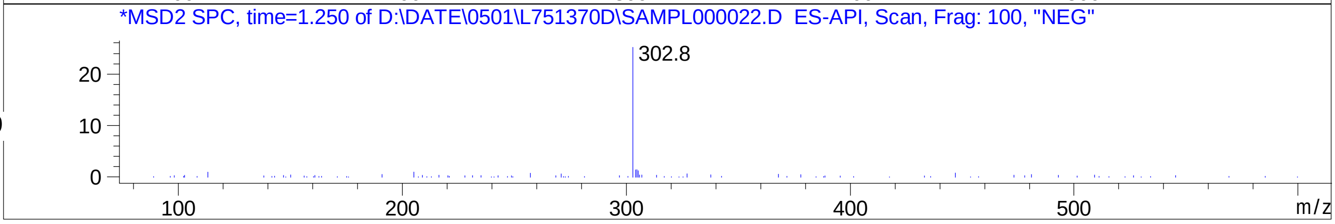

### Z5458838672.PDF

MaxPeak: 96.17%
Ret_Time: 1.178 min

# BE177072$3

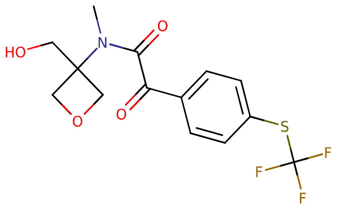


**Mol Wt** 349.32

**Exact Mass** 349.07

| # | Time | Area% |
|---|---|---|
| 1 | 1.178 | 96.17 |
| 2 | 1.268 | 3.83 |

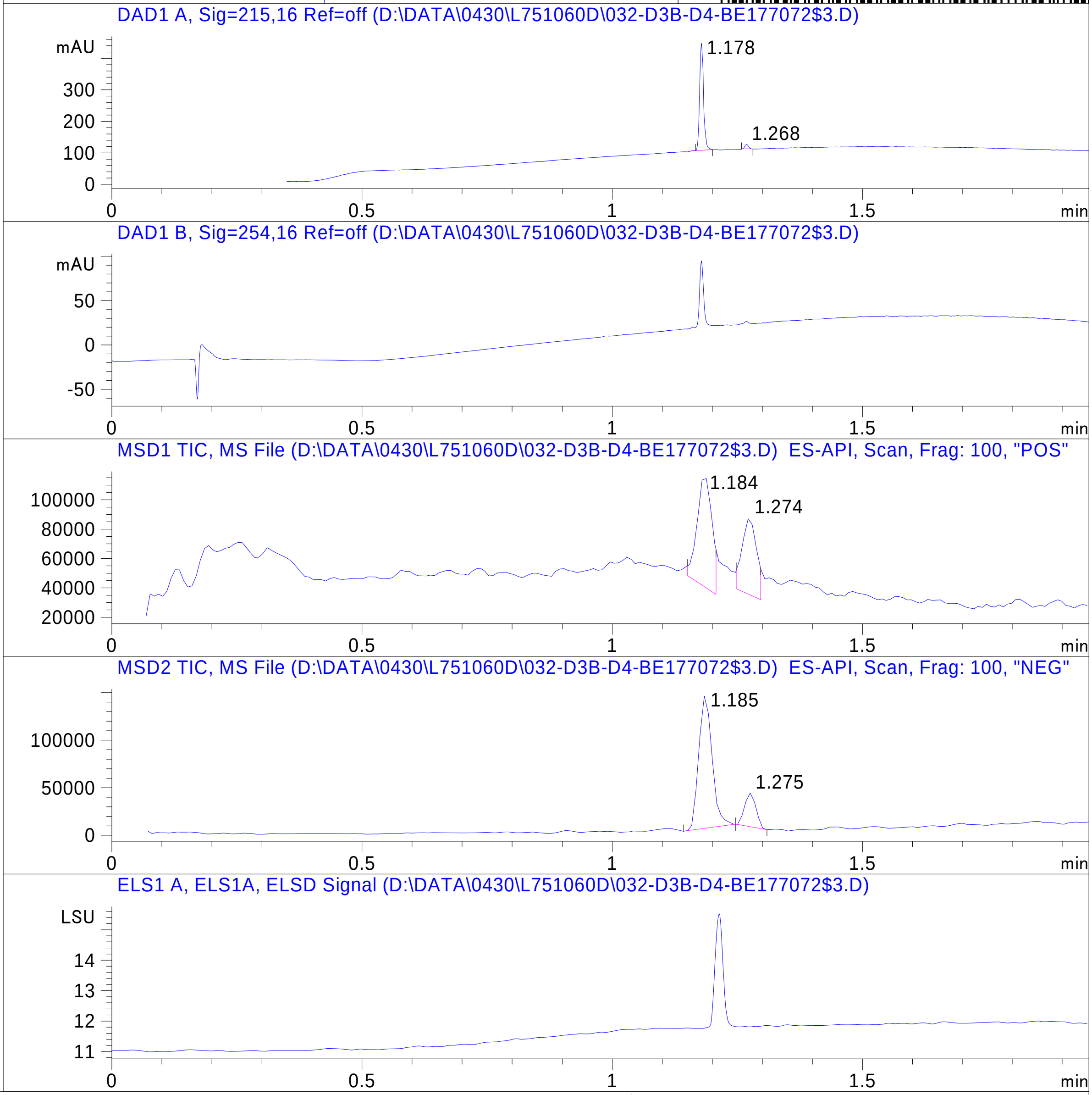


RT 1.184

RT 1.274

RT 1.185

RT 1.275

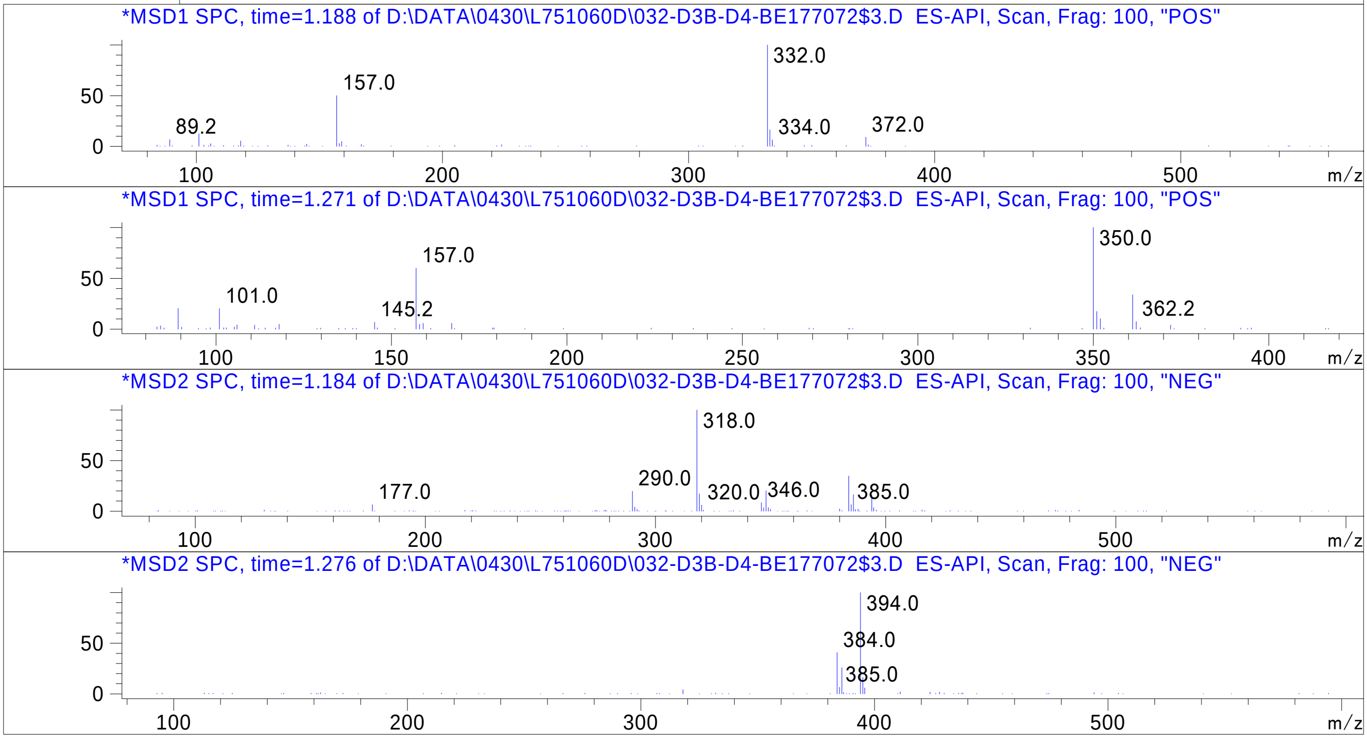


Inj.Date 4/29/2024 H -15-

C:\Chem32\1\Data\04_29\L751060D\SUPOR_30.M

### Z5530630474.PDF

# BD252365$2

MaxPeak: 100.00%
Ret_Time: 0.950 min

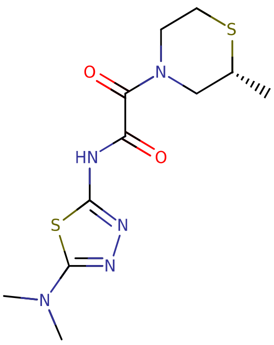

**Mol Wt** 315.42
**Exact Mass** 315.09

| # | Time | Area% |
|---|---|---|
| 1 | 0.950 | 100.00 |

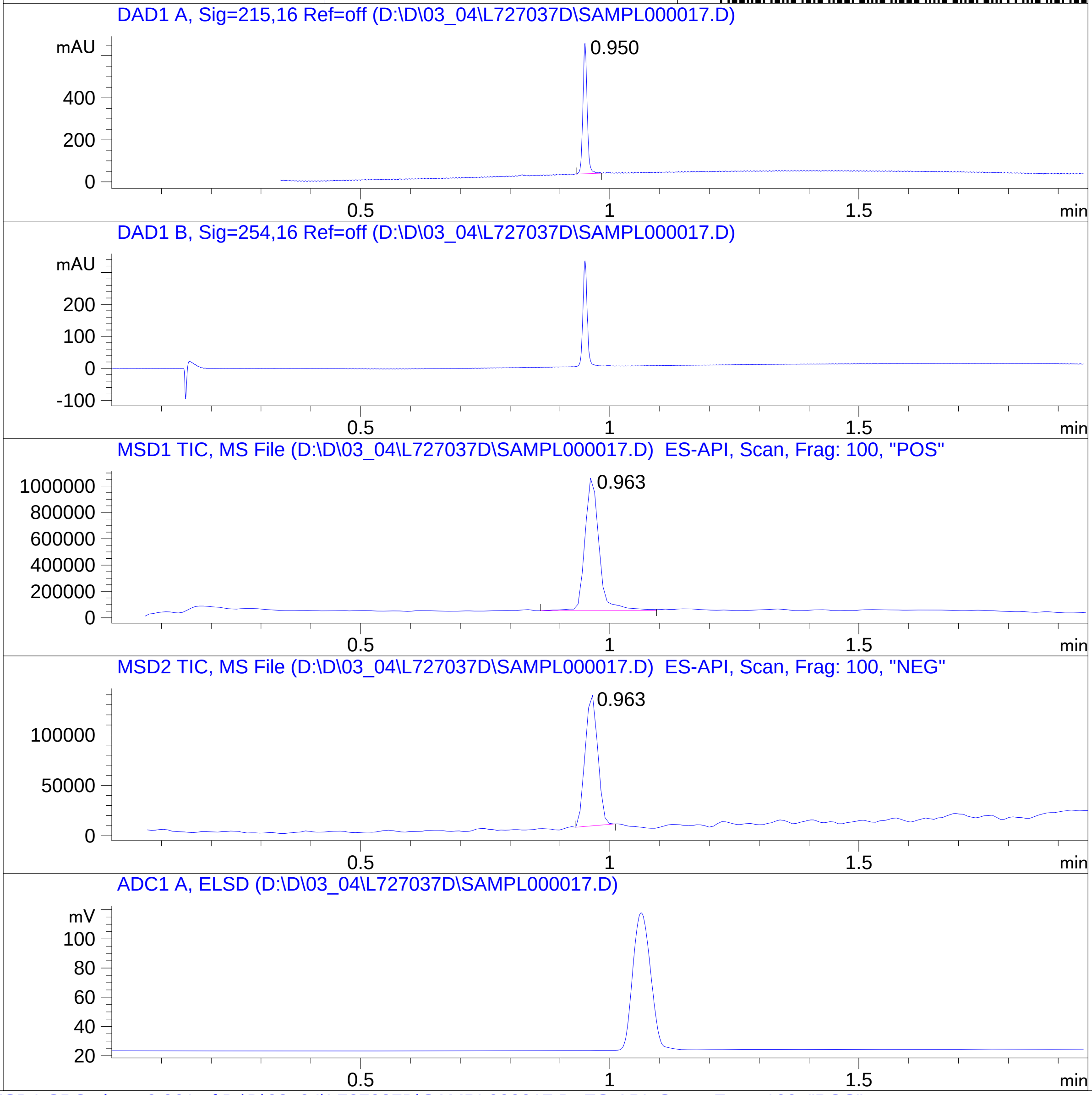


RT 0.963

RT 0.963

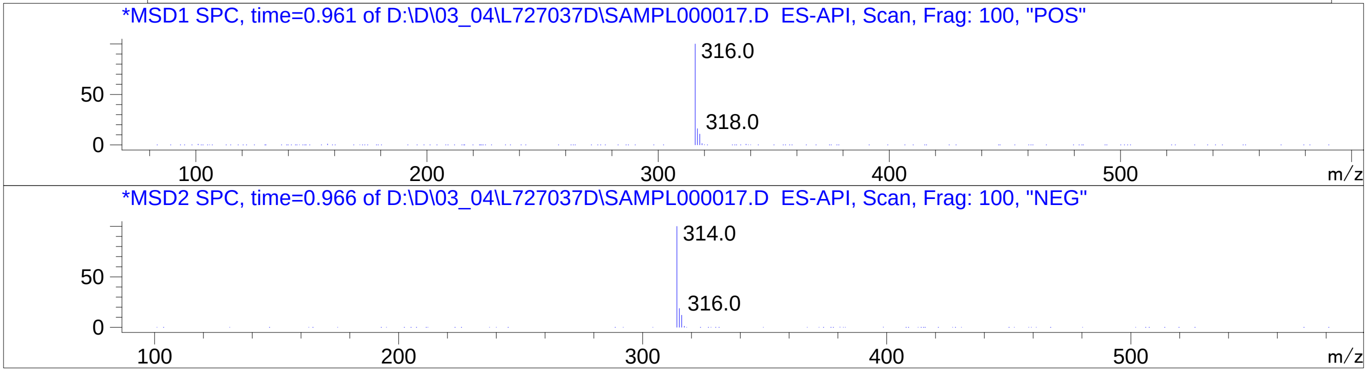


Inj.Date 3/4/2024
C:\CHEM32\1\METHODS\SUPOR_30.M

N

### Z5887206953.PDF

MaxPeak:  100.00%
Ret_Time: 0.896 min

# BE177015$1

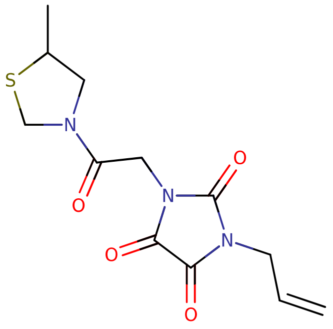

**Mol Wt** 297.33

**Exact Mass** 297.08

| # | Time | Area% |
|---|---|---|
| 1 | 0.896 | 100.00 |

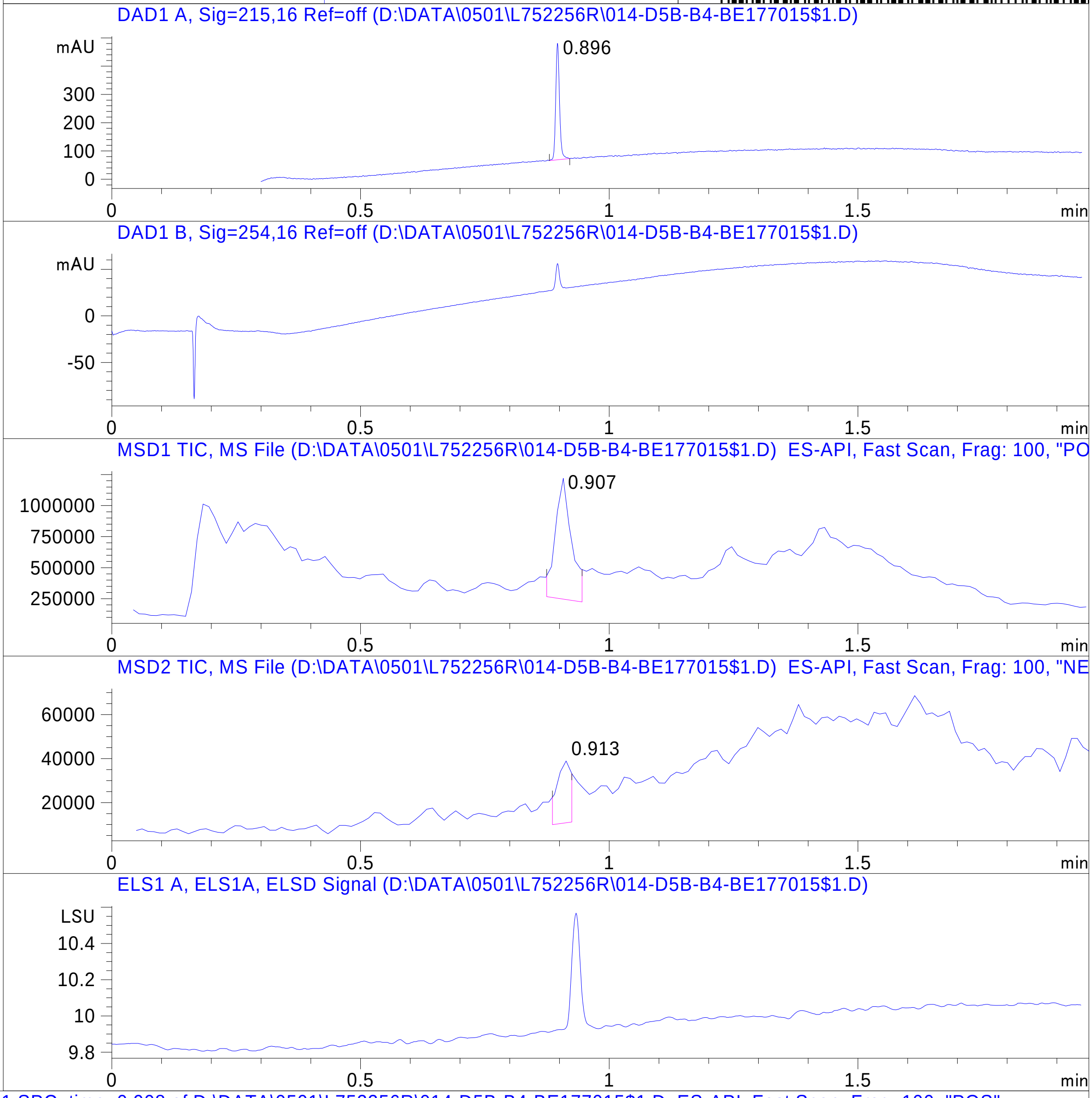


RT 0.907

RT 0.913

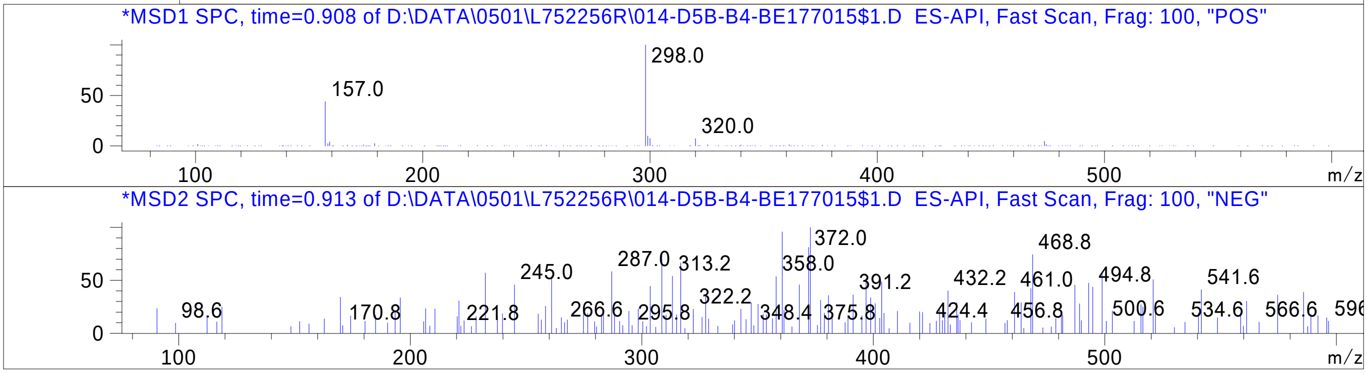

### Z5887209772.PDF

# BD252419$1

MaxPeak:  97.04%
Ret_Time: 0.970 min

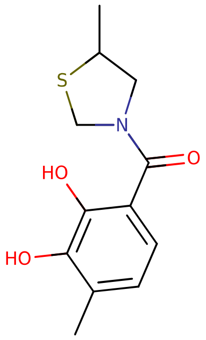


**Mol Wt** 253.32

**Exact Mass** 253.09

| # | Time | Area% |
|---|---|---|
| 1 | 0.710 | 1.37 |
| 2 | 0.970 | 97.04 |
| 3 | 1.224 | 1.59 |

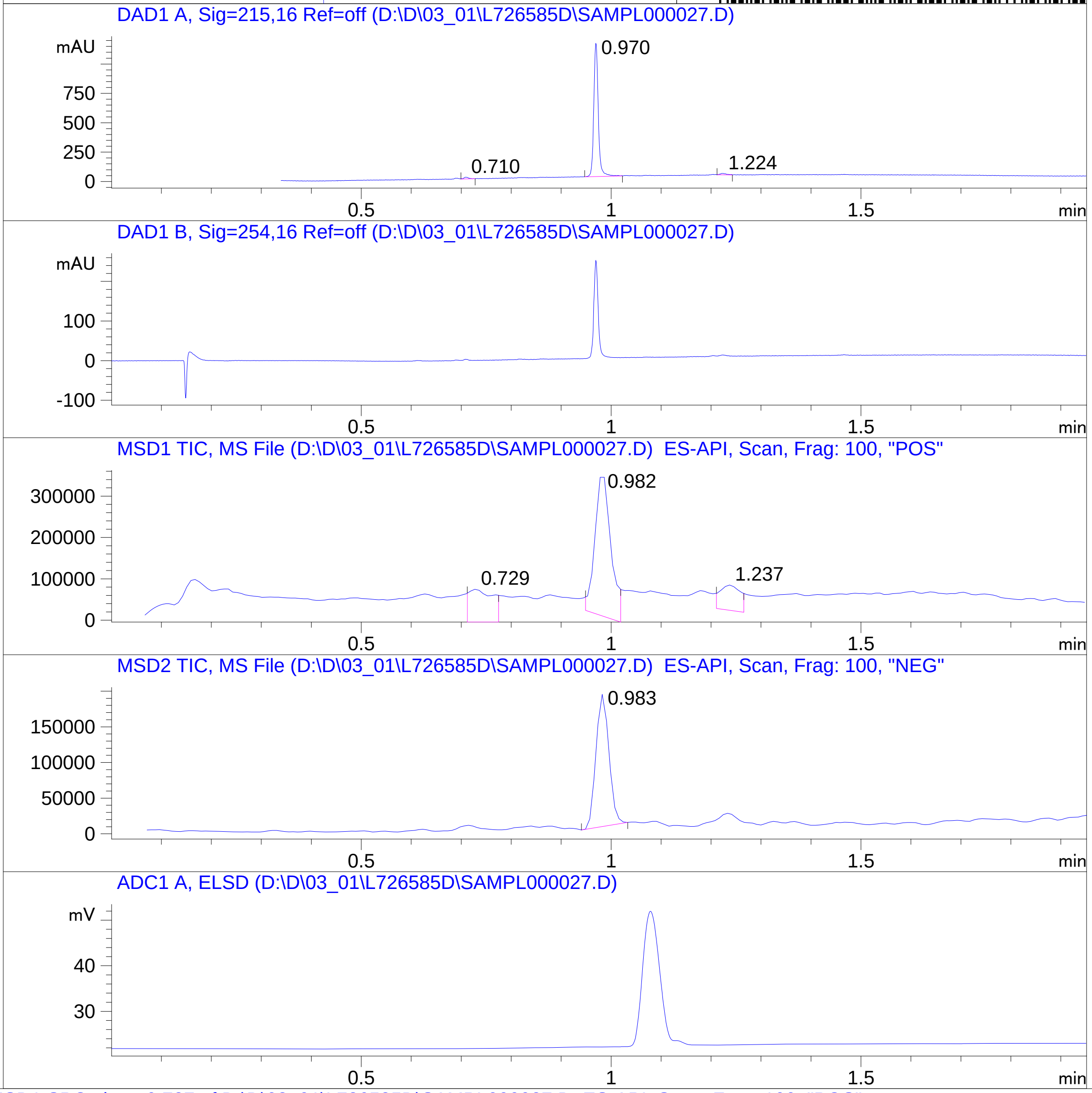


RT 0.729

RT 0.982

RT 1.237

RT 0.983

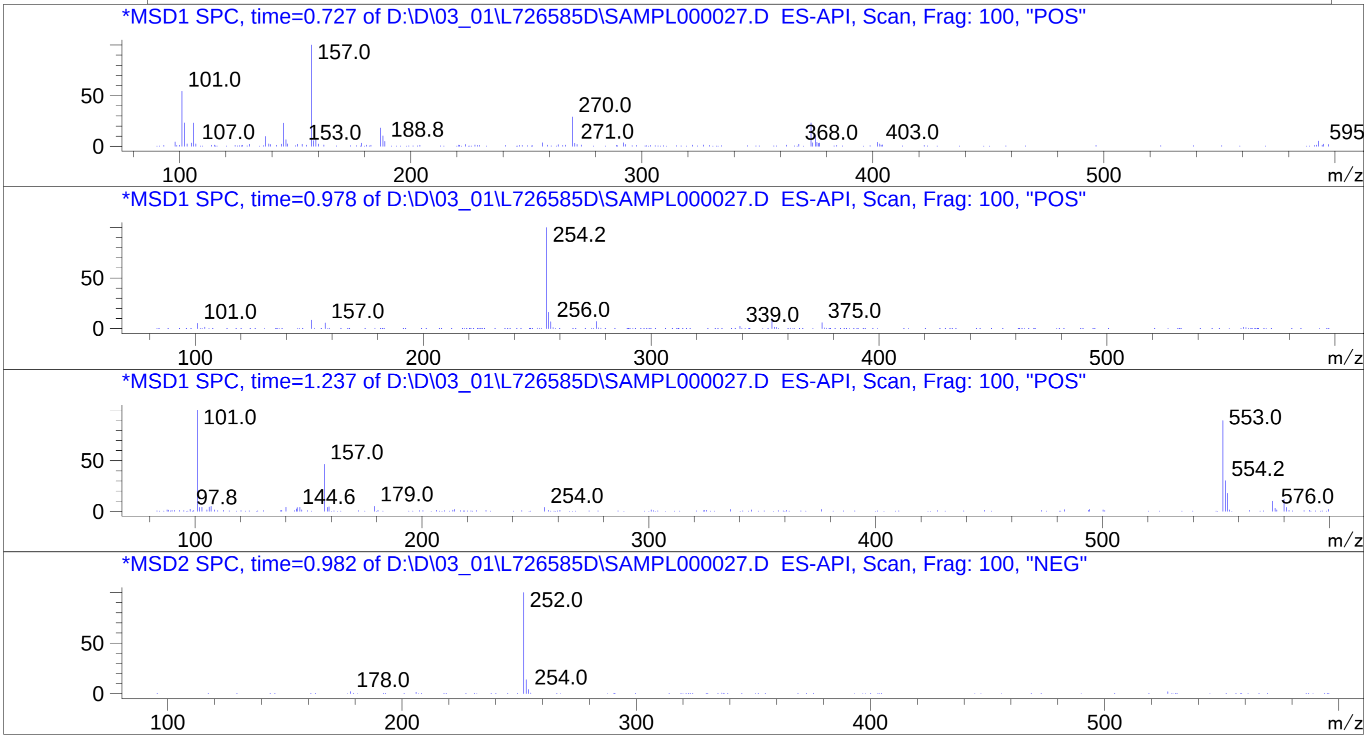

### Z5887220253.PDF

MaxPeak: 100.00%
Ret_Time: 0.966 min

# BE177038$1

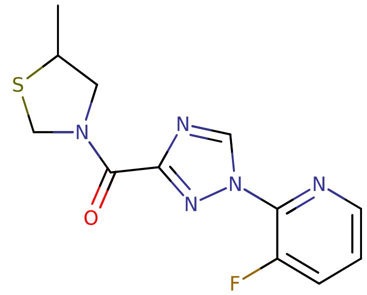

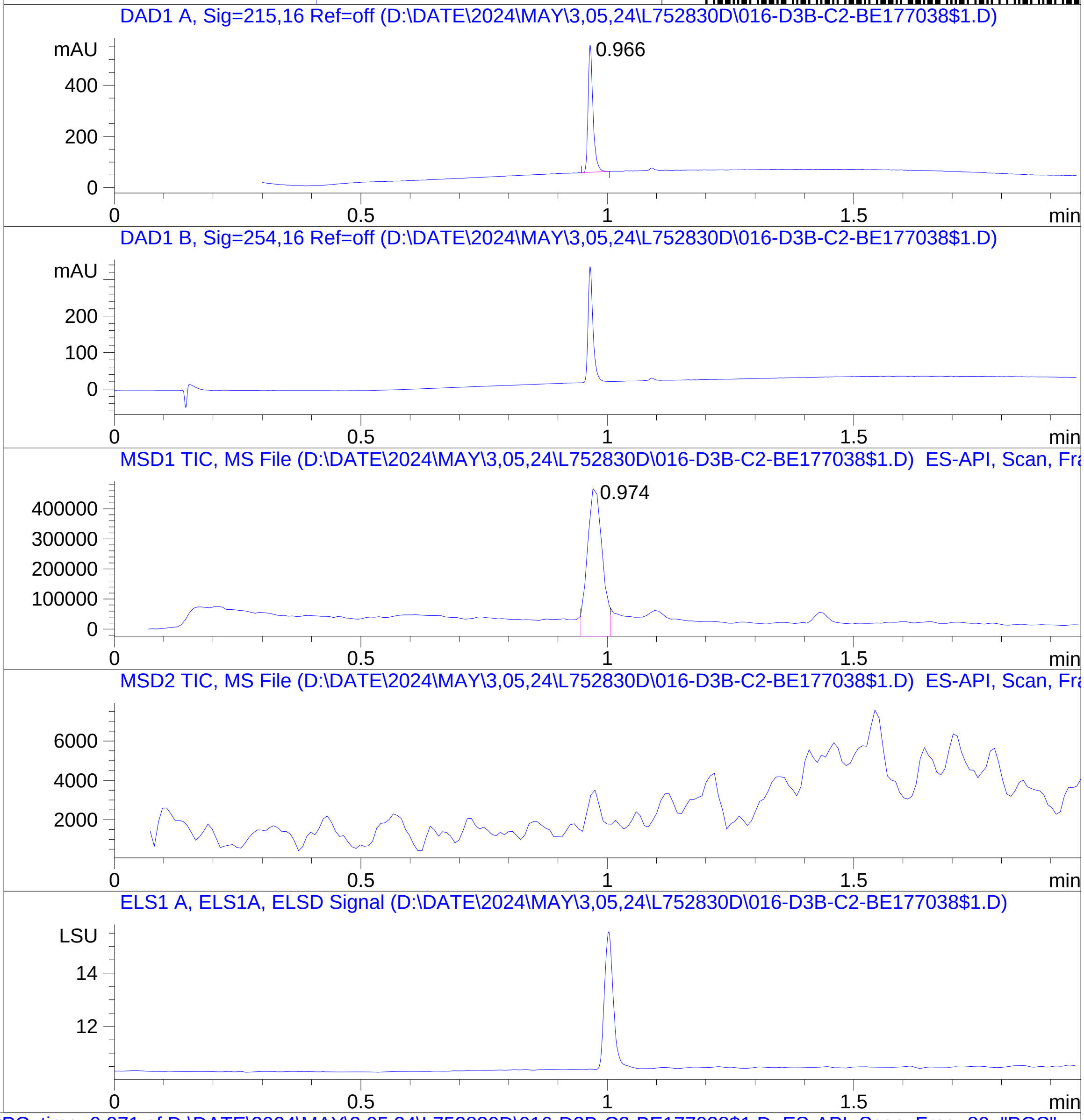


**Mol Wt** 293.32

**Exact Mass** 293.08

| # | Time | Area% |
|---|---|---|
| 1 | 0.966 | 100.00 |

RT 0.974

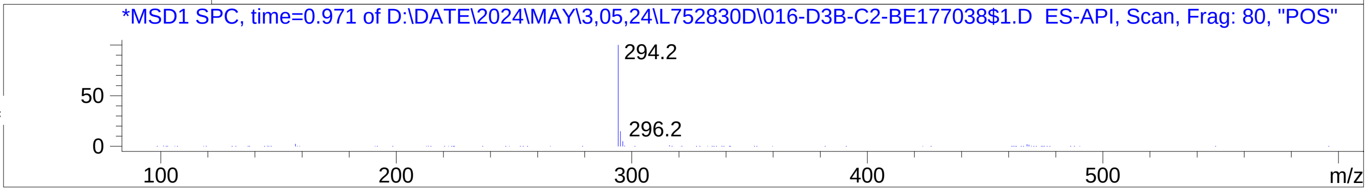


Inj.Date 5/2/2024 T -16- ->
C:\Chem32\1\Data\05_02\L752830D\SUPOR_30.M

### Z5887221669.PDF

MaxPeak:  100.00%
Ret_Time: 1.082 min

# BE177027$6

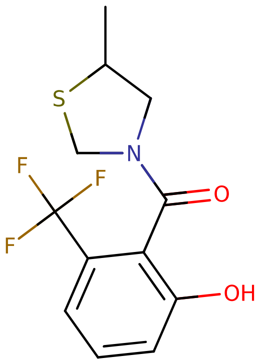


**Mol Wt** 291.29

**Exact Mass** 291.07

| # | Time | Area% |
|---|---|---|
| 1 | 1.082 | 100.00 |

RT 1.090

RT 1.091

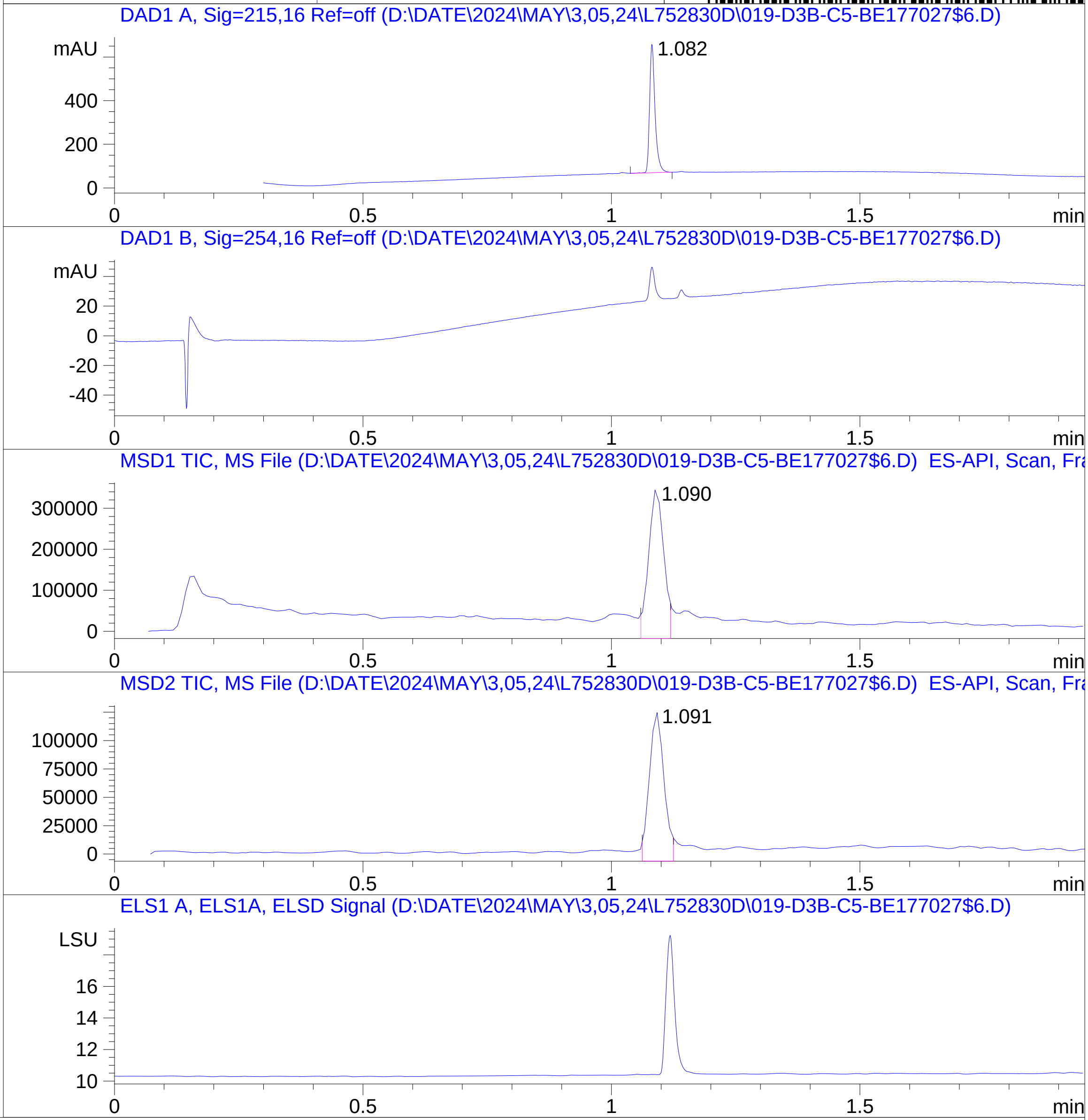


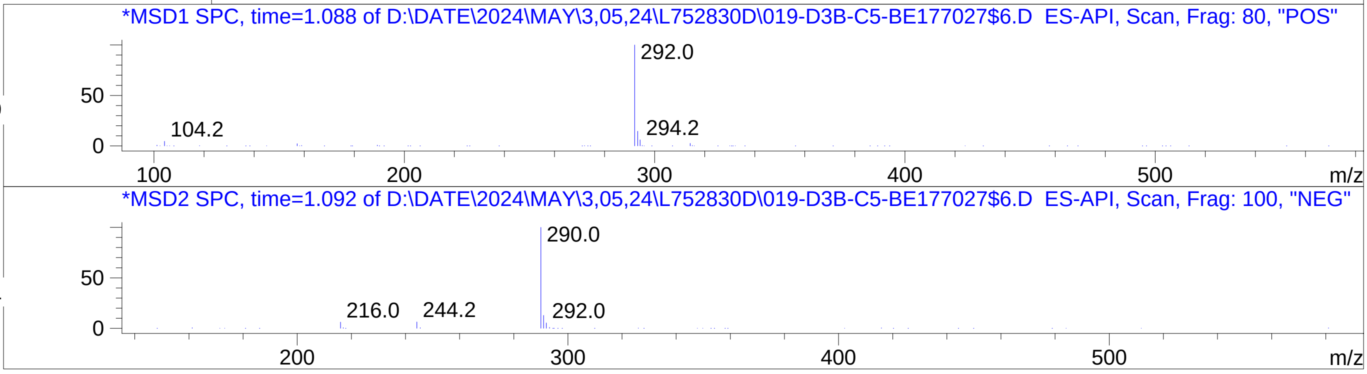

### Z5891107923.PDF

MaxPeak: 97.91%
Ret_Time: 0.816 min

# BE177087$2

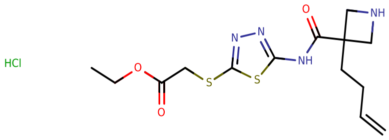


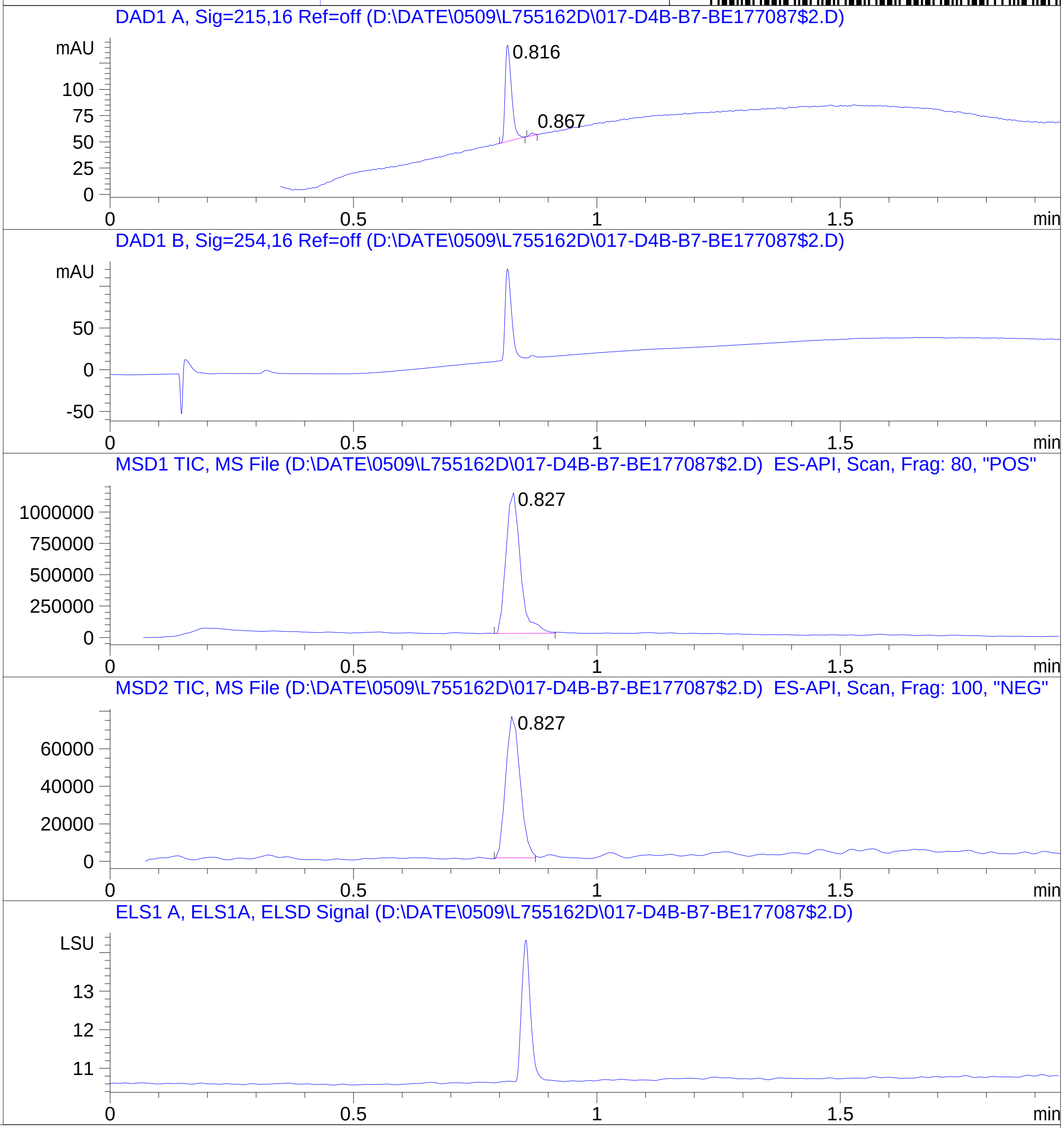


**Mol Wt** 392.92

**Exact Mass** 356.11

| # | Time | Area% |
|---|---|---|
| 1 | 0.816 | 97.91 |
| 2 | 0.867 | 2.09 |

RT 0.827

RT 0.827

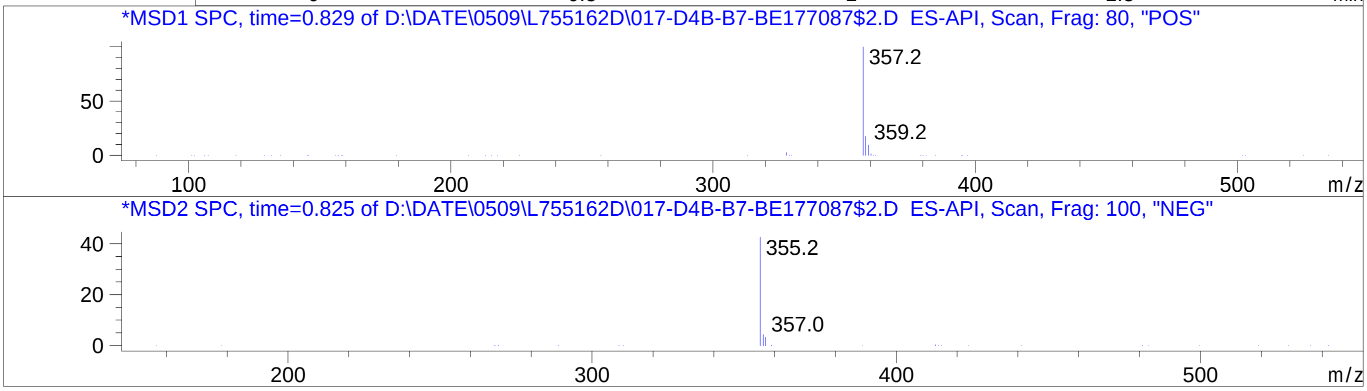

### Z6122202104.PDF

MaxPeak:  100.00%
Ret_Time: 0.767 min

# BE177073$1

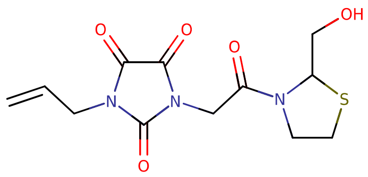


**Mol Wt** 313.33

**Exact Mass** 313.07

| # | Time | Area% |
| --- | --- | --- |
| 1 | 0.767 | 100.00 |

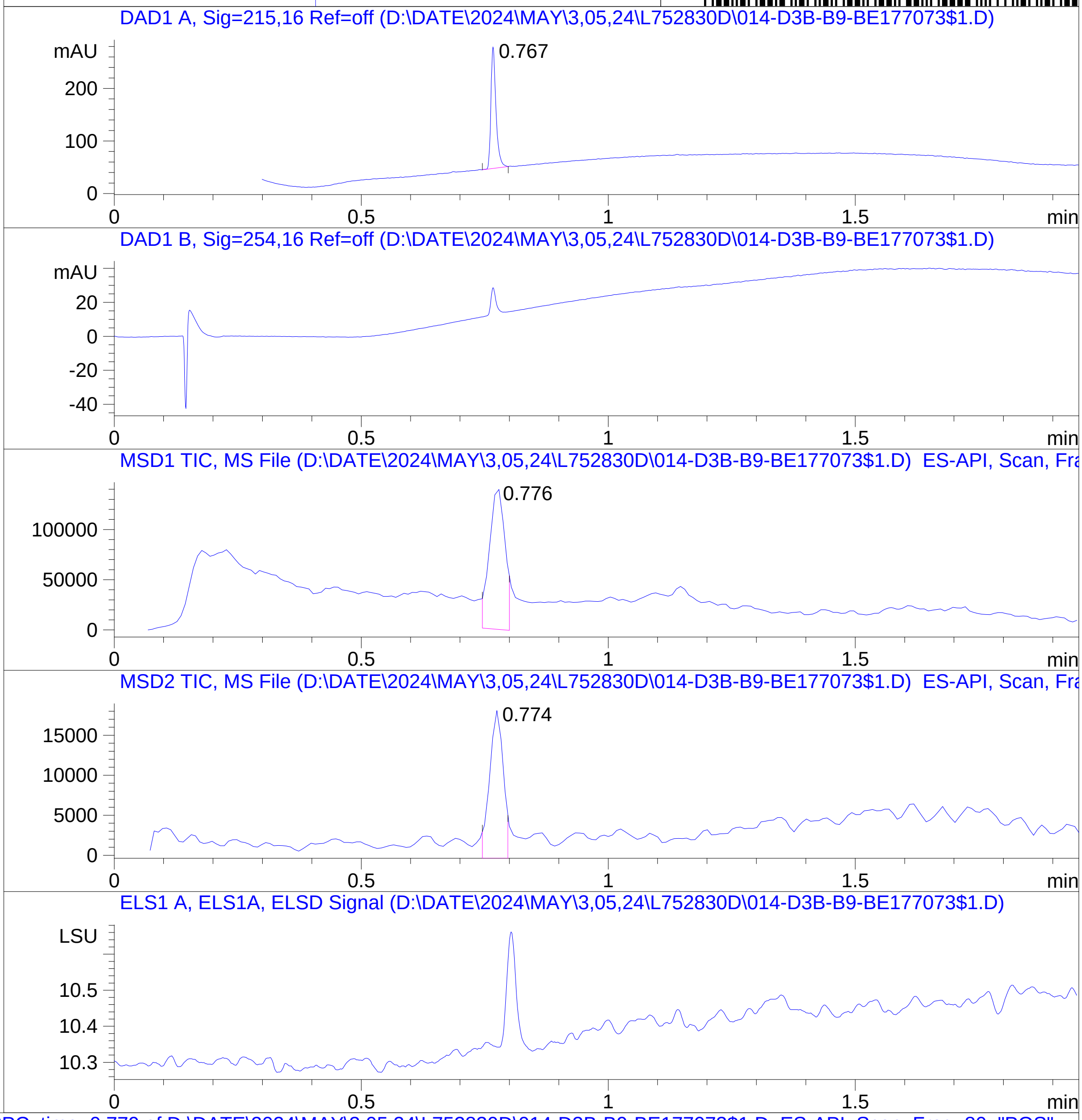


RT 0.776

RT 0.774

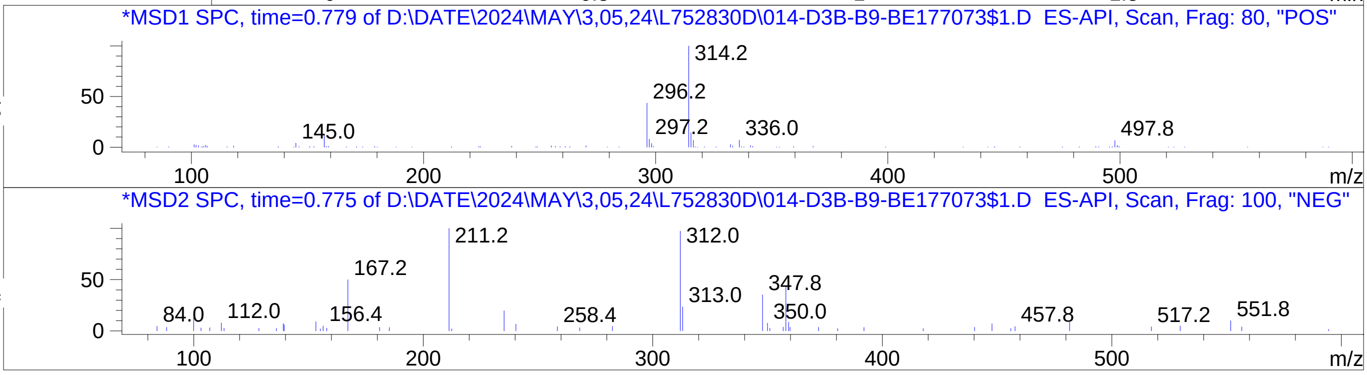

### Z6996322771.PDF

BE177006$3

MaxPeak: 51.67%
Ret_Time: 0.855 min

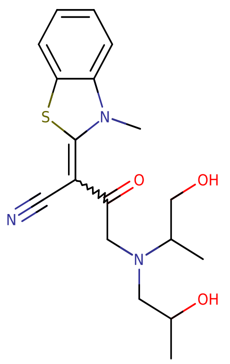


**Mol Wt** 361.46

**Exact Mass** 361.17

| # | Time | Area% |
|---|---|---|
| 1 | 0.855 | 51.67 |
| 2 | 0.868 | 42.01 |
| 3 | 0.953 | 2.08 |
| 4 | 1.188 | 2.48 |
| 5 | 1.242 | 1.75 |

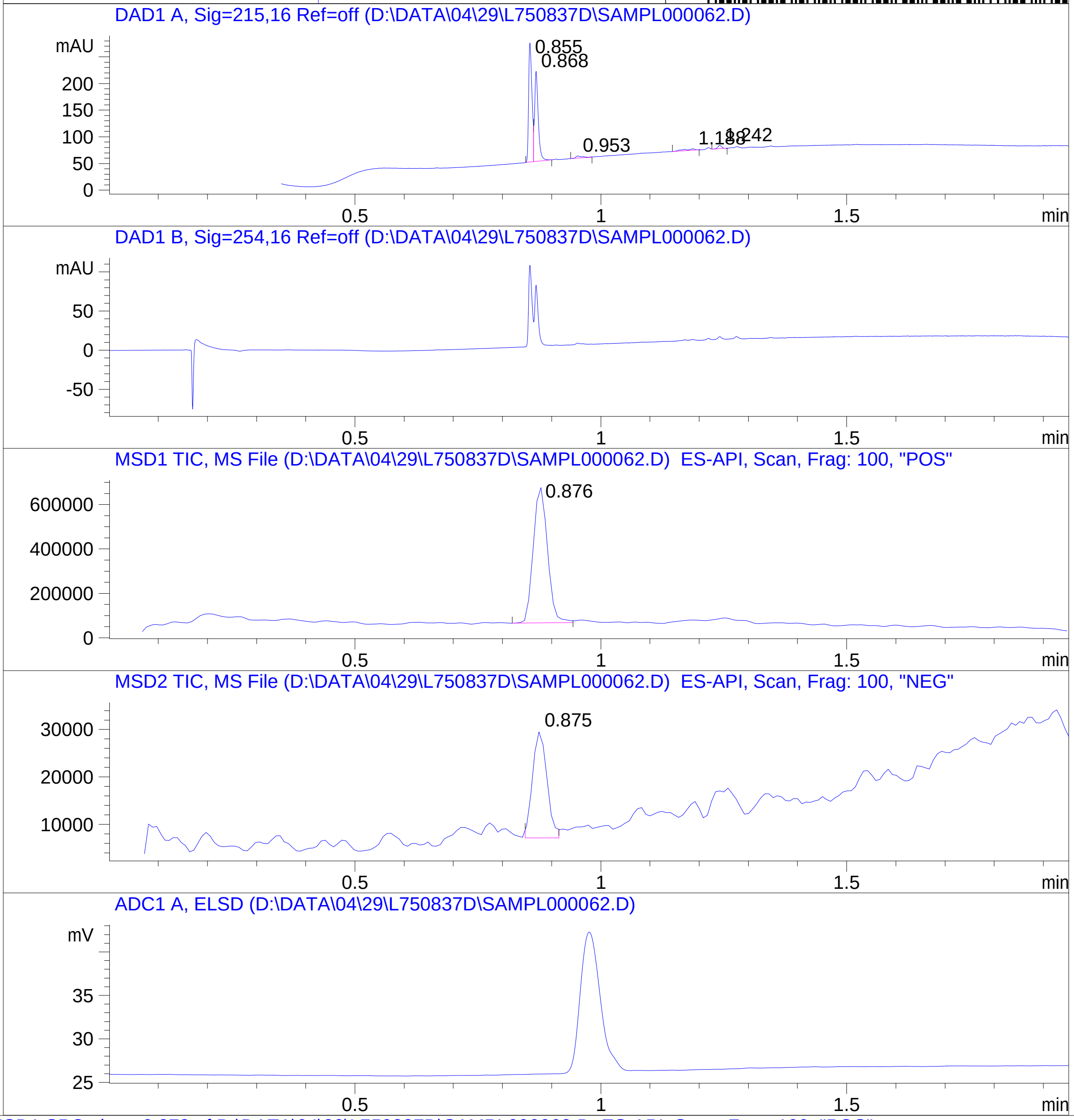


RT 0.876

RT 0.875

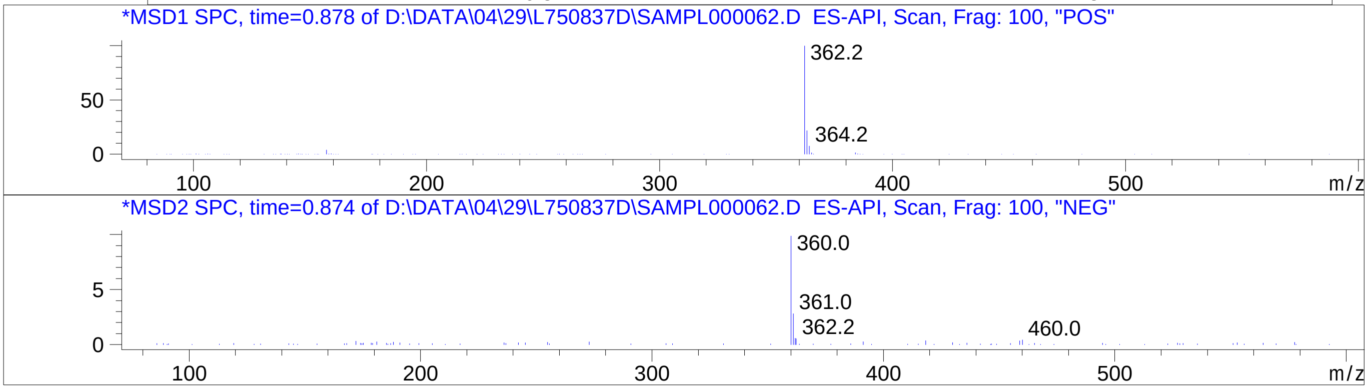


Inj.Date 4/29/2024 CH -6-
C:\CHEM32\1\METHODS\SUPOR_30.M

### Z7445715318.PDF

# BE177045$3

MaxPeak:  48.42%
Ret_Time: 0.879 min

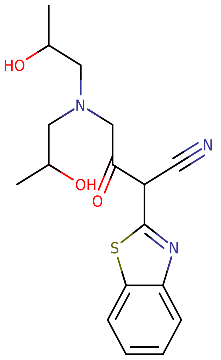


**Mol Wt** 347.43

**Exact Mass** 347.15

| # | Time | Area% |
|---|---|---|
| 1 | 0.854 | 2.37 |
| 2 | 0.873 | 45.43 |
| 3 | 0.879 | 48.42 |
| 4 | 1.116 | 3.78 |

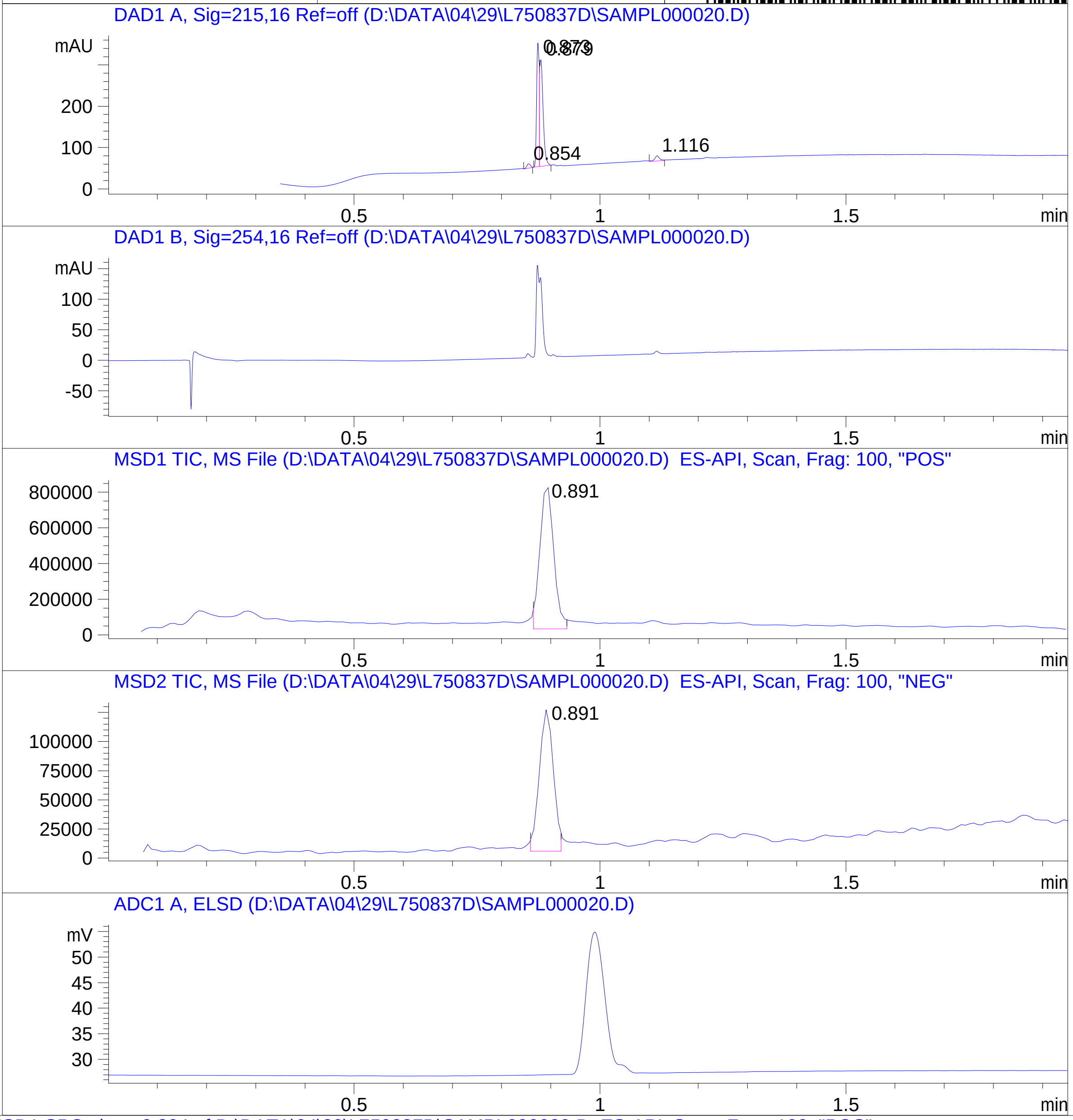


RT 0.891

RT 0.891

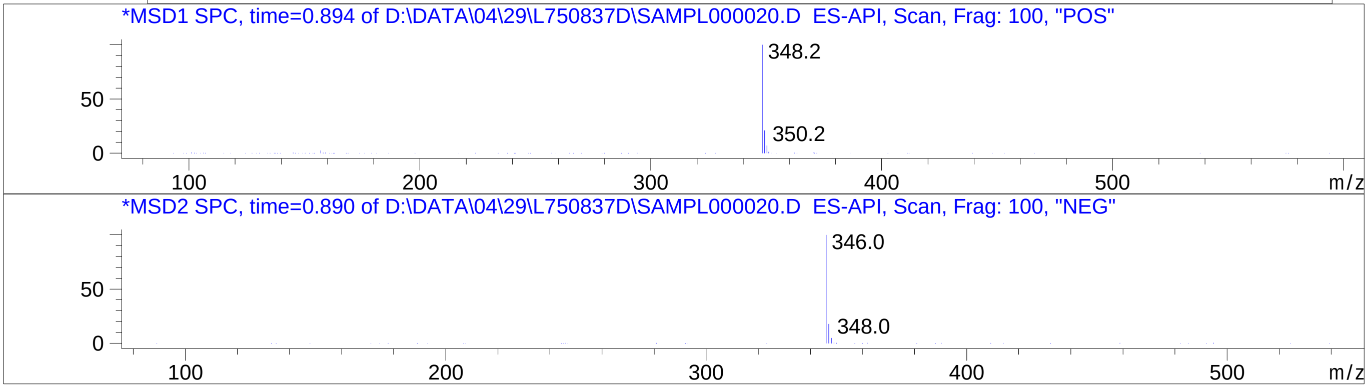


Inj.Date 4/29/2024 CH -6-
C:\CHEM32\1\METHODS\SUPOR_30.M

### Z7718740232.PDF

MaxPeak: 100.00%
Ret_Time: 1.060 min

# BE177074$2

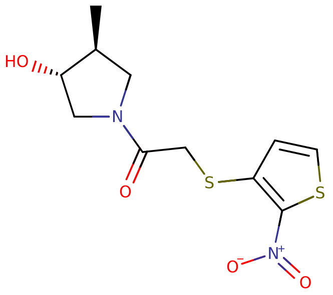


**Mol Wt** 302.37

**Exact Mass** 302.04

| # | Time | Area% |
|---|---|---|
| 1 | 1.060 | 100.00 |

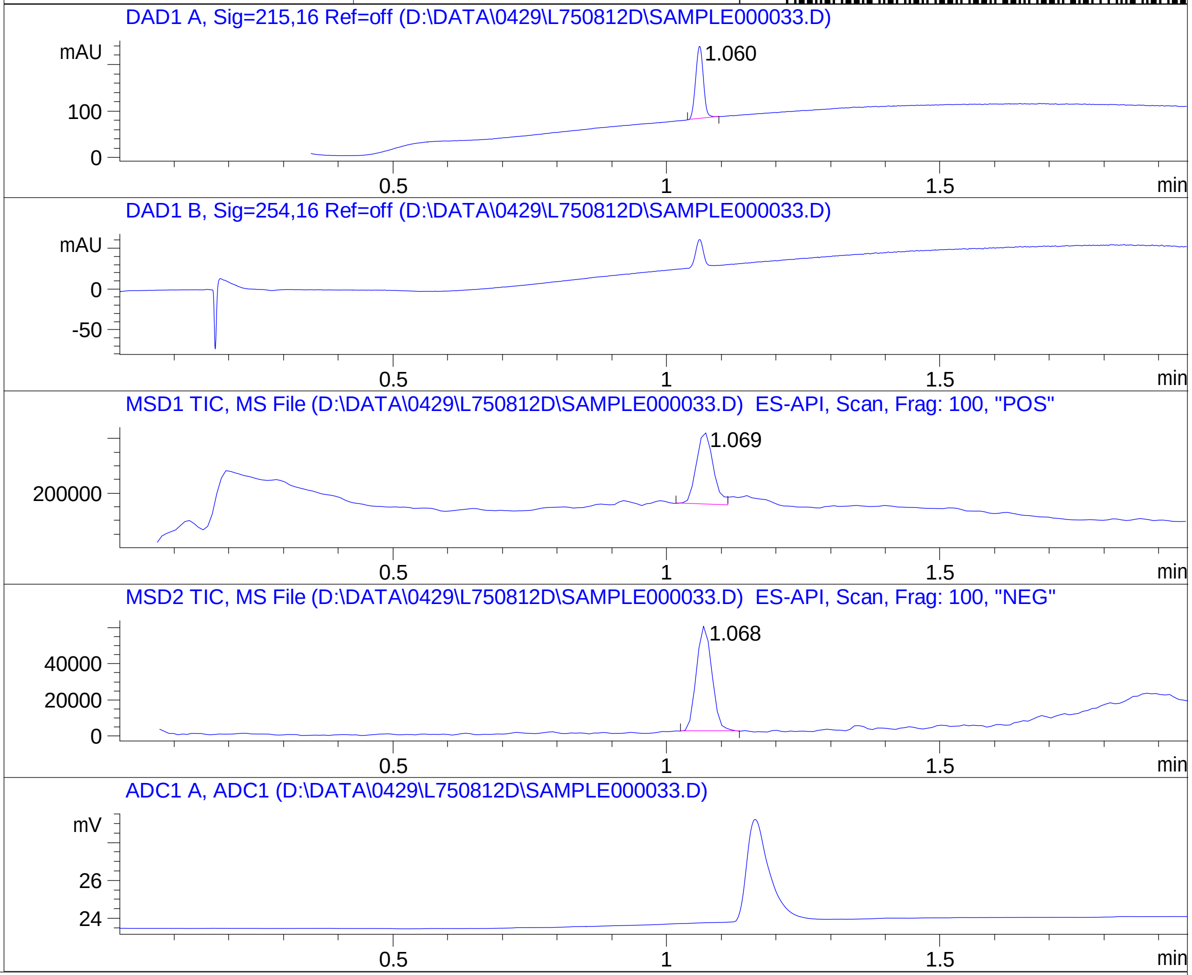


RT 1.069

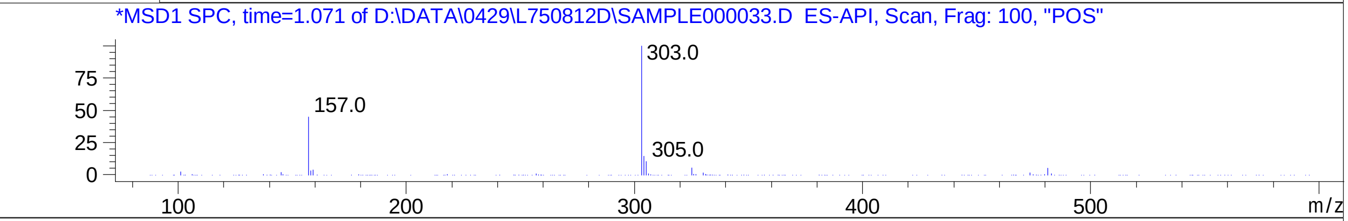


RT 1.068

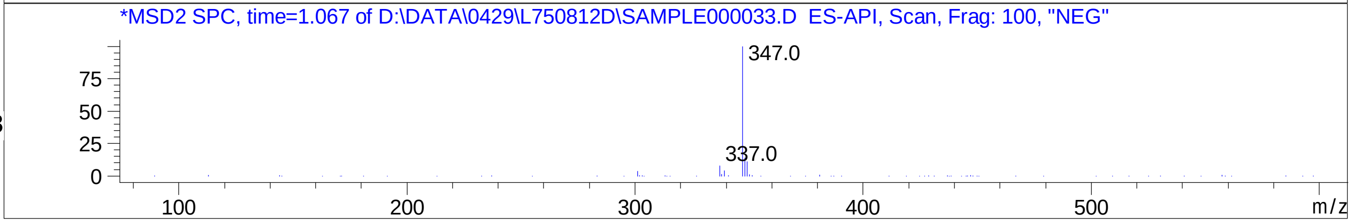

### Z8823222222.PDF

MaxPeak: 91.33%
Ret_Time: 0.874 min

# BE177046$2

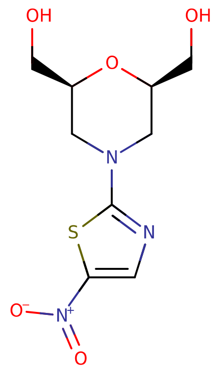


**Mol Wt** 275.28

**Exact Mass** 275.05

| # | Time | Area% |
|---|---|---|
| 1 | 0.874 | 91.33 |
| 2 | 1.627 | 8.67 |

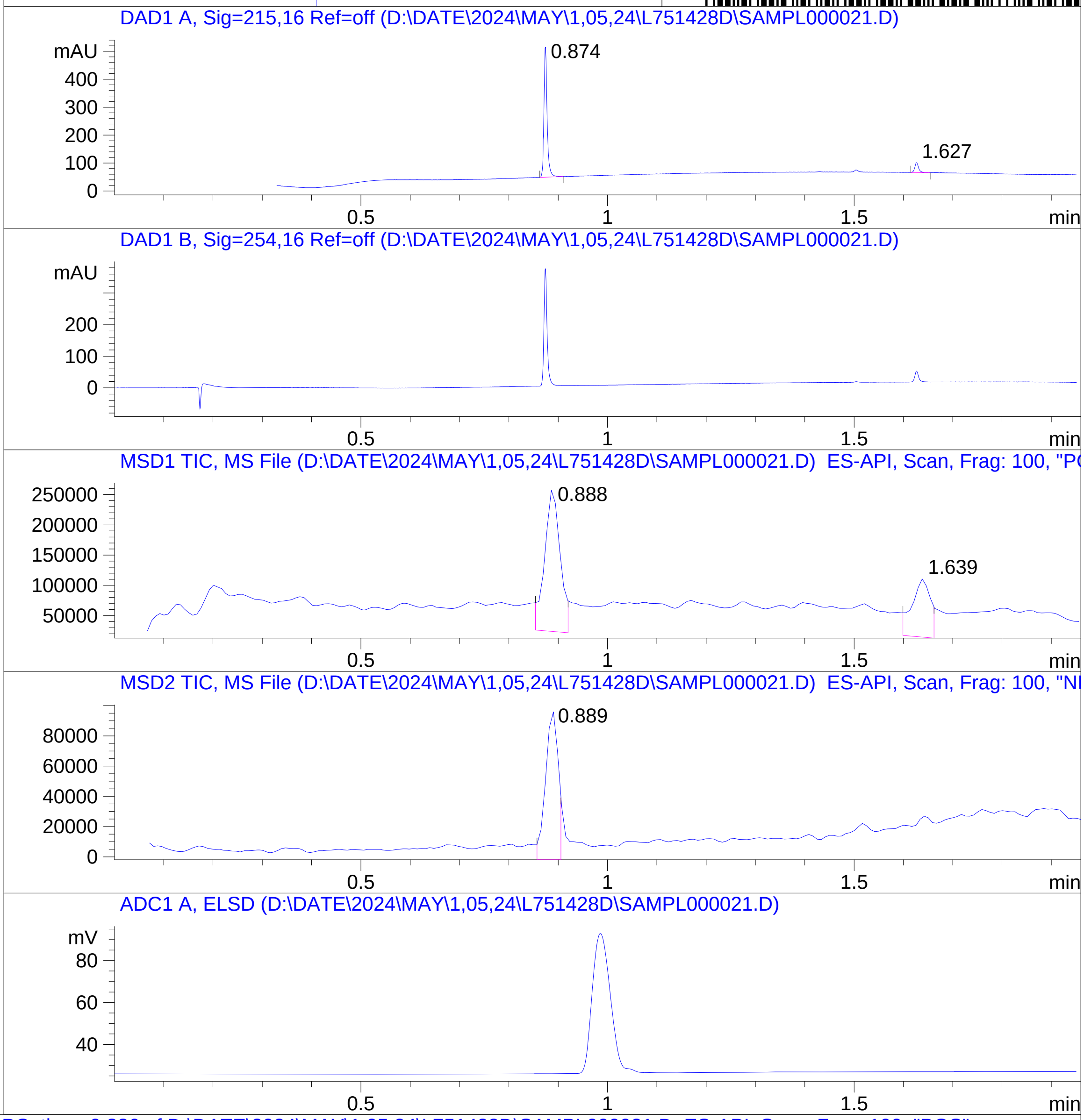


RT 0.888

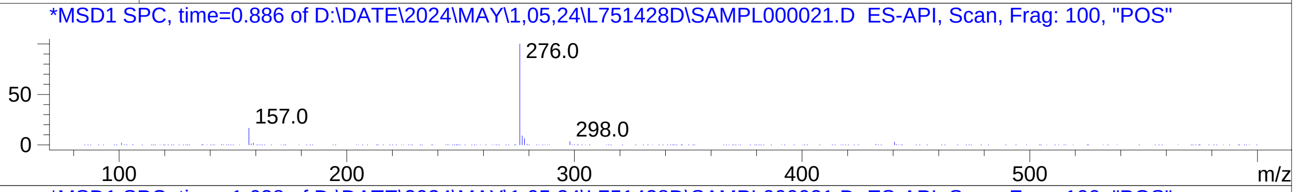


RT 1.639

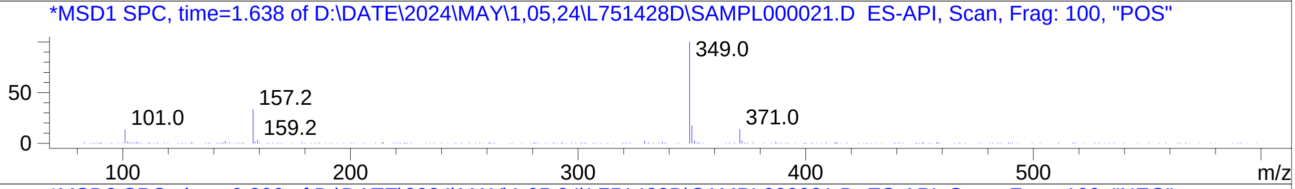


RT 0.889

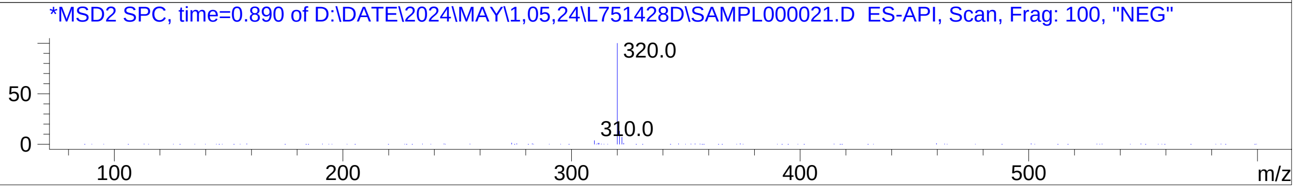

### Z8921960433.PDF

MaxPeak: 100.00%
Ret_Time: 1.158 min

# BD252929$2

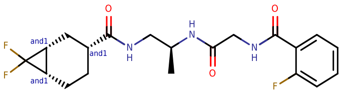


**Mol Wt** 411.42

**Exact Mass** 411.21

| # | Time | Area% |
| --- | --- | --- |
| 1 | 1.158 | 100.00 |

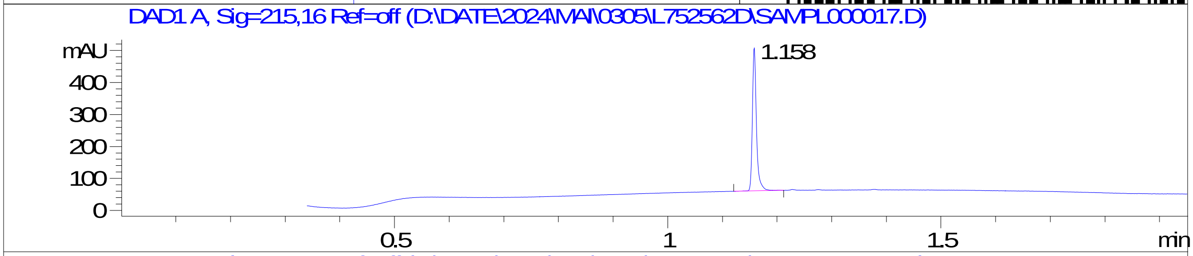


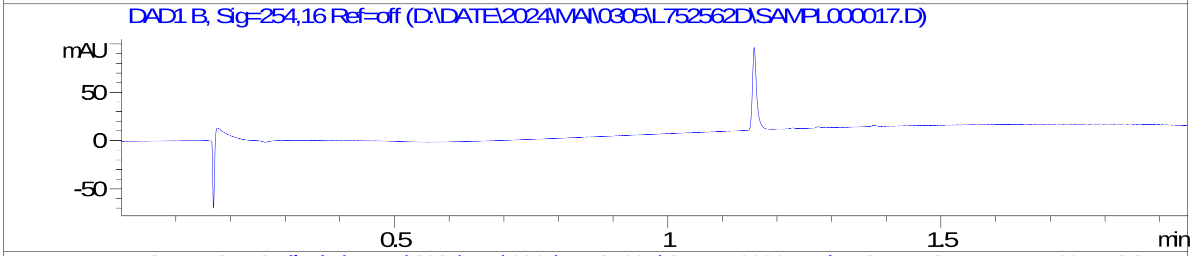


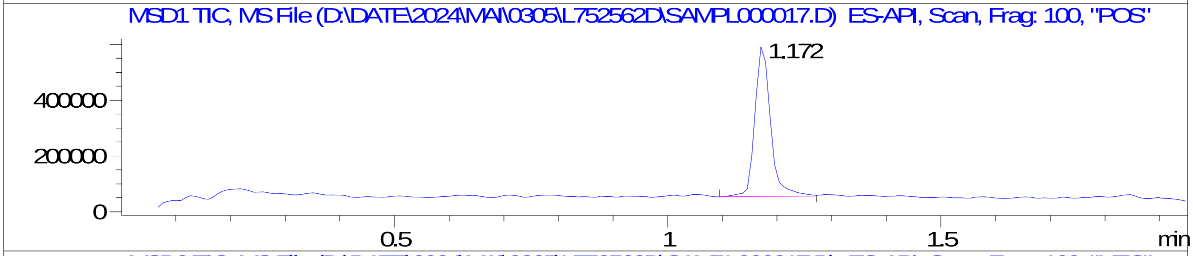


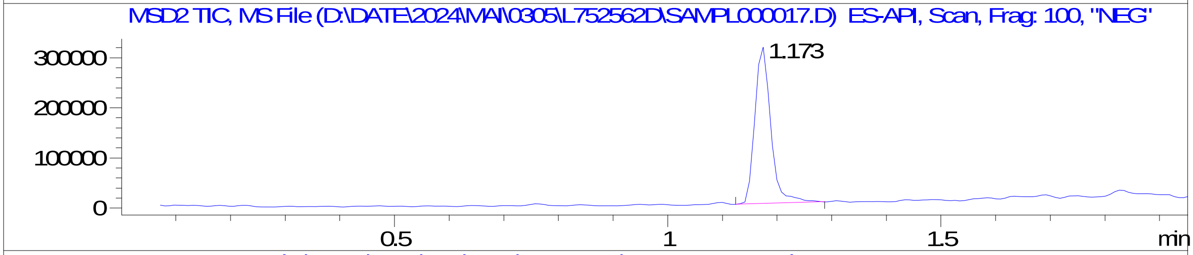


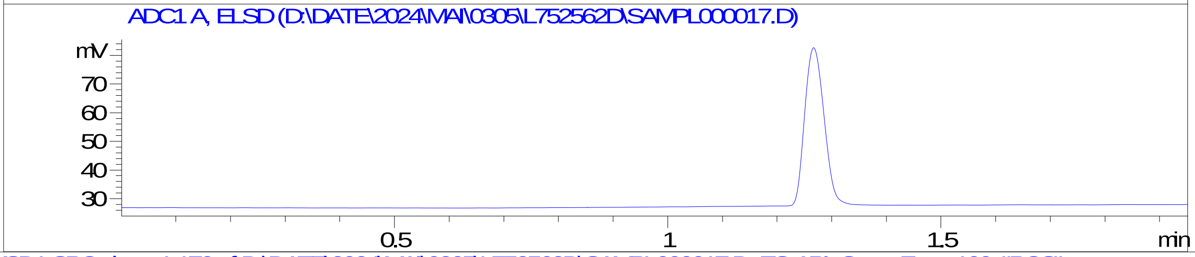


RT 1.172

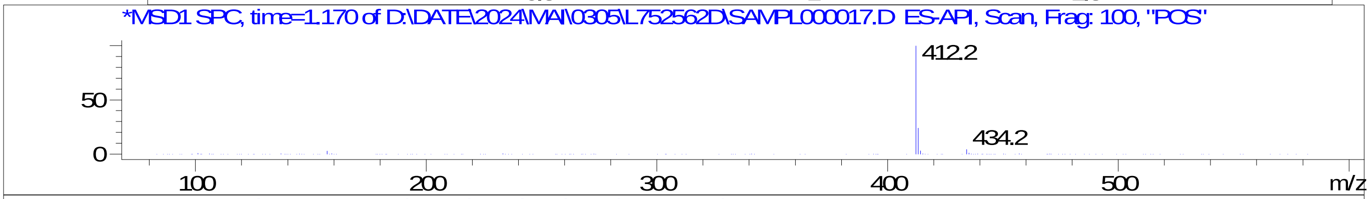


RT 1.173

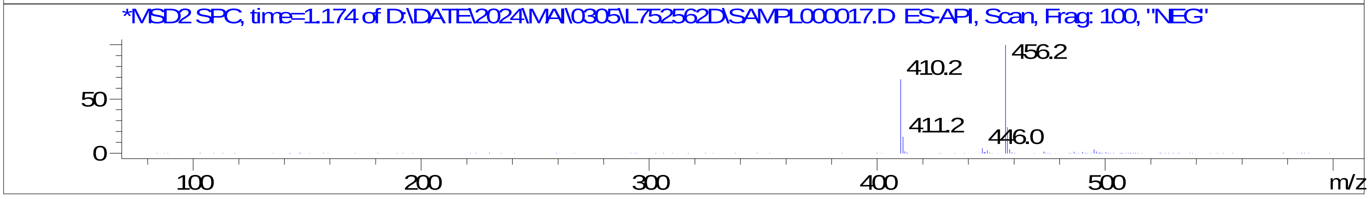

### Z8921960445.PDF

MaxPeak: 100.00%
Ret_Time: 1.172 min

# BD252745$2

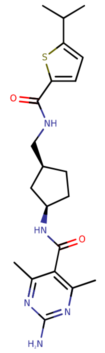

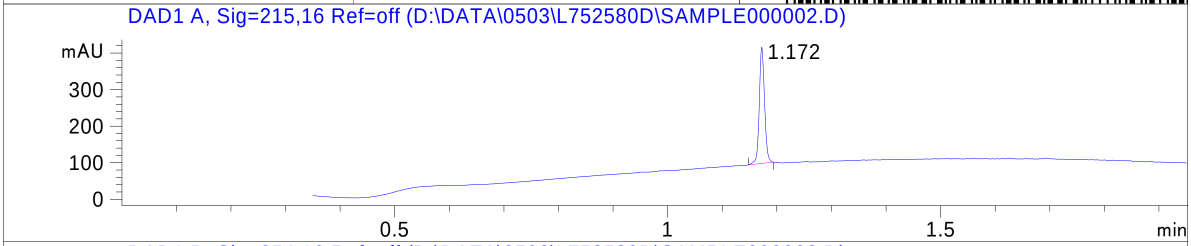


**Mol Wt** 415.55

**Exact Mass** 415.24

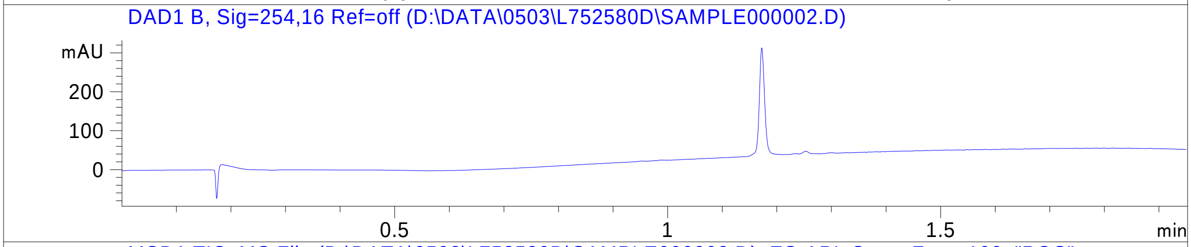


| # | Time | Area% |
|---|---|---|
| 1 | 1.172 | 100.00 |

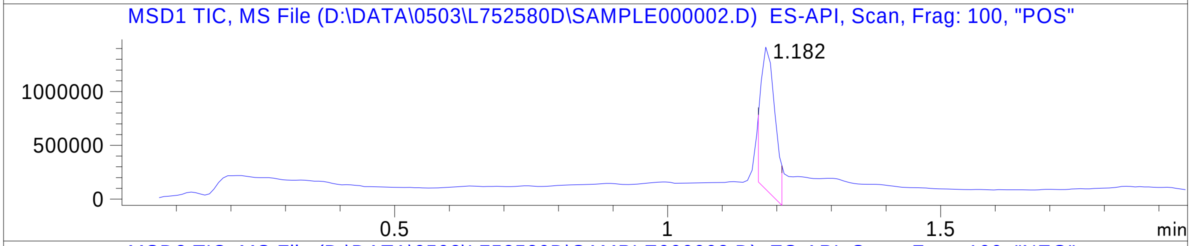


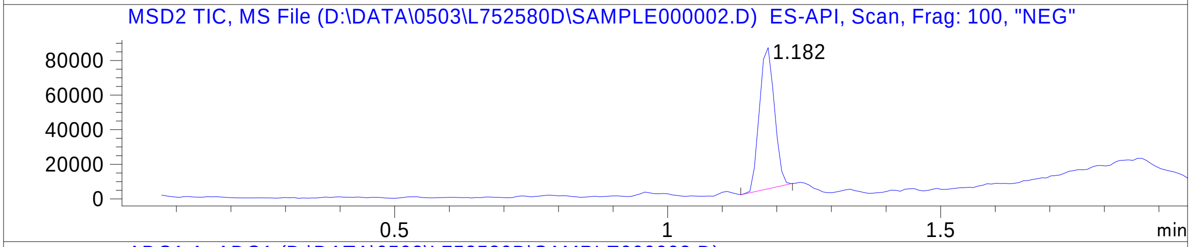


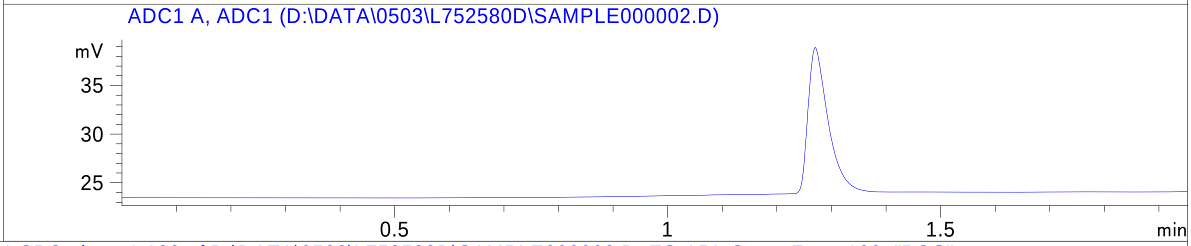


RT 1.182

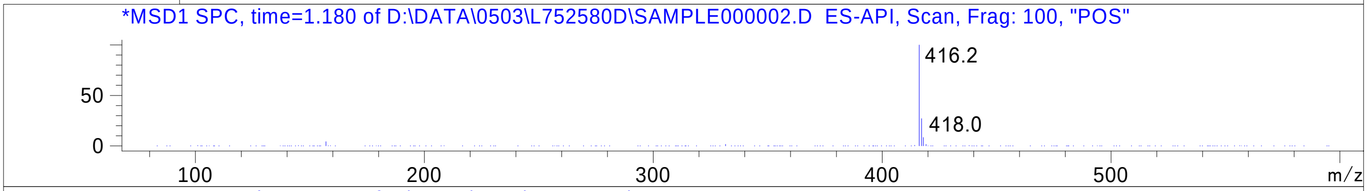


RT 1.182

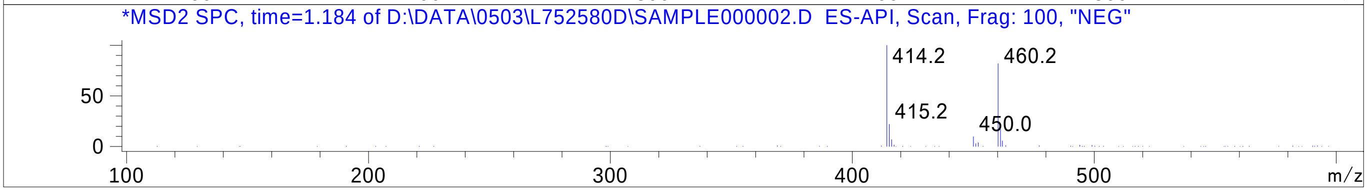

### Z8921960499.PDF

MaxPeak:  100.00%
Ret_Time: 1.326 min

# BD253003$2

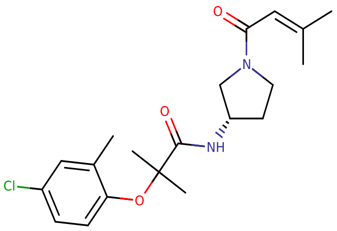


**Mol Wt** 378.89

**Exact Mass** 378.21

| # | Time | Area% |
|---|---|---|
| 1 | 1.326 | 100.00 |

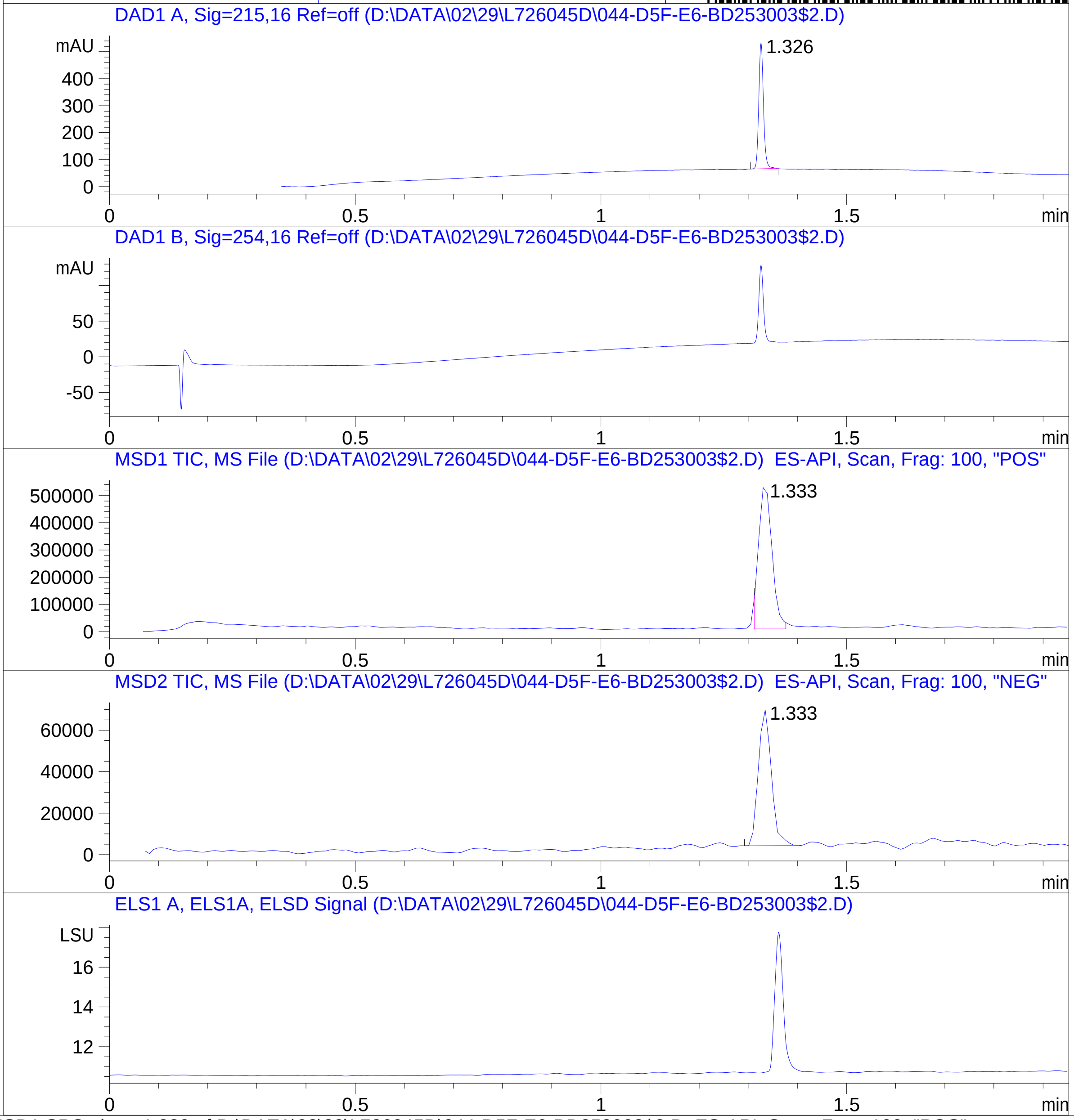


RT 1.333

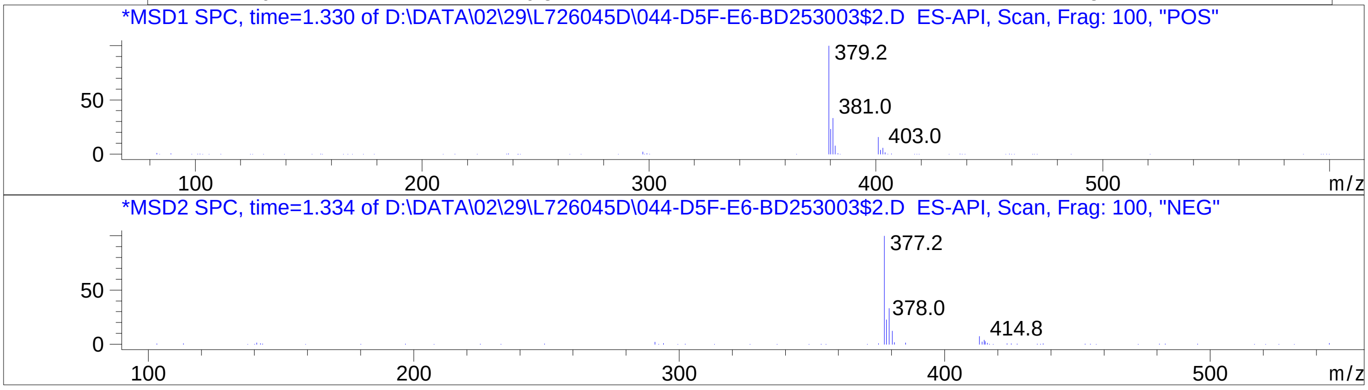


RT 1.333

Inj.Date 2/29/2024 CH -16-
C:\Chem32\1\Data\02_29\L726045D\SUPOR_30.M

### Z8921960552.PDF

MaxPeak: 100.00%
Ret_Time: 1.339 min

# BD252791$1

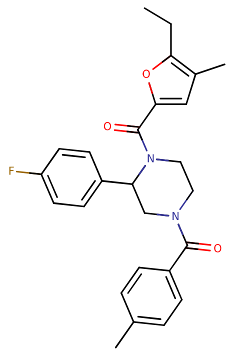


**Mol Wt** 434.5

**Exact Mass** 434.24

| # | Time | Area% |
|---|---|---|
| 1 | 1.339 | 100.00 |

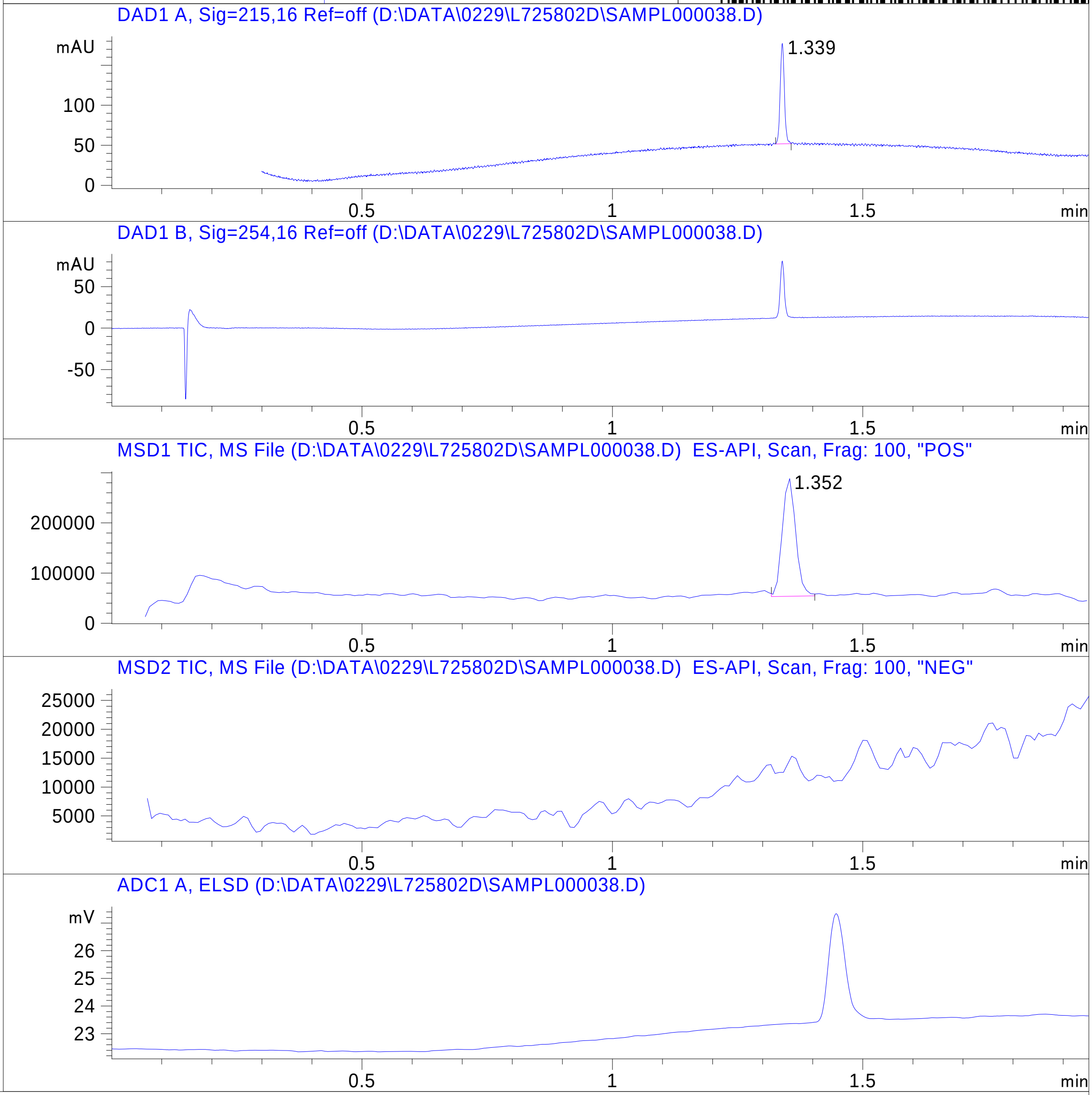


RT 1.352

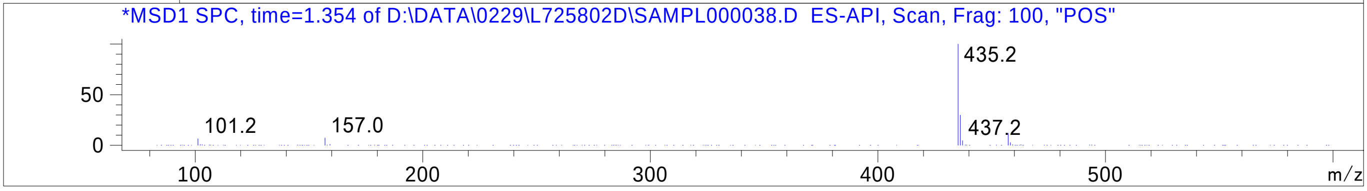


Inj.Date 2/29/2024 H -6-

C:\CHEM32\1\METHODS\SUPOR_30.M

### Z8921960554.PDF

MaxPeak: 93.22%
Ret_Time: 1.258 min

# BD252792$3

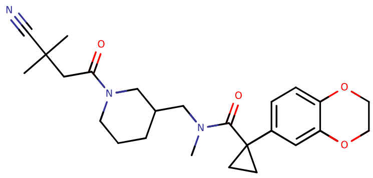

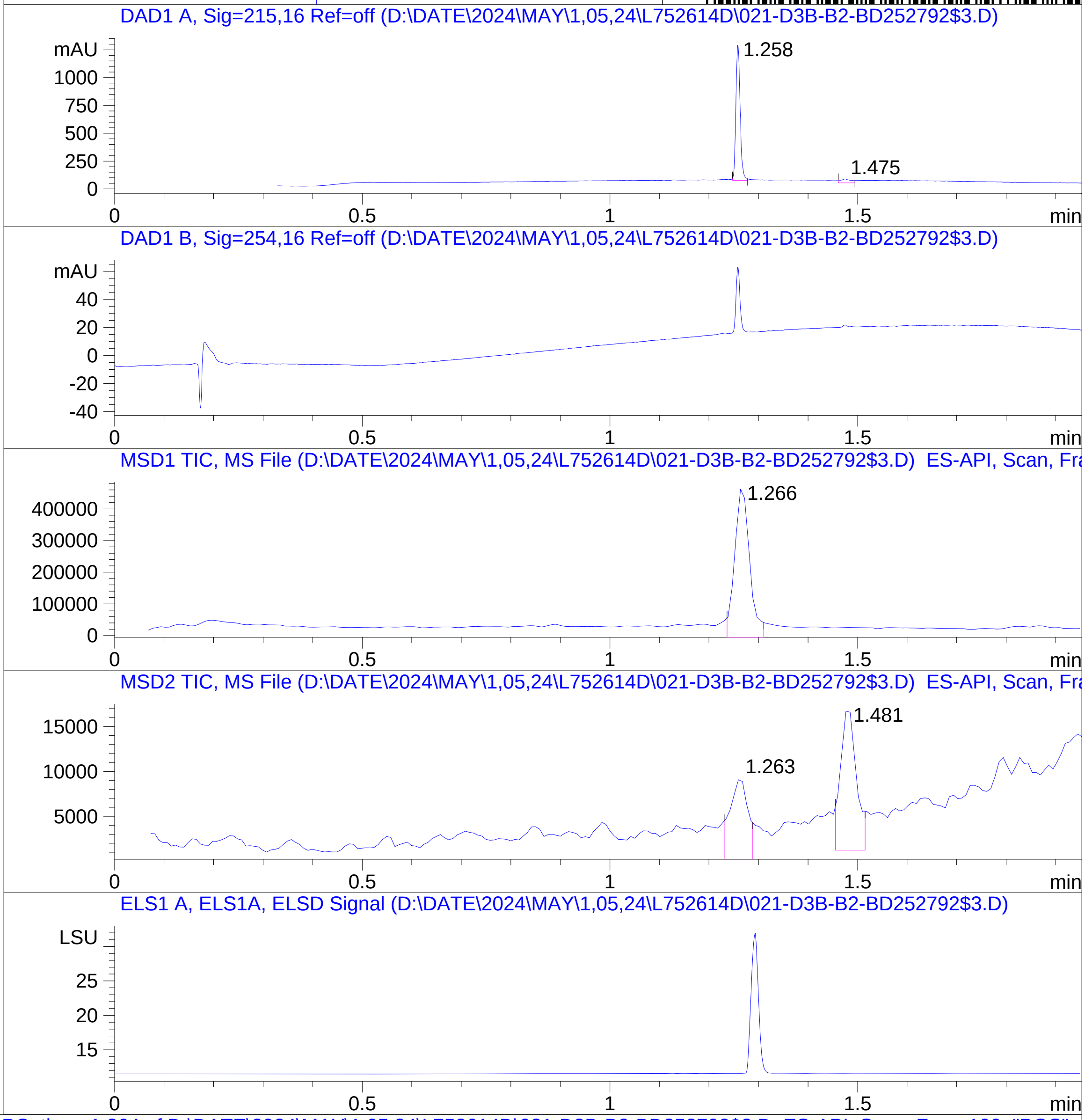


**Mol Wt** 439.55

**Exact Mass** 439.29

| # | Time | Area% |
|---|---|---|
| 1 | 1.258 | 93.22 |
| 2 | 1.475 | 6.78 |

RT 1.266

RT 1.263

RT 1.481

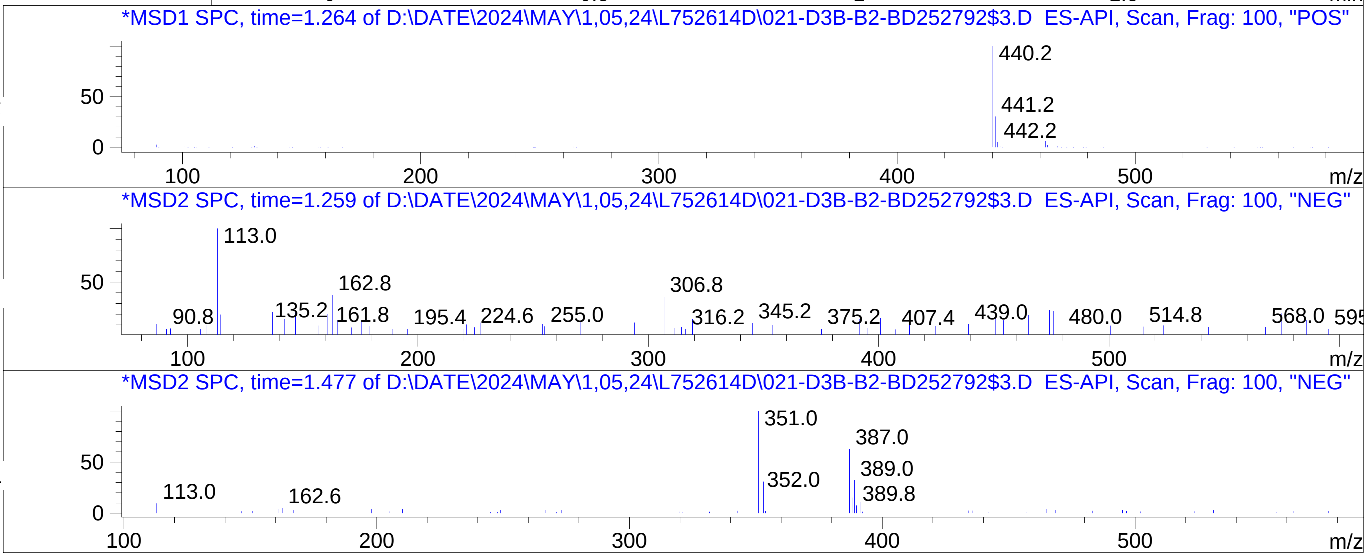



C:\Chem32\1\Data\05_02\L752614D\SUPOR_30.M

### Z8921960585.PDF

# BD252529$2

MaxPeak: 98.94%
Ret_Time: 1.361 min

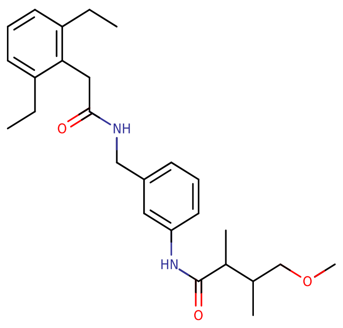


**Mol Wt** 424.58

**Exact Mass** 424.33

| # | Time | Area% |
|---|---|---|
| 1 | 1.361 | 98.94 |
| 2 | 1.419 | 1.06 |

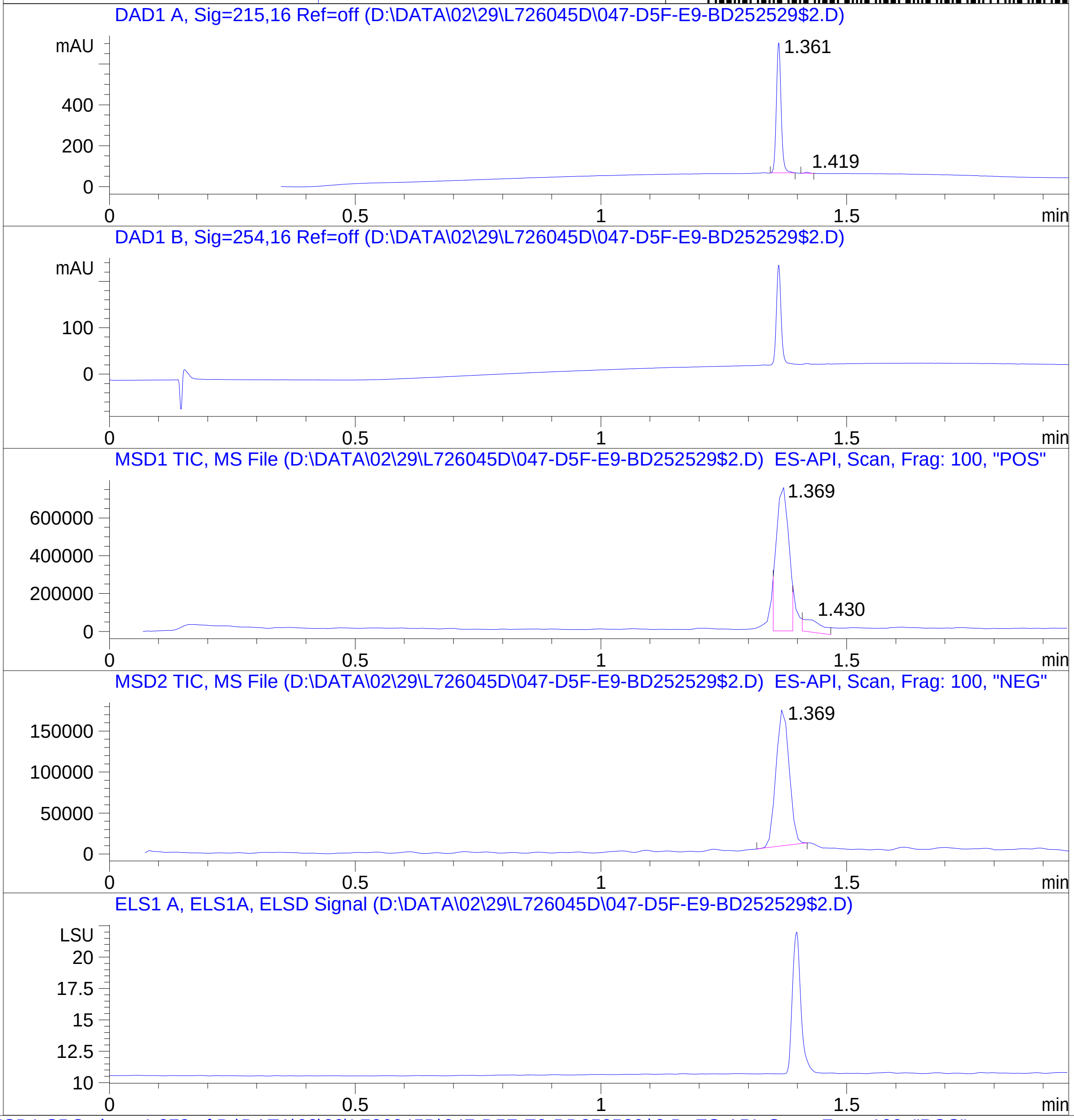


RT 1.369

RT 1.430

RT 1.369

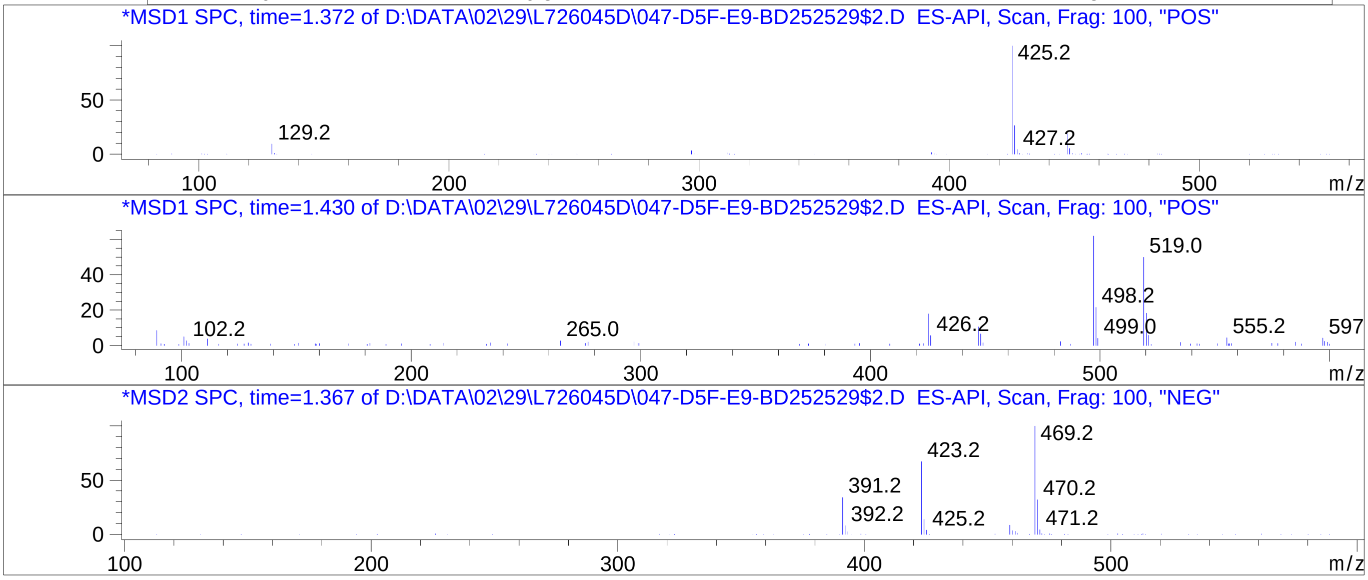


Inj.Date 2/29/2024 CH -16-
C:\Chem32\1\Data\02_29\L726045D\SUPOR_30.M

### Z8921960588.PDF

MaxPeak: 100.00%
Ret_Time: 1.108 min

# BD252512$2

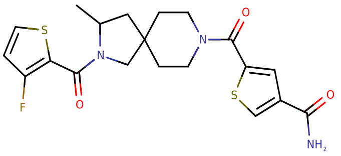


**Mol Wt** 435.54

**Exact Mass** 435.13

| # | Time | Area% |
|---|---|---|
| 1 | 1.108 | 100.00 |

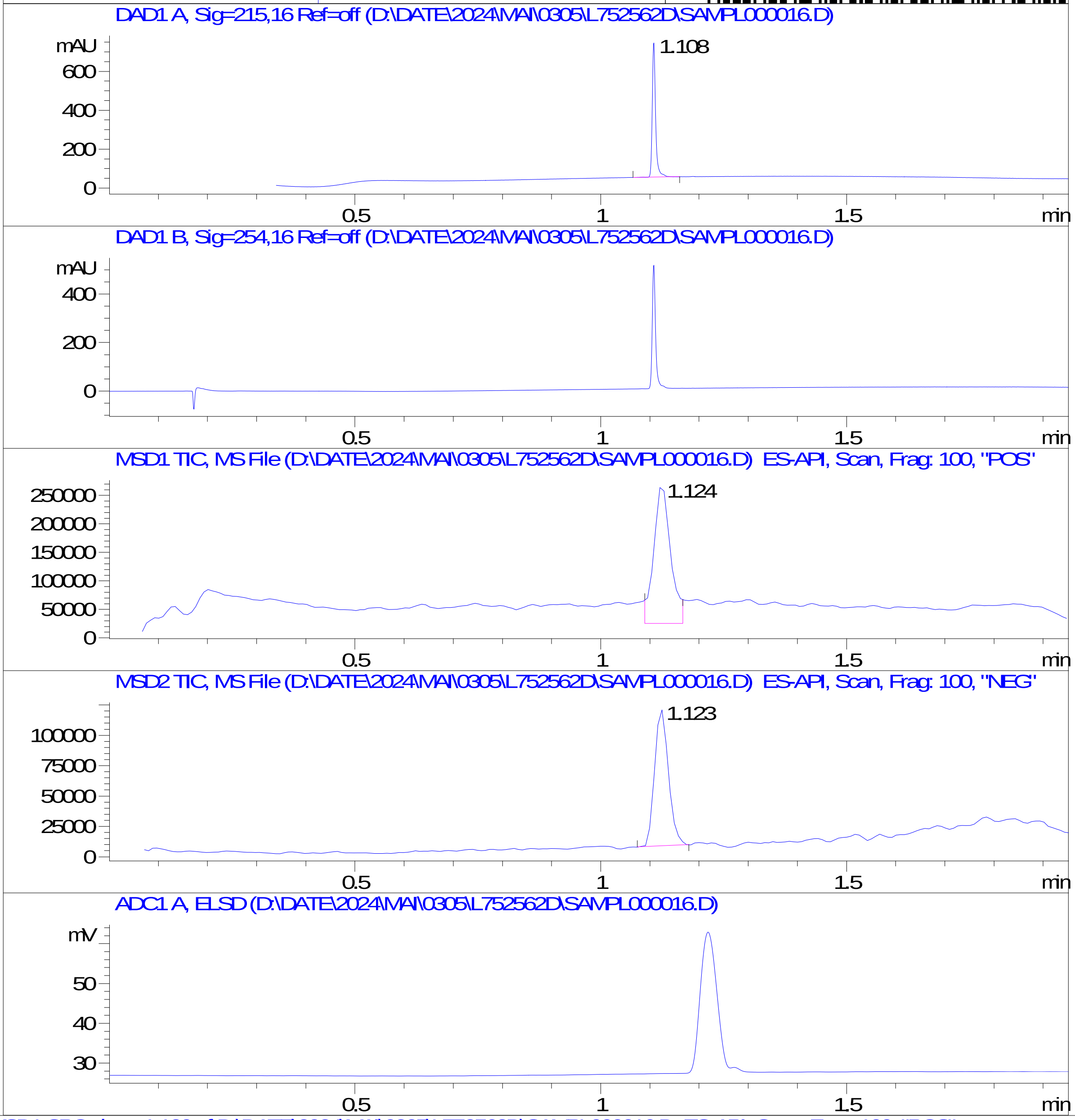


RT 1.124

RT 1.123

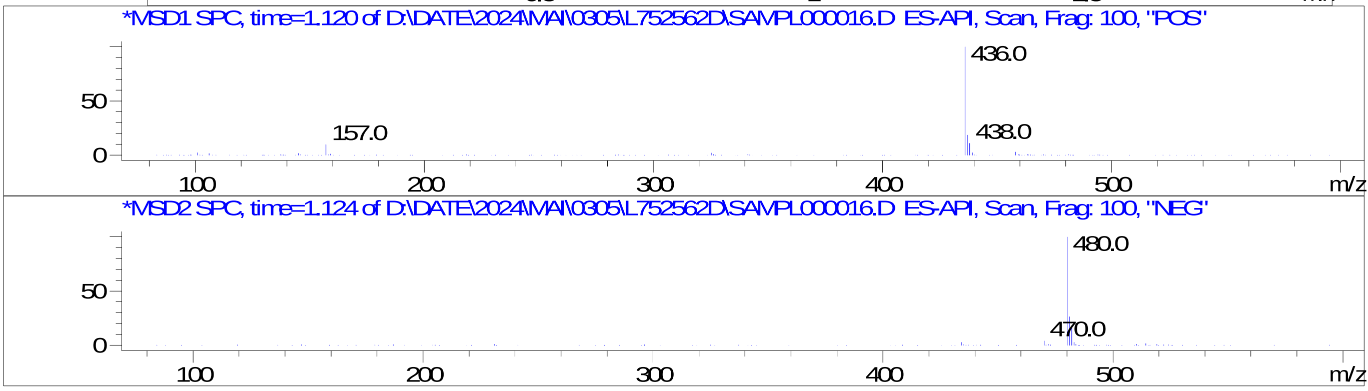

### Z8921960612.PDF

MaxPeak: 98.16%
Ret_Time: 1.213 min

# BD252966$1

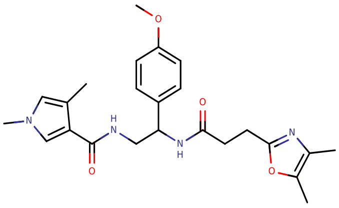


**Mol Wt** 438.52

**Exact Mass** 438.26

| # | Time | Area% |
| --- | --- | --- |
| 1 | 1.180 | 1.84 |
| 2 | 1.213 | 98.16 |

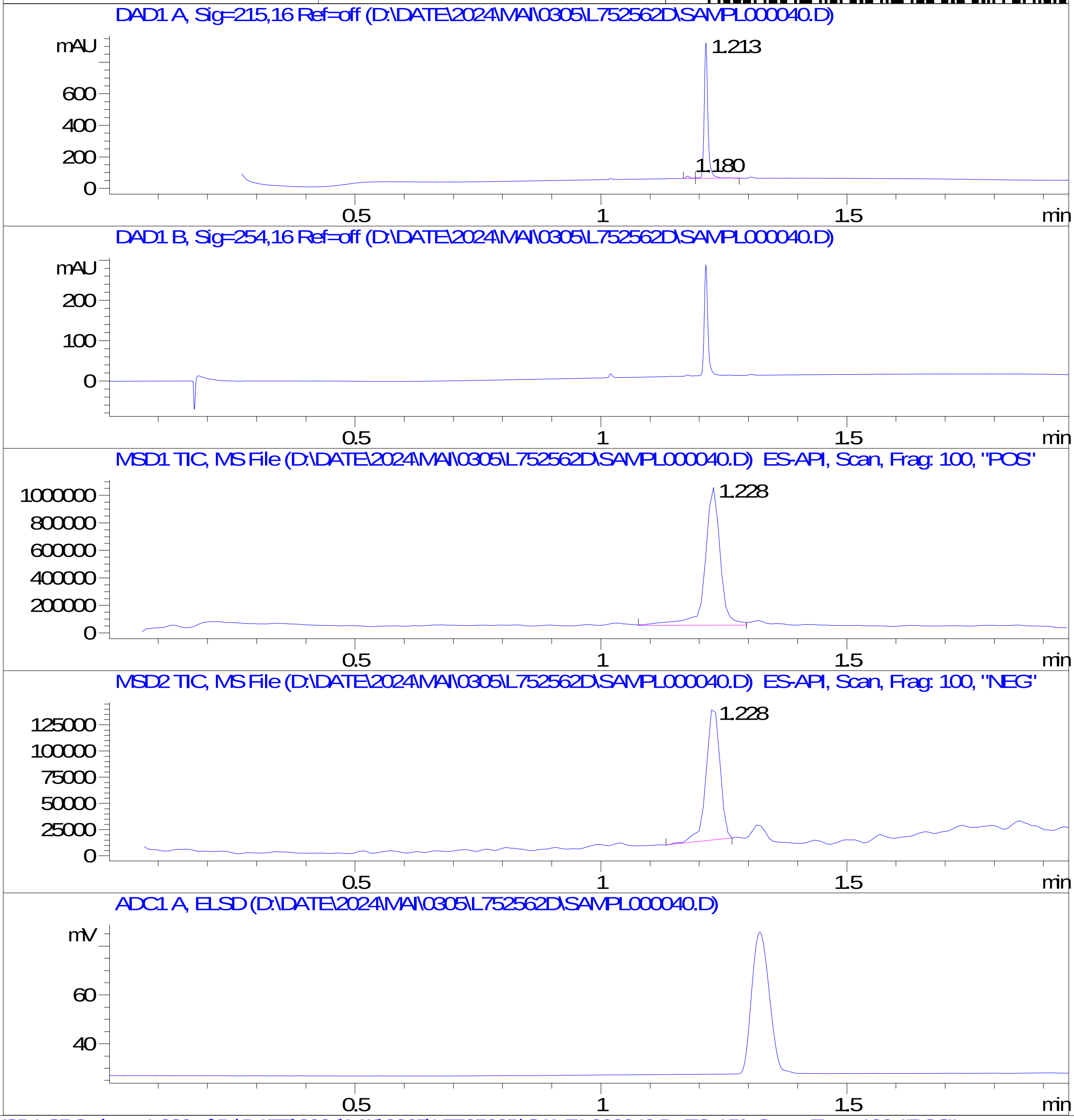


RT 1.228

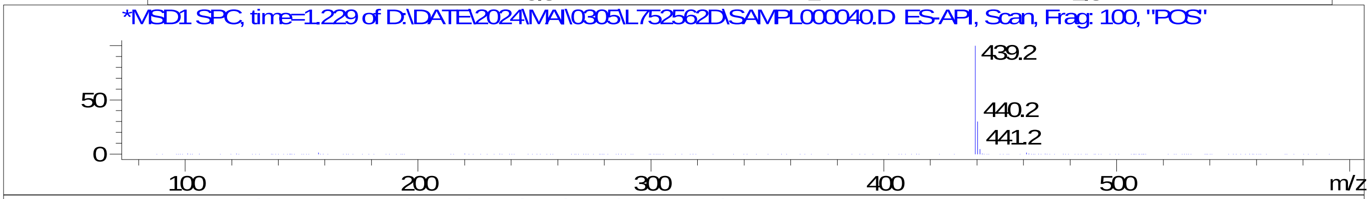


RT 1.228

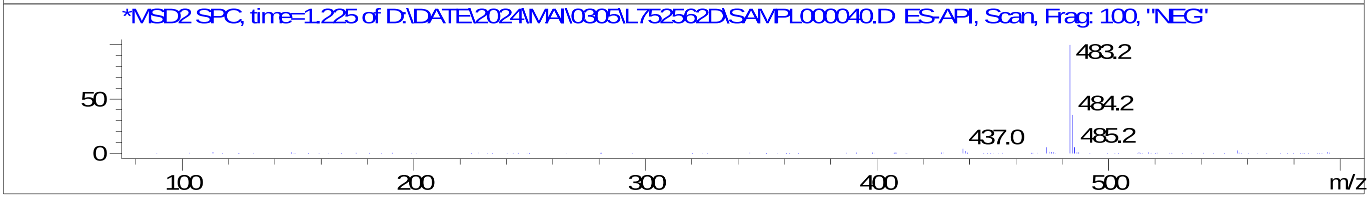

### Z8921960677.PDF

# BD252819$1

MaxPeak: 100.00%
Ret_Time: 0.903 min

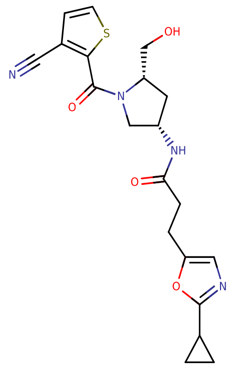


**Mol Wt** 414.48

**Exact Mass** 414.15

| # | Time | Area% |
|---|---|---|
| 1 | 0.903 | 100.00 |

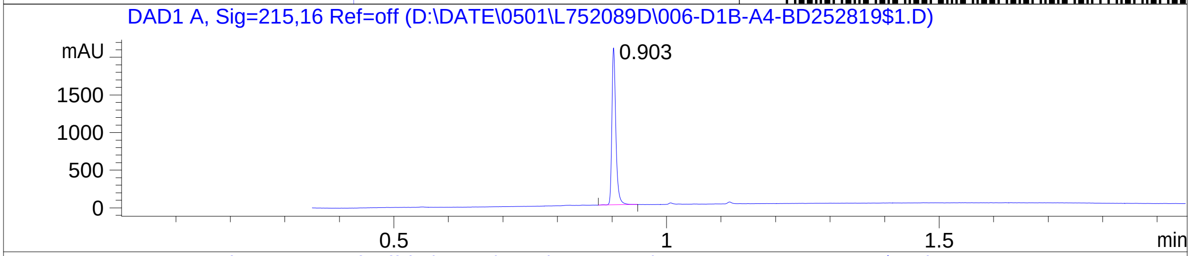


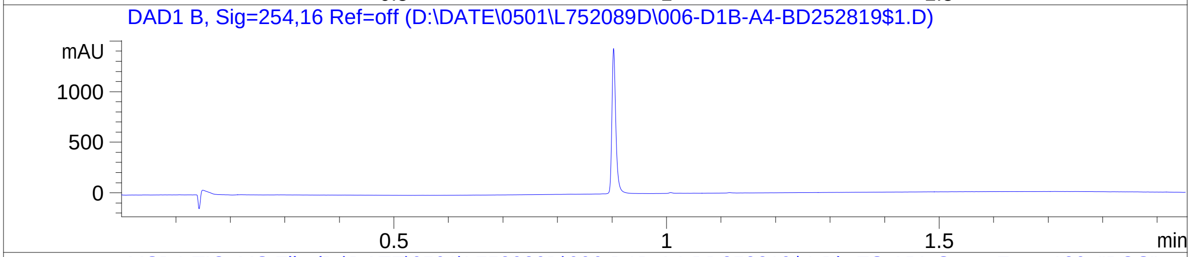


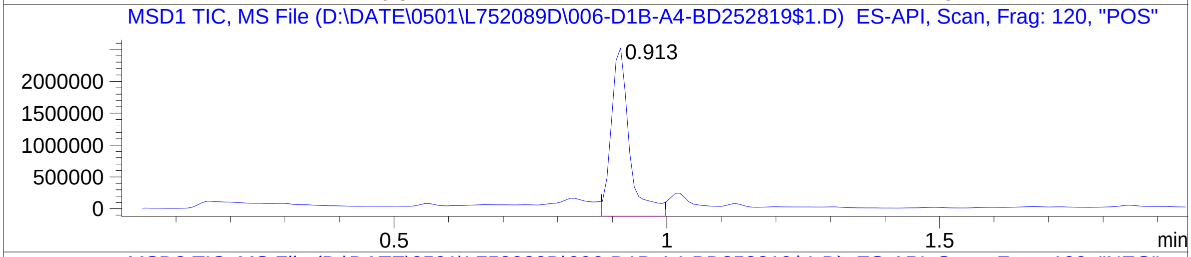


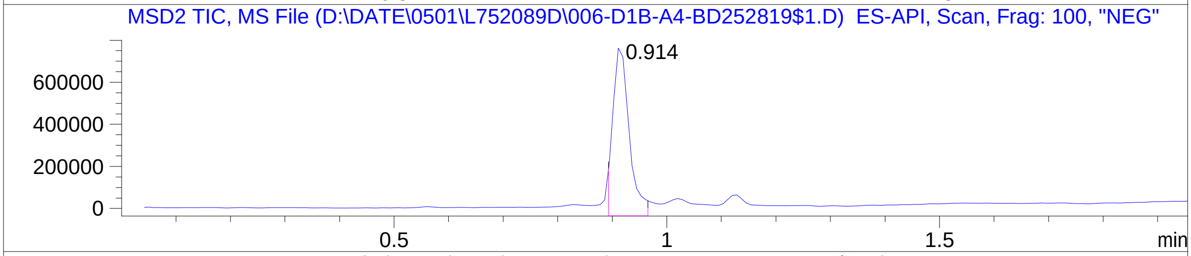


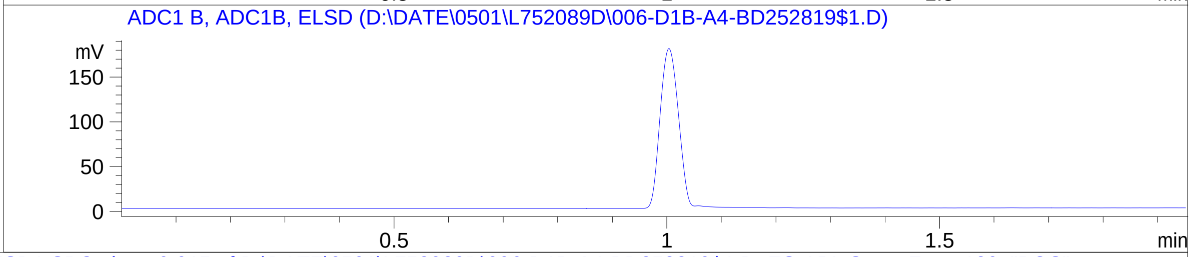


RT 0.913

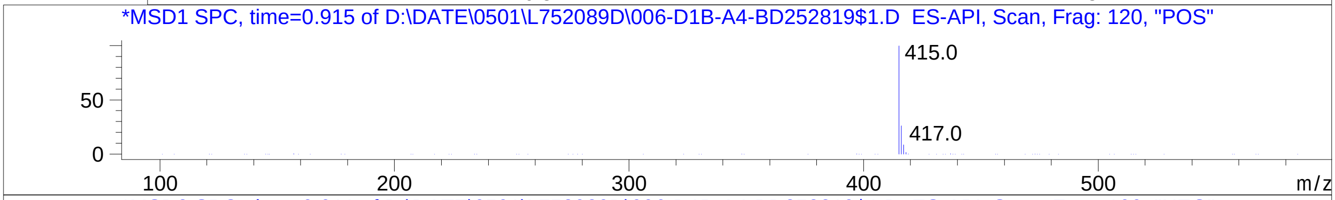


RT 0.914

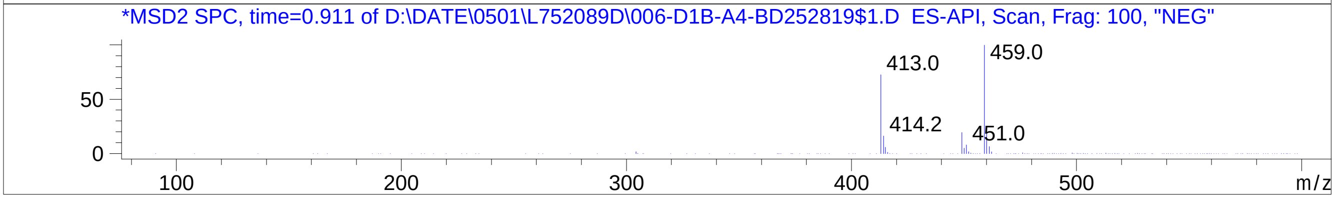

### Z8921960716.PDF

# BD252821$3

MaxPeak: 100.00%
Ret_Time: 1.172 min

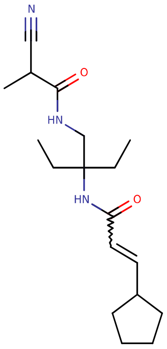


**Mol Wt** 319.44

**Exact Mass** 319.27

| # | Time | Area% |
|---|---|---|
| 1 | 1.172 | 100.00 |

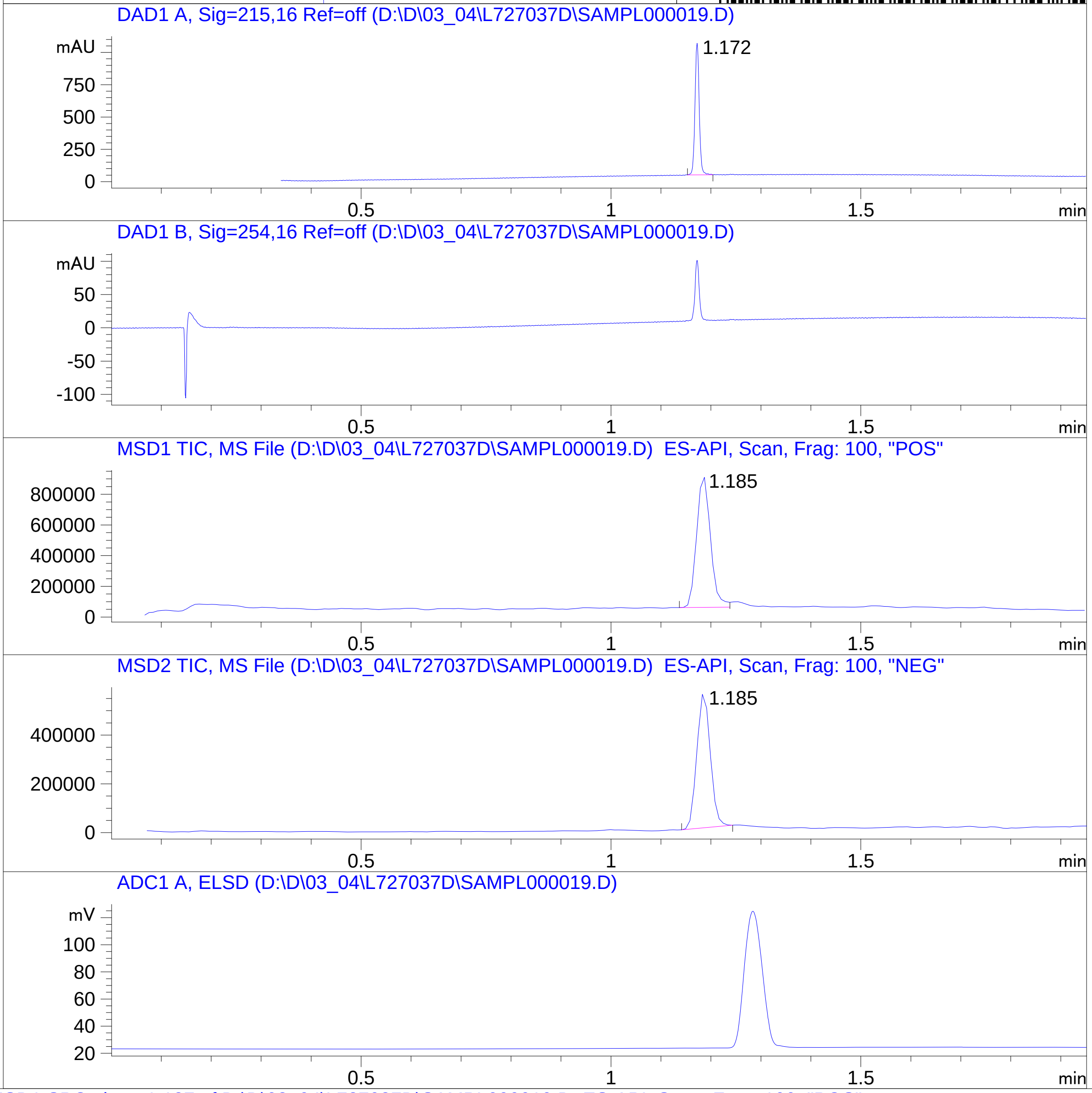


RT 1.185

RT 1.185

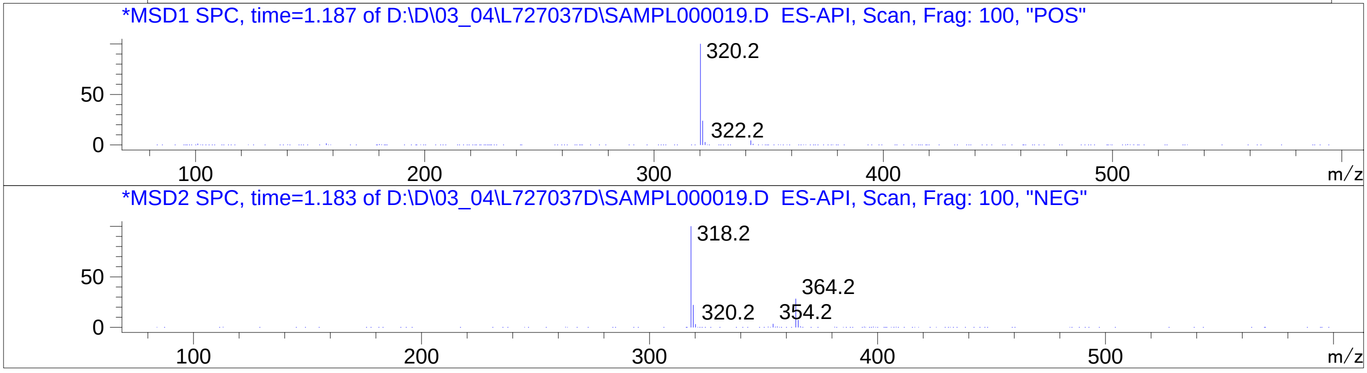

### Z8921960725.PDF

# BD252973$2

MaxPeak: 100.00%
Ret_Time: 1.011 min

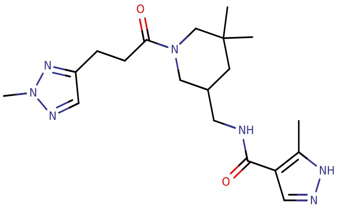

**Mol Wt** 387.48
**Exact Mass** 387.27

| # | Time | Area% |
|---|---|---|
| 1 | 1.011 | 100.00 |

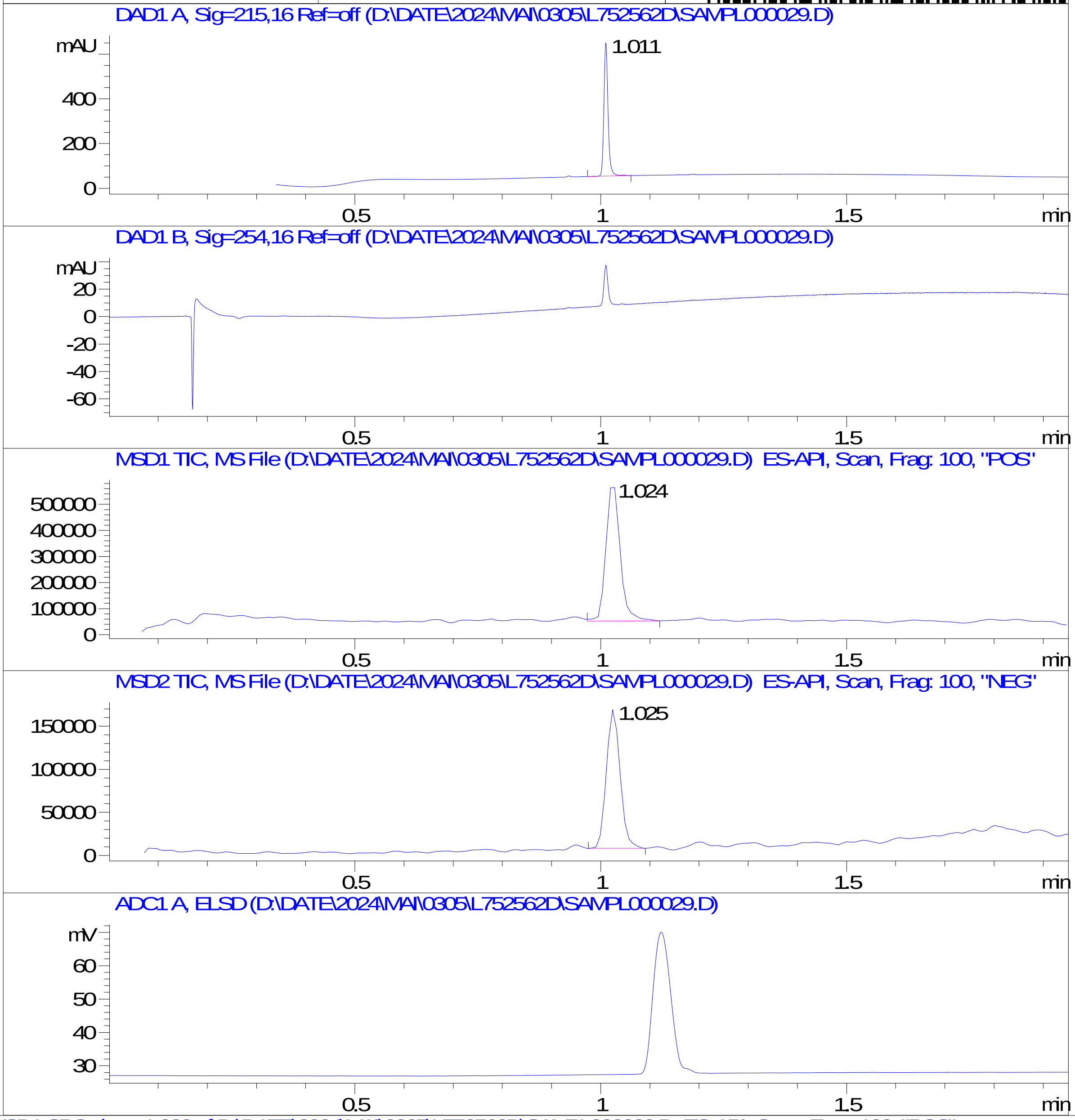


RT 1.024

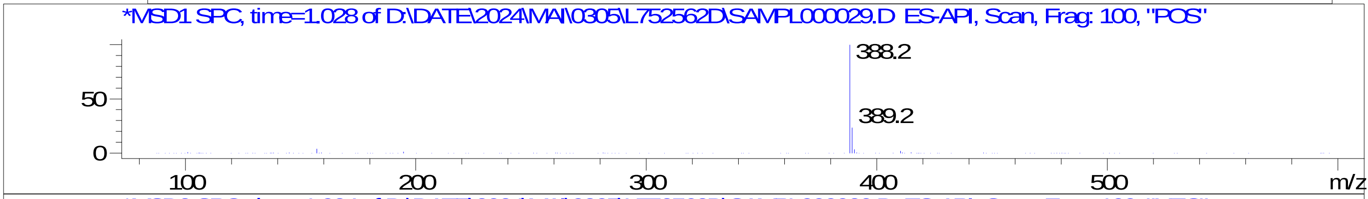


RT 1.025

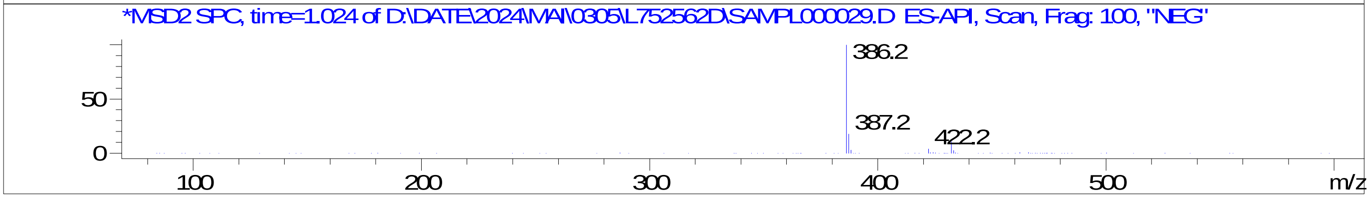

### Z8921960729.PDF

MaxPeak: 91.80%
Ret_Time: 1.191 min

BD252795$2

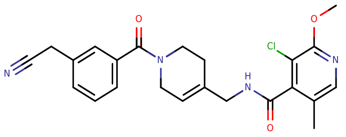


**Mol Wt** 438.91

**Exact Mass** 438.17

| # | Time | Area% |
|---|---|---|
| 1 | 1.150 | 5.52 |
| 2 | 1.191 | 91.80 |
| 3 | 1.208 | 2.68 |

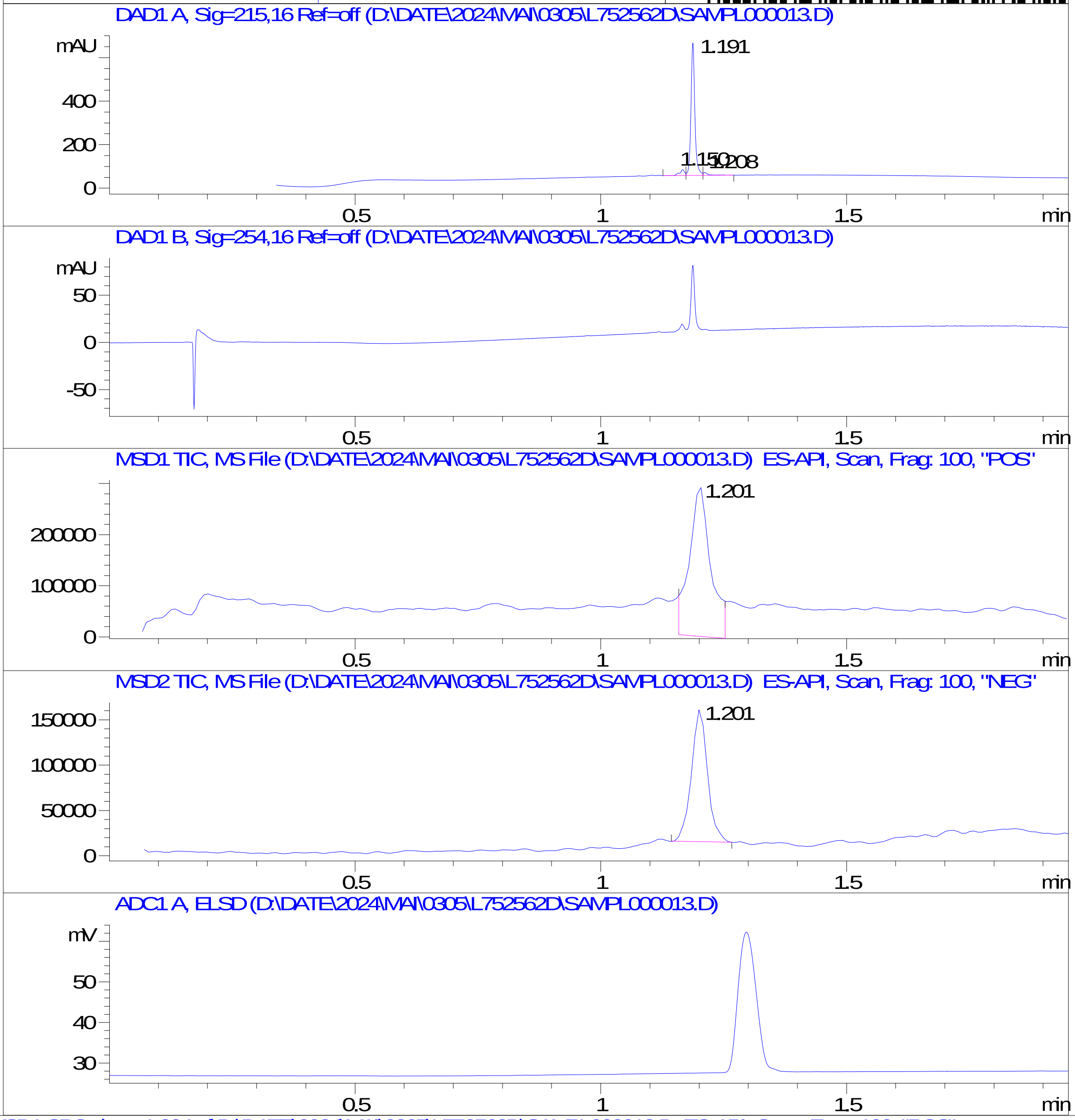


RT 1.201

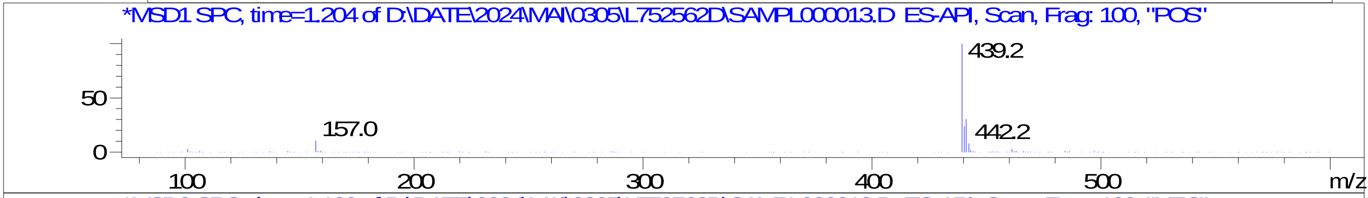


RT 1.201

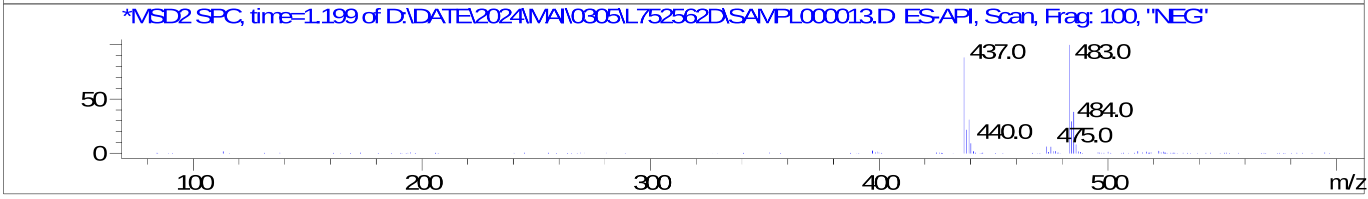

### Z8921960753.PDF

MaxPeak: 90.80%
Ret_Time: 0.733 min

# BD252974$22

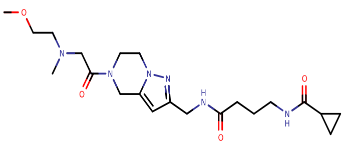

**Mol Wt** 434.53

**Exact Mass** 434.3

| # | Time | Area% |
|---|---|---|
| 1 | 0.707 | 2.15 |
| 2 | 0.733 | 90.80 |
| 3 | 0.757 | 1.38 |
| 4 | 0.905 | 5.67 |

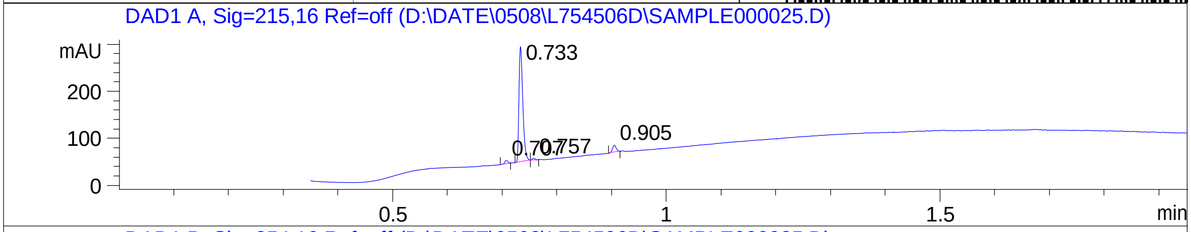


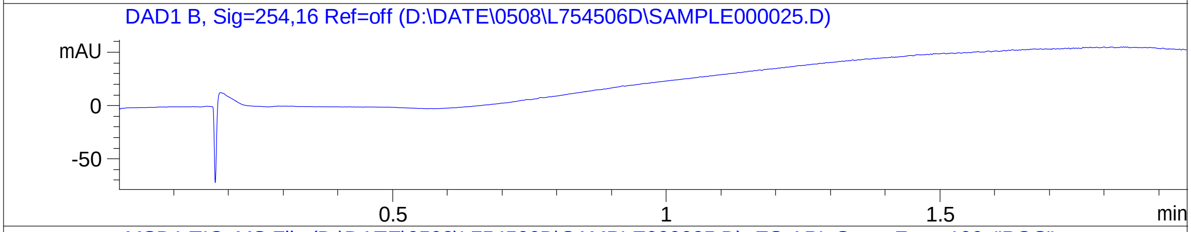


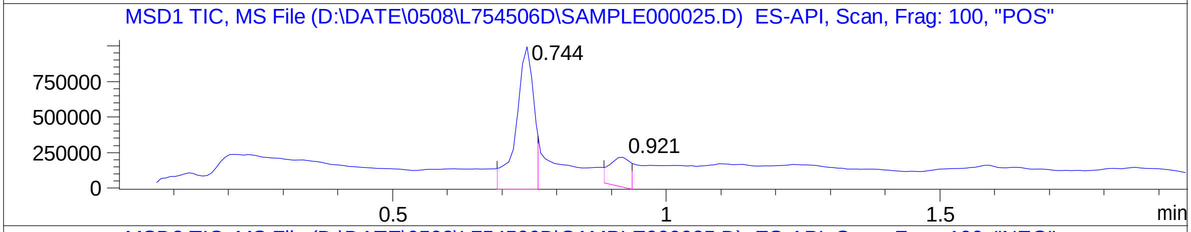


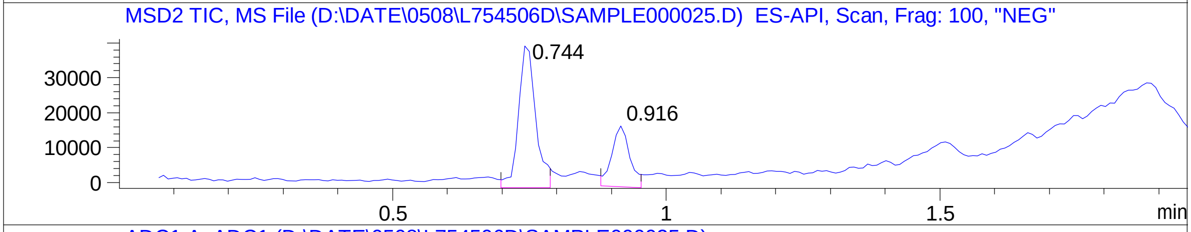


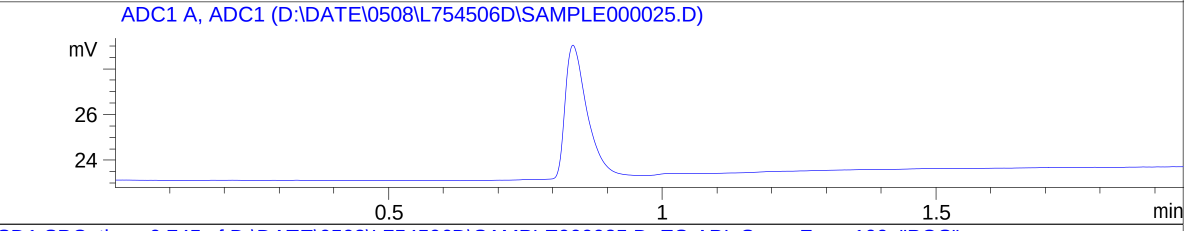


RT 0.744

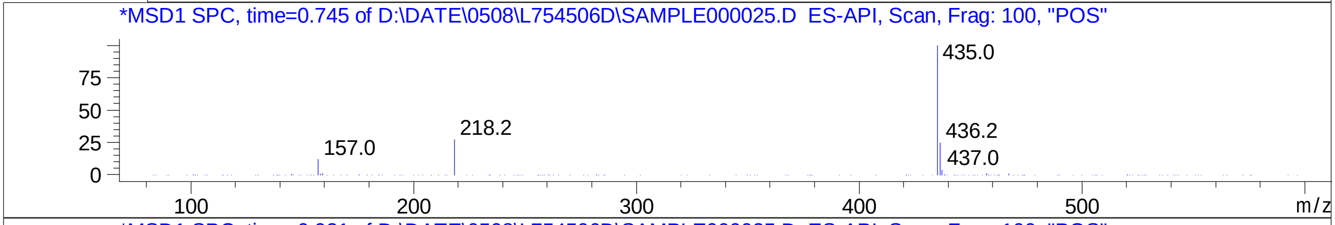


RT 0.921

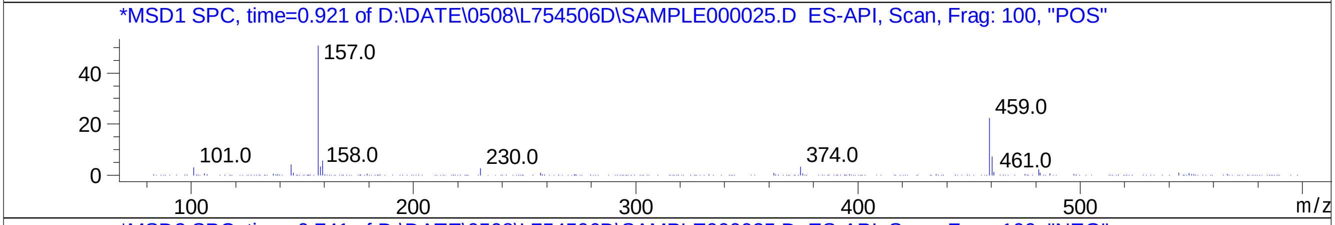


RT 0.744

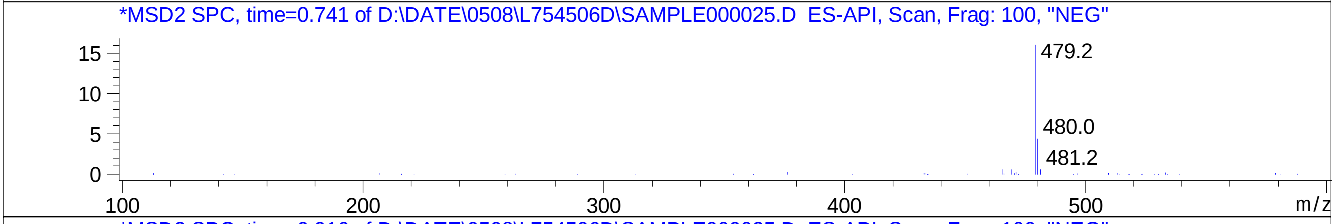


RT 0.916

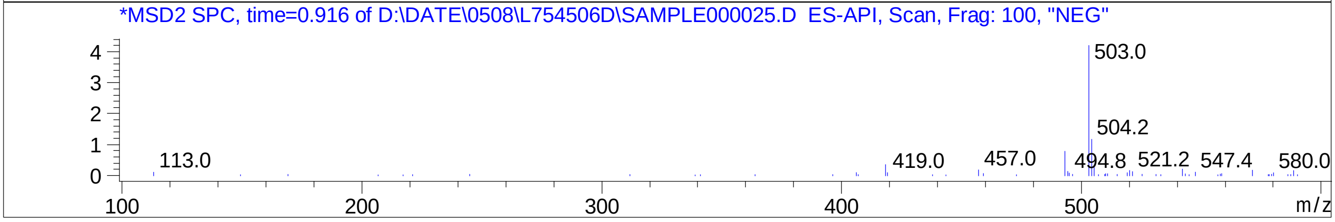

### Z8921960754.PDF

MaxPeak: 51.02%
Ret_Time: 1.495 min

# BD252856$1

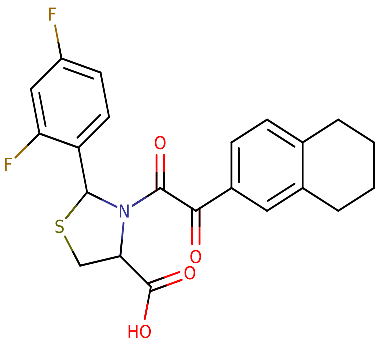


**Mol Wt** 431.45

**Exact Mass** 431.12

| # | Time | Area% |
|---|---|---|
| 1 | 1.180 | 2.01 |
| 2 | 1.423 | 1.18 |
| 3 | 1.473 | 2.37 |
| 4 | 1.495 | 51.02 |
| 5 | 1.504 | 43.41 |

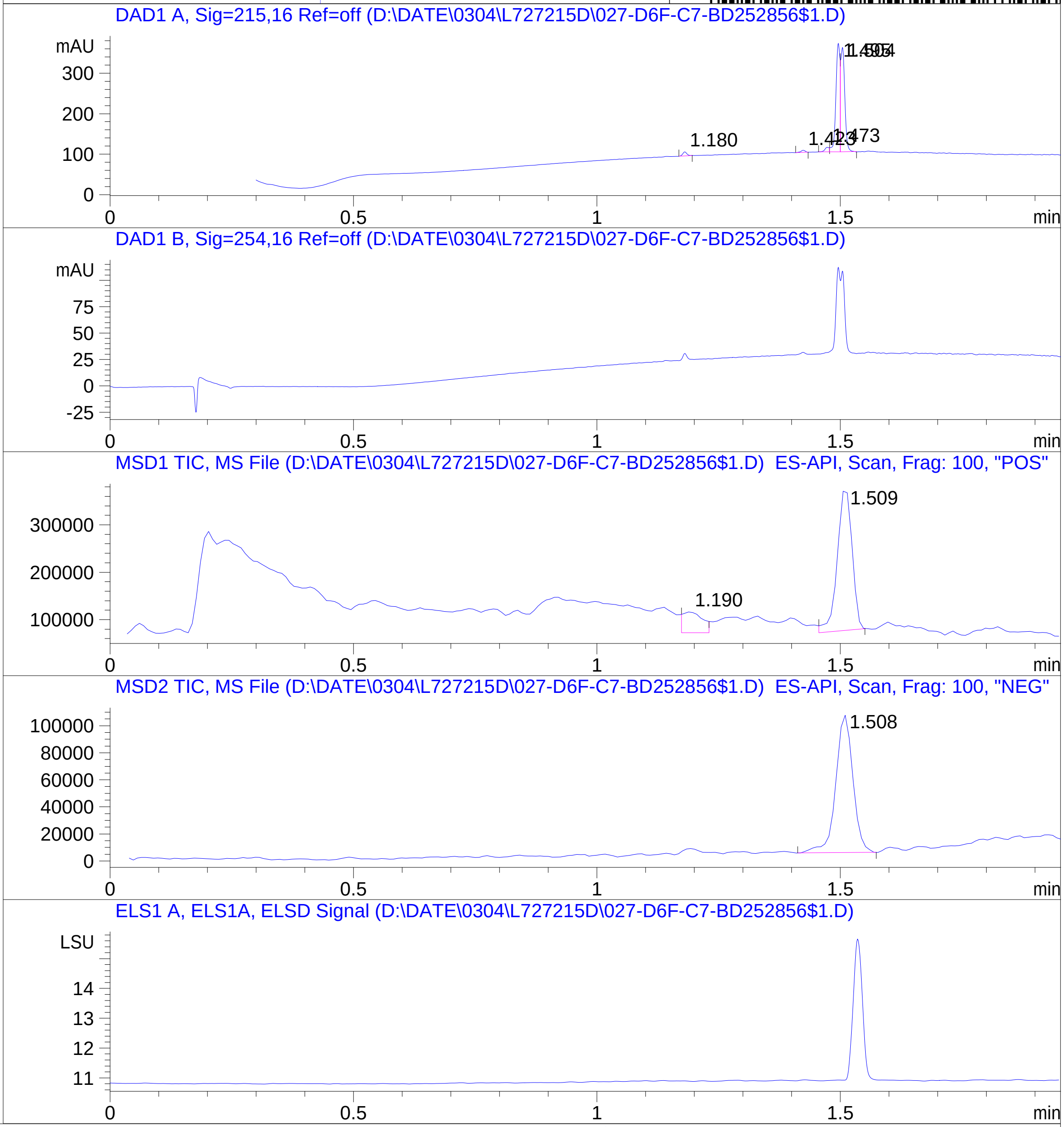


RT 1.190

RT 1.509

RT 1.508

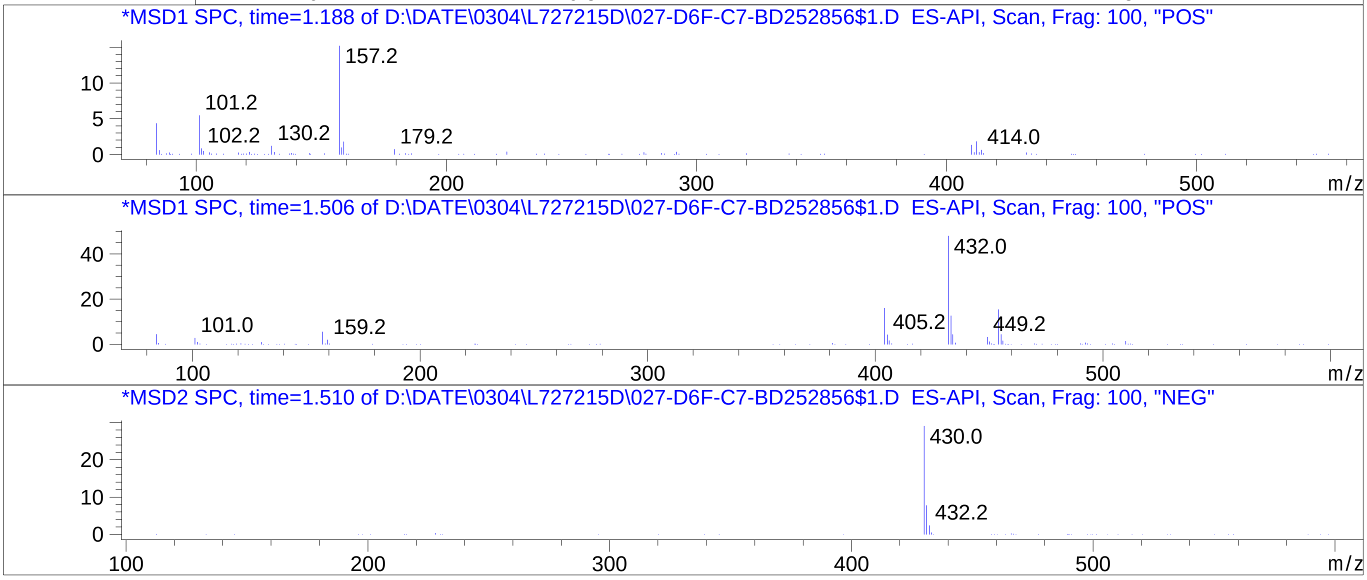

### Z8921960758.PDF

# BD252857$4

MaxPeak: 100.00%
Ret_Time: 1.170 min

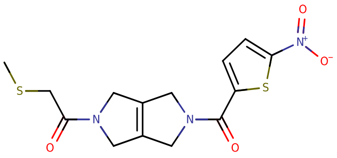

**Mol Wt** 353.42

**Exact Mass** 353.05

| # | Time | Area% |
| --- | --- | --- |
| 1 | 1.170 | 100.00 |

RT 1.180

RT 1.180

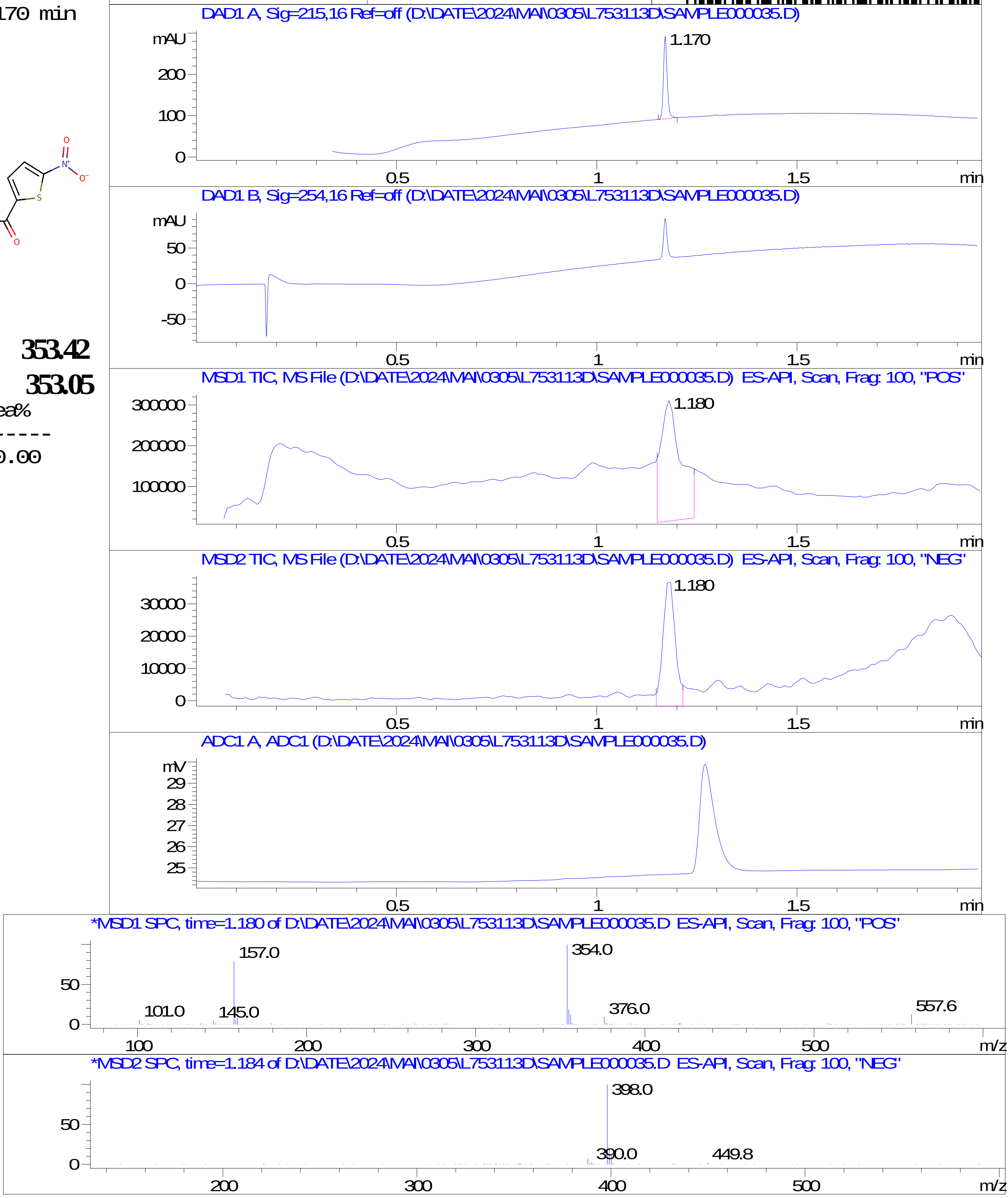

### Z8921960766.PDF

MaxPeak:  95.55%
Ret_Time: 1.039 min

# BD252975$2

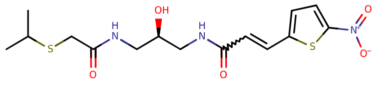

**Mol Wt** 387.47

**Exact Mass** 387.1

| # | Time | Area% |
|---|---|---|
| 1 | 0.994 | 4.45 |
| 2 | 1.039 | 95.55 |

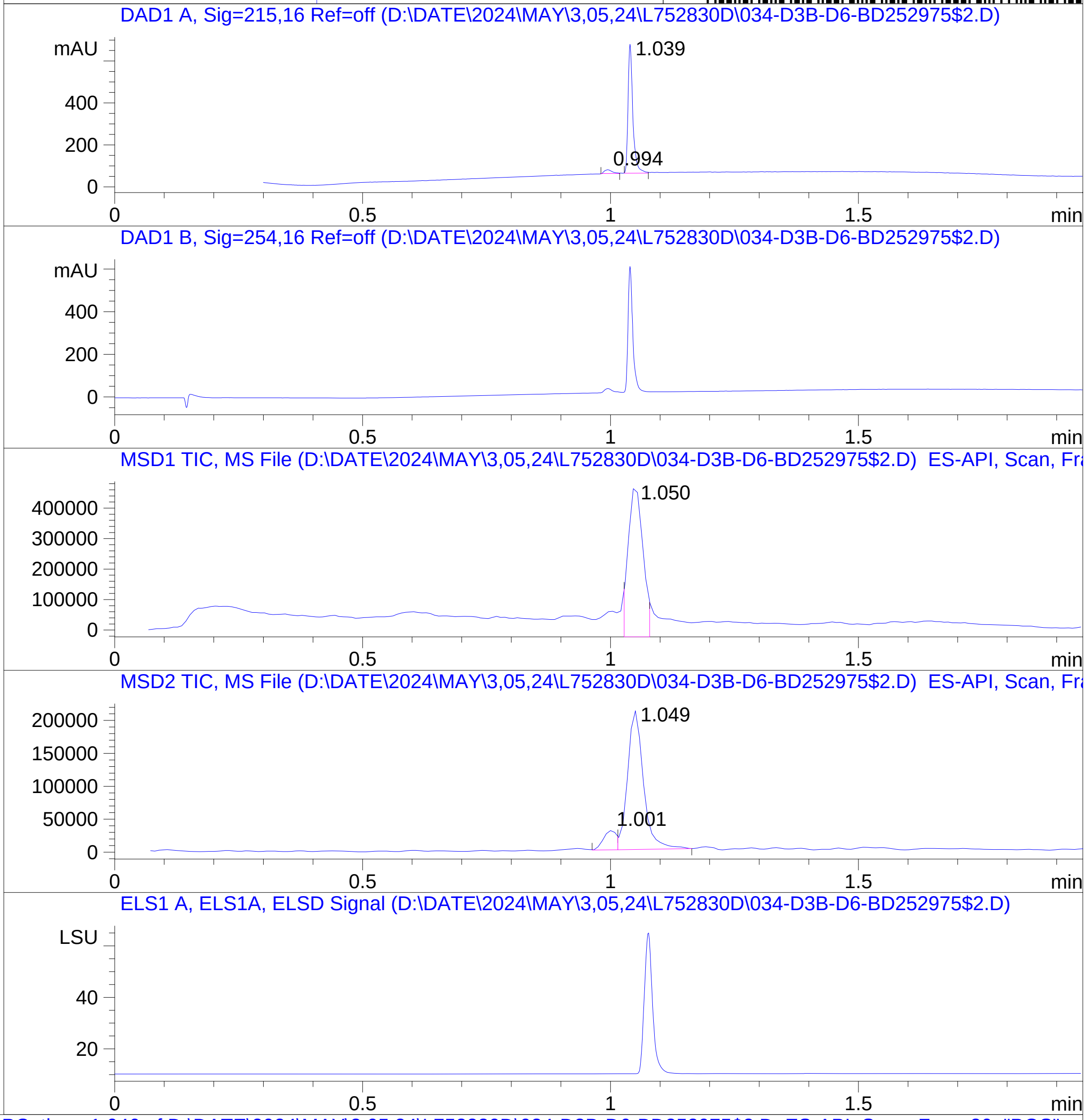


RT 1.050

RT 1.001

RT 1.049

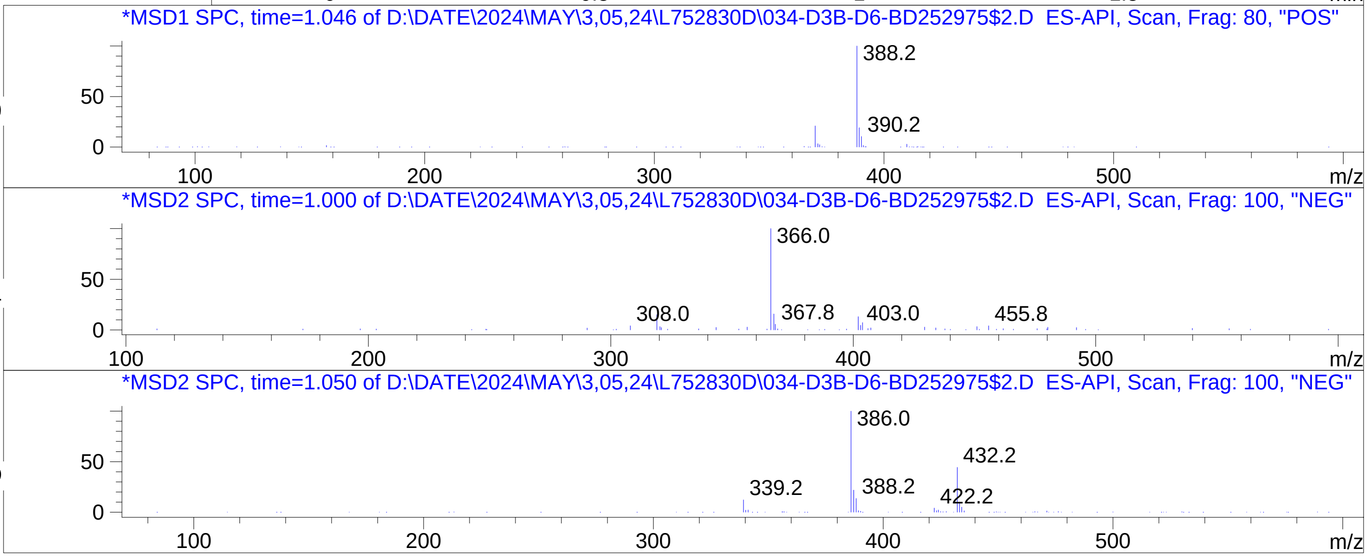


Inj.Date 5/2/2024 T -16- ->
C:\Chem32\1\Data\05_02\L752830D\SUPOR_30.M

### Z8921960806.PDF

MaxPeak: 100.00%
Ret_Time: 1.114 min

# BD252859$2

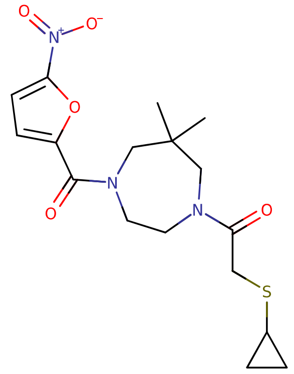


**Mol Wt** 381.45

**Exact Mass** 381.15

| # | Time | Area% |
|---|---|---|
| 1 | 1.114 | 100.00 |

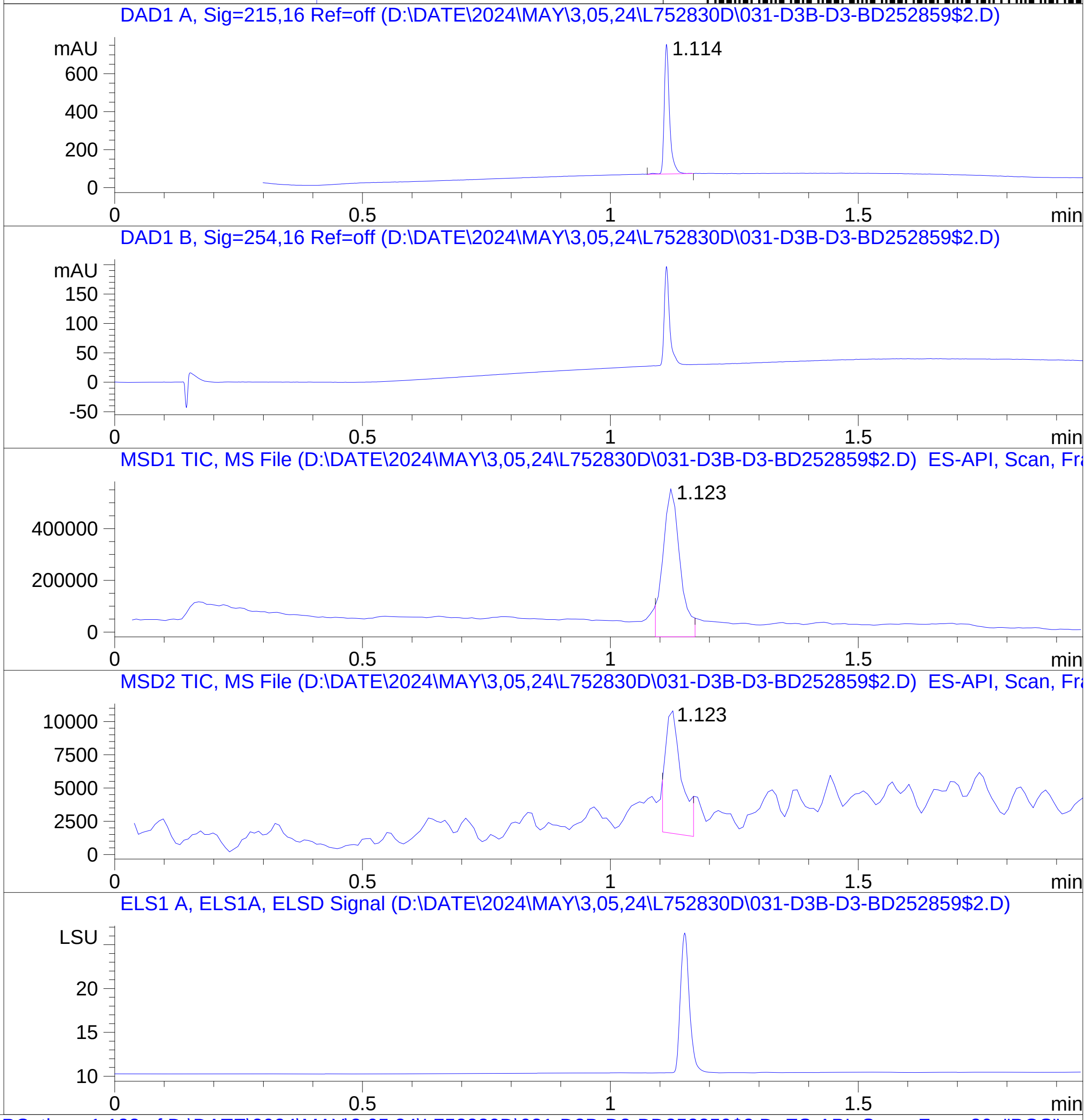


RT 1.123

RT 1.123

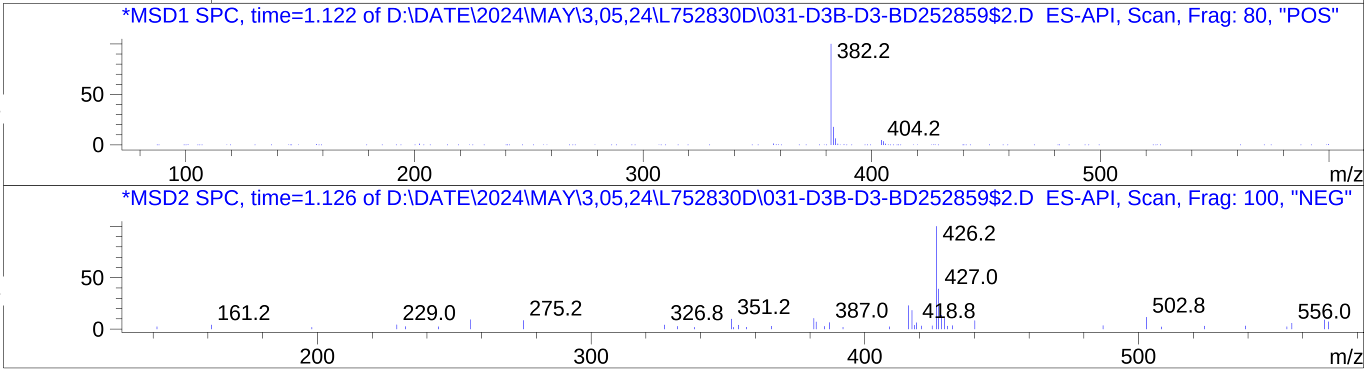

### Z8921960811.PDF

MaxPeak: 100.00%
Ret_Time: 0.945 min

# BD252799$1

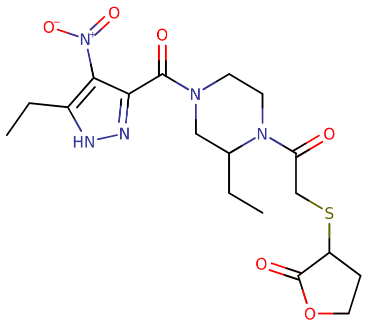

**Mol Wt** 439.49

**Exact Mass** 439.16

| # | Time | Area% |
|---|---|---|
| 1 | 0.945 | 100.00 |

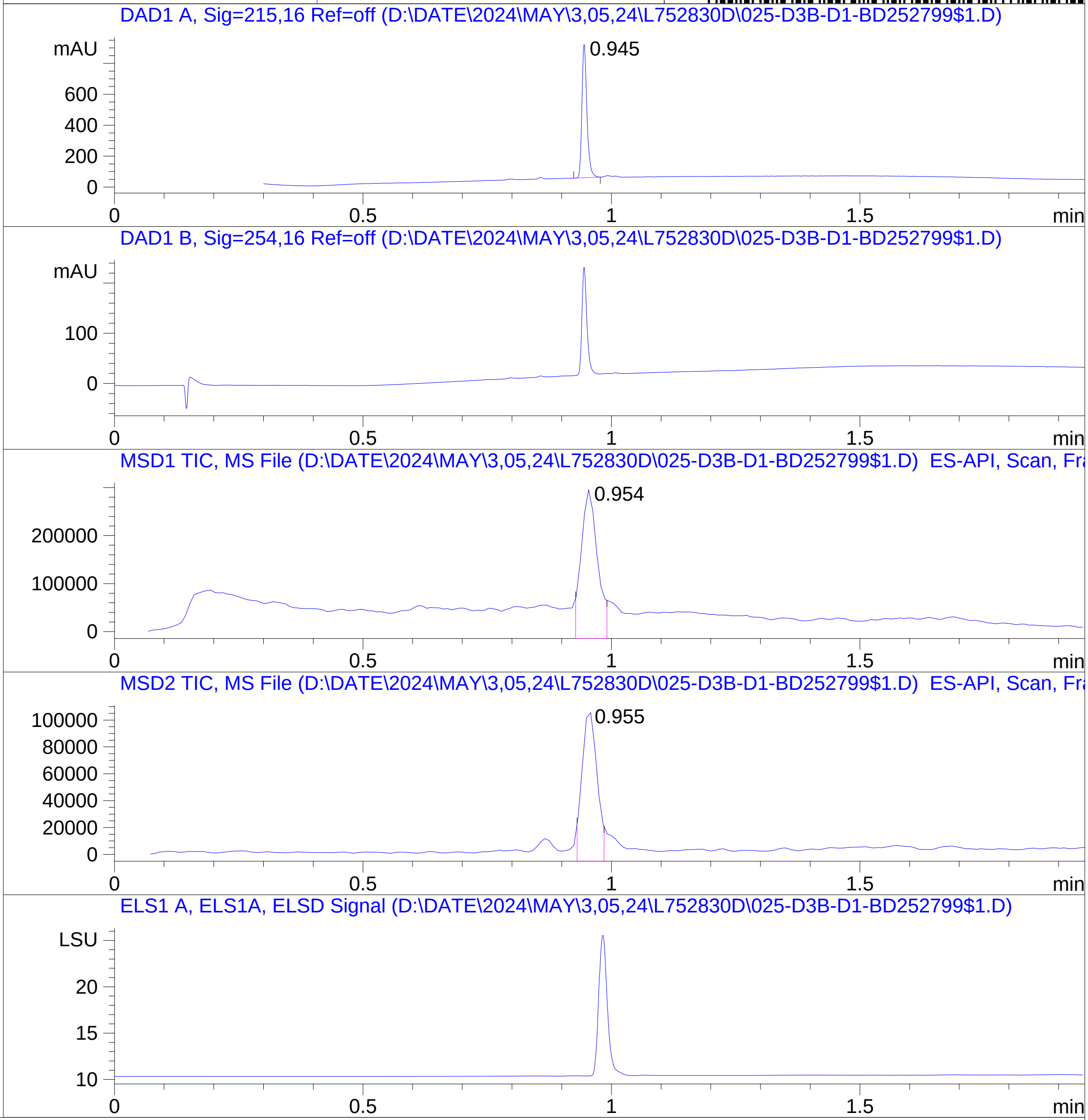


RT 0.954

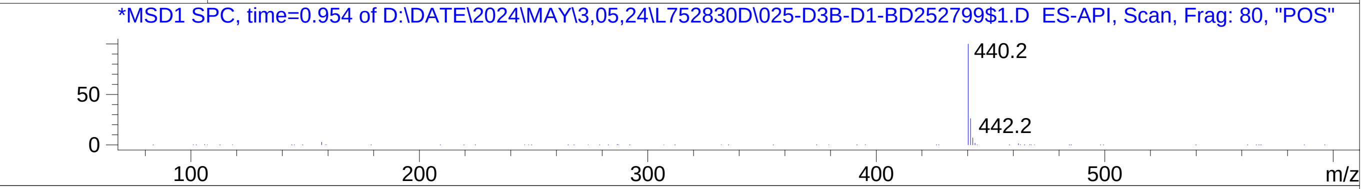


RT 0.955

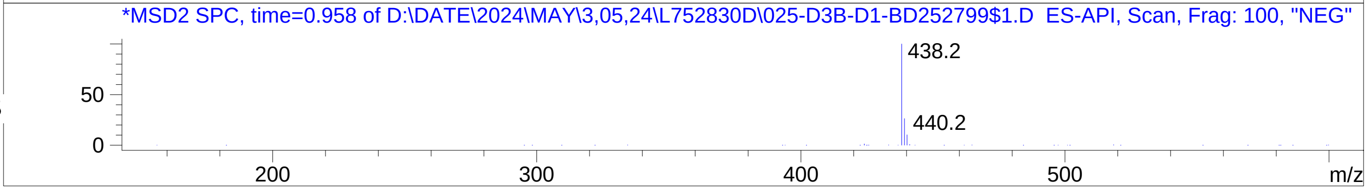


Inj.Date 5/2/2024 T -16- ->
C:\Chem32\1\Data\05_02\L752830D\SUPOR_30.M
