## Supplementary material for "SyntheMol-RL: a flexible reinforcement learning framework for designing novel and synthesizable antibiotics": Table S1

**Supplementary Table 1. Bacterial counts 24-hpi for Vehicle and Treatment skin samples.**

| Vehicle (10% DMSO) Bacterial Burden 24-hpi (CFU/g) | Treatment (2% Synthecin) Bacterial Burden 24-hpi (CFU/g) |
|---|---|
| 6214689266 | 46623794.21 |
| 7890743551 | 41522491.35 |
| 4656084656 | 35043804.76 |
| 5973715651 | 76131687.24 |
| 7822410148 | 69466882.07 |
